# A Symmetric Prior for the Regularisation of Elastic Deformations: Improved Anatomical Plausibility in Nonlinear Image Registration

**DOI:** 10.1101/664227

**Authors:** Frederik J Lange, John Ashburner, Stephen M Smith, Jesper L R Andersson

## Abstract

Nonlinear registration is critical to many aspects of Neuroimaging research. It facilitates averaging and comparisons across multiple subjects, as well as reporting of data in a common anatomical frame of reference. It is, however, a fundamentally ill-posed problem, with many possible solutions which minimise a given dissimilarity metric equally well. We present a novel regularisation method that aims to selectively drive solutions towards those which would be considered *anatomically plausible* by penalising unlikely lineal, areal and volumetric deformations. In addition, our penalty is symmetric in the sense that geometric expansions and contractions are penalised equally, which encourages inverse-consistency. We demonstrate that our method is able to significantly reduce volume and shape distortions compared to state-of-the-art elastic (FNIRT) and plastic (ANTs) registration frameworks. Crucially, this is achieved whilst matching or exceeding the registration quality of these methods, as measured by overlap scores of labelled cortical regions. Furthermore, extensive use of GPU parallelisation has allowed us to implement what is a highly computationally intensive optimisation strategy while maintaining reasonable run times of under half an hour.

## 1. Introduction

Nonlinear registration is commonly used in neuroimaging to deform images of individual brains into some common space (normalising both size and shape). Often the registration is based on structural, e.g., T_1_-weighted, images. The resulting warp is subsequently applied to both structural images as well as images depicting some function of interest such as BOLD or diffusion-derived connectivity. The rationale behind this is typically to facilitate statistical analysis of those functional data across subjects and populations.

An important distinction in this context is between volume- and surface-based methods. The former attempt to find the inter-subject mappings in the original (3D) space. The latter, in contrast, are typically only concerned with the cortex and attempt to inflate the folded 2D-manifold (the cortex) onto a sphere before finding the (2D) inter-subject mapping on that sphere (e.g., Fischl (2012); Robinson et al. (2014)). In this paper we present a novel volume based method.

Existing volume based methods can broadly be categorised based on:

### Construction of warps

As part of the iterative process a warp can be updated by adding an update, or by warping the previous warps by an update. The former is often referred to as an *elastic* or a *small deformation* framework, as opposed to a *plastic* or *large deformation* framework for the latter (Miller et al., 2003). Note that for infinitesimally small displacements, composition of warps can be extended to the concept of integrating a velocity field (Beg et al., 2005).

### Regularisation of warps

Warps can be *regularised*, i.e., have smoothness imposed on them, by adding a penalty term to the cost-function, or by explicitly smoothing the updates.

### Similarity measure

This is a scalar measure that assesses the similarity/difference between the images and whose maximisation/minimisation drives the registration.

The present paper is concerned mainly with the second of these, the regularisation, though we will touch upon the construction of warps and the effect of similarity metric choice as well. A through review of these categorisations can be found in Sotiras et al. (2013).

Historically, for the small deformation framework, one important aspect of the regularisation was to try to ensure that the warps stayed one-to-one and onto i.e., that each point in image A mapped to a unique point in image B, and that each point in image B could be reached from image A. This also means that the mapping is invertible. This was often attempted by adding a penalty term to the cost-function such that one simultaneously minimised the difference between the images (dissimilarity) and some function of the warps (regularisation). That function would often be borrowed from mechanics, such as bending energy (Bookstein, 1997), membrane energy (Amit et al., 1991) or linear-elastic energy (Miller et al., 1993) (whose formulations are summarised in Ashburner and Friston (1999)). However, none of those functions had any specifically biological relevance to the problem of mapping individual brains to each other. Moreover, they do not explicitly guarantee invertibility. So it was common practice to either ignore the issue of invertibility, or to empirically calibrate the relative weights of the image difference and the warp penalty so that the resulting warps were (almost) always invertible. But that typically meant a quite high weight for the penalty term which resulted in an inability to properly model large deformations.

This limitation led to the development of the large deformation framework where warps are updated by warping (resampling) the previous warp by an update warp (equivalent to a composition of multiple small warps). If both the previous and the update warps are invertible, this guarantees that the composed warp is also invertible. Therefore, as long as one ensures that each update warp is invertible the end result will be too. Furthermore, certain methods (such as LDDMM (Beg et al., 2005) and Geodesic Shooting (Ashburner and Friston, 2011)) seek solutions where the overall set of composed warps (or integrated velocity field) are as “smooth” (according to some criterion) as possible, rather than greedily smoothing at each step. The result is then that these methods are able to model large deformations while remaining invertible by taking many small update steps, leading to better matching than could be achieved within the small deformation framework.

However, invertibility is not the be-all and end-all for regularising warps. It does restrict the space of allowed warps, but some form of regularisation is still needed for choosing the optimal warp within that space. Moreover, it is no more clear in the large than in the small deformation framework how exactly that regularisation of the warps/velocity fields should be performed. Even when maintaining invertibility one can, and frequently does, still obtain warps that correspond to implausibly large local changes in volume and/or shape (Ashburner and Ridgway, 2013; Mang and Biros, 2016; Coalson et al., 2018). For a more detailed discussion of some of the tools which address invertibility, see Section 4.2.

In this paper we suggest using a small deformation framework together with a penalty function that explicitly penalises changes in both volume and shape. Furthermore, it approaches infinity as relative volumes approach zero (which would break invertibility), which means that it can be given a small weight in order to allow large deformations and still guarantee diffeomorphic warps. Hence, it addresses the same issue of invertibility as LDDMM based methods *and additionally* also the issue of how to find a particular set of warps within that space. It should be stressed at this point that the penalty we propose is not the same, or even particularly similar to, previously suggested penalties based on some function of the Jacobian determinant (see for example Rohlfing et al. (2003) or Leow et al. (2007)). We should also be clear that even though we refer to this as an elastic deformation, the prior is not based on any linear elastic model with the ensuing small deformation assumption. We will refer to our penalty as the Symmetric Prior for the Regularisation of Elastic Deformations (SPRED). The mathematical formulation and theoretical basis of the penalty are explained in Section 2.3. It has been used in the past (Ashburner et al., 1999, 2000), but the computational cost is such that at the time it was not practically feasible. With the advent of *general-purpose computing on graphics processing units* (GPGPUs) that is no longer the case, and we describe our implementation using NVIDIA’s CUDA framework (NVIDIA, 2019) of a registration algorithm based on that penalty function in Section2.4.

The rationale behind the suggested penalty function is twofold,

- To minimise geometric distortions, *i.e.* changes in both shape and volume, and to ensure the warps are one-to-one and onto.
- To be symmetric in the sense that a geometric (lineal, areal or volumetric) change by a factor of 2 is penalised the same as one by a factor of 0.5.

The first desideratum comes from the notion that brain tissue implements function, be that function integration of signal in the grey matter or transmission of signal in the white matter. The implementation of that function will necessarily occupy some physical space and it would seem unlikely that a given function can be implemented in a very small fraction of space in one brain compared to in another. A penalty should therefore favour no volume changes at all. It should also rapidly increase as the relative volumes become unrealistic, approaching infinity as the volume changes get close to breaking the one-to-one and onto criteria, thereby never allowing negative relative volumes. But it should also prevent unrealistic shape changes. Given what we know about the structure of both grey and white matter it is hard to picture a situation where a function implemented in a 1×1×1 mm^3^ cube in one brain might occupy a 0.01×10×10 mm^3^ sheet or a 0.1×0.1×100 mm^3^ stick in another. Note that this can all be reduced to penalising lineal changes since that would automatically also penalise shape and volume changes.

The second property, symmetry, comes from the desire that a registration algorithm should be inverse consistent, i.e., that the transform that is obtained from registering image A to image B is the inverse of that obtained from registering B to A (Christensen, 1999). A necessary, but not sufficient, condition for this is a prior that is symmetric in terms of penalising an expansion from A to B equally to the corresponding contraction from B to A.

## 2. Methods

### 2.1. Registration Framework

We now broadly describe the registration framework within which our SPRED penalty is implemented in order that specific details of the implementation are more readily understandable.

SPRED forms part of the development of our new MultiModal Registration Framework (MMORF) tool. MMORF is a volumetric registration tool whose underlying transformation model is a 3D cubic B-spline parametrised free-form deformation. The parameter space over which the optimisation is performed therefore represents the spline coefficients of three displacement fields (*x, y* and *z*-warps). A mean-squares (MSQ) similarity (or rather *dissimilarity*) metric is the data-consistency term optimised during registration. Regularisation penalties are calculated in the reference image space. A multi-smoothing, multi-resolution approach is employed to encourage convergence towards a globally optimal solution. From the outset MMORF has been designed to leverage GPU parallelisation, which is a largely why SPRED is computationally tractable within this framework.

The preferred optimisation strategy is the Levenberg variant of Gauss-Newton (Levenberg, 1944; Press et al., 2007). In this scheme, the Gauss-Newton Hessian **H**_*GN*_ is replaced with **H**_*L*_, where:

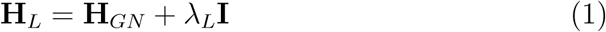

*λ*_*L*_ is used to ensure that the update step always leads to a decrease in the cost function. If an update would successfully reduce the cost function *λ*_*L*_ is decreased, otherwise it is increased and the update step re-evaluated. Additionally, the same strategy is used to ensure no update step is ever taken which would lead to non-diffeomorphic warps. This is achieved by checking that the resulting Jacobian determinants are all positive before accepting any update step, and modifying *λ*_*L*_ accordingly. Due to memory constraints, the Hessian can only be stored on the GPU until a warp resolution of 5 mm isotropic. Beyond this resolution our implementation offers a choice between two optimisation methods that do not require explicit representation of the Hessian. One is the Scaled Conjugate Gradient (Moller, 1993) method, which is related to other quasi-Newton methods and which uses that history of cost-function changes from earlier iterations to calculate a step-length along the next direction, thereby eliminating the need for line minimisations. The other is the Majorise-Minimisation (MM) method (Hunter and Lange, 2004) which replaces the Hessian with a diagonal matrix where the the *i*th value on the diagonal is the sum of absolute values of the elements in the *i*th column of the Gauss-Newton Hessian.

### 2.2. Intuition

In the following section we will detail how to calculate the suggested penalty, and why it is an ideal, albeit challenging, candidate for Single Instruction Multiple Data (SIMD) style parallelisation on a GPU (NVIDIA, 2019). For the estimation of the warp parameters we use the Gauss-Newton method, so we will also need to calculate the gradient of the penalty with respect to the spline coefficients of the warps and a Gauss-Newton approximation to the Hessian.

We attempt now to give an intuitive outline of the technical/mathematical formalism which follows in Section 2.3. At each voxel one can calculate a Jacobian matrix using the values of the surrounding warp spline-coefficients. The singular values of this matrix represent the orthogonal scalings of that voxel due to the warp. The SPRED penalty (described in Section 2.3.2) is a function of these singular values, and we can therefore calculate each voxel’s contribution to the total penalty. Those calculations are a perfect match for SIMD parallelisation with one computational thread per *voxel*, where each thread performs identical calculations but on different data. The individual voxel contributions are subsequently summed (a *reduction* in GPU parlance) to obtain the total penalty for the warp.

In order to calculate the gradient of the penalty, one needs to apply the chain rule, which says that:

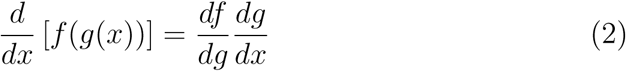

Remembering that for example *g* might be a vector-valued function of a vector-valued argument *x* which would make *df/dg* a vector and *dg/dx* a matrix. In our case the contribution to an element of the gradient from a given voxel is a scalar valued function of the vector of singular values of the local Jacobian matrix. These in turn are each a function of the vector of elements of the Jacobian matrix. Finally, these elements are functions of the vector of spline-coefficients whose support includes that voxel. The first two factors are again calculated using one thread per *voxel*, while the third factor uses one thread per *spline-coefficient*. Similarly each element of the total gradient is calculated by one thread per *spline-coefficient*.

To understand how the Hessian matrix is calculated consider a matrix **A** where each row corresponds to a voxel and each column to a warp spline-coefficient, and where **A**_*ij*_ is the rate of change of the penalty contribution from voxel *i* with respect to coefficient *j*. The Gauss-Newton approximation to the Hessian then becomes **A**^T^**A**. However, for practical reasons, **A** is never explicitly calculated or stored and instead the Hessian **H** is calculated directly. In this instance we use one thread per non-zero *element of* **H** for the final calculation, but there are a number of preceding steps which employ the same one thread per *voxel* approach as in the gradient calculations.

### 2.3. Theory

We now formalise the concepts introduced in Section 2.2 in terms of the mathematics required for incorporating the SPRED penalty into the optimisation strategy of our registration framework.

#### 2.3.1. Jacobian Matrix

Consider the generic problem of registering some 3D moving image *g* to a reference image *f*, where both are defined on ℝ^3^. Let *x, y* and *z* be the 3 orthogonal directions in ℝ^3^. We define a transformation 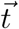 on the domain of *f* with parameters 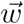, such that 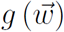 becomes our transformed moving image. At each point (*x, y, z*) :

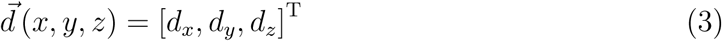

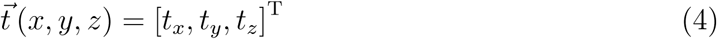

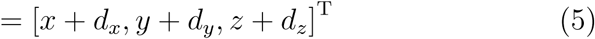

Where 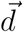 is a displacement field. If 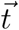 defines a continuous, differentiable function on ℝ^3^, then we may calculate a local Jacobian matrix **J** at any point in *f*, with:

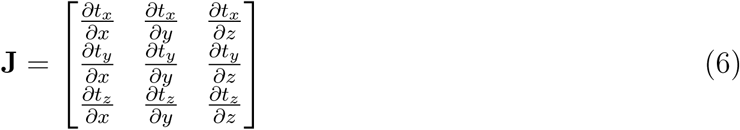

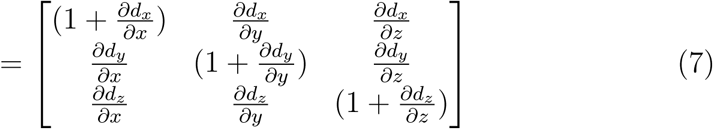

Where *∂d*_*i*_*/∂j* is the partial derivative of the *i* component of 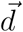 in the *j* direction.

**J** then describes how the image is locally scaled and rotated by the action of 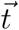. These actions can be decoupled by applying singular value decomposition (SVD) to **J**, such that:

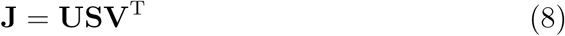

**U** and **V** are unitary matrices describing the rotational effect of **J**, and will not be considered further as they do not geometrically distort the image in any way. **S** is a diagonal matrix containing the singular values and represents the orthogonal scaling effect of **J**. It is the individual elements *s*_*i*_ of **S** which will be penalised in our regularisation, and we will refer to them as the Jacobian Singular Values (JSVs).

#### 2.3.2. Penalty Function

The penalty is based on the prior belief that the JSVs are drawn from a lognormal distribution. Such a penalty meets the biological plausibility arguments set out in Section 1 as:

- *s*_*i*_ = 1 is most likely
- *s*_*i*_ = 0 and *s*_*i*_ = ∞ are infinitely unlikely
- *s*_*i*_ = *a* and 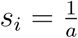 are equally likely

Now, we define *C* 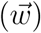 as the penalty (or *cost*) associated with with 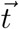:

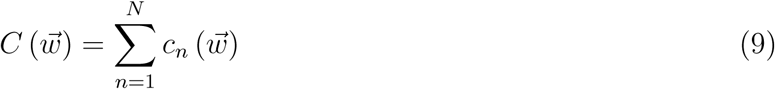

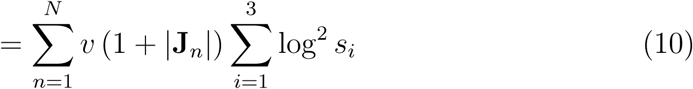

Where:

*C* = total cost/penalty

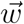 = transformation parameters

*N* = total voxels in image

*c*_*n*_ = cost/penalty at voxel *n*

*v* = voxel volume

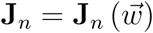 = Jacobian matrix at voxel *n*

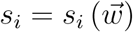 = *i*^th^ singular value of **J**_*n*_

From Equation 10 we recognise that the log^2^ *s*_*i*_ term represents our log-normal prior on the JSVs. However this prior is based on the distribution of singular values only in *f*. Therefore we have also included a term *v* (1 + |**J**_*n*_|) to ensure that our penalty is truly symmetric in *g* by accounting for the total volume in both images being penalised.

In practice, computation of the logarithm in Equation 9 (and its associated effect on the gradient and Hessian) may add a significant computational overhead (Ashburner et al., 2000). We therefore apply the following approximation:

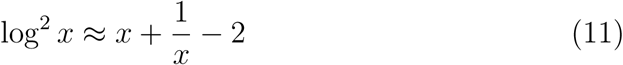

And, recognising that:

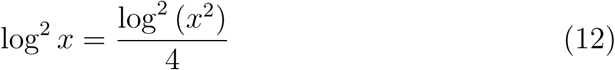

and that:

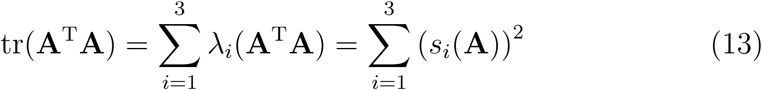

where *λ*_*i*_(**A**^T^**A**) denotes the *i*^th^ eigenvalue of **A**^T^**A** and where *s*_*i*_(**A**) denotes the *i*^th^ singular value of **A**. We may then write:

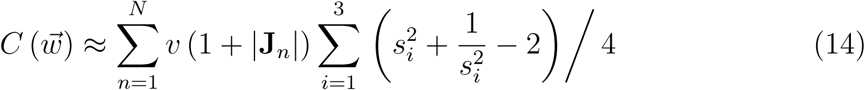

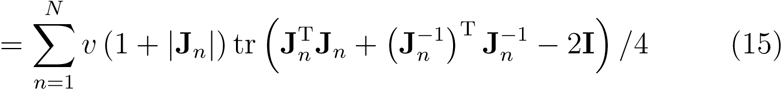

An important benefit of this approximation is that our cost function is now an explicit function of only the elements of **J**, and therefore we never need to calculate the actual SVD of **J**.

#### 2.3.3. Transformation Model

The previous sections have used a generic transformation 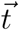, but at this point it becomes necessary to define the actual transformation model used. As in Equation 5, we utilise a displacement field 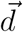 where each of the three directional components (*d*_*x*_, *d*_*y*_ and *d*_*z*_) in ℝ^3^ is composed of a basis set of uniformly spaced, 3D, cubic B-splines (**B**_*x*_, **B**_*y*_ and **B**_*z*_). Our transformation parameters 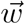 are then defined to be the coefficients of each of the B-splines in our basis set. If *M* B-splines are required to fully cover the *N* voxels in our reference image *f*, then we require 3*M* parameters to fully describe 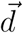 (i.e. *M* parameters for each displacement direction).

#### 2.3.4. Gradient and Hessian

The penalty function as defined in Section 2.3.2 will be included into a Gauss-Newton style optimisation framework, and therefore we require the gradient and the Gauss-Newton approximation to the Hessian of Equation 14. We note that for a valid application of Gauss-Newton (i.e., for being able to use the Gauss-Newton approximation to the Hessian) the function being minimised must be of the form 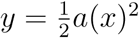, which Equation 15 is not (Chen, 2011). However, by making the substitution:

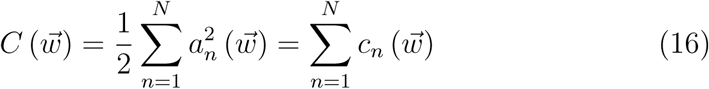

it becomes of that form. This means that:

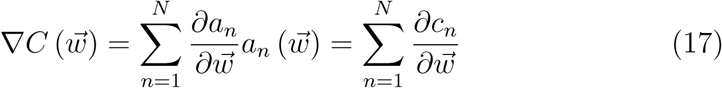

and therefore that the Gauss-Newton Hessian approximation becomes:

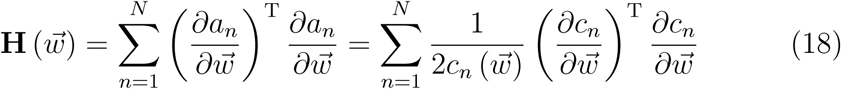

i.e., it introduces a factor 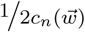 compared to how one would “normally” calculate the Gauss-Newton Hessian.

The gradient 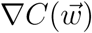 is a 3*M* × 1 column vector (where *M* is the number of B-splines used to represent one directional component of the warp-field), whose *m*^th^ element is of the form:

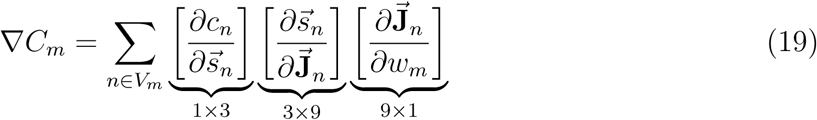

Where:

*V*_*m*_ = {voxels in *f* where B-spline *m* has support}

where *n* ∈ *V*_*m*_ means that the summation is over all voxels *n* for which the B-spline *m* has support, and where 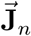 is a vectorised version of the Jacobian at the *n*th voxel and 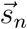 is a vector with the three singular values of that Jacobian.

Similarly the Gauss-Newton Hessian is a 3*M* × 3*M* matrix whose *jk*^th^ element is:

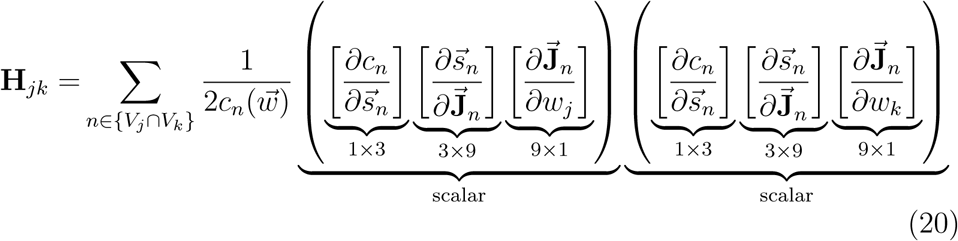

Where:

*V*_*j*_ ={voxels in *f* where B-spline *j* has support}

*V*_*k*_ ={voxels in *f* where B-spline *k* has support}

where *n* ∈ {*V*_*j*_ ∩ *V*_*k*_} denotes summation over all voxels *n* for which both B-spline *j* and *k* have support. It should be noted that although 3*M* × 3*M* can be a very large number, especially for high warp-resolutions/small knot-spacings, the vast majority of elements are zero because most spline combinations *jk* have no overlap in support.

We will demonstrate the practicalities of actually calculating these entities in the sections which follow.

### 2.4. Implementation

We now present how we achieved computational tractability of the SPRED penalty, its gradient, and GN Hessian.

#### 2.4.1. Parallelising Calculation of Gradient and Hessian

As with all but the simplest optimisation strategies, calculating the value of the SPRED penalty itself is not the computationally costly part of this algorithm. Rather, it is the corresponding gradient and Hessian calculations which are massively computationally complex. See, for example, Equation 19, where we have multiple applications of the chain rule to vector valued functions of vectors, leading to intermediate matrices. These intermediates then need to be multiplied together, further increasing the complexity. It is easy to see that this complexity is compounded even further in Equation 20 when calculating the Hessian.

If these calculations were all performed sequentially the problem would become computationally intractable, and the suggested form of regularisation would be a mathematical nicety with no practical impact.

However we will demonstrate how the problem can be framed in terms of the application of multiple massively parallelised SIMD operations, rendering it realistic to use.

As an example, let us consider how one might parallelise the gradient calculation in Equation 19. Firstly we note that the part:

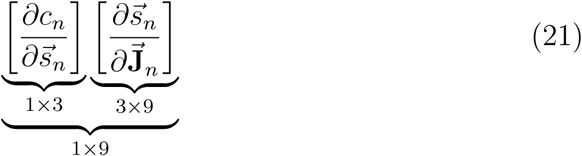

depends only on entities dependent on which voxel *n* is currently being considered. Each of these 1 × 9 vectors can therefore be calculated and stored independently by one GPU thread. We represent these pre-calculated vectors as an *N* × 9 matrix (where *N* is the total number of voxels in image **f**) which we denote [*∂c/∂***J**] where the *n*th row is:

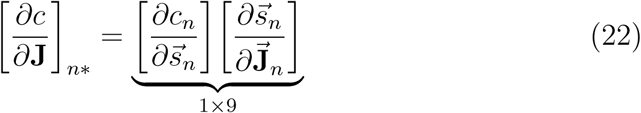

The columns of [*∂c/∂***J**] can be thought of as *partial derivative images* and can be visualised as demonstrated in Appendix B. Note also that in these calculations one can use the approximation given in equation 15 to avoid explicit computation of the JSVs.

The next part:

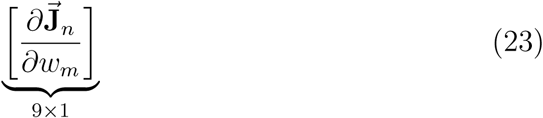

is “constant” in the sense that its elements are given by the spatial derivatives of the B-spline basis-functions and can be calculated (analytically) once and for all for a given knot-spacing. If we denote a column-vectorised version of a single 3D B-spline basis-function (whose dimensions are given by the knot-spacing) by **B** we can create a matrix [*∂***J***/∂w*] containing all the elements we need as:

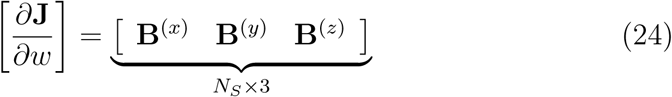

where **B**^(*)^ denotes the spline basis-function spatially differentiated in the *i*th direction and *N*_*S*_ denotes the total size of the 3D spline (determined by the knot-spacing).

Finally each element ∇*C*_*m*_ of the gradient can be calculated by independent threads as:

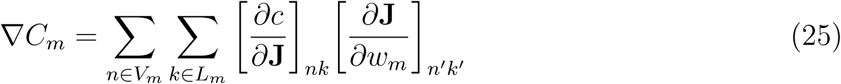

Where:

*V*_*m*_ = {voxels in *f* where B-spline *m* has support}

*L*_*m*_ = {elements of **J**_*n*_ where B-spline *m* has an effect}

*k* ∈ *L*_*m*_ denotes that *k* will select the three columns of [*∂c/∂***J**] that contribute to the *m*^th^ element of the gradient. To be concrete, if 1 ≤ *m* ≤ *M* the element pertains to the *x*-component of the warp-field and *k* will select the columns of [*∂c/∂***J**] corresponding to the first row of **J** (see equation 6), and if *M* < *m* ≤ 2*M* it will select the columns of [*∂c/∂***J**] corresponding to the second row etc. The ′ superscripts on *n*′ and *k*′ indicate transformed versions of *n* and *k* such that *n*′ takes into account the offset of the *m*^th^ spline into the image, and *k*′ will select the corresponding column (regardless of row) in **J**.

Equation 25 can also serve as an illustration of some of the challenges of SIMD parallelisation. We have chosen a strategy where each element of ∇*C* is calculated by one thread. That choice means that we avoid having to synchronise writes to ∇*C* or having to perform reductions, both of which reduce occupancy and speed. The downside of that choice is that the different threads within a warp will access global memory in a non-coalesced fashion. This is because each thread will access a 3D sub-volume in each of the nine “partial derivative images” defined in equation 22, where these sub-volumes are offset by an integer multiple of the knot-spacing in one or more dimensions. In the supplementary material there are timings and more detailed examples of the challenges of parallelising the proposed regulariser.

So, in summary the calculation of the gradient can be divided into three parts:

- One which is separable over voxels (or more generally “sample points”) where the calculations for one voxel can be performed and stored independently by one GPU thread.
- One which is “constant” and can be performed once and for all and stored where the results can be accessed by all threads.
- One which is separable over warp parameters (spline coefficients) where calculations for one warp parameter can be performed and stored independently by one GPU thread.

Following this line of reasoning, we may similarly divide the Hessian calculation of Equation 20 into portions which are constant, separable over samples, and separable over warp parameters.

As it is this process which is central to how using the SPRED penalty is made tractable, we provide an intuitive understanding of what implementing the parallelisation of equation 19 actually looks like in practice by considering a simple 2D example. This can be found in Appendix B.

### 2.5. Testing

As assessing the performance of a registration tool is non-trivial, we here try to strike a balance between considering some measure of data-consistency, and some measure of anatomical plausibility. It is known that image similarity is a poor choice for measuring registration accuracy as it is just a proxy for what we are truly interested in, i.e., alignment of common anatomical structures (Rohlfing, 2012). Image-wide tissue maps are not a good option either, as they are global in nature and only really test tissue classification rather than anatomical alignment (Rohlfing, 2012). We therefore choose the overlap of manually segmented cortical regions as our data-consistency measure, which is an established method by which to assess the quality of registration (Klein et al., 2009).

The focus of this work is not primarily on assessing our method’s ability to maximise data consistency. However, if a set of warps was unable to produce high quality overlap results then it would not be of interest irrespective of how plausible the deformation is. High registration accuracy therefore simply allows us to evaluate the anatomical plausibility of the warps in a meaningful way.

For any method, regardless of the specifics of the regulariser, there is a trade-off between the image similarity term and regularisation term. That trade-off is determined by their relative weights, something that is typically calibrated empirically (however see Simpson et al. (2012) for an attempt to determine it probabilistically). Therefore, in order to compare two methods in terms of plausible warps it is crucial to calibrate both methods so that they achieve (close to) identical overlap scores.

We ensured a high registration accuracy by using ANTs (Avants et al. (2008)) with similar settings to those used in Klein et al. (2009) as a yardstick for registration accuracy. To make sure warp comparisons were meaningful we calibrated the other methods to yield very similar overlap scores. If we could not achieve that for a method we did not include that in the comparison of warps.

We use two main indices for assessing what we consider to be anatomical plausibility. One is the Jacobian determinant that quantifies local volume change. We consider both the histogram of Jacobian-determinants within the brain, which provides information about how aggressively the warp is squeezing or expanding the volume on average, and how the Jacobian-determinants are spatially distributed, i.e., whether the deformation appears to be spatially sensible.

The second index is the cube-volume aspect ratio (CVAR, Smith and Wormald (1998)) which quantifies shape changes. In three dimensions it is the cube root of the ratio of the volume of the smallest possible enclosing cube to the actual volume. Similarly to the Jacobian determinant above we consider both the histograms and the spatial distributions of the CVAR.

#### 2.5.1. Registration Tools

In order to make a meaningful and relevant evaluation of the effect of the SPRED penalty on registration performance, four volumetric registration tools were included in this comparison, namely:

**FLIRT (Jenkinson and Smith, 2001)** We chose to include a linear registration method in order to provide a yardstick by which to compare the overlap scores, and in particular the differences in overlap, of the other methods.

**ANTs (Avants et al., 2008, 2014)** ANTs has been shown to perform well in a direct comparison with other methods (Klein et al. (2009), Ou et al. (2014)). A method performing significantly worse would not be of any practical interest, so it provides a minimum accuracy bar for the other methods in the comparison. We ran ANTs both with cross correlation (Avants et al. (2011)) and mean sum of squares image similarity metrics. In both cases the greedy SyN transformation model was used.

**FNIRT (Andersson et al., 2007)** We included FNIRT because it uses the same image similarity and warp representation as MMORF, but employs a different strategy to ensure invertibility (Karaçali and Davatzikos (2004)) and a different regulariser (bending energy) compared to MMORF.

**MMORF** Implements the regulariser (SPRED) that we propose in the present paper. In order to demonstrate just how different to a Jacobian Determinant regulariser (for example Rohlfing et al. (2003) or Leow et al. (2007)) SPRED is, we also implemented a regulariser based on the sum of squares of log-determinants of the Jacobians in the exact same framework.

Whilst there are myriad other tools that may also have been included in this comparison (e.g., SPM Unified Segmentation (Ashburner and Friston, 2005), DARTEL (Ashburner, 2007), elastix (Klein et al., 2010), MIRTK (Rueckert et al., 1999; Schnabel et al., 2001), Diffeomorphic Demons (Vercauteren et al., 2009), ART (Ardekani et al., 2005)), the chosen methods represent a broad enough overview of the types of tools available for the purposes of characterising the performance of MMORF.

#### 2.5.2. Registration Parameter Selection

An important aspect when comparing registration algorithms is ensuring that the user-selectable parameters are as optimal as possible (Klein et al., 2009).

FNIRT was run with an optimised strategy proposed by the tool’s author. This consisted of a multi-resolution, multi-smoothing level registration, to a final knot spacing of 1 mm isotropic, and is explained in detail in Andersson et al. (2019).

ANTs-CC was run using the antsRegistrationSyn.sh script, but with the gradient step and smoothing values changed to match those supplied by the tool’s author for use in previously published comparative studies (Klein et al., 2009). ANTs-MSQ was run with hand tuned parameters. Both versions of ANTs employed a multi-resolution, multi-smoothing level approach, to a final resolution of 1 mm isotropic.

MMORF was run using parameters that were shown to perform well during the development of the tool. Again, a multi-resolution, multi-smoothing level approach was used, but with a final knot spacing of 1.25 mm.

Note that whilst significant effort has been made to ensure that each tool performs as well as possible, the overarching goal was not to definitively classify which tool performed best in terms of overlap scores. But rather to investigate how the SPRED penalty affects the nature of the deformations when compared to similar performing tools. Hence, the main goal was to calibrate their respective performances in terms of overlap accuracy so that they were not significantly different. Further details regarding parameter selection can be found in Appendix C.

#### 2.5.3. Test Dataset

We have chosen to use the Non-rigid Image Registration Evaluation Project (NIREP) dataset (Christensen et al., 2006) in testing the registration methods. This dataset consists of the T_1_-weighted scans of 16 healthy subjects, eight male (average age 32.5 years) and eight female (average age 29.8 years). Each subject’s scan has been brain extracted, bias corrected, and 32 cortical regions (16 left hemisphere, 16 right) have been expertly hand-segmented. The segmented regions provide a *ground truth* for anatomical correspondence of a finer granularity than simple tissue maps.

#### 2.5.4. Evaluation Strategy

The registration tools were evaluated using pairwise registrations of each combination of the 16 subjects (240 warps in total).

Each of the 32 segmented cortical regions were transformed through these warps and various overlap metrics calculated, namely: Jaccard Coefficient, Dice Coefficient, Specificity, Sensitivity. The distributions of these overlaps in each region were then compared between registration methods.

Additionally, distributions of Jacobian determinants and of cube-volume aspect ratios (CVAR, see Appendix D for a definition) were calculated in order to gain some insight into how aggressive each method is in terms of volume and shape changes. These were inspected and compared as histograms, spatial maps and by statistics such as min, max, mean, 5th and 95th percentiles. The comparisons were performed only for the methods for which we achieved overlap scores that equalled those of ANTs-CC.

## 3. Results

### 3.1. Overlap Scores

Combined left/right hemisphere overlap scores are shown in Figure 1. All overlap measures showed similar trends, and therefore we focus only on the Jaccard Coefficient as our measure of choice. FLIRT, as expected, achieves the lowest levels of overlap, with mean overlaps varying depending on the region being considered. The “gap” in overlap between FLIRT and the non-linear methods also provides a useful yardstick against which to gauge the differences between those. All of the nonlinear methods, again as expected, improve the degree of overlap considerably. ANTs-MSQ has the lowest average overlap of the nonlinear methods. FNIRT has the second lowest overlap scores, although in certain areas it is outperformed by ANTs-MSQ. Note that the FNIRT results are significantly better than in previously reported comparisons (e.g. Ou et al. (2014) who used the parameters from Klein et al. (2009), which were unsuited for skull stripped data), but are in line with those from the tool’s author (Andersson et al., 2019). ANTs-CC and MMORF perform the best and equally well on average, but one or the other performs better in individual areas. These observations are supported when one considers the overall distributions of overlap scores in Figure 2 which display the same trends.

**Figure 1:**
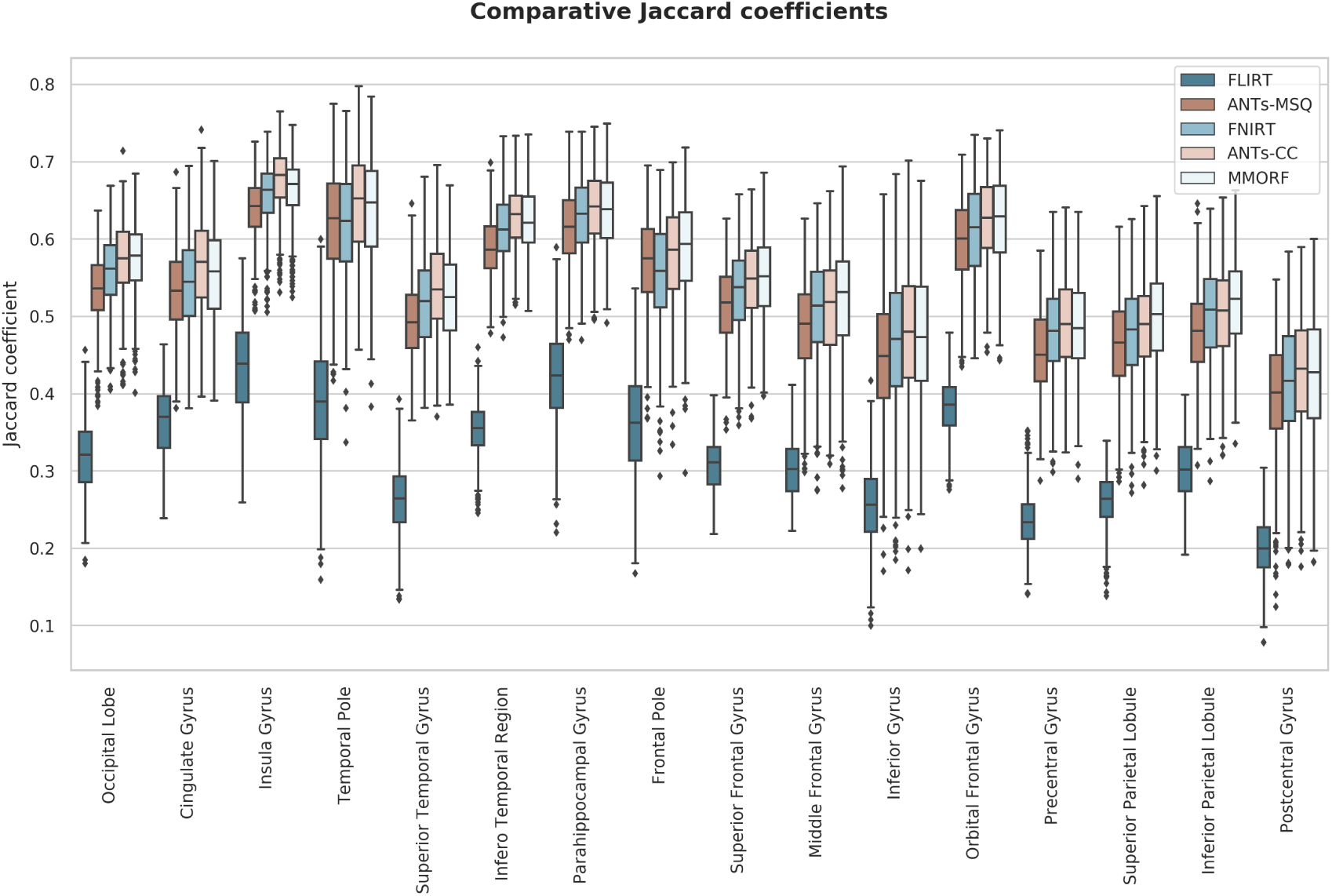
Jaccard coefficients of 16 regions (32 original cortical segmentations combined left/right) for each of the 4 nonlinear and 1 linear registration methods. The FLIRT result provides a baseline for evaluating the improvement in overlap between linear and nonlinear methods. Overall ANTs-CC and MMORF perform similarly. In some areas ANTs-CC produces better results (e.g., cingulate gyrus), and in some area MMORF performs better (e.g., inferior parietal lobule). In all cases, both methods outperformed ANTs-MSQ and, to a lesser extent, FNIRT.

**Figure 2:**
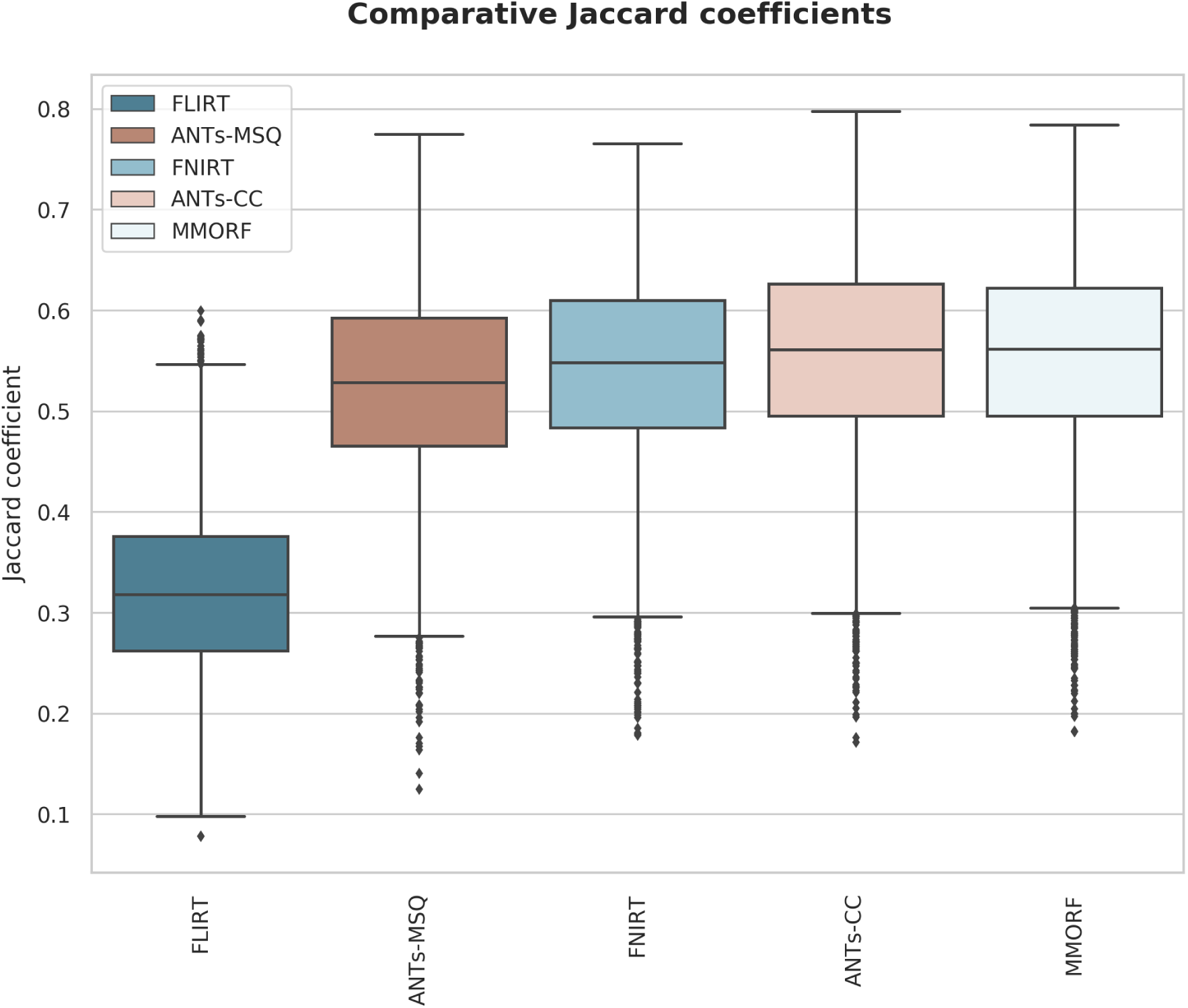
Distribution of Jaccard coefficients combined across all regions and subject pairs for each of the 4 nonlinear and 1 linear registration methods. This confirms the trends observed in Figure 1, with MMORF and ANTs-CC performing best, followed by FNIRT and then ANTs-MSQ. The MMORF and ANTs-CC distributions are remarkably similar, despite the slight region-to-region differences in performance.

It is perhaps surprising that the two versions of ANTs produce such different results, given that they differ only in their choice of similarity metric. However, this is in line with what the creators of ANTs report themselves (Avants et al. (2011)) on the LPBA40 data set, so we believe this to be an accurate representation of their respective performances.

Because the best overlap performance we could achieve with ANTs-MSQ was significantly worse than that of MMORF and ANTs-CC we did not consider it meaningful to include it further in the comparison of warp metrics (Jacobian determinant and CVAR).

We did include FNIRT in one of the warp-comparisons, even though it did perform slightly worse than ANTs-CC and MMORF. That was because the difference was considerably less than for ANTs-MSQ, and also because it used the same image similarity metric and warp-construction as MMORF, differing only with respect to strategy for ensuring invertibility and regularisation.

Our main interest was in the comparisons between ANTs-CC, being a very accurate and widely used method, and MMORF. The fact that we managed to calibrate the two methods so that they have almost identical overlap scores means that the warp distributions can be compared in a fair and meaningful way.

The joint distribution of overlap scores for MMORF and ANTs-CC for all regions and warps shown in Figure 3 confirms that the two methods are on average very comparable.

**Figure 3:**
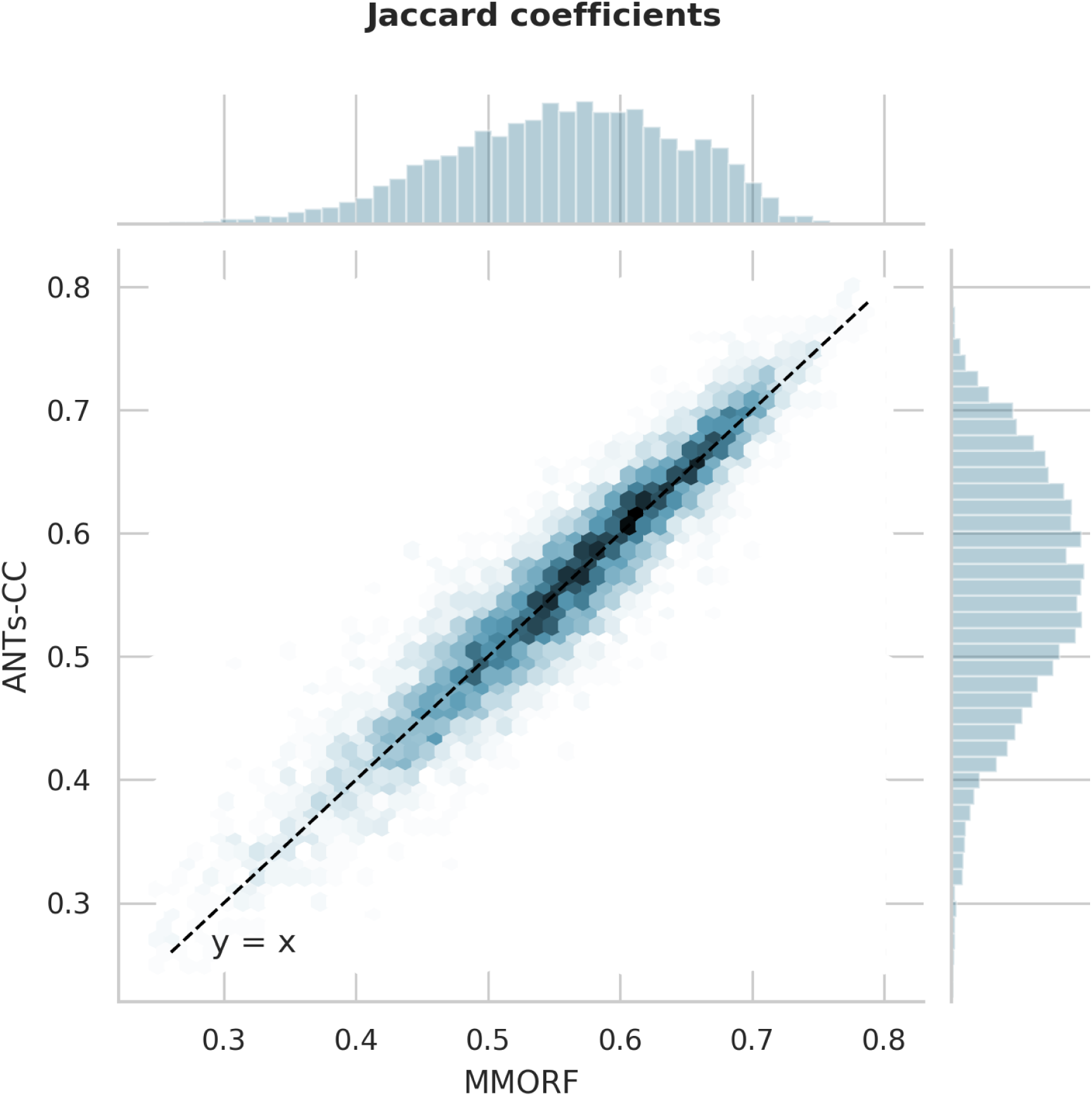
Joint histogram of Jaccard coefficients across all regions and warps for MMORF and ANTs-CC. Note the tight, linear relationship, supporting the observation from Figure 1 that MMORF and ANTs-CC have very comparable overall performance across the NIREP dataset.

### 3.2. Jacobian Determinant Distributions

Figure 4 shows the Jacobian determinant distribution of the warps generated with MMORF, ANTs-CC and FNIRT for a randomly selected image pair. We show both the non-log and log distributions since they highlight slightly different behaviours. It should be noted that although these warps represent a single pair of subjects, we could have chosen any of the 240 warps and the relationship would remain essentially the same.

**Figure 4:**
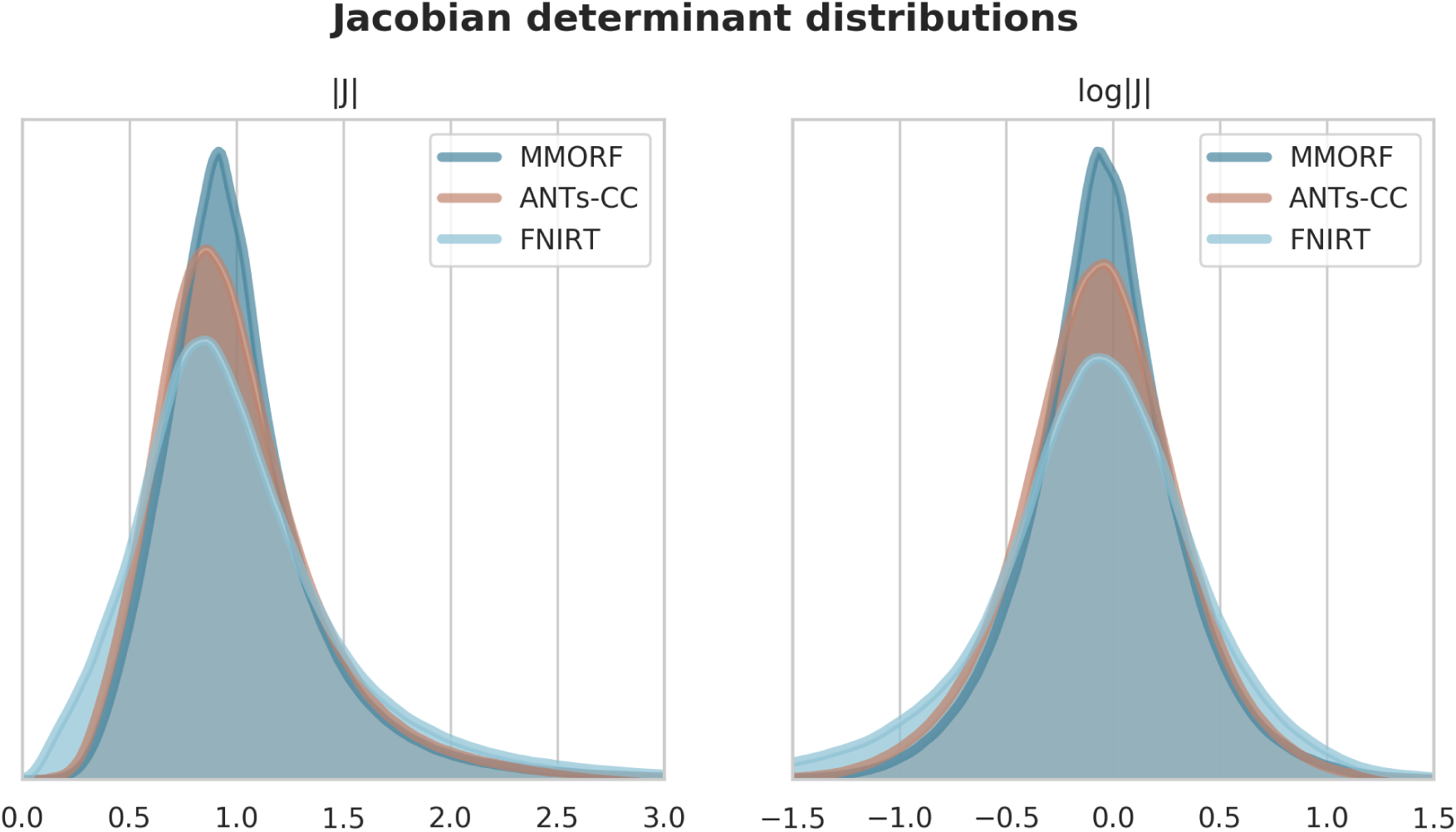
Jacobian determinant and log-Jacobian determinant distribution of a randomly selected registration pair from the NIREP dataset. Only values within the brain itself have been considered. As desired, all four methods produce Jacobian determinants with few to no negative values. There are distinct differences in appearance though. FNIRT is the most different, with a visibly heavier negative log-Jacobian tail. This is an effect of projecting the Bending-Energy regularised registration onto a field without negative Jacobian determinants. ANTs-CC displays the next widest log-Jacobian distribution, but with more evenly weighted tails.

The Jacobian determinant distribution for FNIRT has a shape that is quite distinct from that of MMORF and ANTs-CC. It is seen very clearly in the non-log distribution where instead of tapering off smoothly towards zero there is an almost linear drop. This is most likely due to FNIRT projecting its warps onto the nearest B-Spline field which has no negative Jacobian determinants (Karaçali and Davatzikos (2004)). The shapes of the distributions for MMORF and ANTs-CC are much more similar, but with ANTs-CC having a notably greater dispersion.

Of the three methods that had the most similar overlap scores it is clear that FNIRT causes substantially greater volume distortions, in addition to having slightly lower overlap scores. We therefore drop FNIRT from further analysis at this stage and focus on MMORF and ANTs-CC.

From the log distribution, the range of the 5^th^ to 95^th^ percentile for each method can be calculated. This was done for every warp, and the results are summarised in Figure 5. It can be seen that both the mean and the dispersion of the ranges are considerably greater for ANTs-CC than for MMORF. Figure 6 plots for each of the 240 registrations the log 5-95 range for ANTs-CC against that of MMORF. In all but 4 cases the percentile range is smaller for MMORF.

**Figure 5:**
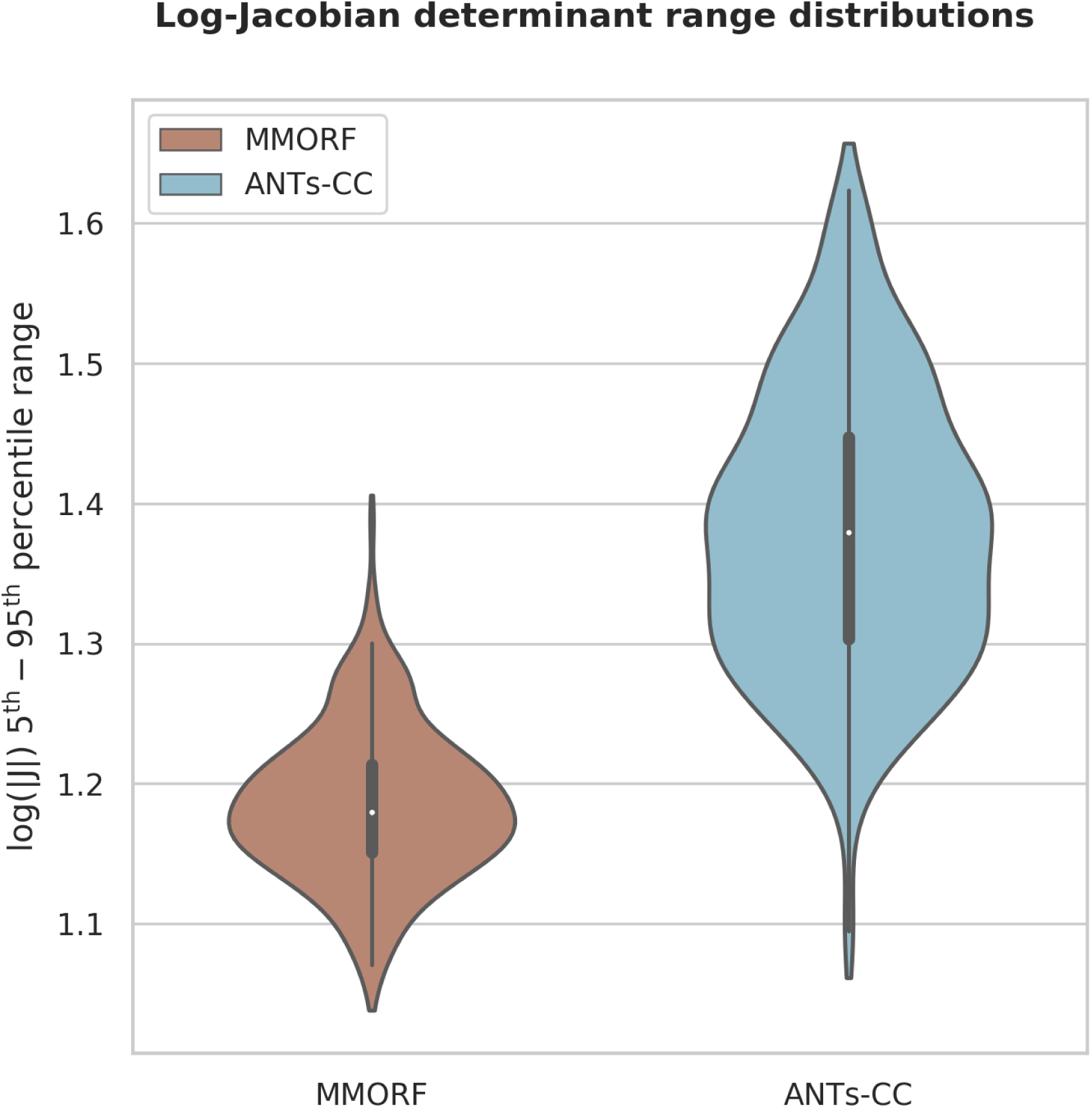
The 5^th^ to 95^th^ percentile is used as a summary measure for the log-Jacobian determinant distributions in Figure 4. The distribution of this percentile range over all 240 registrations is shown for the two best performing methods. ANTs-CC displays a significantly higher range on average, as well as a greater dispersion of ranges when compared to MMORF.

**Figure 6:**
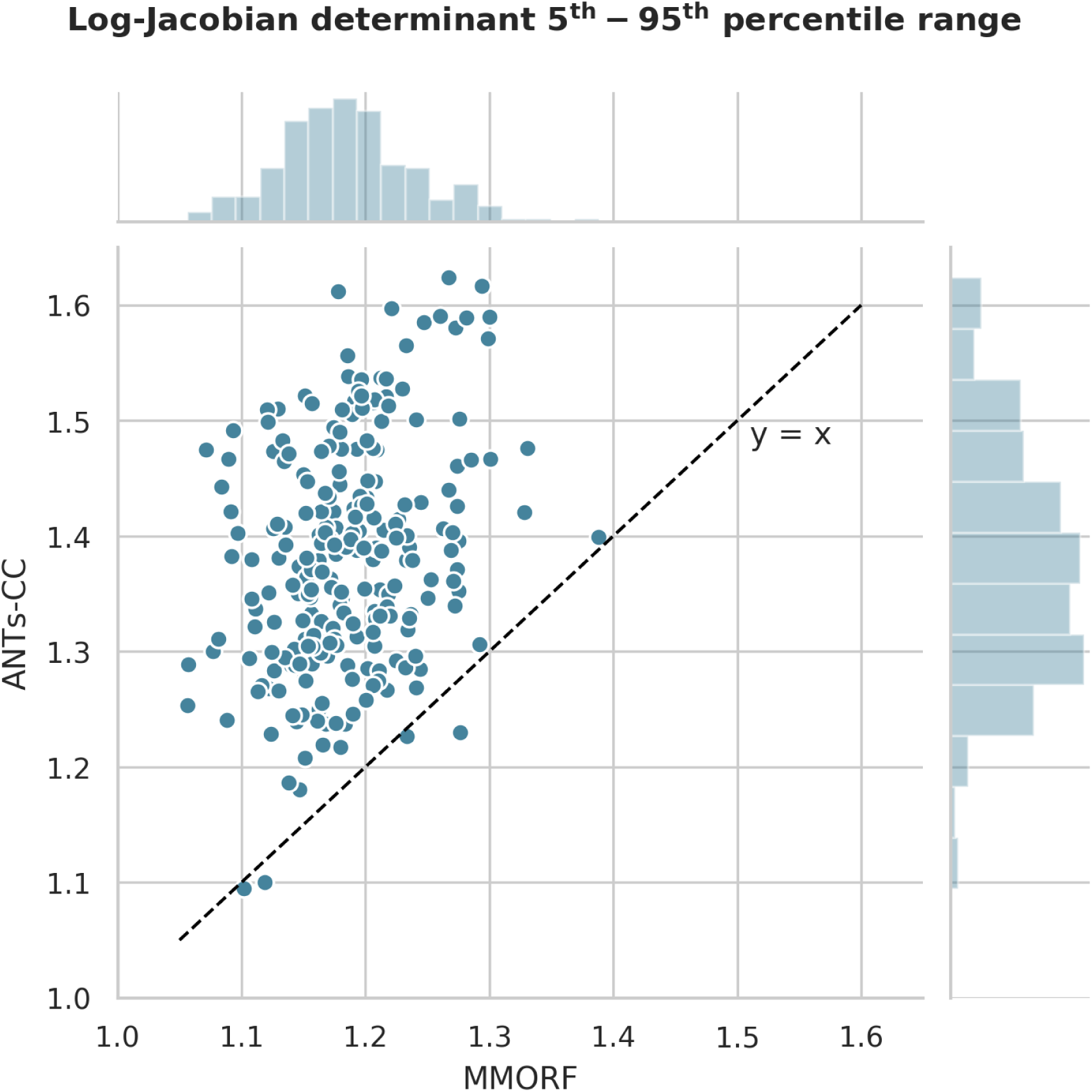
Scatter plot of MMORF and ANTs-CC log-Jacobian determinant 5^th^ to 95^th^ percentile ranges. The key feature here is that ANTs-CC produces systematically wider ranges compared to MMORF, with only 4 of the 240 registrations favouring ANTs-CC. This is despite ANTs-CC and MMORF producing comparable Jaccard coefficients overall, as shown in Figure 3.

### 3.3. CVAR Distributions

Unlike the Jacobian determinant, the mean (across sample points within the brain) of the CVAR is a meaningful statistic. We therefore use that as our summary measure of shape distortions for a given warp. Figure 7 demonstrates that the mean is both lower on average, and has smaller dispersion for MMORF compared to ANTs-CC.

**Figure 7:**
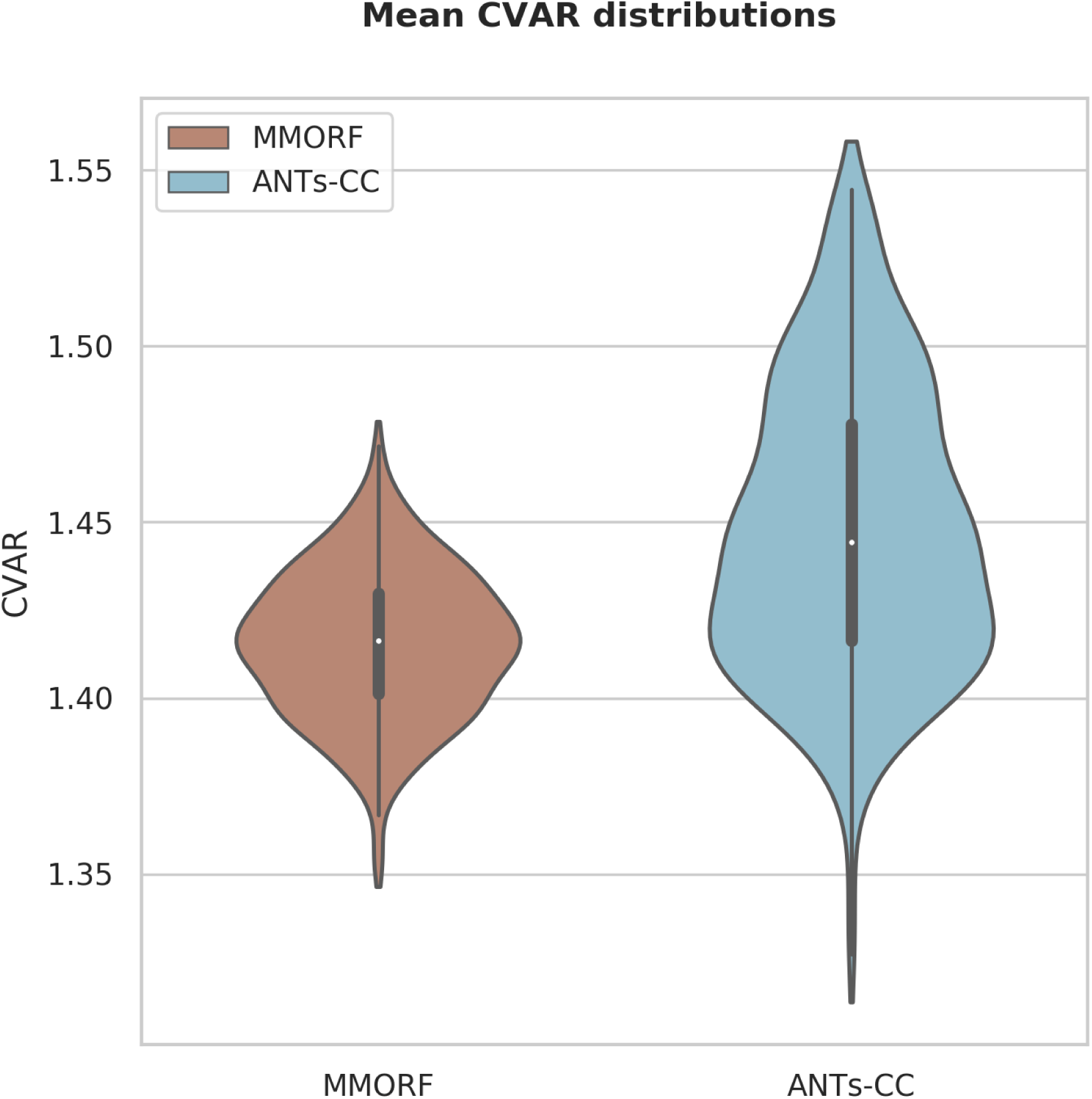
Distribution of mean CVAR within the brain over all 240 registrations is shown for the two best performing methods. ANTs-CC displays a significantly higher mean CVAR on average, as well as greater variance when compared to MMORF.

### 3.4. Spatial Maps

In the previous section we showed that of the methods with comparable overlap scores ANTs-CC caused substantially more volume distortions than MMORF. In order to better understand the source of that difference we look at the spatial maps of the Jacobian determinants and of the CVAR. Figure 8 shows maps of Jacobian determinants and figure 9 of CVAR for the same randomly selected pair of subjects as Figure 4. There are clear visual differences between the results of the two methods. In areas where the T_1_-weighted signal intensity has strong contrast and carries information about the tissue type, such as along the cortex, both methods produce similar looking maps with varying amounts of expansion and compression. Where they differ in appearance is predominantly within areas displaying a relatively flat T_1_-weighted signal profile, such as in white matter. Here MMORF produces smoothly changing values with magnitudes close to 1. ANTs-CC by contrast displays highly oscillatory behaviour, with a higher proportion of values deviating away from 1.

**Figure 8:**
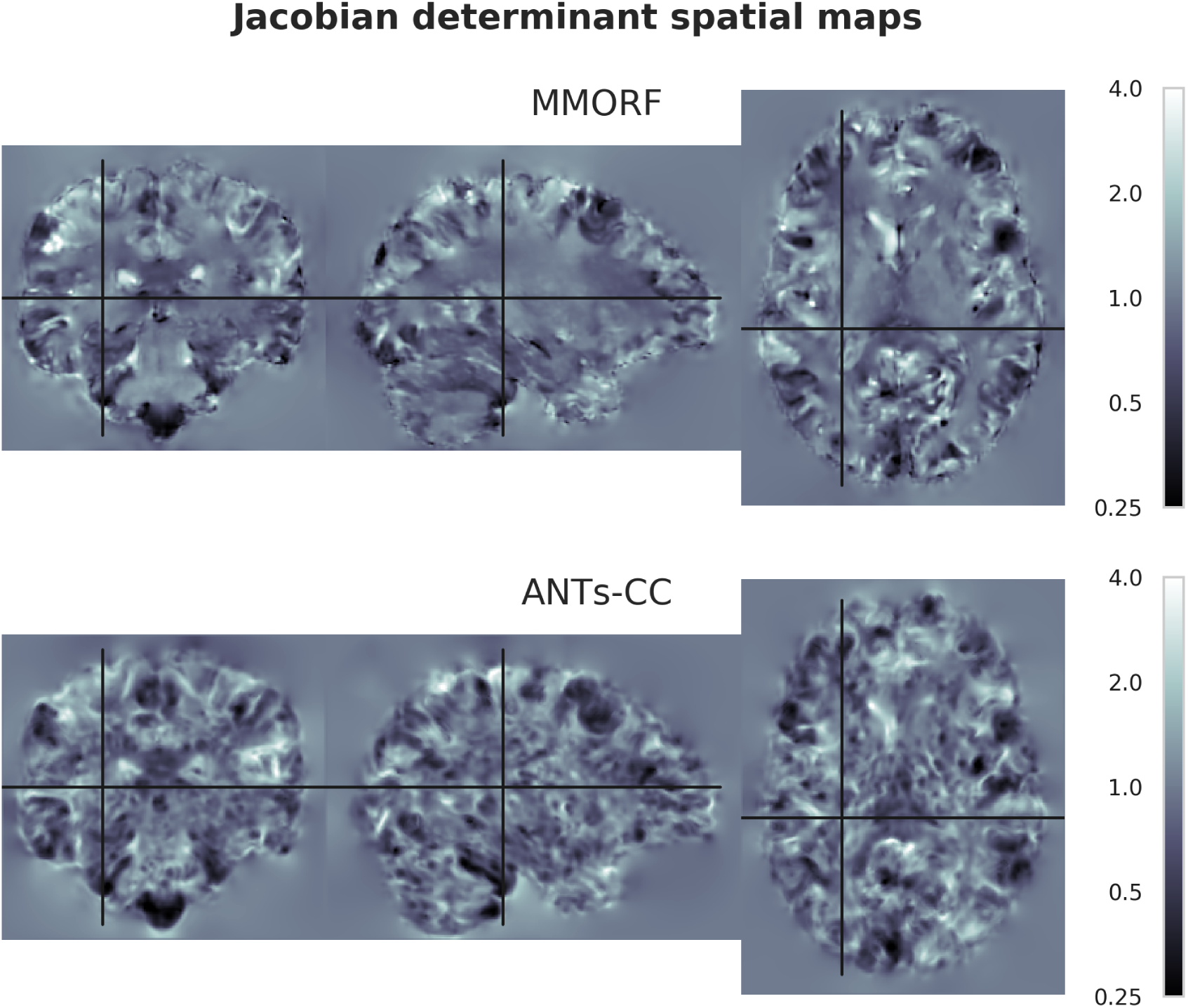
Spatial distribution of Jacobian determinants for MMORF and ANTs. Note the distinct visual differences within the white matter for ANTs-CC. The SPRED penalty has resulted in smooth changes within this region, whilst ANTs-CC has produced high frequency variations. As a T_1_ image contains little information within the white matter, the anatomical plausibility of those rapid variations is not clear. By maintaining smoothness within the white matter, MMORF produces a result that is more anatomically believable given the information present within the images being registered. Importantly, the irregularity introduced by ANTs-CC is not necessary for achieving high overlap scores, as Figures 1 and 3 show. Finally, whilst this may not have any deleterious effect on overlap scores of grey matter regions, the effect of transforming a modality rich in white matter information through such a warp could be significant.

**Figure 9:**
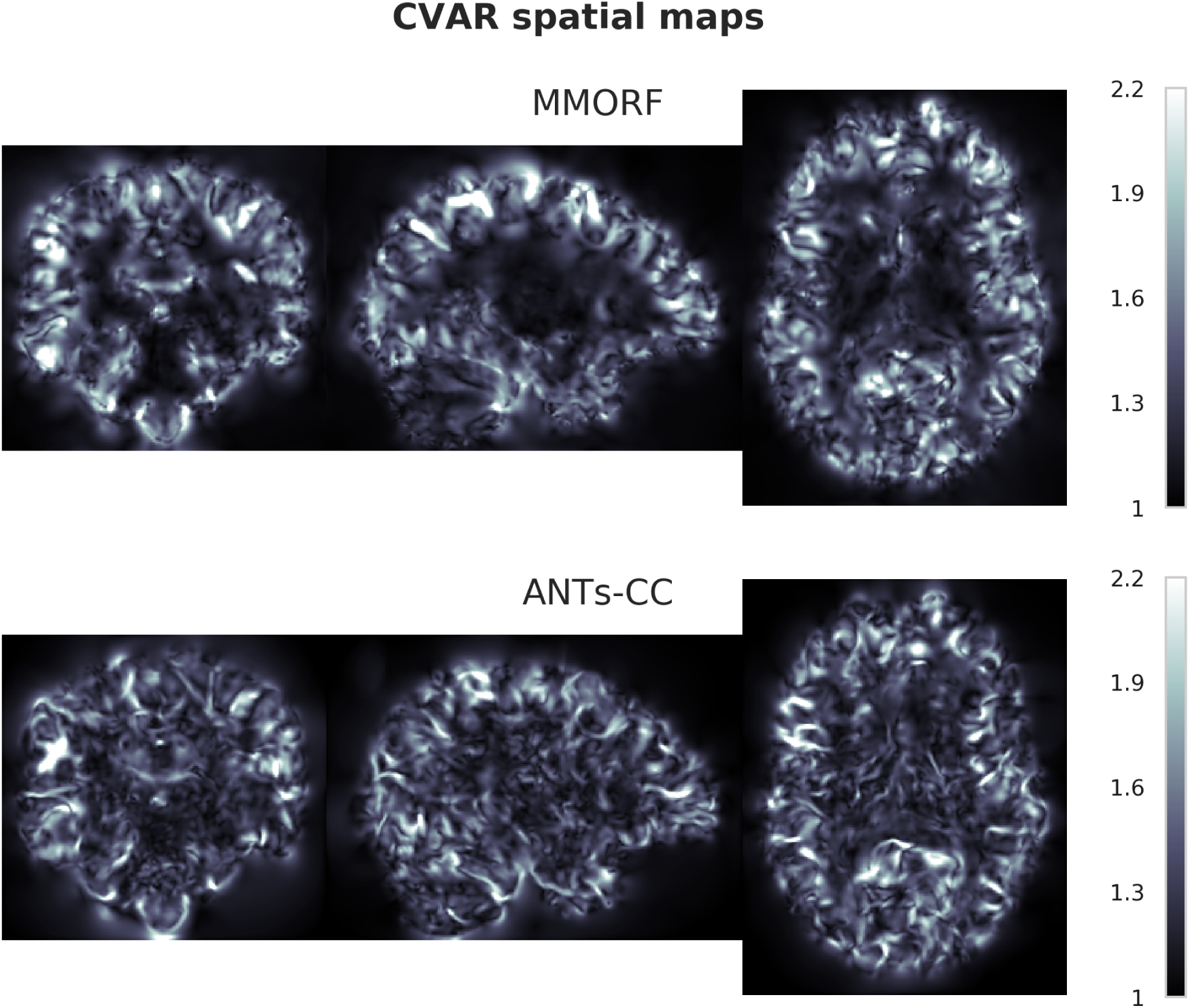
Spatial distribution showing the Cube-Volume Aspect Ratio (CVAR) for MMORF and ANTs-CC. CVAR is a measure of shape change which extends the concept of aspect ratio to arbitrary dimensions, and is described in detail in Appendix E. Values above 1 correspond to increasing deviations from the original shape. It represents the maximum linear relative change along any direction of the deformed voxels. Note that the appearance for MMORF and ANTs-CC is very similar to that in Figure 8.

As a T_1_-weighted volume contains minimal information within the white matter, there is little reason to believe that the deformation should be anything other than smooth within these regions. Clearly this has not detrimentally affected the performance of ANTs-CC in terms of achieving high cortical overlap scores. However, if this warp was used to transform a modality rich in white-matter information (such as diffusion tensor imaging) the results could potentially be very deleterious.

By contrast, the MMORF result is more balanced. In areas where there is a large amount of information to drive the deformation (tissue boundaries) the warps are correspondingly larger and more variable, whereas areas with minimal information are left relatively unchanged.

When looking at figure 9 it should be noted that the particular (randomly selected) registration we chose for displaying spatial maps happens to be one of 65 (out of 240) for which the mean CVAR was greater for MMORF than for ANTs-CC.

## 4. Discussion

We have implemented and investigated a previously suggested method for regularising warps by penalising volume and shape distortions (Ashburner and Friston (1999) and Ashburner et al. (2000)). The regulariser has some very attractive properties, but is computationally very expensive. This meant that the implementation in the original paper used a voxel-by-voxel optimising scheme rather than a global optimiser, which can potentially cause order effects and convergence to a non-optimal minimum. We implemented the regulariser on a GPU, using a B-spline basis to represent the warps and a Gauss-Newton optimiser, resulting in run-times of under half an hour for a full multi-pyramid registration. This has allowed us to run a large number of registrations and compare the results to one of the best performing algorithms in popular use (Avants et al. (2008), Klein et al. (2009)). We have been able to show that it performs as well as ANTs in terms of overlap scores, and that it achieves that with significantly less volume and shape distortions.

### 4.1. Relationship to Earlier Work

Pennec et al. (2005) suggested a very similar penalty function based on the concept of a Riemannian manifold for the allowed (diffeomorphic) warps. Other regularisers (Burger et al. (2013)) with similarities to that investigated by us are based on models for hyperelastic materials. These are models that describe the stress-strain relationships of materials, for example rubber, which are not well described by linear elastic models. These models have also been used as regularisation for correcting susceptibility artefacts in diffusion-weighted MRI (Ruthotto et al. (2012)). Similar hyperelastic models have additionally been used to model cortical growth in the developing brain (Knutsen et al., 2010) and for regularisation of surface based cortical registration (Robinson et al., 2018).

It should be noted that neither the present work, nor that in Ashburner et al. (2000), is explicitly based on hyperelastic models. Nor do we believe that hyperelastic models are necessarily meaningful descriptive generative models for explaining anatomical differences between subjects. Any such model (see for example Van Essen (1997)) is unlikely to be simple enough to be described by a few material constants. The reasoning behind our regularisation function is purely empirical, based on fulfilling our criteria for a useful function and on proving to produce smooth and plausible warps in parts of images with little or no anatomical information.

### 4.2. Beyond Enforcing Diffeomorphic Warps

It is widely agreed that invertibility is a desirable property in a nonlinear spatial transform. There are two principle ways in which this can be enforced

**Warp Construction** Any warp that is a composition (integration in the limit) of diffeomorphic warps is itself diffeomorphic. Hence, methods that estimate warps as compositions of many small updates will by construction ensure that the end result is invertible.

**Warp Penalisation** A highly nonlinear penalty function that goes to infinity as the local Jacobian determinant approaches zero will allow large deformations while maintaining invertibility. It should be noted that functions such as membrane energy or bending energy do not fall into this category.

For completeness both of these approaches will be discussed in more detail below. However, diffeomorphic warps are not the be all and end all. A warp can be invertible and yet highly unrealistic in that it causes very big volume and/or shape distortions. For this reason most methods that enforce invertibility using one of the methods mentioned above will combine that with one or more “traditional” regularisers that enforce smoothness. Ashburner and Ridgway (2013) has shown very convincingly that even within the space of diffeomorphic warps one will obtain very different solutions depending on the exact details and weights of the additional regularisers.

Hence, what we aim to achieve with our choice of regulariser goes much beyond just ensuring diffeomorphic warps. The aim is to find the maximally plausible, in terms of volume and shape distortions, of all possible warps within the space of diffeomorphisms. And to do this with a single penalty function (and a single weight) that achieves the joint objective of ensuring invertibility and simultaneously minimising volume and shape distortions.

We recognise that a judicious choice of forms and weights of additional penalty functions within inherently diffeomorphic frameworks, such as LD-DMM and viscous fluid based methods, could potentially find an equally advantageous solution. But it is nevertheless the case that when comparing our regulariser to a state-of-the-art diffeomorphic method we were able to obtain invertible warps with equally good overlap scores and significantly less volume and shape distortions. Furthermore, there is no intrinsic superiority of inherently diffeomorphic methods over any other method that also guarantees invertibility, but achieves that with equally good, or better, registration accuracy.

#### 4.2.1. Diffeomorphism by Warp Construction

Methods such as ANTs fall into the category of inherently diffeomorphic warps. In particular, ANTs is an example of a greedy approximation of the general LDDMM method. LDDMM methods seek to find a time varying velocity field which minimises a metric based on total path length in the space of diffeomorphisms. Originally introduced by Beg et al. (2005), these methods guarantee that the total deformation remains diffeomorphic when the velocity field is integrated over sufficiently small timesteps. A downside to these methods is the large number of parameters which need to be estimated at each timepoint. Tools such as DARTEL (Ashburner, 2007) seek to overcome this by instead parametrising a stationary velocity field, thereby greatly reducing the parameter space, but at the expense of potentially larger path lengths. Subsequently, Geodesic Shooting (Ashburner and Friston, 2011) based methods have been able to reformulate the original LDDMM problem such that instead of estimating the entire sequence of time-varying velocity fields, only the initial velocity need be estimated, thereby greatly reducing both the parameter space and convergence time of the optimisation.

Whilst all of these methods are capable of ensuring warps remain diffeomorphic, they differ from our method in that the prior on which they are based is that deformations follow the shortest path-lengths, rather than deformations conserving the shape of underlying anatomy. As such, diffeomorphism is an intrinsic rather than a controlled property of the transformation model. It is certainly not impossible to include explicit regularisation of shape changes in these models, however it is not the norm.

#### 4.2.2. Diffeomorphism by Warp Penalisation

An alternative method of enforcing diffeomorphism is by using a penalty function that goes to infinity as the local Jacobian determinant approaches zero.

Most commonly, the Jacobian determinant itself is used as a hard constraint that bounds the allowed range (Haber and Modersitzki, 2004; Sdika, 2008, 2013; Haber et al., 2010), or fixes it to a particular value (Mansi et al., 2011), or a smoothly varying function of the Jacobian determinant is used as a soft constraint (Borz`i et al., 2003; Yanovsky et al., 2007; Leow et al., 2007; Heyde et al., 2016; Mang et al., 2018). It should be noted that when modelling the warps as a velocity field the local divergence can be used as a proxy for the Jacobian determinant.

The crucial difference between our penalty and those based on a function of the Jacobian determinant is that the latter *only* penalises volume distortions. Not only does that mean that shape changes are not penalised, in practice such a function will result in a method where very large shape changes are used to circumvent the volume change limitations. In Appendix E we show the results obtained with a Jacobian determinant penalty function (Σ_*n*_ *v* (1 + |**J**_*n*_|) log^2^ |**J**_*n*_|) where the weight was calibrated so as to yield the same overlap score as our SPRED function. It can be seen that the Jacobian determinant penalty yields equally good overlap scores (figures E.1 and E.2) and results in warps that are diffeomorphic and with a very narrow range of Jacobian determinants (figures E.3 and E.5). But this has been achieved by completely ignoring shape changes, and has resulted in much larger CVARs (figures E.4 and E.5). Looking at a randomly selected warp (figure E.6) it is very clear that the Jacobian determinant penalty has resulted in lots of gratuitous distortions.

This is the reason that many groups have combined a hard or soft constraint on volume changes with an additional regulariser (see for example Haber et al. (2010), Yanovsky et al. (2007), Leow et al. (2007) or Mang et al. (2018)). While that may work well, it means that an additional weight factor needs to be empirically determined.

There is another group of of algorithms (Loeckx et al., 2004; Staring et al., 2007; Modersitzki, 2008; Mang and Biros, 2016) that use functions that penalise deviations from local rigidity (*i.e.* anything beyond local translation and rotation). In that respect they have similarities to our penalty function, but they have mostly been used to enforce (close to) total rigidity in parts of the image for specific applications such as movement of objects that have a mix of rigid (for example bone) and non-rigid tissue (for example muscle).

Of this category of registration methods, Mang and Biros (2016) is the most closely to ours of which we are aware. They acknowledged the issue of excessive shape changes and sought to combat it by explicitly penalising the shear of the velocity field.

### 4.3. Other Parallelised Algorithms

It is increasingly common for registration tools to employ some form of parallelisation, and it is worth contextualising our approach in comparison to some of these methods.

Certain Insight Segmentation and Registration Toolkit (Yoo et al., 2002) methods use CPU multithreading on a single device to accelerate the portions of their code which solve the linear system of equations necessary to compute update steps. This is a simple method to implement, but gains tend to be modest.

A recently described (Mang et al., 2018) parallelised version of Mang and Biros (2016) utilises distributed-memory parallelism on multiple CPU nodes. It uses a Gauss-Newton optimisation strategy, and solves for a stationary velocity field transformation. The primary focus of the parallelism is in the *inversion*, rather than the *calculation*, of the Hessian.

Possibly the most closely related method to ours is NiftyReg (Modat et al., 2010) which employs a B-spline transformation and GPU parallelisation. NiftyReg differs in using a first order optimisation strategy and therefore does not require calculation of the Hessian. Reported performance improvements are very similar to ours, indicating comparable levels of code optimisation. Their impressive sub-minute runtimes are aided by a computationally simpler bending energy regularisation metric. Therefore a direct comparison of runtimes to our method using SPRED is not meaningful.

As discussed in Eklund et al. (2013), most applications of GPU parallelisation to medical image registration focus on speeding up existing methods. However, an interesting alternative is the development of more advanced registration methods which might otherwise have been rejected on the basis of computational complexity.

### 4.4. Registration Accuracy

As we have previously stated, absolute registration accuracy is not the primary focus of this work. Instead it stands as a reference point for our discussion of anatomical plausibility, in that only two methods which are comparable in terms of registration accuracy can be meaningfully differentiated based on anatomical plausibility.

Based on the overlap scores in Section 3.1 we see that MMORF and ANTs-CC are clearly the best performing tools in terms of registration accuracy. Additionally there is very little to differentiate between their performance as a whole. We note that the results for ANTs-CC are slightly better than those which have been previously reported (Ou et al., 2014) for the same validation data set. We therefore believe that the way we have used ANTs-CC has yielded a close to optimal performance on this dataset.

### 4.5. Anatomical Plausibility

Based on the summary 5^th^ to 95^th^ percentile log-Jacobian determinant range metric, MMORF is significantly more anatomically plausible than ANTs-CC. Figure 6 supports this argument, by showing that this is true for a vast majority of the registrations. In other words, for a given registration accuracy one is almost guaranteed to have lower volumetric changes when using MMORF over ANTs-CC. Another way of framing this result is to say that larger volume distortions are not a necessary trade-off for achieving high registration accuracy.

A striking feature of the Jacobian-range (figure 8) and CVAR maps (figure 9) are the seemingly gratuitous warps in white matter. This could potentially be a particular problem if one intends to use the structural registration to transform diffusion data. We believe this to be a strong case for extending the notion of anatomical plausibility beyond that of simply maintaining diffeomorphism and for building that notion into the registration.

Why this behaviour is observed in ANTs-CC is an interesting question in and of itself. At first this might be thought to be a case of insufficient regularisation, however the deformations generated by ANTs-MSQ (data not shown) did not display this same behaviour. Therefore we must posit that this is due to the somewhat *scale-free* nature of the CC metric. In other words, the gradient of CC does not depend on the amount of signal contrast present, only on the local correlation of that signal. What this means in practice is that regularisation on the level of the gradient cannot alter this behaviour, rather regularisation of the metric itself would be required. How to achieve this is beyond the scope of this work, however it highlights the importance of taking a considered approach to evaluating the believability of a deformation.

### 4.6. Limitations of the current work

The aim of the present paper is to introduce the SPRED penalty and to show that it can be used to yield large deformation, diffeomorphic and anatomically plausible warps. It is not yet a finished registration package that deals with differences in contrast, receive bias-field or B1 inhomogeneities. These will be the subject of future work. Neither have we applied it to data where *very* large deformations are needed, such as when registering images of severely atrophied brains or inter-species registration. Finally, whilst our regularisation penalty is symmetric, our similarity measure is not. Thus, for the overall algorithm to be truly symmetric we would ideally either modulate our similarity measure by (1 + |**J**|) (Tagare et al., 2009), or simultaneously register both images to a mid-space (Avants et al., 2008).

### 4.7. Performance Summary

The SPRED penalty allows MMORF to overcome the oft-quoted limitations of small deformation frameworks and achieve levels of registration accuracy comparable to state-of-the-art large deformation tools such as ANTs-CC. MMORF achieves this whilst at the same time producing warps which are systematically less aggressive in terms of both volume and shape distortions. Finally, given the information present in a structural scan, the spatial distribution of the log-Jacobian determinants MMORF produces appear more plausible compared to ANTs-CC, which produces variations that are not obviously necessary.

Overall we conclude that SPRED as implemented within MMORF produces warps which can be used with a high level of confidence. MMORF’s registration accuracy is on par with state-of-the-art volumetric methods, and therefore no sacrifice need be made in terms of maximising data consistency.

## 5. Conclusion and Outlook

We have demonstrated that our SPRED penalty is capable of matching the registration accuracy (as measured using overlap scores of manually segmented cortical regions) of the most well established large-deformation (ANTs-CC) framework available today. Additionally, we find that the results from using the SPRED penalty are consistently more anatomically plausible both in terms of Jacobian determinants and CVARs. We leverage GPU parallelisation in order to make optimisation of the penalty computationally tractable, allowing this method to be of practical benefit as part of a usable registration framework.

We have shown that using a regularisation function such as this can overcome the shortcomings of elastic small deformation frameworks, yielding the best of both worlds: large deformation, diffeomorphic warps with minimal distortions and high registration accuracy. Future work will focus on extending MMORF to include the simultaneous registration of scalar and tensor modalities (see, *e.g.*, Irfanoglu et al. (2016)). Furthermore we wish to investigate the effect of spatially varying the weighting of the SPRED penalty based on prior information regarding tissue variability (*e.g.*, allowing larger deformations within the ventricles).

## Supporting information

Supplemental Code Profiling

Supplemental Registration Framework

Supplemental Jacobian Determinant Comparison

## Acknowledgements

The Wellcome Centre for Integrative Neuroimaging is supported by core funding from the Wellcome Trust [grant number 203139/Z/16/Z]. The Wellcome Centre for Human Neuroimaging is supported by core funding from the Wellcome Trust [grant number 203147/Z/16/Z].

This work was supported by funding from the Engineering and Physical Sciences Research Council (EPSRC) and Medical Research Council (MRC) [grant number EP/L016052/1]; and the NVIDIA Corporation GPU grant program.

FJL is supported by The Oppenheimer Memorial Trust, St Catherine’s College Oxford and The Rotary Foundation.

JLRA is supported by a Wellcome Trust Strategic Award [grant number 098369/Z/12/Z]; and the NIH Human Connectome Project [grant numbers 1U01MH109589-01 and 1U01AG052564–01].

## Conflicts of Interest

We have no conflicts of interest to declare.

## Appendix A. Penalty Approximation

In this section we provide the derivation of the following approximation:

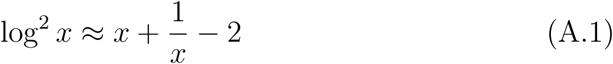

Let:

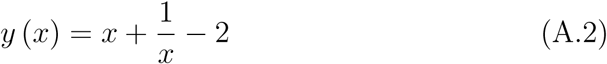

And:

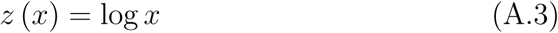

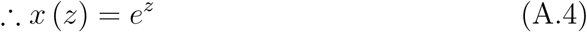

Then:

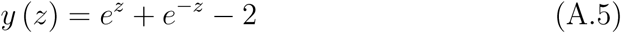

Now, taking the Taylor expansion of *y* (*z*) about *z* = 0:

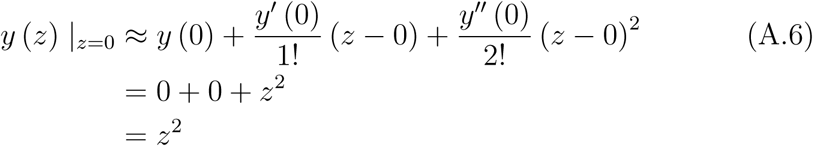

And finally, using Equation A.3:

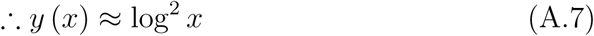

Figure A.1 graphically demonstrates the relationship between the exact and approximate penalty functions.

**Figure A.1:**
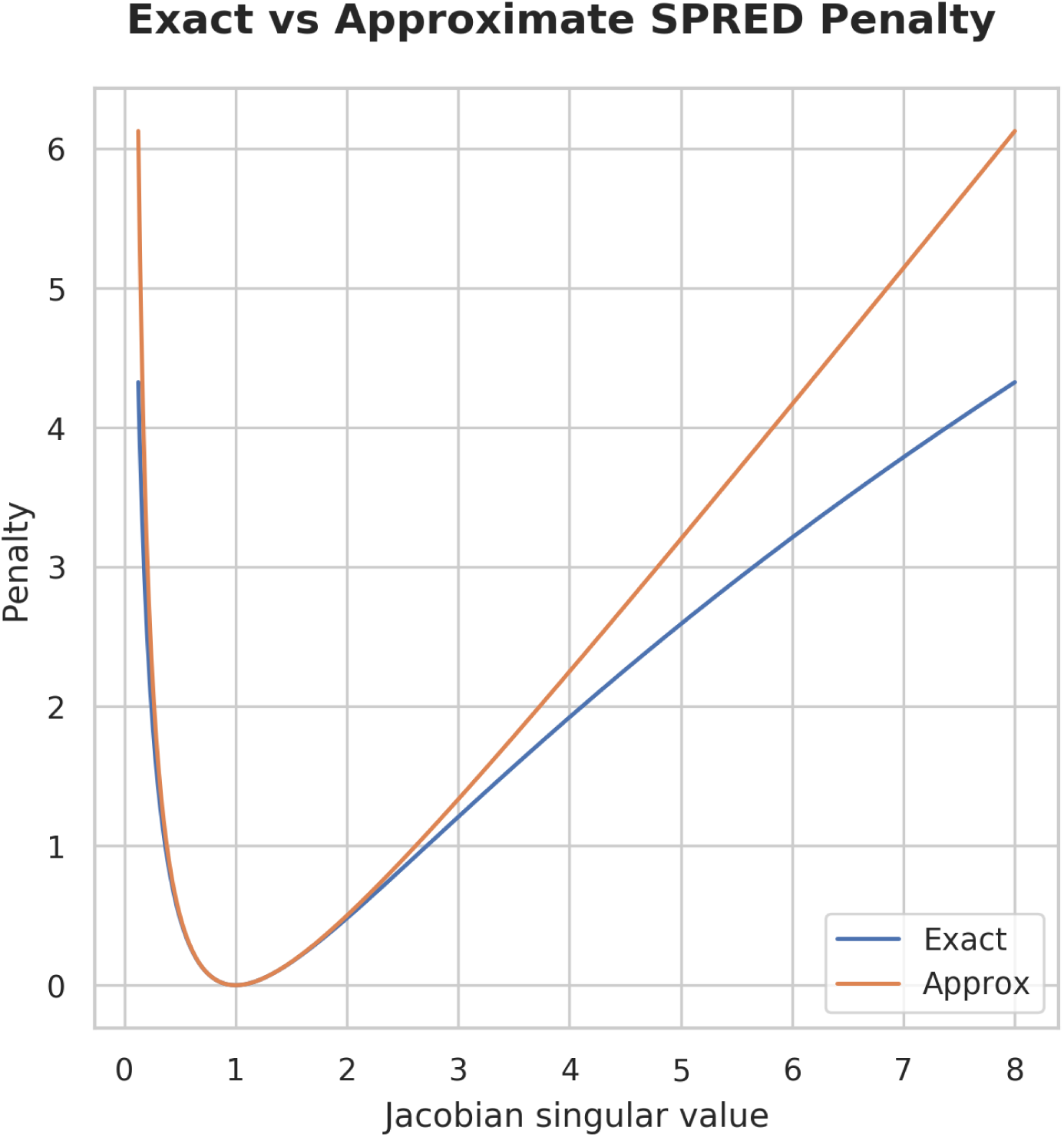
Effect of the approximation in Equation A.1 on the SPRED penalty. The approximation is almost indistinguishable from the exact penalty for singular values near 1. For singular values between 0.25 and 4 the approximation is accurate to within 17 %, and within 42 % between 0.125 and 8. Note that the approximation maintains the desirable symmetry for expansions and contractions. Note too that the approximation majorises the exact penalty (*i.e.*, the approximation is always greater than or equal to the exact penalty), and therefore maintains the exact penalty’s ability to ensure diffeomorphism.

## Appendix B. 2D Example

We begin by defining a displacement warp field and its values in the underlying image space. The image space is a square with dimensions 21 × 21. The cubic B-splines are placed with their control points (knots) 5 samples apart. Each spline has a spatial support radius of 2 knots (10 samples), and therefore we place an extra ring of splines around the perimeter of the image space, such that each point in the image has an equal number of splines with support there. As a result, our *x*-warp and *y*-warp parameters are of dimension 7 × 7. We initialise the warp with a random set of parameters. Figure B.1 shows the warp parameters, the resulting warp in image space, and the gradient of the warp in image space.

Based on the spatial gradients of the displacement fields in Figure B.1, the local Jacobian matrix can be calculated at each point in space according to Equation 7.

The elements of **J** for each point in the sample space, as well as the resulting Jacobian determinant, |**J**|, are shown in Figure B.2, which now contains all of the information necessary to calculate the voxelwise part (Equation 22) of the calculations for the gradient (Equation 19).

Based on the elements of **J**, Figure B.3(a) shows the resulting contribution to the total cost from each point in the image. B.3(b) to B.3(e) then constitute the gradient of the cost with respect to the elements of **J** at each point in the image, i.e., the columns of [*∂c/∂***J**] (Equation 22). Each point in the four partial derivative images can be calculated completely independently of every other point. The calculation of these images therefore constitutes the first parallelised step of our algorithm

We now move onto visualising the result of Equation 25. Note that Equation 24 is constant across the image and identical over the support of every spline. Therefore, when calculating the gradient in a parallel-per-parameter regime, the operation conforms exactly to the SIMD paradigm and may be very efficiently implemented on GPUs. In practice, we perform one such calculation for each combination of **J** element gradient image (Figure B.3(b) to B.3(e)) and warp direction (*x* or *y*), the results of which are shown in Figure B.4(b) to B.4(e). Finally, we sum over these combinations for each warp direction, and the result is then the gradient of our penalty with respect to our warp parameters. This is shown in B.4(f) and B.4(g).

**Figure B.1:**
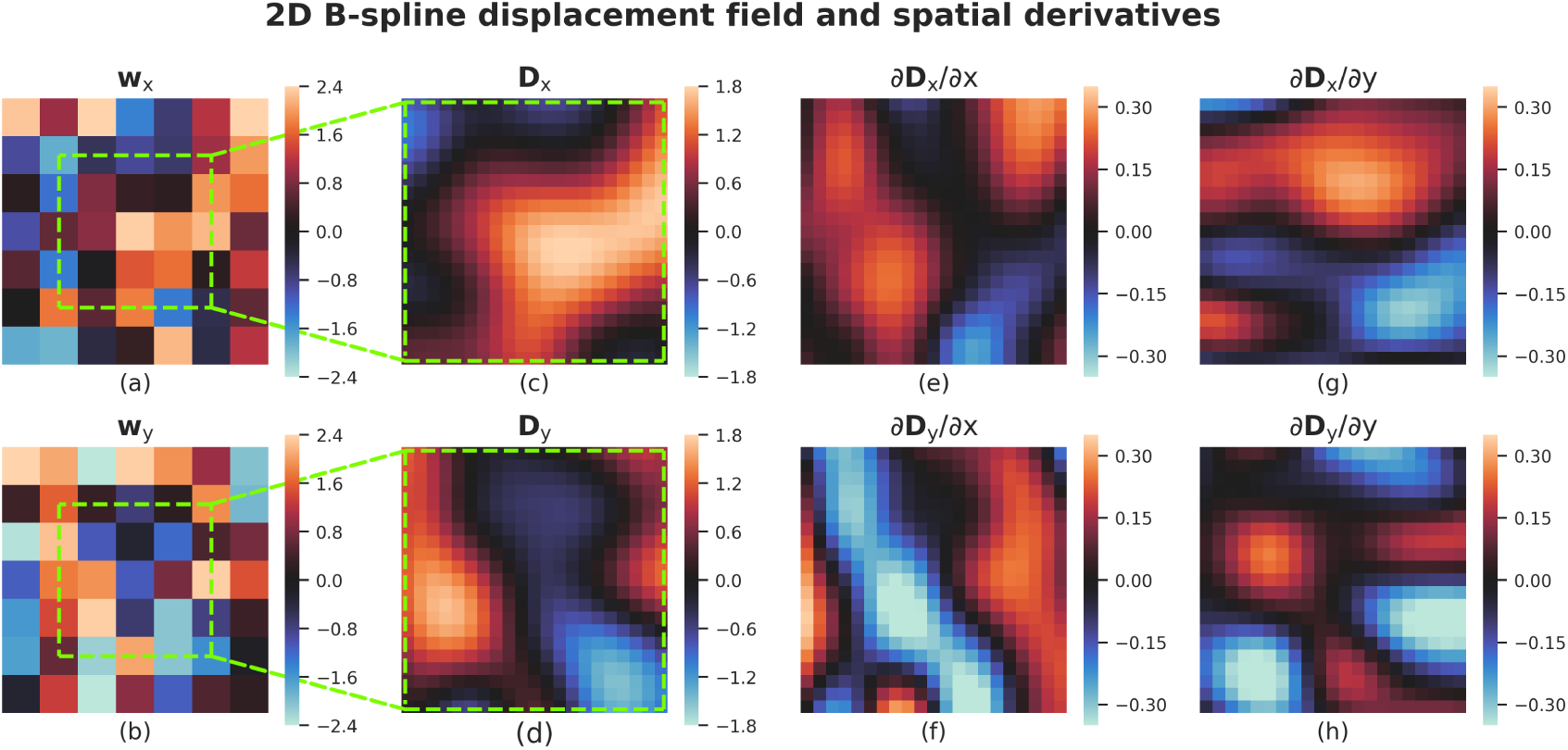
Example of a warp (or technically *displacement*) field for a random set of parameters (cubic B-spline coefficients). The positive *x*-direction is defined from left to right, and the positive *y*-direction from top to bottom. The *x*-warp parameters **w**_*x*_ (a) and *y*-warp parameters **w**_*y*_ (b) are arranged on a regular 7 × 7 grid. The resulting *x*-displacement field **D**_*x*_ (c) and *y*-displacement field **D**_*y*_ (d) corresponding to **w**_*x*_ and **w**_*y*_ are evaluated on a regular grid with knot spacing of 5 × 5. Note that only those parts of the warps with full support of the surrounding B-splines are shown (i.e., the area outside of the green square is considered to be *outside* of the image we are warping). This leaves a square of 21 × 21 samples in image space. The spatial partial derivatives of the *x*-warp in the *x*-direction (e), *y*-warp in the *x*-direction (f), *x*-warp in the *y*-direction (g) and *y*-warp in the *y*-direction (h) are presented, as these values are used in determining the local Jacobian matrices of the warp field. Cubic B-spline parametrisation ensures that both the warps and their spatial derivatives have analytical solutions and are smooth-continuous.

**Figure B.2:**
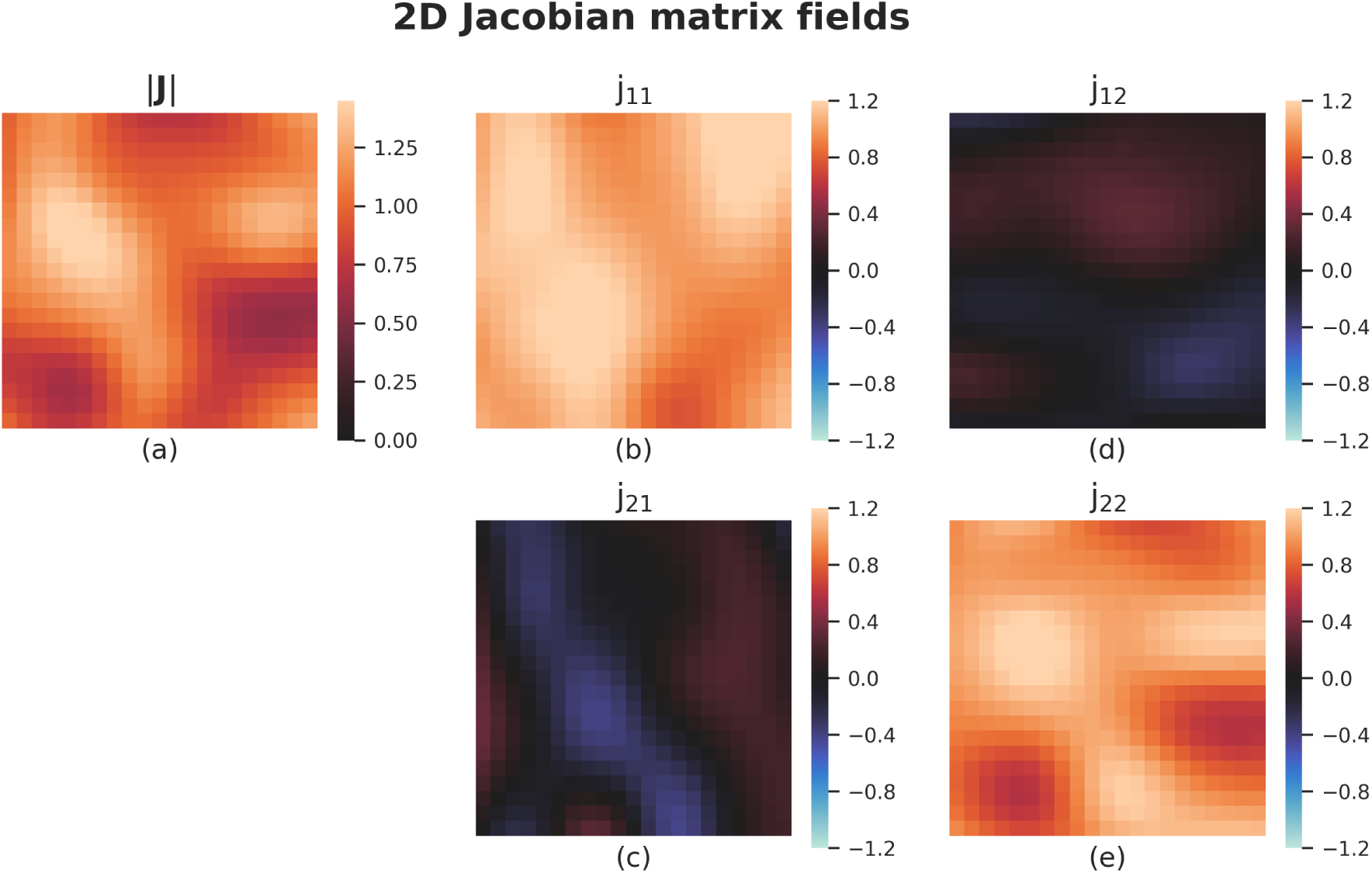
Elements of the local Jacobian matrix of the warp. At each position [*x, y*] in image space a local Jacobian matrix **J** is calculated according to Equation 7 by adding the identity matrix *I*_2_ to the 2 × 2 matrix composed of the elements of Figure B.1 (e) to (h) at that position. The resulting 2 × 2 2D fields (each showing how one element of **J** varies across the image) are presented here in sub-figures (b) to (e). From these a local Jacobian determinant field (a) can be calculated. The SPRED penalty is based on the singular values of **J**, and is therefore a function of fields (b) to (e).

**Figure B.3:**
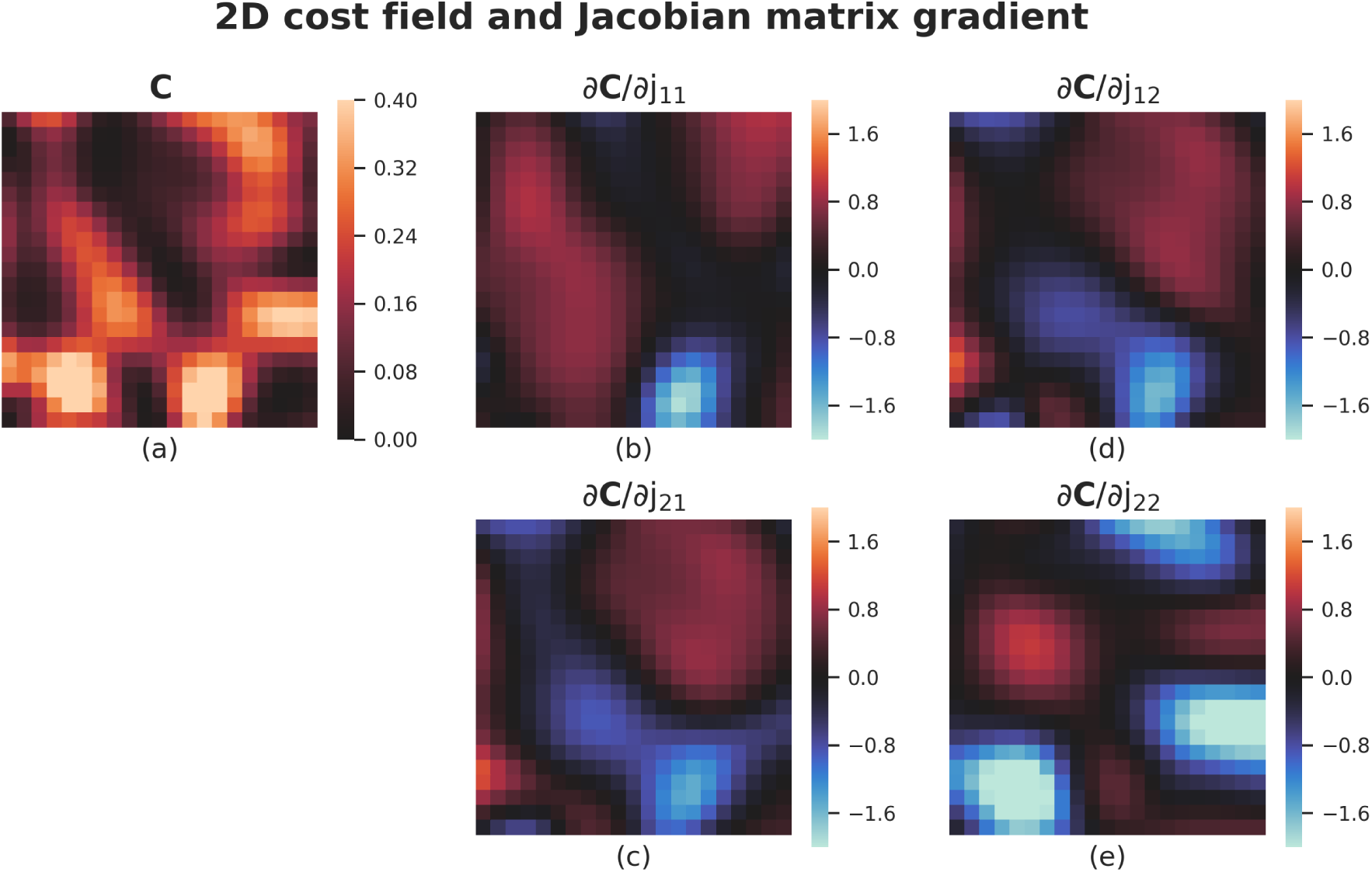
From the singular values of the local Jacobian matrices of Figure B.2 a cost field **C** (a) can be evaluated at each position [*x, y*] of the image. **C** then represents the contribution to the total cost from each point in image space. The partial derivatives of **C** with respect to the elements of **J** are shown in sub-figures (b) to (e). These partial derivative fields (which together make up the overall gradient *∂***C***/∂***J**), can be calculated independently for each position [*x, y*] as they depend *only* on the values of **J** at [*x, y*], making this step ideal for a parallel-per-voxel approach.

**Figure B.4:**
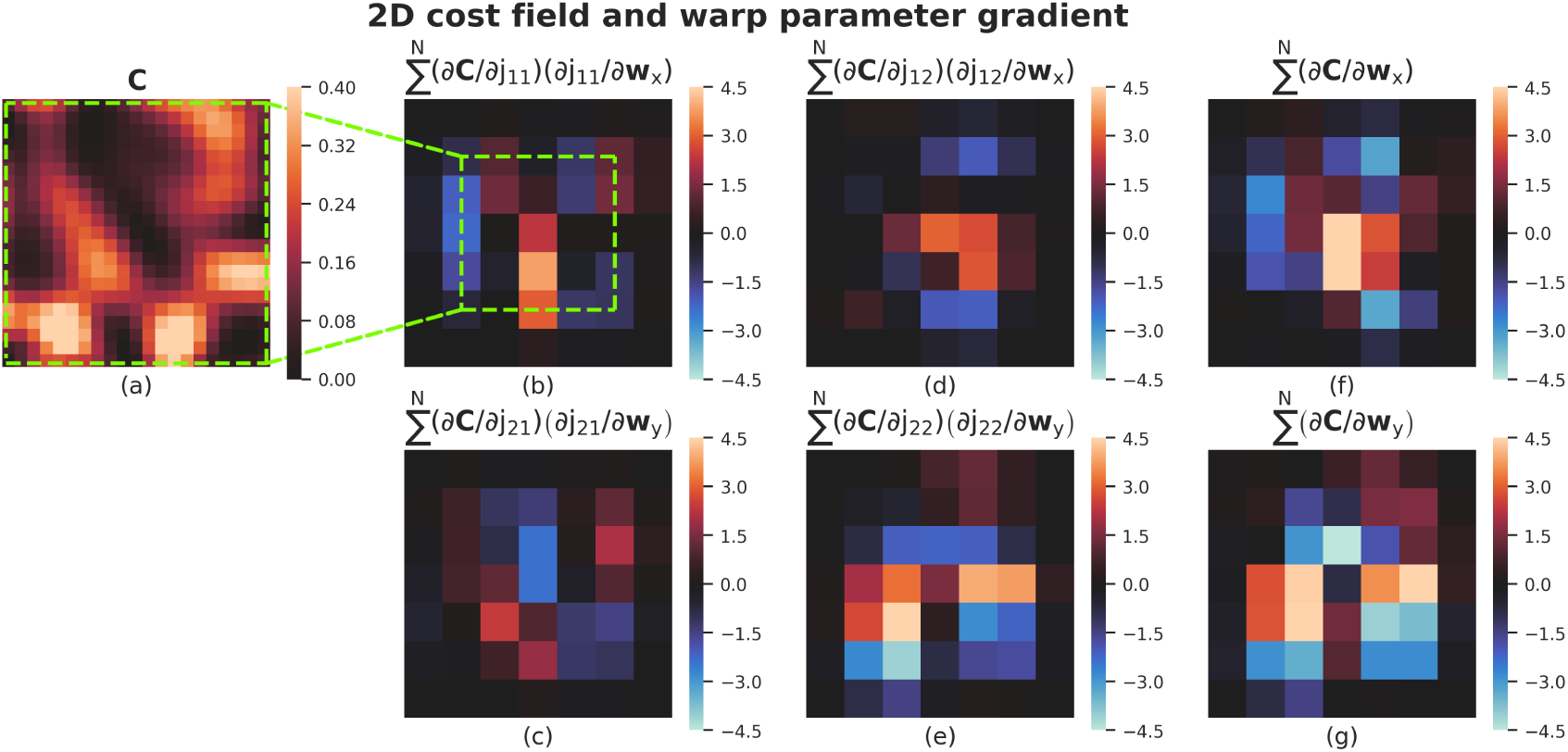
The final, and most computationally expensive, step is calculating the gradient of the total SPRED penalty (being the sum of **C** over all *x* and *y*) with respect to the warp parameters. Given that we have the gradient of **C** with respect to **J**, knowing the gradient of the resulting partial derivative fields (i.e., Figure B.3 (b) to (e)) with respect to the warp parameters would allow us to apply the chain rule to calculate this. We proceed by recognising that the piece-wise polynomial nature of B-splines means that the gradient of **J** with respect to each warp parameter is constant. Additionally, the gradient at position [*x, y*] is exactly 0 for any B-spline which does not have support at that position. Thus, we can calculate the total gradient with respect to each warp parameter by summing the result of the chain rule across each position [*x, y*] where that spline has support. In (b) to (e) we can see this illustrated in terms of the contributions to the overall gradient due to the effect of the warp parameters **w**_*x*_ and **w**_*y*_ on each element of the **J** fields. These partial derivatives can be calculated independently for each warp parameter, making this step ideal for a parallel-per-parameter approach. The end result of the process is then the gradient of the total penalty with respect to **w**_*x*_ (f) and **w**_*y*_ (g).

## Appendix C. Parameter Selection

Parameter selection in image registration is non-trivial and often leads to discussions regarding whether or not the particular parameters selected were optimal. As such, the aim of parameter selection in this work was not necessarily to find the absolutely optimal set of parameters for each method, but rather to find parameters which made the various methods equal in one metric (such as accuracy in terms of overlap scores), in order so that differences in another metric (such as aggressiveness in terms of of Jacobian determinant distributions) can be meaningfully compared.

### Appendix C.1. MMORF

We fixed the number of steps in the warp resolution pyramid (40, 20, 10, 5, 2.5 and 1.25 mm isotropic), and then optimised over three parameters, namely:

- Smoothing as function of warp resolution (*λ*_*s*_)
- Initial regularisation (*λ*_*ri*_)
- Rate of decrease of regularisation (*λ*_*rd*_)

The FWHM of the smoothing kernel was defined as the current warp resolution divided by *λ*_*s*_. The initial regularisation *λ*_*ri*_ is in arbitrary units and its exact value is not of interest. The total regularisation at each step of the resolution pyramid was *λ*_*ri*_ × (*λ*_*rd*_)^*it*^ where *it* is the current position in the resolution pyramid (i.e. 0 to 5).

The parameter values were chosen via a grid search of the parameter space. Half of the NIREP dataset (8 subjects, 56 warps) were used, and overlap scores calculated as per Section 2.5.4. The parameters which resulted in the best overlap scores were then used for the rest of the analysis. The final parameters were:

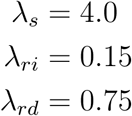

These parameters are relatively aggressive, but as we were aiming for maximum overlap scores this is to be expected.

### Appendix C.2. FNIRT

The parameters for FNIRT were chosen to match those used by the tool’s author in (Andersson et al., 2019).

### Appendix C.3. ANTs

For both ANTs-CC and ANTs-MSQ the number of steps in the resolution pyramid were fixed (10, 6, 4, 2 and 1 mm isotropic). Smoothing was chosen to match MMORF as closely as possible at each resolution level. This left three parameters over which to optimise, namely:

- Gradient step (*λ*_*g*_)
- Update field variance (*λ*_*u*_)
- Total field variance (*λ*_*t*_)

*λ*_*g*_ determines how far each point is allowed to move on each iteration, and increasing this value allows for higher frequency deformations. *λ*_*u*_ determines how much to smooth the gradient field between updates and increasing this value increases smoothness in the velocity field. *λ*_*t*_ determines how much to smooth the total displacement field.

The parameters were selected such that the MSQ after registration was similar to that of MMORF. Note that despite our best efforts, we were unable to get ANTs-MSQ to match MMORF and ANTs-CC in terms of this metric. The best results for ANTs-MSQ were achieved with:

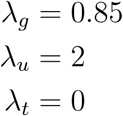

And for ANTs-CC with:

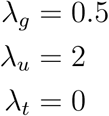

*λ*_*t*_ was found to always increase MSQ and was therefore left at the default value of 0. The final values for *λ*_*g*_ and *λ*_*u*_ are more aggressive than the default (fairly conservative) values of *λ*_*g*_ = 0.1 and *λ*_*u*_ = 3, but we found that this was necessary to achieve the required registration accuracy for a fair comparison.

Note also that the ANTs-CC parameters are very similar to those supplied by the tool’s author for use in Klein et al. (2009).

## Appendix D. Cube-Volume Aspect Ratio

To facilitate a quantitative comparison of registration aggressiveness we require metrics to measure changes in both volume and shape. The Jacobian determinant is useful in that it provides a single number quantifying volumetric changes. Similarly, we use the Cube-Volume Aspect Ratio (CVAR) to describe the deviation in shape of any cuboid from a regular cube (Smith and Wormald, 1998). CVAR in 3 dimensions is defined as: *The cube-root of the ratio of the volume of the smallest regular cube which can fully enclose the cuboid, to the cuboid’s own volume*. Alternatively, for a deformation we may equally well define it as: *The cube-root of the ratio of the largest Jacobian singular value cubed, to the Jacobian determinant* (see Equation D.1).

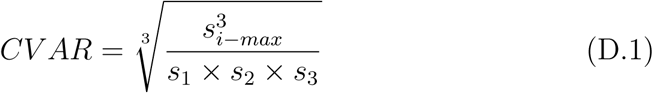

CVAR therefore takes on a value of 1 for a regular cube, and is greater than 1 for any other shape.

## Appendix E. log JSV vs log *|*J*|* regularisation

Given that we believe the Jacobian determinant is a good measure of warp aggressiveness, the question may arise as to why we prefer to penalise the Jacobian singular values rather than the determinant itself. We have stated our reasoning in the main text, that simply penalising the determinant will lead to implausible changes in shape, in an attempt to minimise volumetric changes. Here we provide an example demonstrating this in action.

### Appendix E.1. Testing

We registered all subjects of the NIREP dataset to subject NA01 only (i.e. 15 warps in total) using two different regularisation methods. In the first instance we used the SPRED penalty as described in the main text, and in the second we used the following simpler penalisation of the Jacobian determinant:

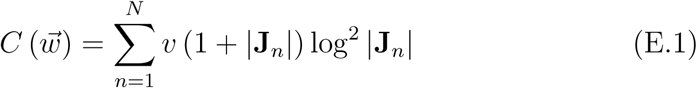

The regularisation weighting was chosen such that the average overlap scores were comparable between the two methods. We then calculated overlap scores as in Section 2.5.4 of the main text. For each warp we then calculated the minimum, maximum, mean, 5^th^ and 95^th^ percentile for both the Jacobian determinant and CVAR metrics. Overlap scores are presented in Figures E.1 and E.2. Jacobian determinant ranges are presented in Figure E.3, and mean CVAR in Figure E.4. Spatial maps of both Jacobian determinants and CVAR are presented in Figure E.5. Finally, Figure E.6 shows a grid deformed under both regularisation schemes.

**Figure E.1:**
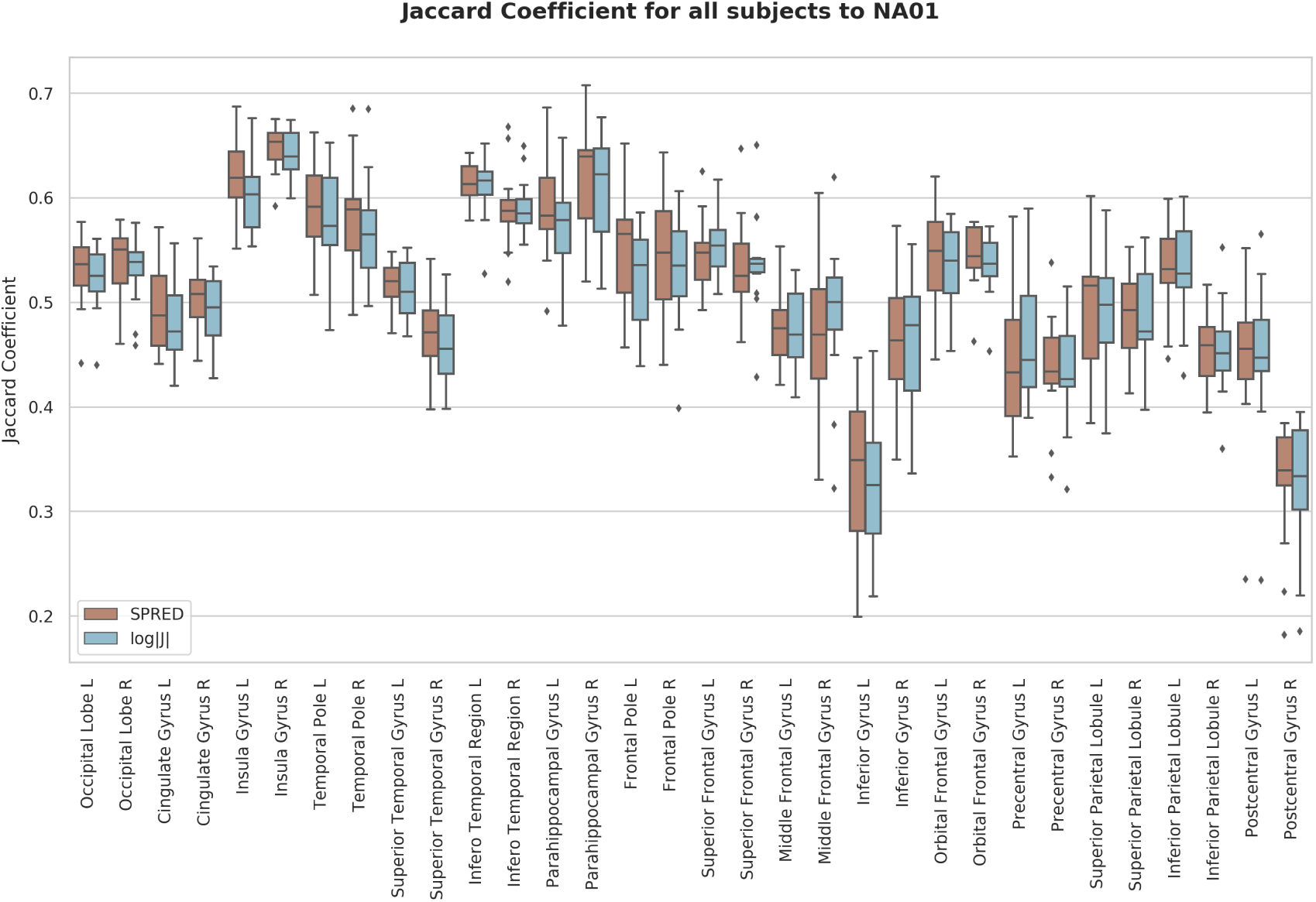
Jaccard coefficients of 32 cortical segmentations for registration to subject NA01 of the NIREP dataset. We compare the SPRED penalty to a simpler log |**J**| penalty. The results are comparable between the methods in terms of average overlap score.

### Appendix E.2. Discussion

From Figures E.1 and E.2 we see that the choice of regularisation metric has not affected the overall registration accuracy. Additionally, from Figure E.3 we note that, as expected, penalising the Jacobian determinant directly leads to warps with less volumetric changes. However, Figure E.4 shows that this comes at the cost of increasing the distortions in shape as measured by CVAR. Adding an additional penalty term (such as Bending Energy) to the Jacobian determinant regularisation could reduce these shape changes, however this would introduce a second regularisation weighting to tune. In this respect the SPRED penalty is far more elegant, as it effectively controls both volumetric and shape changes, preserves topology, and requires only a single tuning parameter.

Figure E.5 provides an intuitive appreciation of the information in Figures E.3 and E.4, with the extent to which volumetric and shape changes are traded off by the different regularisation schemes becoming clearly evident. Finally, from E.6 we see the extent to which higher CVAR values translate into more convoluted warps with reduced spatial smoothness as compared to SPRED.

In conclusion, this example shows that simply penalising the Jacobian determinant is insufficient to ensure plausible deformations.

**Figure E.2:**
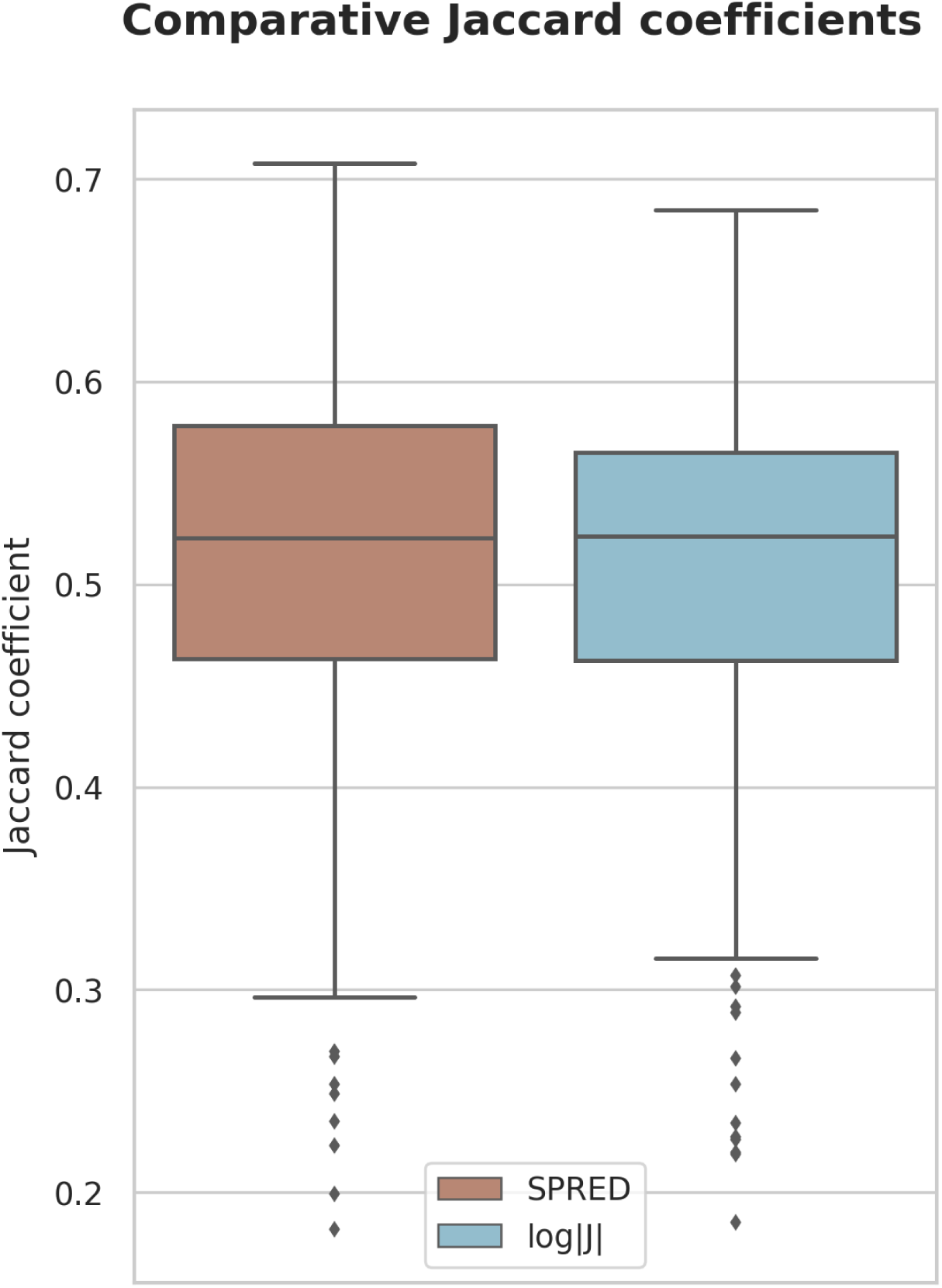
Distribution of Jaccard coefficients combined across all regions and subjects pairs for both regularisation penalties. This confirms what is seen in Figure E.1, that both methods have similar overlap accuracy on average.

**Figure E.3:**
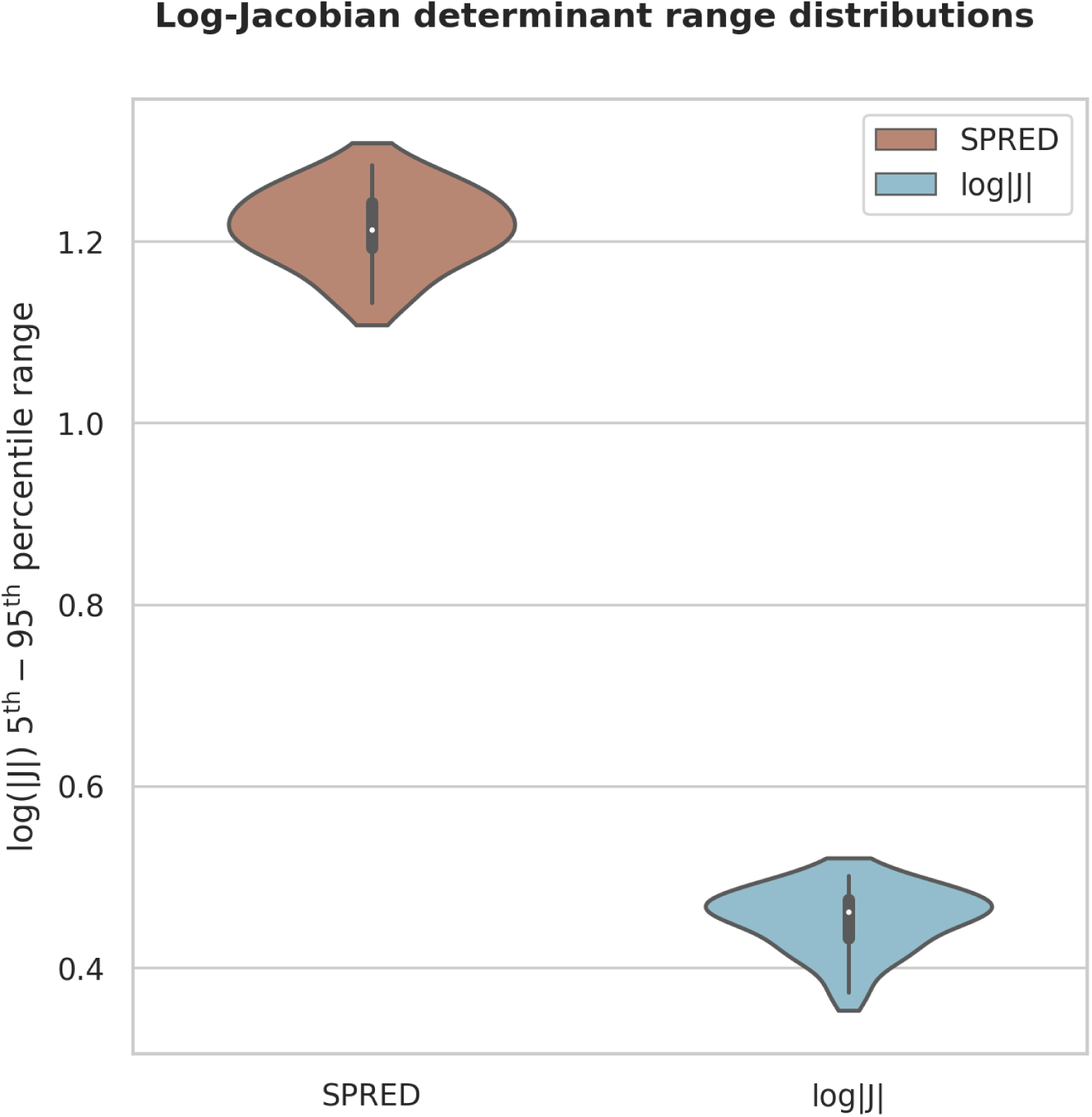
5^th^ to 95^th^ percentile range of Jacobian determinants for warps from all subjects to NA01 of the NIREP dataset. As would be expected penalising the log-Jacobian determinant directly leads to a much tighter distribution of Jacobian determinants about 1 than penalising the JSVs.

**Figure E.4:**
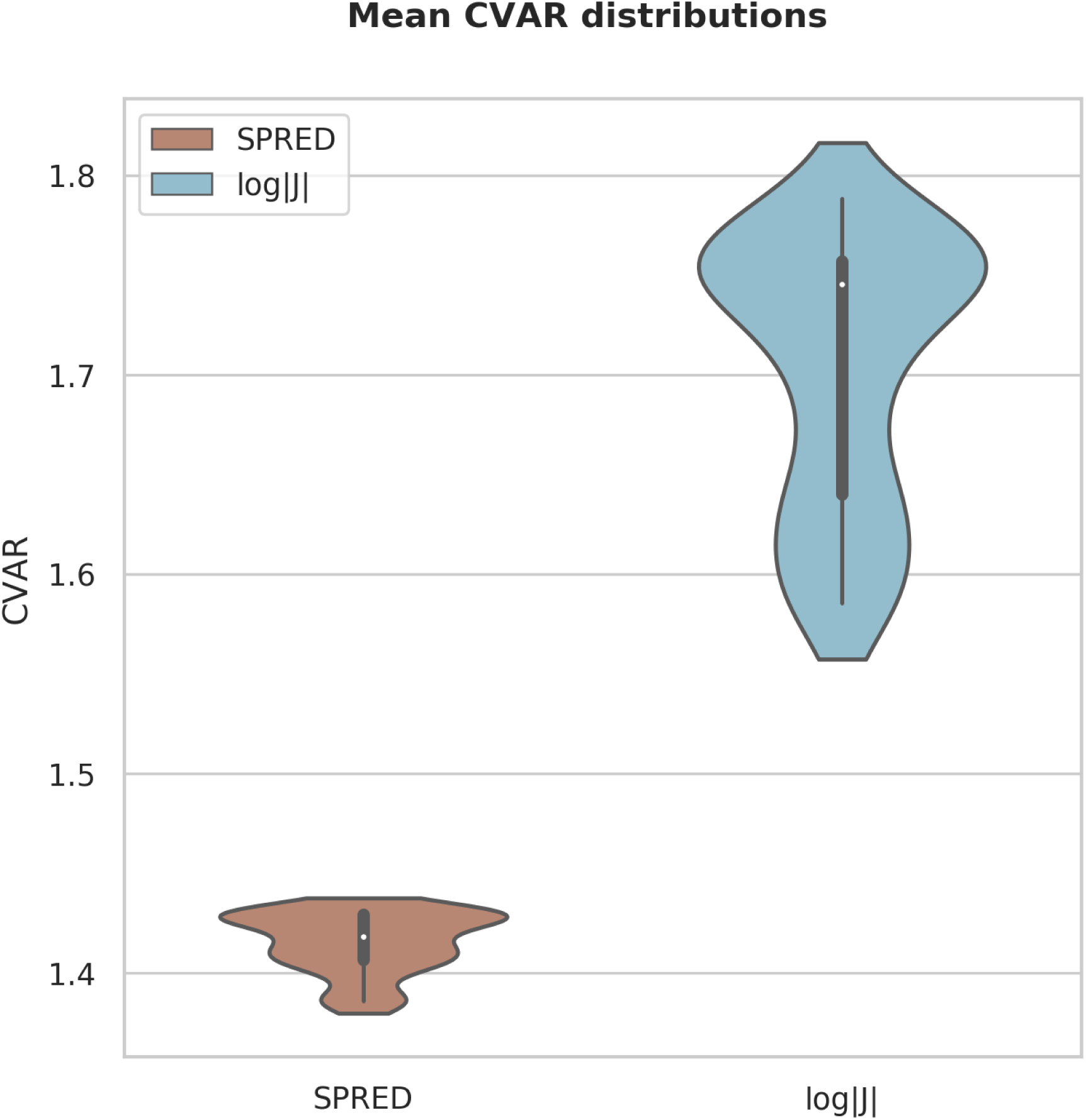
Distribution of mean CVAR within the brain for warps from all subjects to NA01 of the NIREP dataset. We observe that SPRED clearly better preserves the original shape of the underlying anatomy during deformation.

**Figure E.5:**
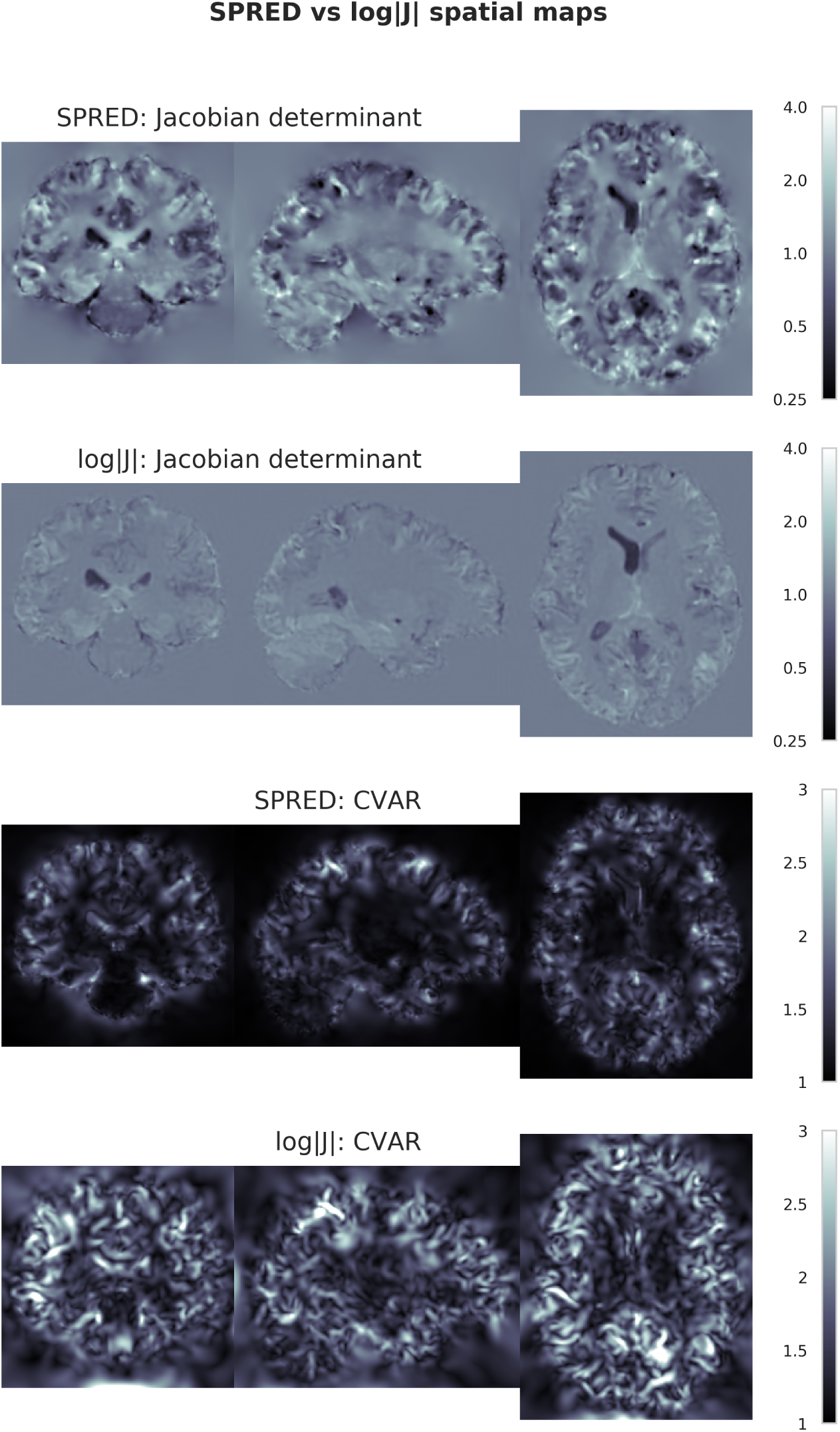
Spatial distribution of Jacobian determinants and CVAR for SPRED and log |**J**| regularisation. Penalising the volumetric changes directly clearly leads to flatter Jacobian determinant maps, however this is accompanied by a marked increase in CVAR confirming that this volumetric control comes at the cost of extreme shape changes.

**Figure E.6:**
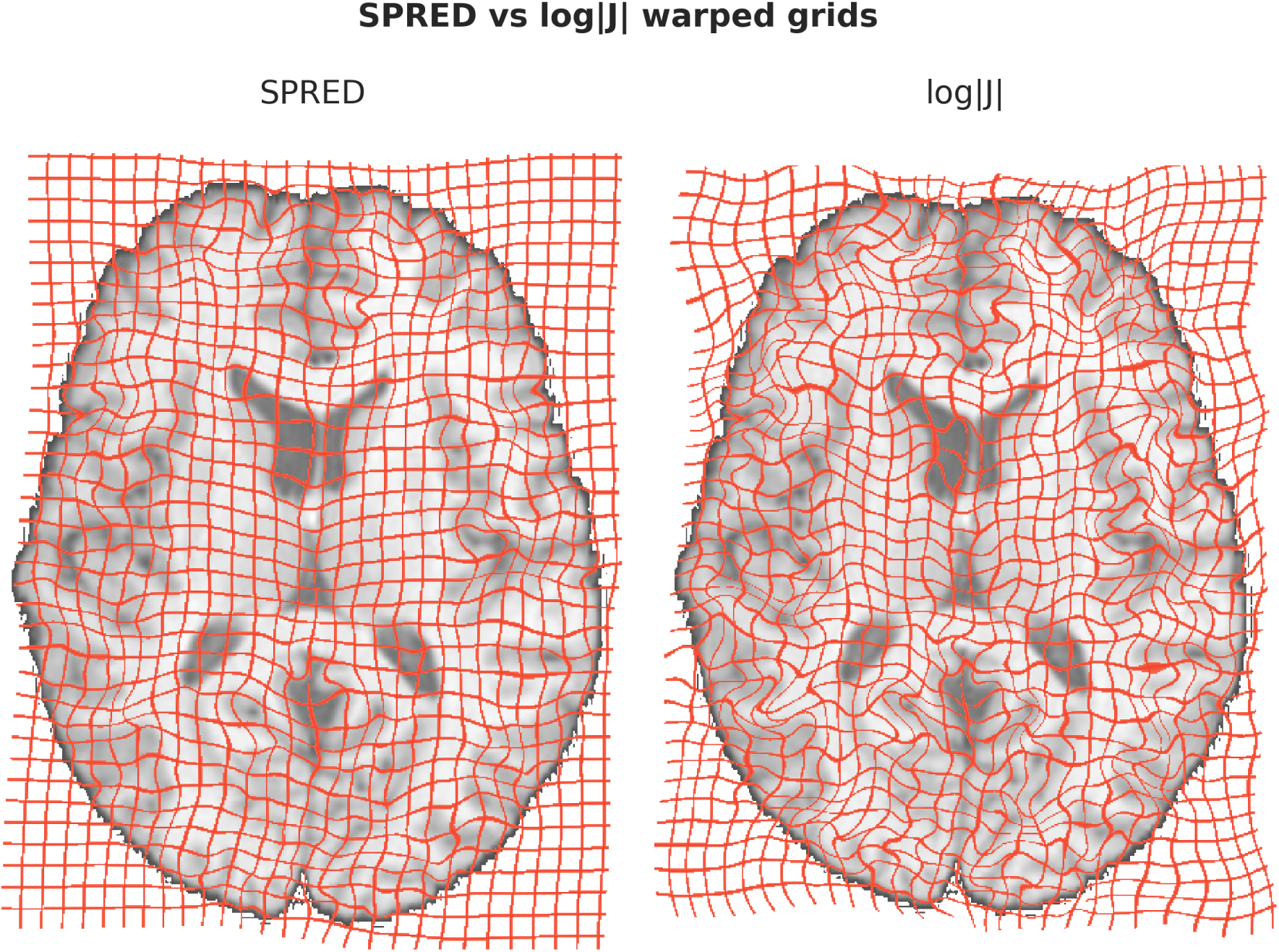
Warped grid showing the effect of SPRED and log |**J**| regularisation. Penalising the Jacobian determinant alone has led to extremely convoluted warps, whereas penalising the Jacobian singular values results in smoothly varying warps which are far more plausible.

## Notes

#### Summary of Updates

Differences between our method and those that penalise only the Jacobian determinant are clarified. Added comparison of shape distortion as measured by cube-volume aspect ratio. Add supplemental files covering code profiling and a general overview of the registration framework.

