## Supplemental Code Profiling for "A Symmetric Prior for the Regularisation of Elastic Deformations: Improved Anatomical Plausibility in Nonlinear Image Registration"

### A Symmetric Prior for the Regularisation of Elastic Deformations: Improved Anatomical Plausibility in Nonlinear Image Registration - *Supplementary Material: GPU Considerations and Code Profiling*

Frederik J Lange[a,*], John Ashburner[b], Stephen M Smith[a], Jesper L R Andersson[a]

[a] *Centre for Functional MRI of the Brain (FMRIB), Wellcome Centre for Integrative Neuroimaging, Nuffield Department of Clinical Neurosciences, University of Oxford, John Radcliffe Hospital, Headley Way, Oxford, OX3 9DU, UK*

[b] *Wellcome Centre for Human Neuroimaging, UCL Institute of Neurology, University College London, 12 Queen Square, London, WC1N 3BG, UK*

---

#### 1. Code Optimisation

In Section 2.4.1 of the main text we presented an approach to parallelising the calculations associated with incorporating the SPRED penalty into a Gauss-Newton optimisation framework, as well as providing an intuitive 2D example in Appendix B. However, there are a number of GPU-specific considerations that must be taken into account for the parallelisation to actually execute efficiently. Here we will demonstrate how we have addressed those considerations, as well as characterise the resulting code performance. It would benefit the reader to at least have a cursory familiarity with the CUDA C Programming Guide (NVIDIA, 2019) in order to understand the terminology used.

##### *1.1. GPU Specific Considerations*

We begin with some basic definitions:

*Corresponding Author

*Email addresses:* `` (Frederik J Lange), `` (John Ashburner), `` (Stephen M Smith), `` (Jesper L R Andersson)

**Host:** The CPU and its associated memory

**Device:** The GPU and its associated memory

**Kernel:** The code to be executed in parallel

**Thread:** The smallest logical execution unit of the device

**Block:** A group of threads

**Warp:** A set of 32 threads from the same block that are executed simultaneously on the device

**Latency:** The time taken to execute an instruction or read from memory

**Global Memory:** The largest but slowest memory space on the device, accessible to all threads

**Register Memory:** The fastest, most local memory, owned by a single thread

**Shared Memory:** Fast access memory shared between all threads within a block

**Texture Memory:** Fixed latency global memory cache that leverages 1D, 2D or 3D spatial locality of data

**Occupancy:** The ratio of the number of active warps to the maximum number of warps the device can handle

**Memory Coalescing:** Threads within a warp accessing consecutive location within memory

**Warp Divergence:** When threads within a warp do not follow the same branching pattern and must therefore have their execution serialised

**Warp Efficiency:** The average percentage of threads within a warp that are not serialised, and which are therefore simultaneously active

###### *1.1.1. Occupancy*

Occupancy is a critical factor in overall GPU code performance, as it directly determines the upper bound on how efficiently the GPU's compute elements are being utilised. A dominant factor in determining occupancy is the amount of register memory required by a particular kernel. If the kernel required less than 32 registers per thread, then theoretical occupancy will always be 100 %. In such cases, kernels can be executed executed with the maximum number of threads per block supported by a particular device (usually 1024).

Complications arise when register counts exceed 32 though. In such cases, executing a kernel with the maximum number of threads per block results in only a single block being able to execute on the device at any one time, as two blocks would require more registers than are physically present on the device, and all threads within a block must be loaded at once, or not at all. This dramatically reduces occupancy, and subsequently code performance. It is therefore better to reduce the number of threads per block thereby allowing maximal use of the available device resources.

Kernel register counts can be determined at compile time, and block sizes set explicitly to account for this, however hard-coding this may lead to brittle code. There is no guarantee that future versions of compilers and devices might not alter what the optimum configuration is. We therefore made use of the CUDA function `cudaOccupancyMaxPotentialBlockSize` to determine the best configuration at runtime.

Using this approach, all kernels have a maximum occupancy of between 62.5 % and 100 %, with achieve occupancies near the theoretical limits. Note that we did experiment with forcing the compiler to never exceed 32 registers per thread, however the increased latencies due to register spillage (i.e. using global memory as a swap space) nullified any gains in occupancy. Note also that our lowest occupancy kernel still had a factor of 300 speedup, and our most critical kernel in terms of execution times had a theoretical occupancy of 75 %.

###### *1.1.2. Memory Access*

Considerable time was spent on ensuring memory access was as optimised as possible, and we will therefore present some of the key design decisions that were found to improve performance.

The first consideration was memory coalescing when saving results of gradient and Hessian calculations. This is trivial when writing the result of gradient calculations if the parallelism is performed in a "thread per gradient element" regime and therefore this was an obvious approach to take. Calculating the Hessian is considerably more complicated problem though. As the Hessian is incredibly sparse, the choice of sparse matrix format will have a significant impact on the pattern of memory access. We selected a sparse diagonal format, where values along any non-zero diagonal of the Hessian are stored sequentially in memory. By assigning each diagonal to a block, we could then ensure that the threads within that block were always writing to sequential memory addresses resulting in near-perfectly coalesced memory write-access.

The second consideration was reading of the transformation basis functions. Many portions of the code require a representation of the cubic B-splines, or potentially a spatially differentiated version thereof. Calculating the gradient of the mean-squared error in a "thread per element" approach is equivalent to a strided convolution of the B-spline basis function with a spatially differentiated moving image. As such the B-spline need only be stored once per block as its values are identical irrespective of where the convolution is occurring. This is a perfect use case for shared memory, allowing the values of the spline to be read from global memory only once (in a perfectly coalesced manner) and then shared by all threads. Furthermore, the separable nature of B-splines was used to reduce memory usage by storing only 1D splines and then performing the multiplication on-the-fly. The same approach was used in the Hessian calculation, where the values along any diagonal can be similarly modelled as a strided convolution between the spatial overlap of two splines and a gradient image.

The final consideration was reading in the volumetric gradient images during the convolutions. Strided memory access immediately precludes coalescence unless the stride is accounted for pre-emptively in the way the data is stored. For example, for 3 sequential threads to access the true sequence:

$$\{1, 2, 3, 4, 5, 6, 7, 8, 9\}$$

in a coalesced manner with a stride of 3, it would need be stored in memory as:

$$\{1, 4, 7, 2, 5, 8, 3, 6, 9\}$$

Whilst this is doable, we chose to employ a more elegant approach using texture memory. Texture memory is a fixed latency cache that is optimised for spatially local data, and therefore well suited to the problem of strided convolutions The fixed latency means that for coalesced memory access texture memory would always be slower than global memory, however for non-coalesced memory access this is offset by the fact that the cache reduces the burden on global memory bandwidth (which would become the limiting factor in most of our kernels). Furthermore, by launching sufficient warps (*i.e.* when combined with high occupancy), latency can effectively be automatically hidden by allowing other warps to execute whilst one warp waits for data. We tested the effectiveness of this approach by replacing all texture memory reads with constant values (effectively eliminating the latency, but maintaining the number of computations) and saw minimal impact on the overall performance, indicating that the latency/bandwidth trade-off had the desired effect. Note that for some more recent GPU architectures, L1 caching may provide similar performance improvements for non-coalesced data access, however the method we have employed here will be effective across the vast majority of GPUs currently in use.

###### *1.1.3. Warp Efficiency*

Warp divergence is a critical aspect to minimise in order to attain high levels of computational throughput, and the decision to use a sparse diagonal matrix format for the Hessian benefits this. Consider the non-zero elements of one row of the Hessian. Each one can be mapped to a pattern of spatial overlap of the two B-splines corresponding to that point in the matrix. When traversing a row, the spatial overlap pattern changes for each non-zero element, and therefore would require a different pattern of data access. If threads within a warp were arranged per-row, then the control flow within each thread would differ, and the calculations would be serialised. In the worst case scenario, this could lead to an up to 32 times reduction in performance. However, it can be noticed that the pattern of spatial overlap along any diagonal of the Hessian is constant. Therefore, if the threads within a warp are assigned to a diagonal then they require exactly the same control flow to correctly index into the B-splines and warp divergence is effectively eliminated. By employing this strategy we achieve average warp efficiencies close to the theoretical optimum.

###### *1.1.4. Additional Considerations*

There are a number of smaller optimisations that have been made that we will now briefly describe for completeness.

**Image sampling and interpolation:** All image sampling and interpolation (including image gradients and warp sampling) is performed on the device using the `CubicInterpolationCuda` code by Ruijters and Thevenaz (2012).

**Data transfer:** The images being registered, the warp representation and the Hessian are all stored on the device reducing the need for expensive host-to-device and device-to-host data transfers.

**Linear Solver:** Gauss-Newton update steps are calculated in a parallelised manner on the device, and employ the CUSP (Dalton et al., 2014) iterative linear solver to invert the Hessian. Note that CUSP uses a compressed sparse row matrix representation, and we therefore implemented a kernel for converting to this format from our sparse diagonal Hessian.

##### *1.2. Profiling Results*

It may be of interest to quantify some aspects of our tool's performance through code profiling. To this end, we utilised the NVIDIA Visual Profiler (NVVP) to time the execution of kernels. CPU code was executed on an Intel Core i7-6700HQ processor with 16GB of RAM. GPU code was executed on an NVIDIA 960M GPU with 2GB of RAM.

We assessed both the overall proportion of time spent on each calculation (gradient, Hessian etc.) as well as the speedup in performance.

###### *1.2.1. Overall Performance*

As we have used a multiresolution registration approach, the proportion of time spent on each calculation is different depending on the current warp resolution. We present here the overall performance for isotropic knot spacings of 40, 30, 20 and 10 mm in Table 1. The key points to take away from these results are:

- The high GPU utilisation (particularly for higher resolution warps) suggests that the major computational bottlenecks have been effectively assigned to the GPU.
- The SPRED penalty (and particularly the Hessian calculation) is by far the most computationally intensive part of the algorithm.
- The algorithm is not being bottlenecked by excessive data transfers.

- Inverting the Hessian to determine the Gauss-Newton update step (via a GPU parallelised iterative linear solver) is not a computational bottleneck.

| Knot Spacing (mm) | Runtime (s) | GPU Utilisation (%) | Data Transfer (%) | SPRED Hessian (%) | SSD Hessian (%) | Gradient (%) | Sparse Matrix Convert (%) | Linear Solver (%) |
|---|---|---|---|---|---|---|---|---|
| 40 | 3.5 | 46.0 | 22.8 | 91.2 | 0.2 | 1.3 | 3.5 | 1.4 |
| 30 | 5.8 | 59.0 | 14.4 | 91.6 | 0.1 | 0.7 | 3.8 | 1.4 |
| 20 | 9.4 | 76.9 | 7.4 | 96.5 | 0.1 | 0.4 | 0.8 | 0.7 |
| 10 | 45.0 | 83.2 | 3.3 | 93.2 | 0.1 | 0.2 | 3.0 | 1.3 |

Table 1: Performance for 5 Gauss-Newton iterations of optimisation at varying isotropic warp resolutions (or knot spacings). Key points to note are that the GPU utilisation increases with warp resolution, whilst the proportion of time needed for data transfer decreases. This indicates that the GPU is being used most effectively when it is likely to have the biggest impact. Additionally calculating the Hessian of the SPRED penalty is by far the most computationally intensive portion of the algorithm. Iterative inversion of the Hessian (via a GPU parallelised iterative solver) accounts for less than 2 % of the GPU compute time. This rises to approximately 5 % of the compute time when factoring in a necessary conversion from our own internal sparse diagonal format to the solver's required compressed sparse row format. Note that the somewhat different 20 mm performance is a result of, entirely by chance, requiring fewer Levenberg iterations at that resolution, and consequently fewer sparse matrix conversions and linear solver iterations.

*1.2.2. Kernel Specific Performance*

Given an understanding of the relative importance of the GPU kernels within the entire registration, we examined the performance of selected kernels compared to reference CPU implementations. The CPU testing was coordinated in MATLAB, but utilises highly optimised compiled C code to perform the actual calculations. We have verified that the outputs of both methods are identical. Note that we did not implement the entire registration algorithm for the CPU, merely selected functions for testing purposes.

The first speedup we present in Figure 1 is the calculation of $\partial \mathbf{C}/\partial \mathbf{J}$. It is immediately evident that this calculation has benefited greatly from parallelisation, with over two orders of magnitude speedup. This is as expected, as the parallel per-voxel calculation is perfectly suited to the single instruction multiple data GPU parallelisation paradigm. Consequently, this particular calculation's contribution to the overall runtime has become negligible.

Secondly, we observe in Figure 2 that the overall gradient calculation (for both the similarity and regularisation metrics) has a speedup in the region of an order of magnitude. It should be noted that the overall performance is not overly dependent on this calculation in a Gauss-Newton framework.

Finally we consider the major computational bottleneck identified above, namely the calculation of the Hessian. Note that we have used two separate algorithms, one for the MSQ metric and one for the SPRED metric. The difference between the methods is due to the fact that we are able to exploit additional levels of symmetry in the MSQ Hessian that are not present in the SPRED Hessian. We observe in Figure 3 speedups in the region of $30 - 40\times$ for the MSQ Hessian calculation. Consequently the MSQ Hessian is not rate limiting to the registration.

There are a number of reasons why the SPRED Hessian is by far the most important calculation in terms of execution time. One, as seen in Figure 4, is that the speedup is about half that of the MSQ Hessian at around $15 - 20\times$ for practical combinations of knot spacing and coefficient dimensions. This is largely attributable to two reasons, namely:

- The MSQ Hessian has a theoretical 100 % GPU utilisation that is reduced to 75 % for the SPRED Hessian due to the second algorithm requiring more registers per thread.
- The SPRED Hessian requires more memory reads per element of the Hessian, which introduces additional latency costs on the GPU.

Secondly, the SPRED Hessian calculation is called $9\times$ more often than the MSQ Hessian, due to the cross-terms introduced by ${}^{\partial \mathbf{C}}/_{\partial \mathbf{J}}$.

Finally, the reduced symmetry introduces $8\times$ more calculations per SPRED Hessian. In future, we will focus on increasing the speedup of the SPRED Hessian further, potentially by deconstructing the algorithm into multiple smaller computational units that would be better able to hide the inherent latencies in the calculations.

Overall the GPU implementations in MMORF achieve their goal of targeting the bottleneck calculations of the algorithm, leading to a method that is computationally tractable.

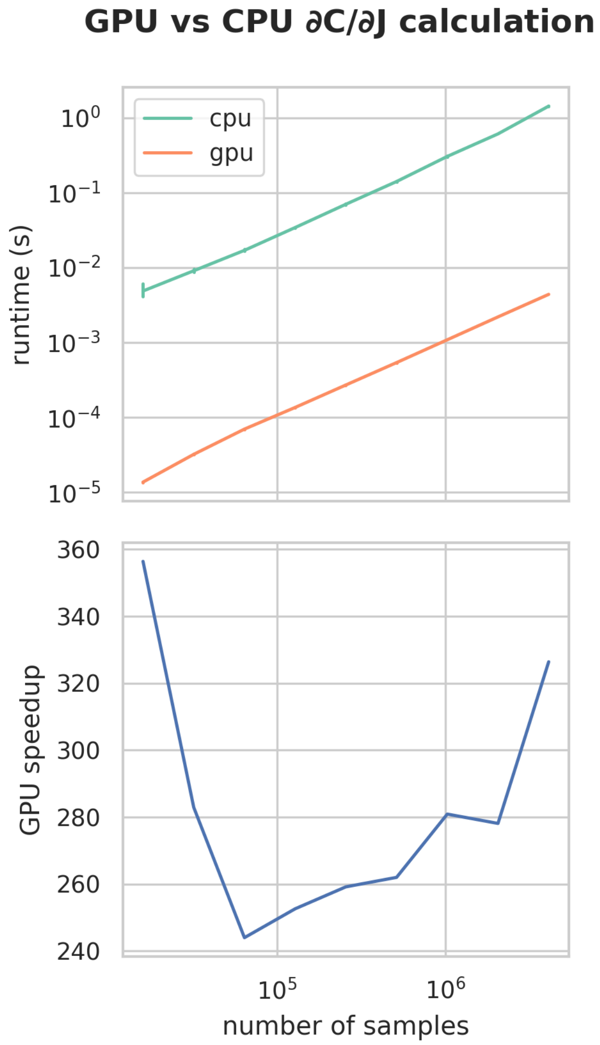


Figure 1: Runtime and speedup of GPU vs CPU implementations of the $\partial\mathbf{C}/\partial\mathbf{J}$ calculation that forms part of both the gradient and Hessian calculations.

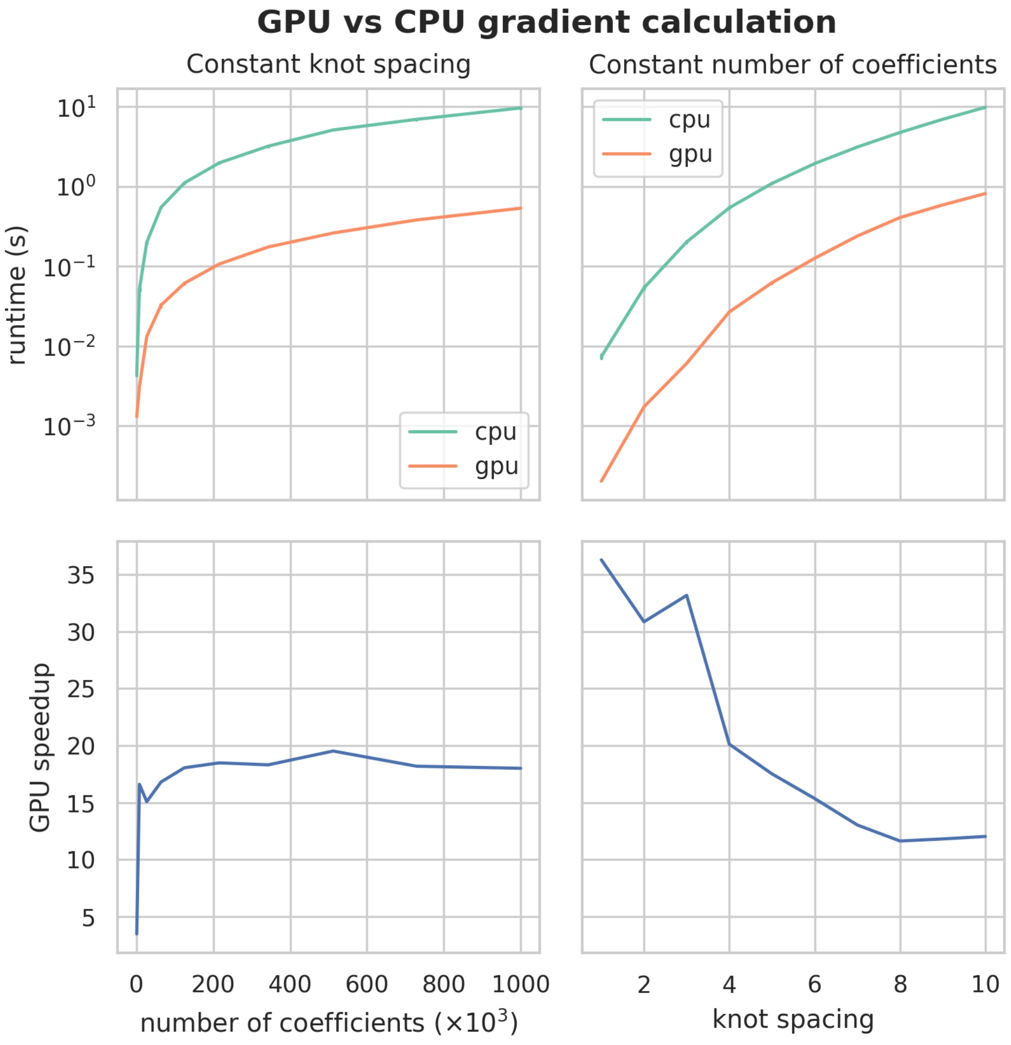


Figure 2: Runtime and speedup of GPU vs CPU implementations of the gradient calculation with respect to the B-spline warp parameters. The left column shows results when knot spacing is kept constant at 5 mm isotropic, and the number of spline coefficients in each dimension are varied. The right column shows results when the number of spline coefficients in each dimension is held constant at 50 isotropic, and the knot spacing is varied. In all cases the images are samples at 1 mm isotropic.

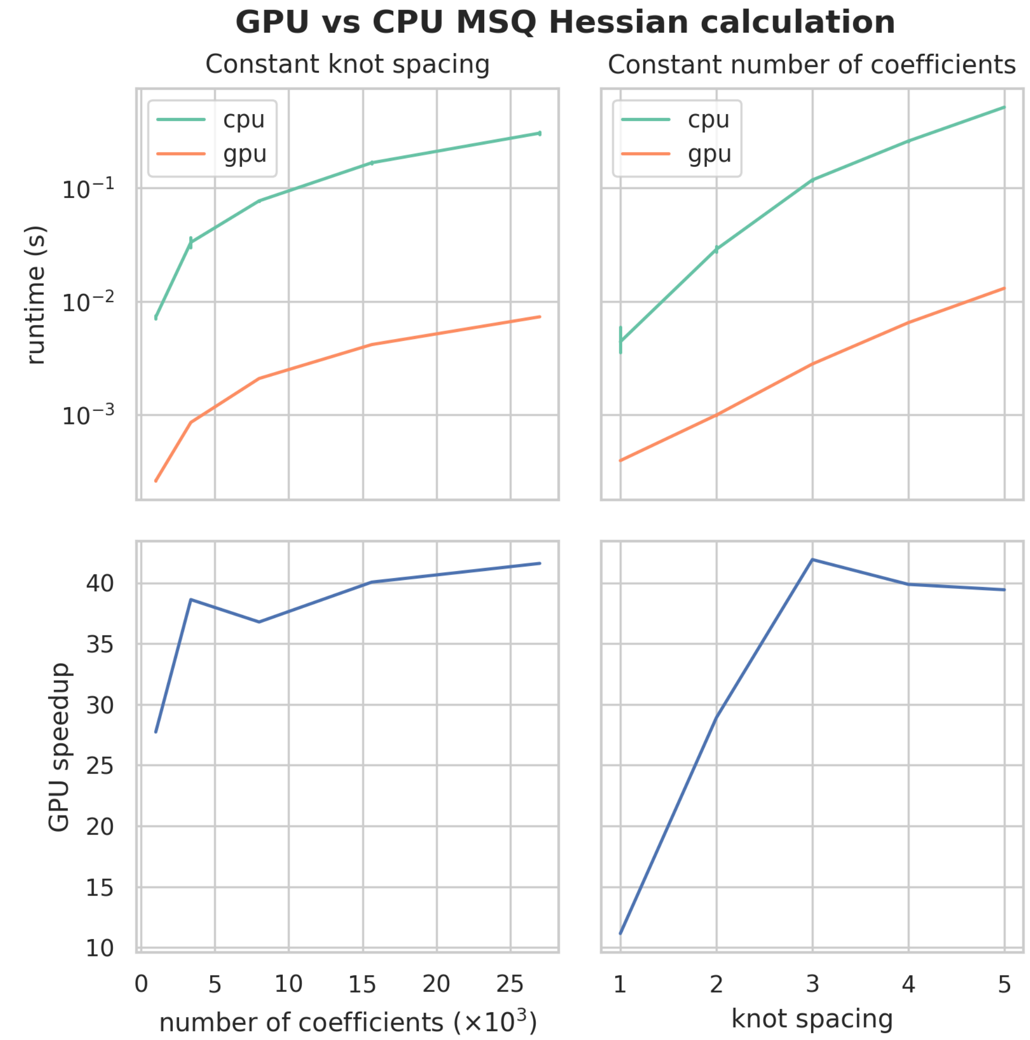


Figure 3: Runtime and speedup of GPU vs CPU implementations of the Hessian calculation for the MSQ metric. The left column shows results when knot spacing is kept constant at 2 mm isotropic and the number of spline coefficients in each dimension are varied. The right column shows results when the number of spline coefficients in each dimension is held constant ant 15 isotropic and the knot spacing is varied. In all cases the images are samples at 1 mm isotropic.

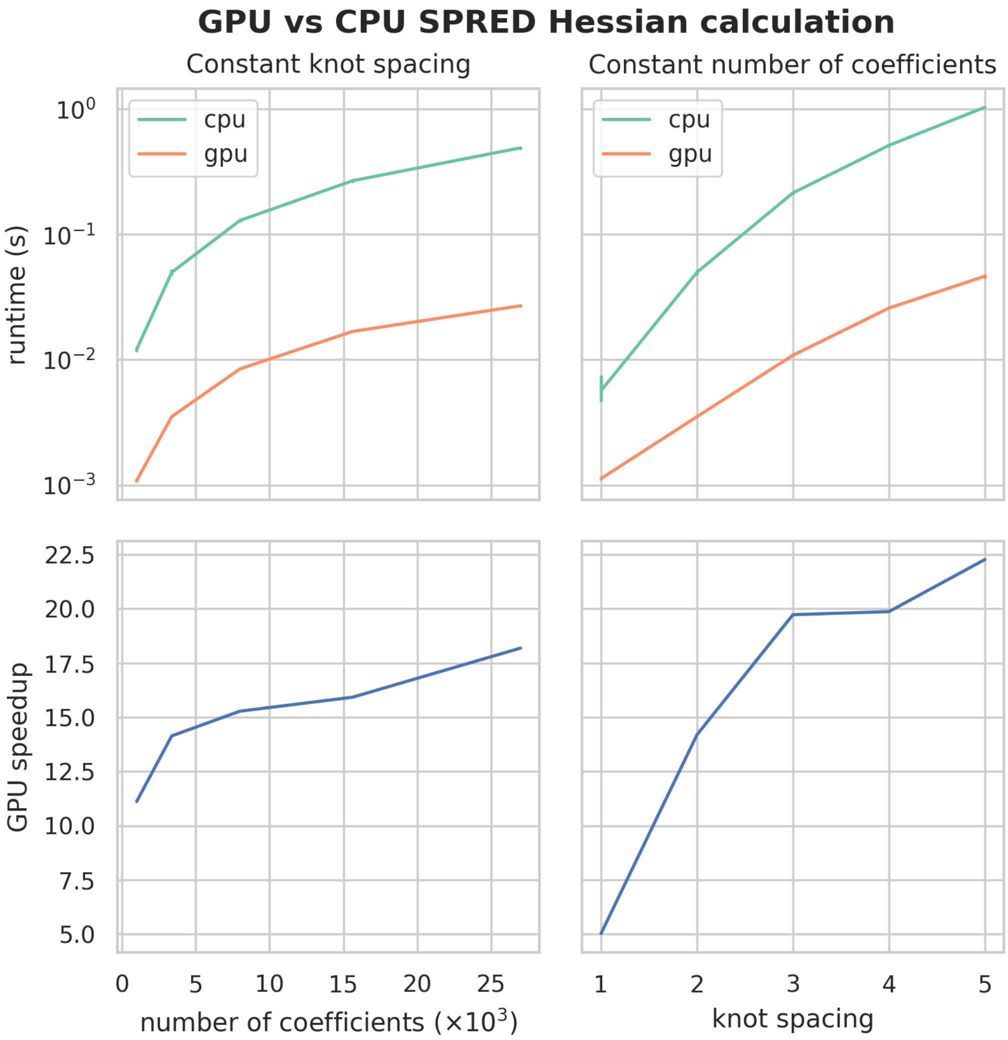


Figure 4: Runtime and speedup of GPU vs CPU implementations of the Hessian calculation for the SPRED regularisation metric. The left column shows results when knot spacing is kept constant at 2 mm isotropic and the number of spline coefficients in each dimension are varied. The right column shows results when the number of spline coefficients in each dimension is held constant ant 15 isotropic and the knot spacing is varied. In all cases the images are samples at 1 mm isotropic.
