## Supplemental Registration Framework for "A Symmetric Prior for the Regularisation of Elastic Deformations: Improved Anatomical Plausibility in Nonlinear Image Registration"

### 1. Parametrisation of the Transform

The defining characteristic of small-deformation registration frameworks is that they solve for the transform in terms of a displacement field directly, rather than a velocity field which is then integrated (or exponentiated) to form the displacement field. As stated in Section 2.3.3 of the main text, we employ a cubic B-spline free-form deformation to parametrise the transform.

Cubic B-splines possess a number of desirable characteristics, such as:

- C2 continuity

  *i.e.*, continuous, analytic derivatives up to second order

- Compact support

  *i.e.*, each spline has only a finite non-zero extent

*Corresponding Author

*Email addresses:* `` (Frederik J Lange), `` (John Ashburner), `` (Stephen M Smith), `` (Jesper L R Andersson)

- Spatial separability

  *i.e.*, an $n$-dimensional spline (and its spatial derivatives) can be decomposed into the product of $n$ 1-dimensional splines (or differentiated splines)

By treating the volumes to be registered as continuous functions on $\mathbb{R}^3$, we define the continuous case of the transformation in the following manner:

$$\mathbf{f}\left(x, y, z\right) = \text{Reference volume} \tag{1}$$

$$\mathbf{g}\left(x, y, z\right) = \text{Moving volume} \tag{2}$$

$$M = \text{Number of splines per warp direction} \tag{3}$$

$$\vec{w}_x/\vec{w}_x/\vec{w}_z = x,\ y \text{ and } z \text{ direction warp parameters} \tag{4}$$

$$\vec{w} = \underbrace{\begin{bmatrix}\vec{w}_x \\ \vec{w}_y \\ \vec{w}_z\end{bmatrix}}_{3M\times 1} \tag{5}$$

$$d_{x/y/z} = x,\ y \text{ and } z \text{ displacements} \tag{6}$$

$$\vec{\mathbf{t}}\left(x, y, z, \vec{w}\right) = \text{Coordinate transformation/warp} \tag{7}$$

$$= \begin{bmatrix}x & y & z\end{bmatrix} + \begin{bmatrix}d_x & d_y & d_z\end{bmatrix} \tag{8}$$

$$\mathbf{g}\left(\vec{\mathbf{t}}\left(x, y, z, \vec{w}\right)\right) = \text{Transformed moving volume} \tag{9}$$

For simplicity, we use the same set of B-splines to define the warps in all 3 directions (*i.e.*, the number and spatial extent of splines defining the displacements in each direction is the same, and they are located identically in space). $M$ is chosen to be all splines whose spatial support overlaps with the domain over which $\mathbf{f}$ is defined. The warp parameters $\vec{w}$ are then the coefficients of each B-spline, and each parameter only affects displacement in a single direction.

In practice though, we only ever deal with discrete samples of our continuous images. We can therefore describe the process of transforming a set of sample coordinates in $\mathbf{f}$ to their corresponding coordinates in $\mathbf{g}$ as follows:

$$
\begin{aligned}
N &= \text{Number of sampled voxels in } \mathbf{f} && (10)\\
\vec{x}/\vec{y}/\vec{z} &= x,\ y \text{ and } z \text{ coordinates of samples in } \mathbf{f} && (11)\\
\mathbf{X} &= \underbrace{\begin{bmatrix}\vec{x} & \vec{y} & \vec{z}\end{bmatrix}}_{N\times 3} && (12)\\
&= \begin{bmatrix} x_0 & y_0 & z_0\\ x_1 & y_1 & z_1\\ \vdots & \vdots & \vdots\\ x_N & y_N & z_N \end{bmatrix} && (13)\\
\vec{b}_m &= \text{Vectorised } m^{\text{th}} \text{ B-spline at sample positions } \mathbf{X} && (14)\\
\mathbf{B} &= \underbrace{\begin{bmatrix}\vec{b}_0 & \vec{b}_1 & \cdots & \vec{b}_M\end{bmatrix}}_{N\times M} && (15)\\
&= \underbrace{\left.\begin{bmatrix} b_{00} & b_{01} & \cdots & b_{0M}\\ b_{10} & \ddots & & \vdots\\ \vdots & & \ddots & \vdots\\ b_{N0} & \cdots & \cdots & b_{NM} \end{bmatrix}\right\}}_{\text{1 B-spline per column}} \text{1 Voxel per row} && (16)\\
\mathbf{W} &= \underbrace{\begin{bmatrix}\vec{w}_x & \vec{w}_y & \vec{w}_z\end{bmatrix}}_{M\times 3} && (17)\\
\mathbf{T}\left(\mathbf{X}, \vec{w}\right) &= \text{Transformed sample coordinates} && (18)\\
&= \mathbf{X} + \mathbf{BW} && (19)
\end{aligned}
$$

The compact support of B-splines means that **B** can be very sparse. Additionally, (disregarding edge cases) each column of **B** is simply a shifted version of any other column. We therefore never store **B** explicitly, and instead compute **BW** using simple convolution.

### 2. Cost Function

Registration can be seen as an optimisation problem, and we therefore require an objective function to optimise. We have chosen to formulate this as the minimisation of a cost function. Our cost function combines a dissimilarity metric and a regularisation penalty. The regularisation is the topic of the main text, but we present here the formal definition of our mean-squares (MSQ) dissimilarity metric, its gradient and Gauss-Newton Hessian:

$$
\begin{aligned}
C_{MSQ} &= \frac{1}{N}\sum_{n=1}^{N}\left(\mathbf{f}(x_n, y_n, z_n) - \mathbf{g}\left(\vec{t}(x_n, y_n, z_n, \vec{w})\right)\right)^2 && (20)\\
&= \frac{1}{N}\left[\vec{f}\left(\mathbf{X}\right) - \vec{g}\left(\mathbf{T}\left(\mathbf{X}, \vec{w}\right)\right)\right]^{\mathrm{T}}\left[\vec{f}\left(\mathbf{X}\right) - \vec{g}\left(\mathbf{T}\left(\mathbf{X}, \vec{w}\right)\right)\right] && (21)\\
&= \frac{1}{N}\left[\vec{f} - \vec{g}\left(\vec{w}\right)\right]^{\mathrm{T}}\left[\vec{f} - \vec{g}\left(\vec{w}\right)\right] && (22)\\
&= \frac{1}{N}\vec{e}\left(\vec{w}\right)^{\mathrm{T}}\vec{e}\left(\vec{w}\right) && (23)
\end{aligned}
$$

$$
\begin{aligned}
\nabla_{MSQ} &= -\frac{2}{N}\left[\frac{\partial\vec{g}\left(\vec{w}\right)}{\partial\vec{w}}\right]^{\mathrm{T}}\left[\vec{f} - \vec{g}\left(\vec{w}\right)\right] && (24)\\
&= -\frac{2}{N}\mathbf{J}\left(\vec{w}\right)^{\mathrm{T}}\vec{e}\left(\vec{w}\right) && (25)
\end{aligned}
$$

$$
\begin{aligned}
\mathbf{H}_{MSQ} &= \frac{2}{N}\left[\frac{\partial\vec{g}\left(\vec{w}\right)}{\partial\vec{w}}\right]^{\mathrm{T}}\left[\frac{\partial\vec{g}\left(\vec{w}\right)}{\partial\vec{w}}\right] && (26)\\
&= \frac{2}{N}\mathbf{J}\left(\vec{w}\right)^{\mathrm{T}}\mathbf{J}\left(\vec{w}\right) && (27)
\end{aligned}
$$

### 3. Optimisation

Optimisation is based on Levenberg's extension to the iterative Gauss-Newton method. Update steps for Newton-based methods can be derived from the second order Taylor expansion of a function $C\left(\vec{w}\right)$ as follows:

$$
\begin{aligned}
Q\left(\vec{w}_{n+1}\right) &= Q\left(\vec{w}_n + \Delta\vec{w}\right) && (28)\\
&= C\left(\vec{w}_n\right) + \nabla_C\left(\vec{w}_n\right)\cdot\Delta\vec{w} + \Delta\vec{w}^{\mathrm{T}}\mathbf{H}_C\left(\vec{w}_n\right)\Delta\vec{w} && (29)
\end{aligned}
$$

We note that:

$$
\begin{aligned}
&\min_{\Delta\vec{w}} Q\left(\vec{w}_n + \Delta\vec{w}\right) && (30)\\
&\implies \nabla_Q\left(\vec{w}_n + \Delta\vec{w}\right) = 0 && (31)
\end{aligned}
$$

And therefore:

$$\nabla_C(\vec{w}_n) + \mathbf{H}_C(\vec{w}_n)\,\Delta\vec{w} = 0 \tag{32}$$

$$\Delta\vec{w} = -\mathbf{H}_C^{-1}(\vec{w}_n)\,\nabla_C(\vec{w}_n) \tag{33}$$

The Gauss-Newton update step is not guaranteed to reduce the cost function at every iteration, and we therefore employ Levenberg's extension in which the Hessian is replaced by:

$$\mathbf{H}_{Levenberg} = \mathbf{H}_{Gauss-Newton} + \lambda\mathbf{I} \tag{34}$$

### 4. Linear Solver

Newton-based methods require inverting the Hessian to calculate the update step. For problems with many parameters, actually inverting the Hessian is infeasible, and we instead use an iterative conjugate gradient method to solve the system of linear equations defined by:

$$\mathbf{H}\Delta\vec{w} = -\nabla \tag{35}$$

In order to speed up the optimisation we use the Jacobi preconditioner:

$$\mathbf{P} = \mathrm{diag}(\mathbf{H}) \tag{36}$$

to solve the left-preconditioned system:

$$\mathbf{P}^{-1}\mathbf{H}\Delta\vec{w} = -\mathbf{P}^{-1}\nabla \tag{37}$$

This method is effective due to the diagonal dominance of $\mathbf{H}$.
