## Supplemental Jacobian Determinant Comparison for "A Symmetric Prior for the Regularisation of Elastic Deformations: Improved Anatomical Plausibility in Nonlinear Image Registration"

### A Symmetric Prior for the Regularisation of Elastic Deformations: Improved Anatomical Plausibility in Nonlinear Image Registration - *Supplementary Material: Comparison of Log-Jacobian Determinant Spatial Maps for all Subject Pairs in the NIREP Dataset*

Frederik J Lange, Stephen M Smith, John Ashburner, Jesper L R Andersson

---

---

#### 1. Reference Subject 01

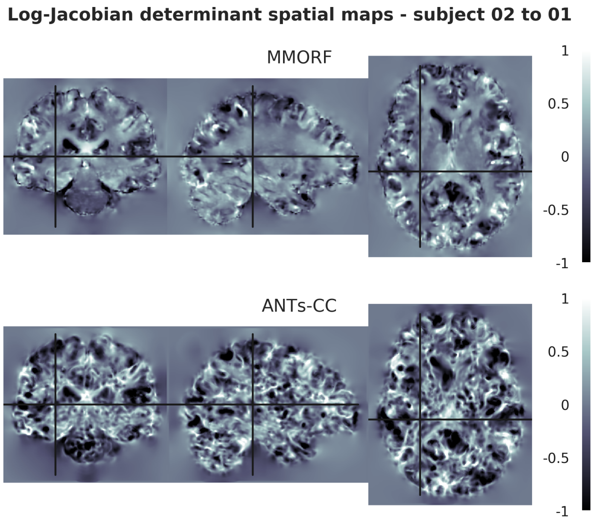


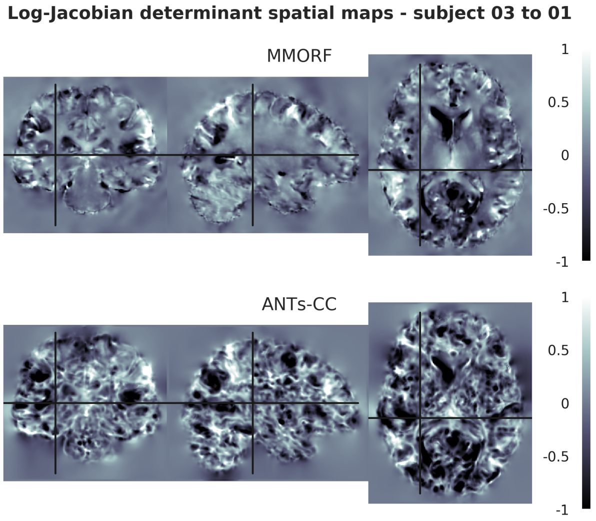


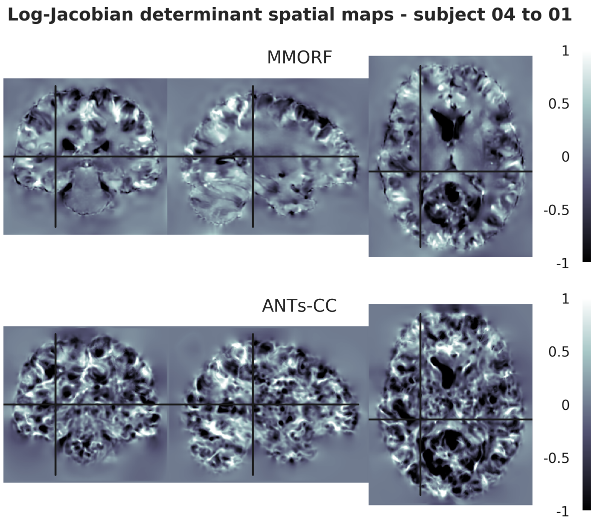


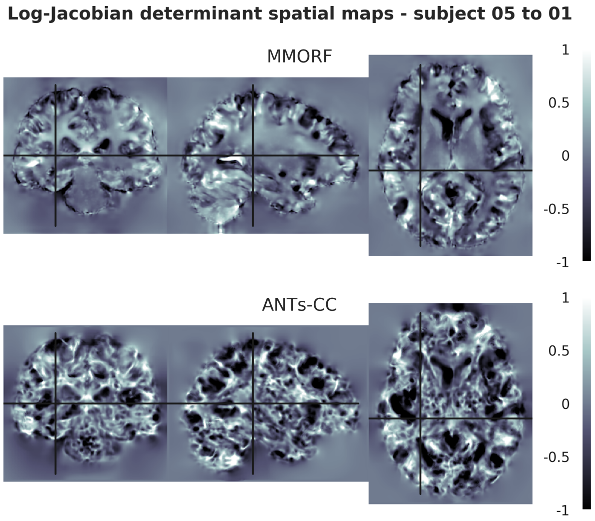

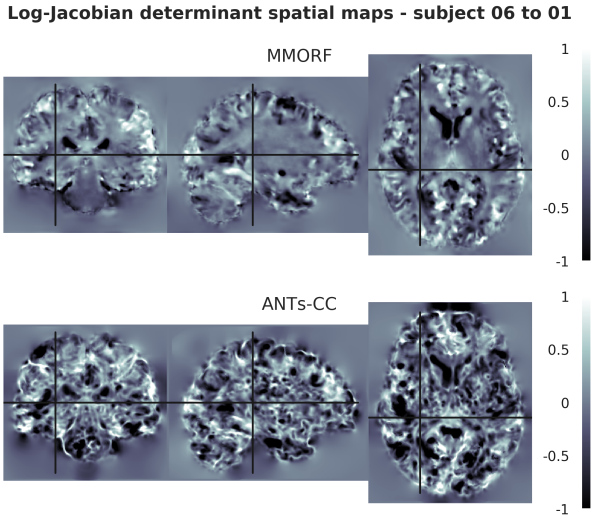
Log-Jacobian determinant spatial maps - subject 06 to 01
MMORF
1
0.5
0
-0.5
-1
ANTs-CC
1
0.5
0
-0.5
-1

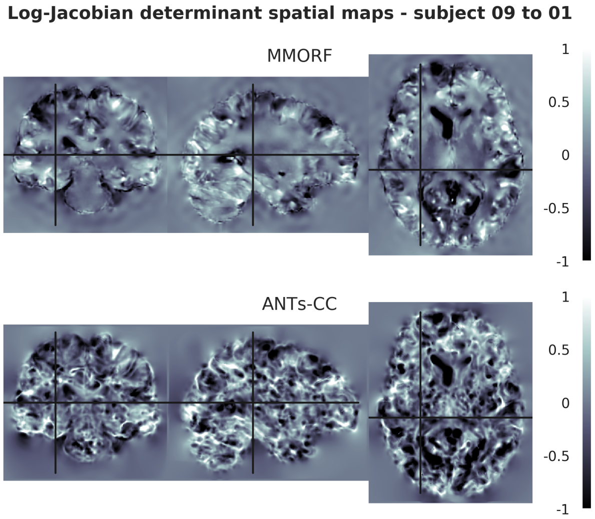
Log-Jacobian determinant spatial maps - subject 09 to 01
MMORF
1
0.5
0
-0.5
-1
ANTs-CC
1
0.5
0
-0.5
-1

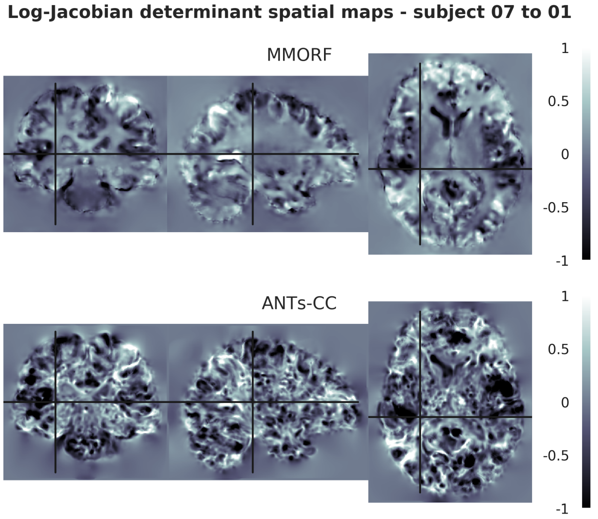
Log-Jacobian determinant spatial maps - subject 07 to 01
MMORF
1
0.5
0
-0.5
-1
ANTs-CC
1
0.5
0
-0.5
-1

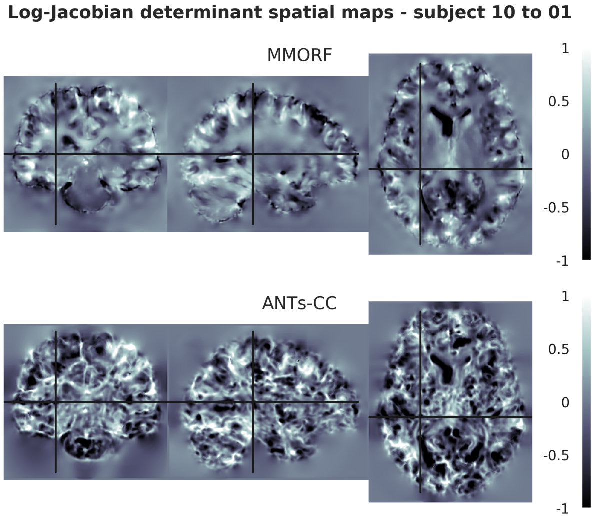
Log-Jacobian determinant spatial maps - subject 10 to 01
MMORF
1
0.5
0
-0.5
-1
ANTs-CC
1
0.5
0
-0.5
-1

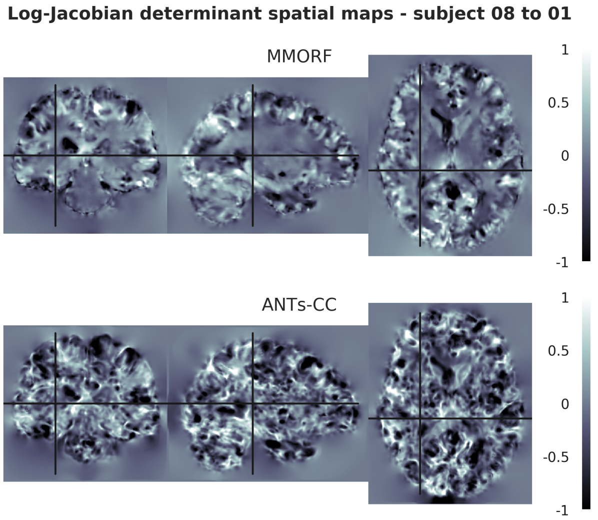
Log-Jacobian determinant spatial maps - subject 08 to 01
MMORF
1
0.5
0
-0.5
-1
ANTs-CC
1
0.5
0
-0.5
-1

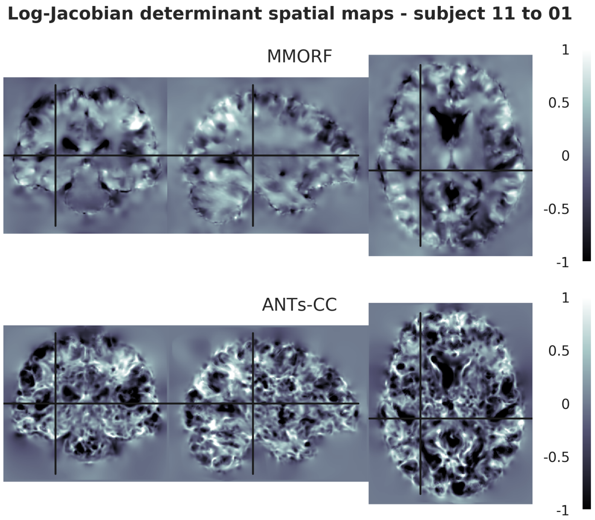
Log-Jacobian determinant spatial maps - subject 11 to 01
MMORF
1
0.5
0
-0.5
-1
ANTs-CC
1
0.5
0
-0.5
-1

**Log-Jacobian determinant spatial maps - subject 12 to 01**

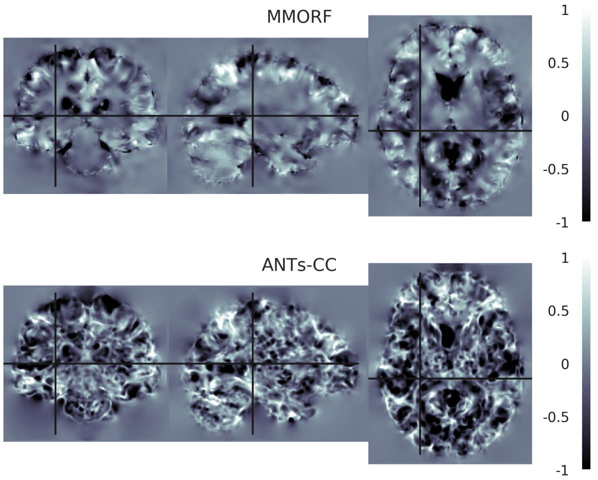


**Log-Jacobian determinant spatial maps - subject 13 to 01**

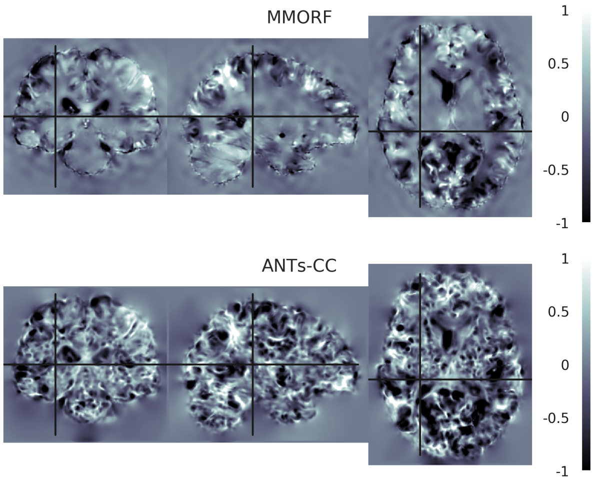


**Log-Jacobian determinant spatial maps - subject 14 to 01**

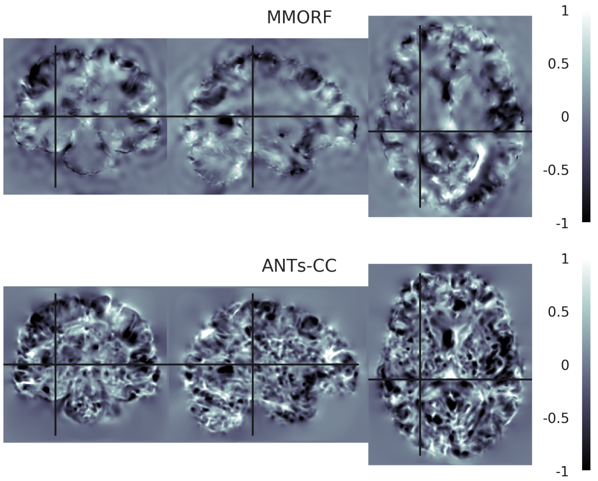


**Log-Jacobian determinant spatial maps - subject 15 to 01**

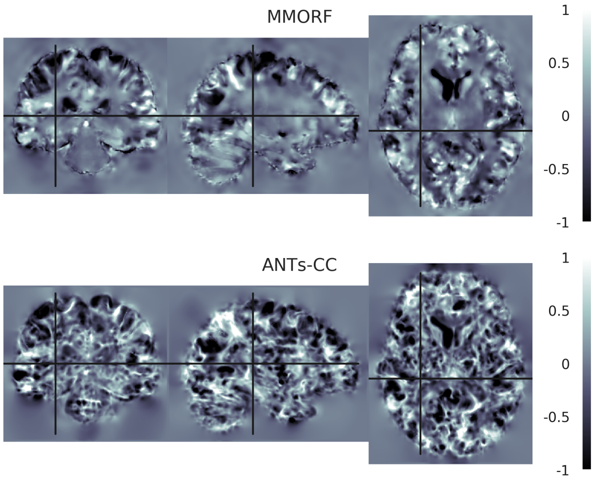


**Log-Jacobian determinant spatial maps - subject 16 to 01**

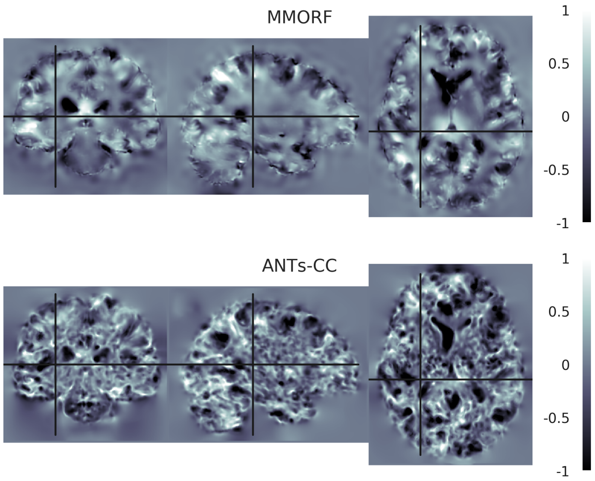

#### 2. Reference Subject 02

**Log-Jacobian determinant spatial maps - subject 01 to 02**

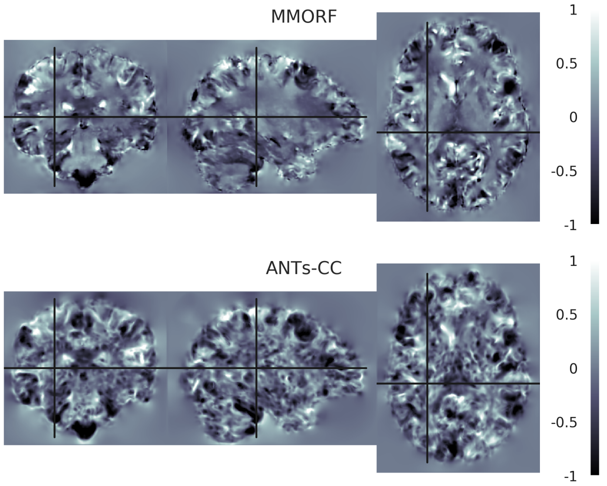


**Log-Jacobian determinant spatial maps - subject 03 to 02**

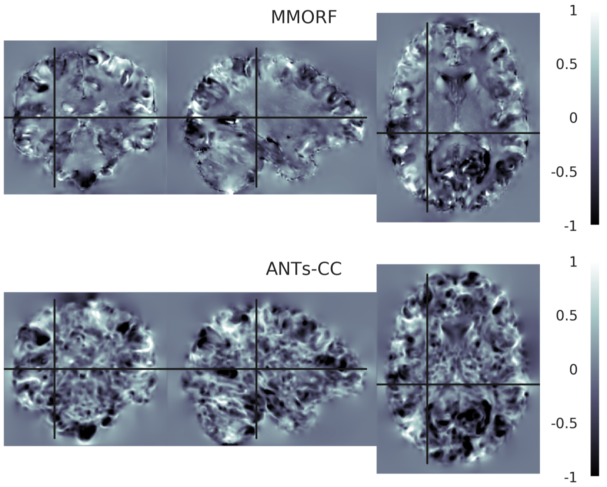


**Log-Jacobian determinant spatial maps - subject 04 to 02**

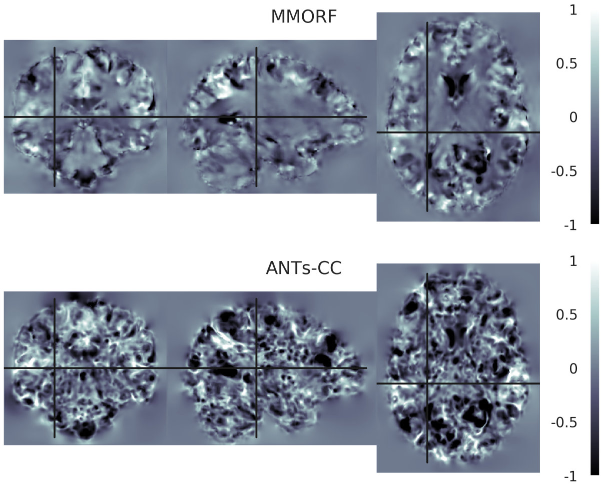


**Log-Jacobian determinant spatial maps - subject 05 to 02**

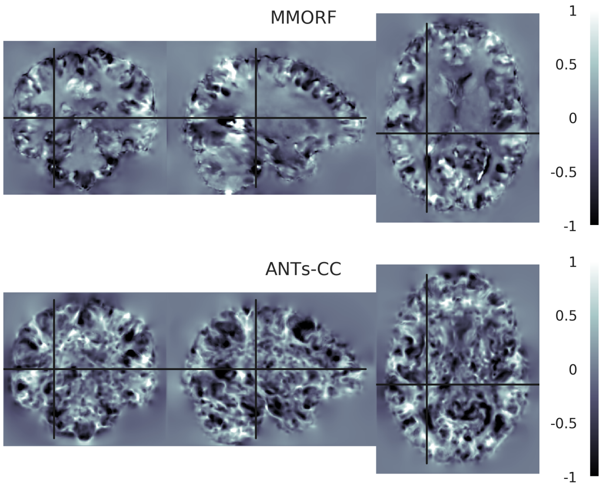


**Log-Jacobian determinant spatial maps - subject 06 to 02**

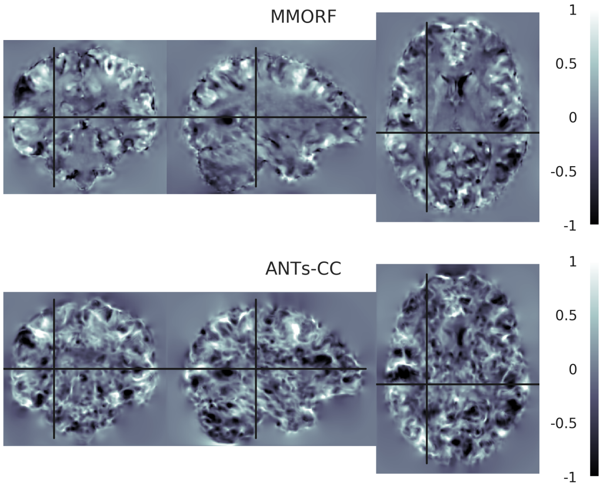


**Log-Jacobian determinant spatial maps - subject 07 to 02**

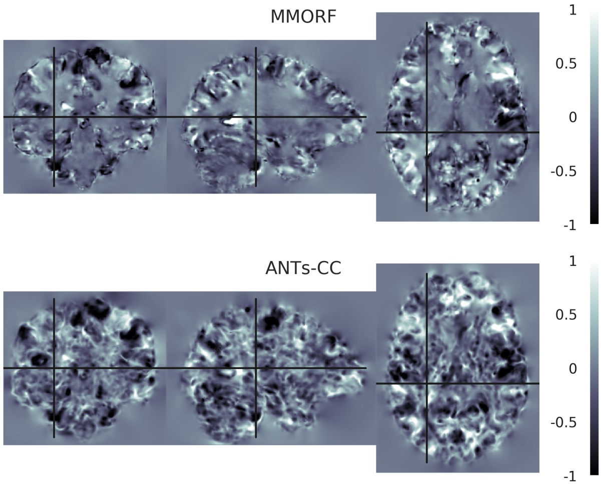

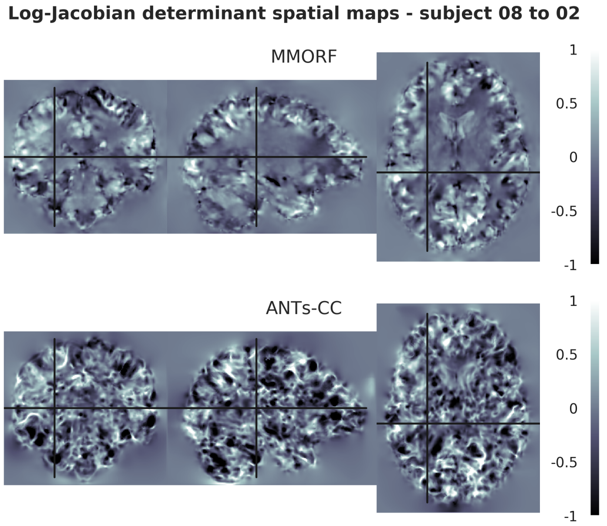
Log-Jacobian determinant spatial maps - subject 08 to 02
MMORF
ANTs-CC
1
0.5
0
-0.5
-1

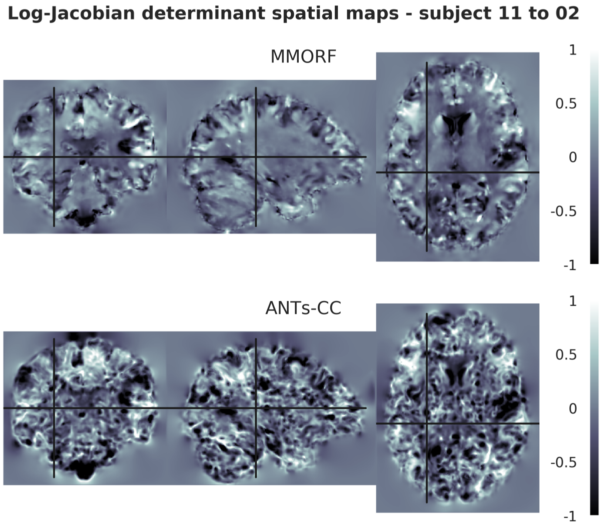
Log-Jacobian determinant spatial maps - subject 11 to 02
MMORF
ANTs-CC
1
0.5
0
-0.5
-1

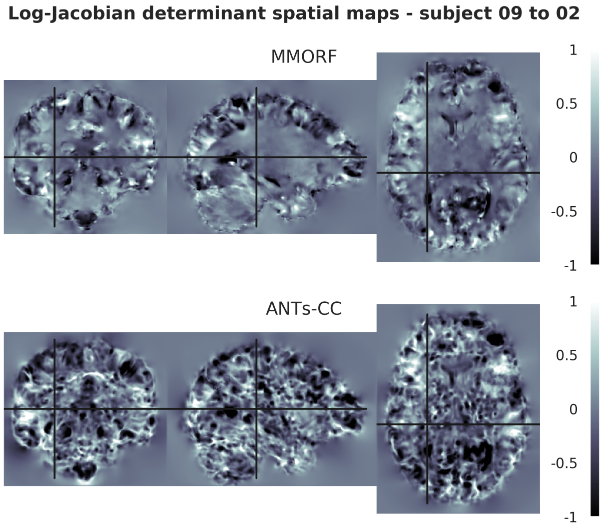
Log-Jacobian determinant spatial maps - subject 09 to 02
MMORF
ANTs-CC
1
0.5
0
-0.5
-1

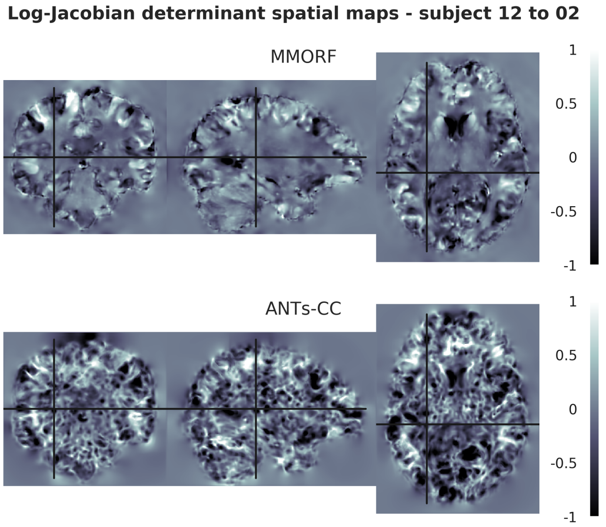
Log-Jacobian determinant spatial maps - subject 12 to 02
MMORF
ANTs-CC
1
0.5
0
-0.5
-1

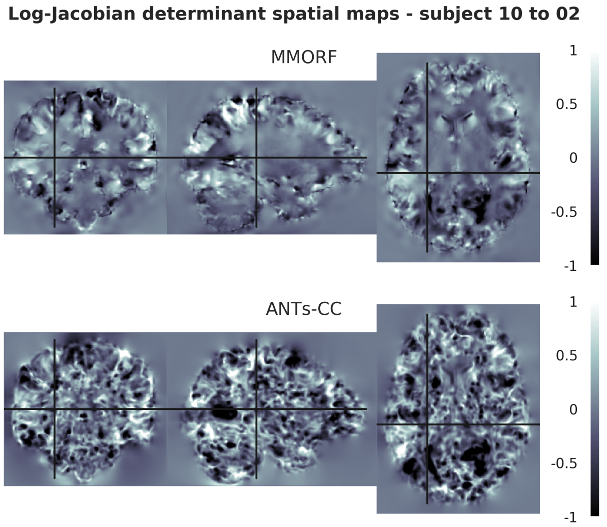
Log-Jacobian determinant spatial maps - subject 10 to 02
MMORF
ANTs-CC
1
0.5
0
-0.5
-1

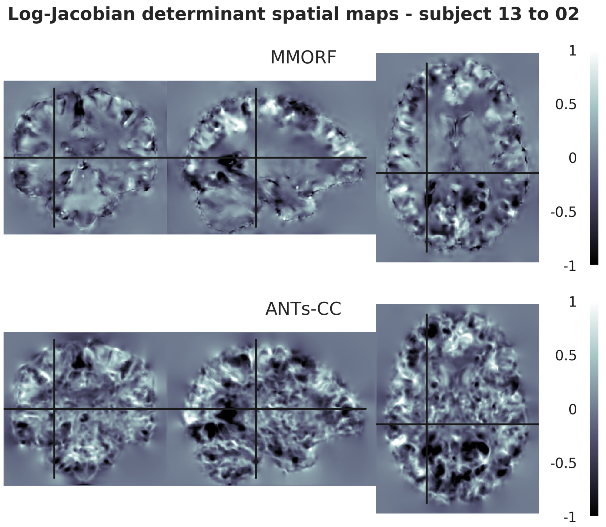
Log-Jacobian determinant spatial maps - subject 13 to 02
MMORF
ANTs-CC
1
0.5
0
-0.5
-1

**Log-Jacobian determinant spatial maps - subject 14 to 02**

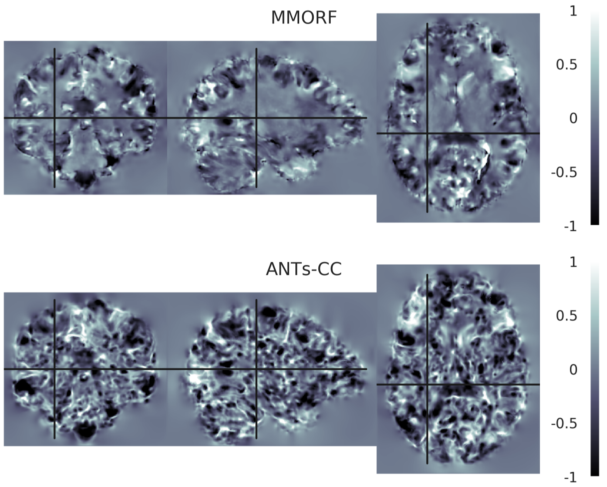


**Log-Jacobian determinant spatial maps - subject 15 to 02**

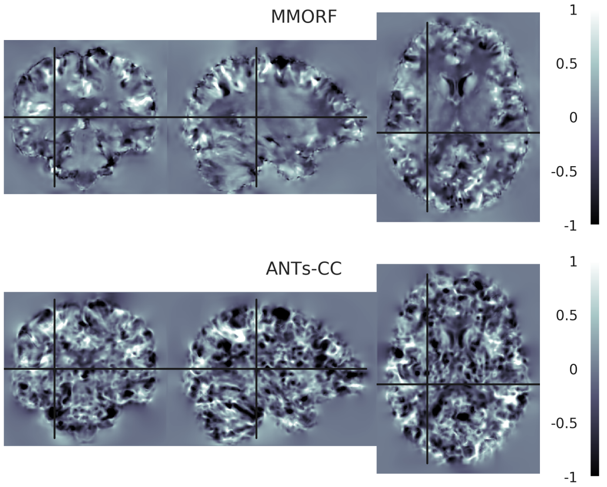


**Log-Jacobian determinant spatial maps - subject 16 to 02**

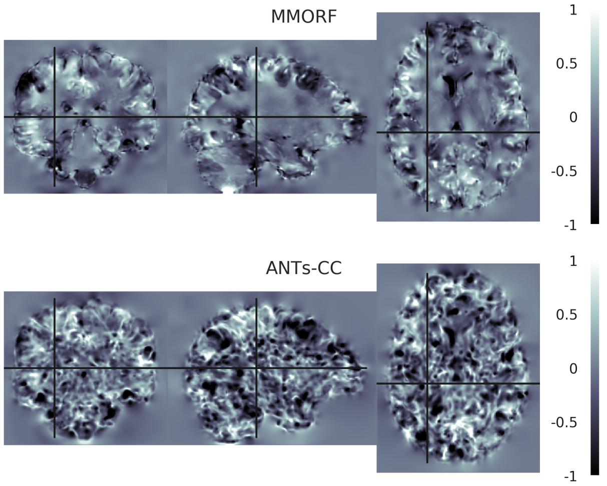

#### 3. Reference Subject 03

**Log-Jacobian determinant spatial maps - subject 01 to 03**

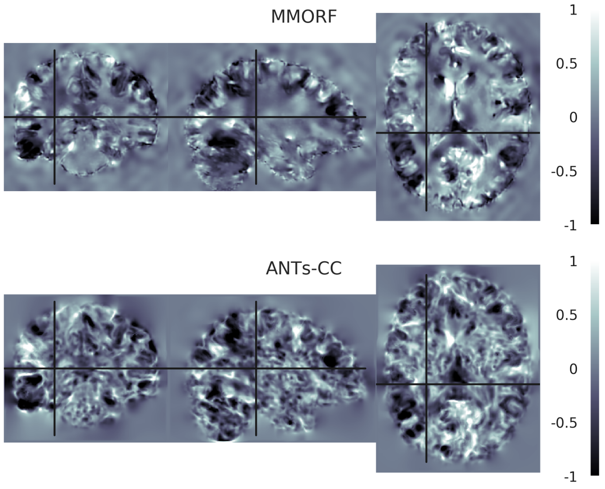


**Log-Jacobian determinant spatial maps - subject 02 to 03**

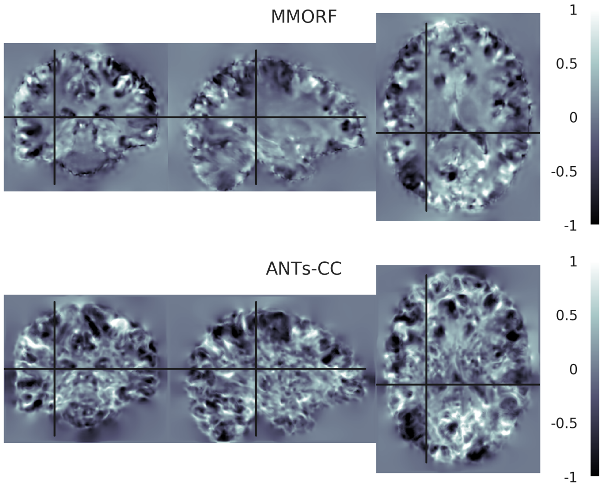


**Log-Jacobian determinant spatial maps - subject 04 to 03**

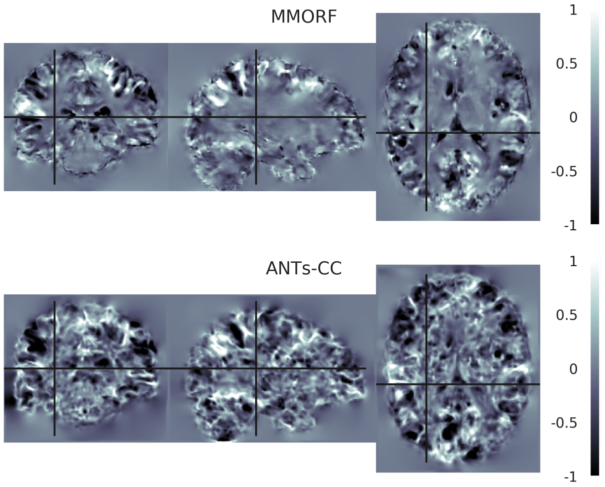


**Log-Jacobian determinant spatial maps - subject 05 to 03**

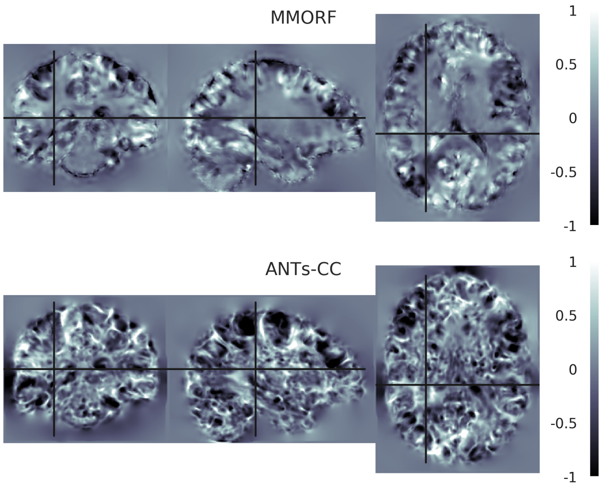


**Log-Jacobian determinant spatial maps - subject 06 to 03**

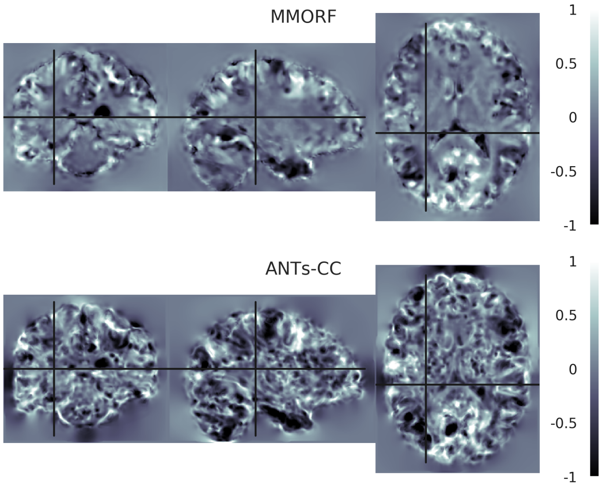


**Log-Jacobian determinant spatial maps - subject 07 to 03**

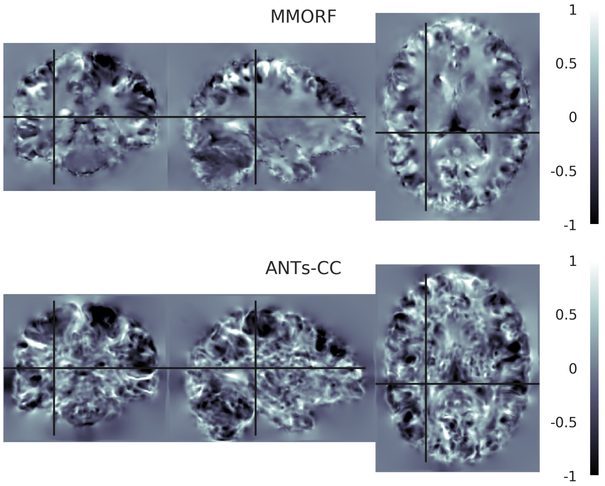

**Log-Jacobian determinant spatial maps - subject 08 to 03**

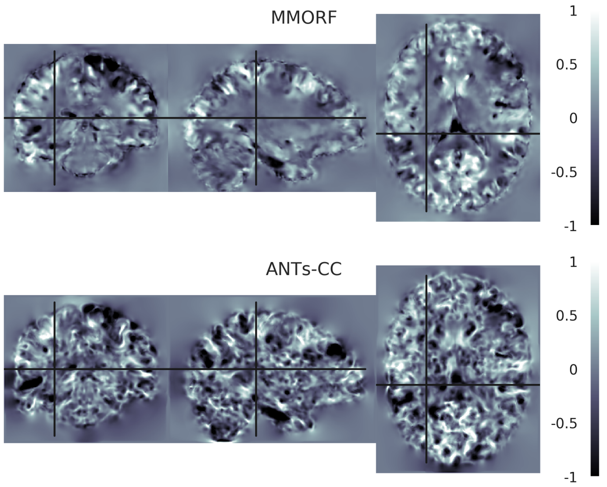


**Log-Jacobian determinant spatial maps - subject 09 to 03**

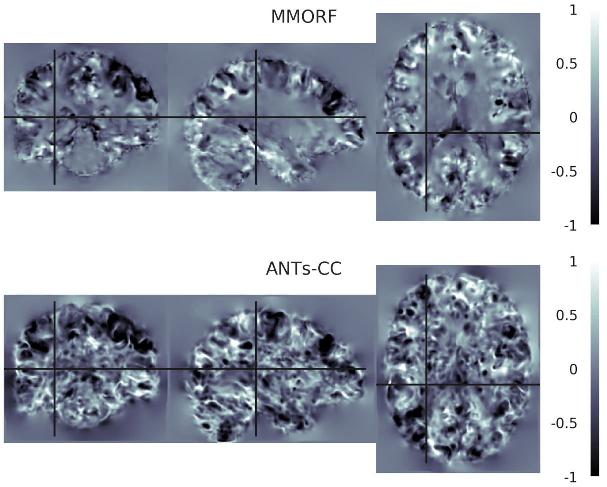


**Log-Jacobian determinant spatial maps - subject 10 to 03**

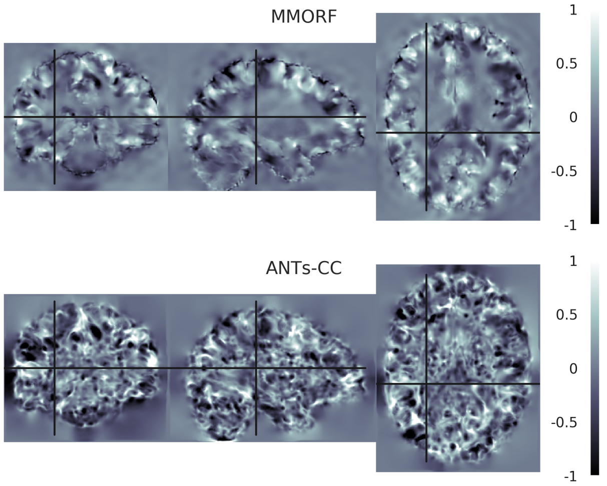


**Log-Jacobian determinant spatial maps - subject 11 to 03**

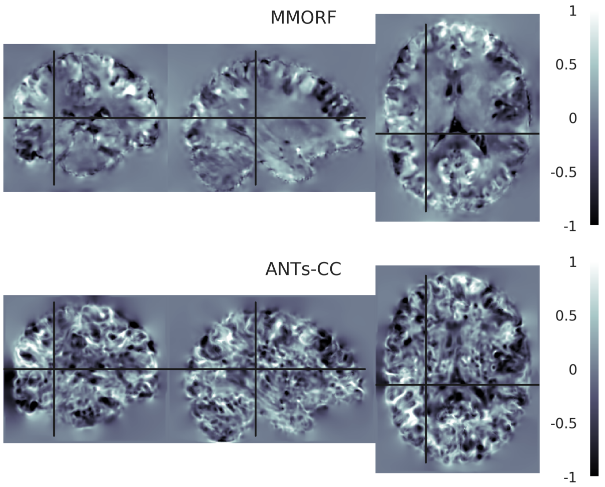


**Log-Jacobian determinant spatial maps - subject 12 to 03**

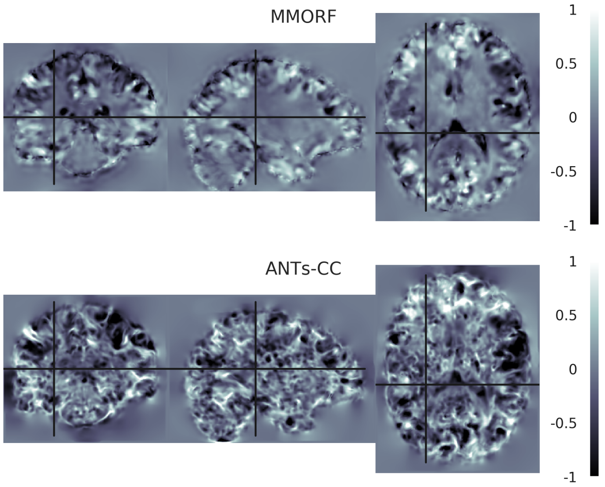


**Log-Jacobian determinant spatial maps - subject 13 to 03**

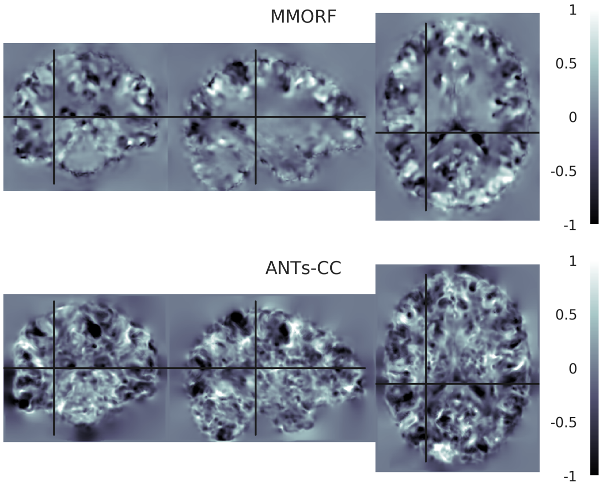

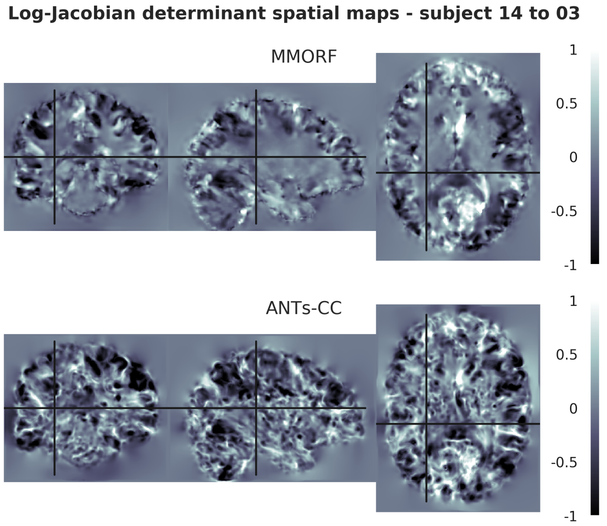
Log-Jacobian determinant spatial maps - subject 14 to 03
MMORF
ANTs-CC
1
0.5
0
-0.5
-1

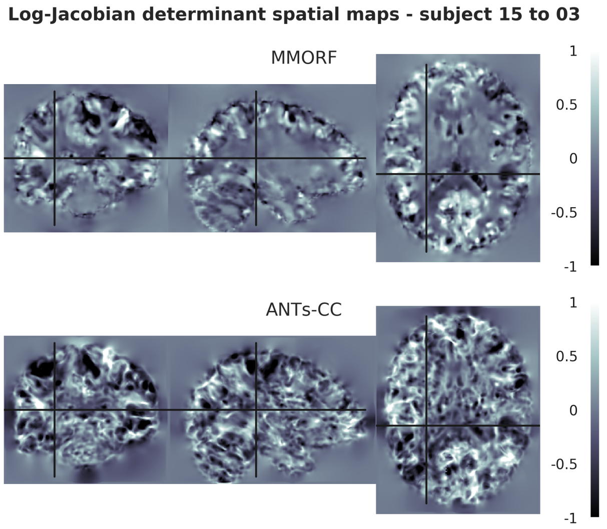
Log-Jacobian determinant spatial maps - subject 15 to 03
MMORF
ANTs-CC
1
0.5
0
-0.5
-1

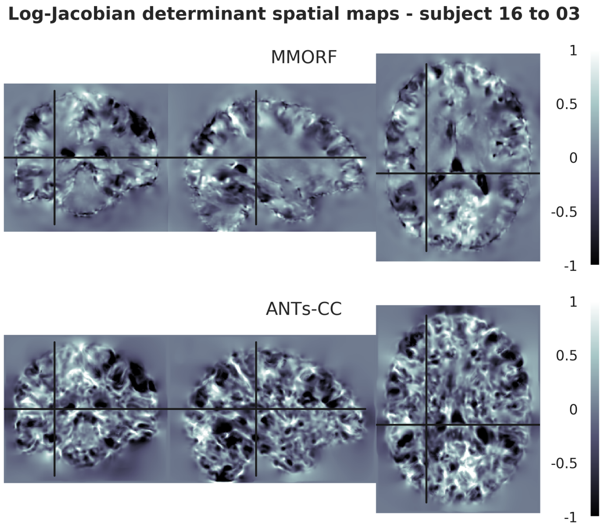
Log-Jacobian determinant spatial maps - subject 16 to 03
MMORF
ANTs-CC
1
0.5
0
-0.5
-1

#### 4. Reference Subject 04

**Log-Jacobian determinant spatial maps - subject 01 to 04**

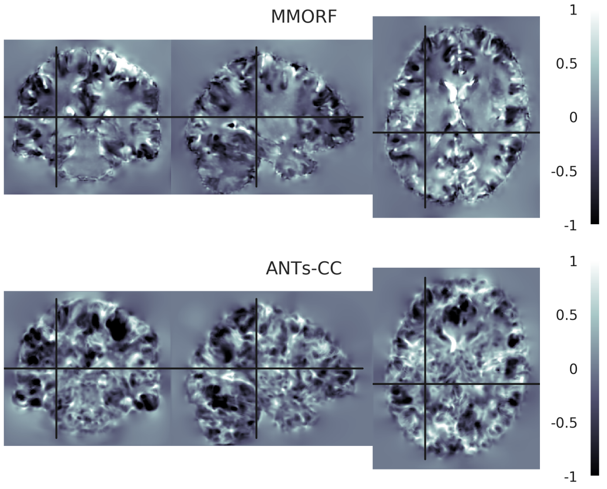


**Log-Jacobian determinant spatial maps - subject 02 to 04**

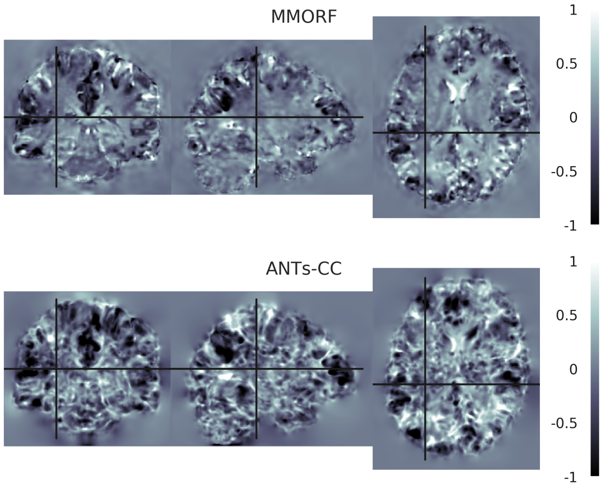


**Log-Jacobian determinant spatial maps - subject 03 to 04**

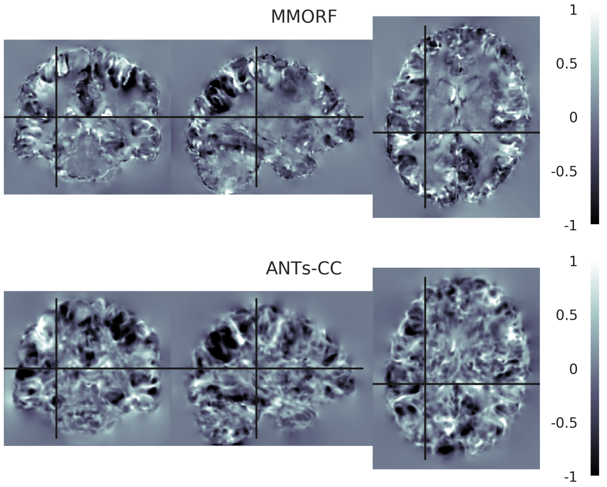


**Log-Jacobian determinant spatial maps - subject 05 to 04**

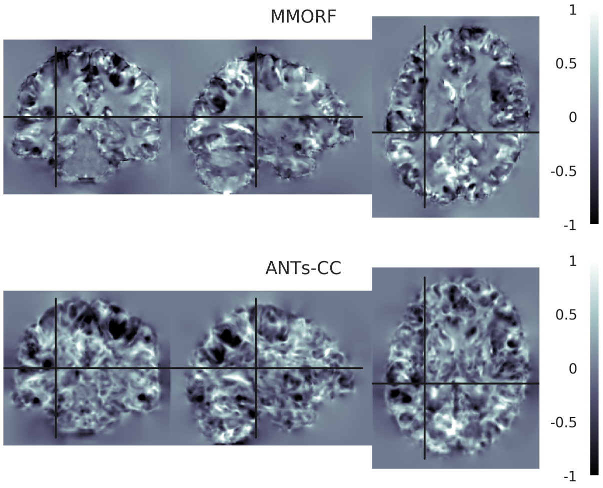


**Log-Jacobian determinant spatial maps - subject 06 to 04**

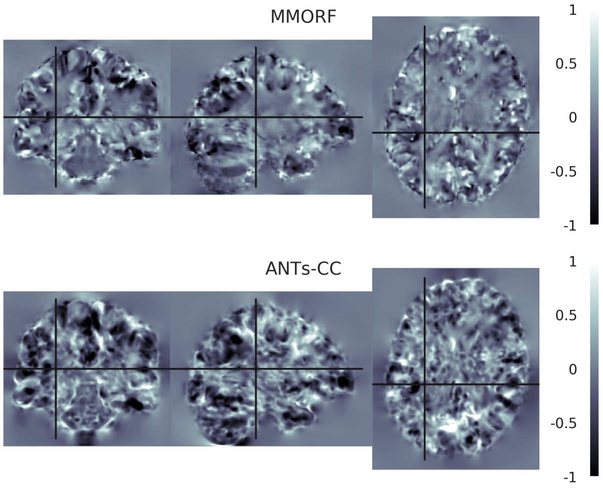


**Log-Jacobian determinant spatial maps - subject 07 to 04**

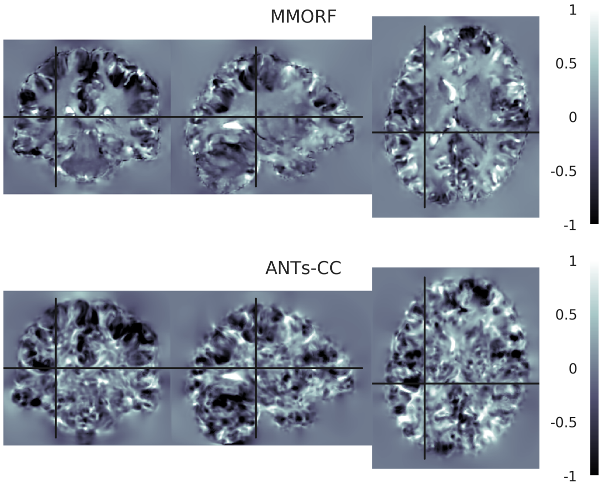

**Log-Jacobian determinant spatial maps - subject 08 to 04**

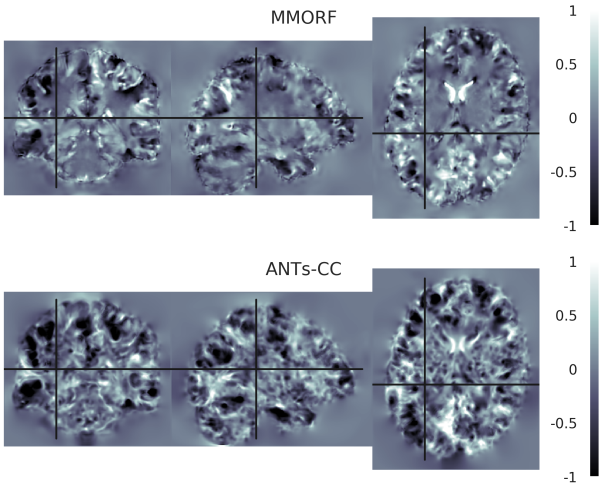


**Log-Jacobian determinant spatial maps - subject 11 to 04**

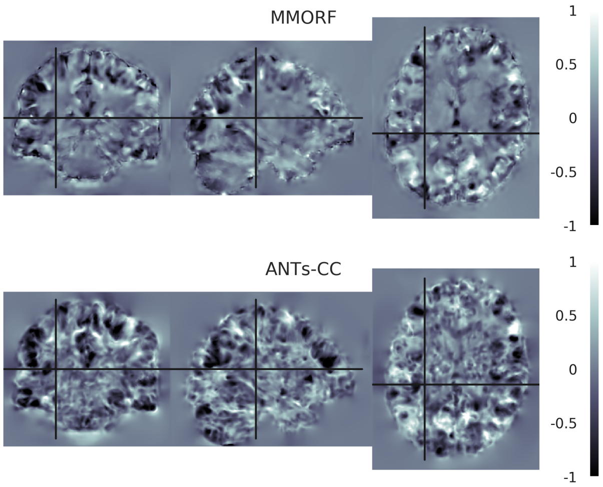


**Log-Jacobian determinant spatial maps - subject 09 to 04**

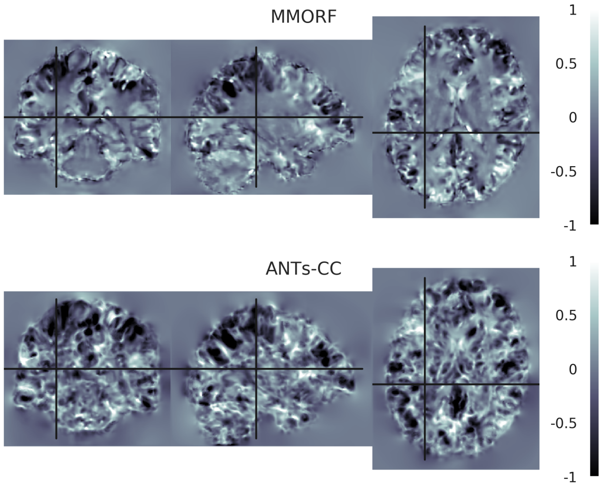


**Log-Jacobian determinant spatial maps - subject 12 to 04**

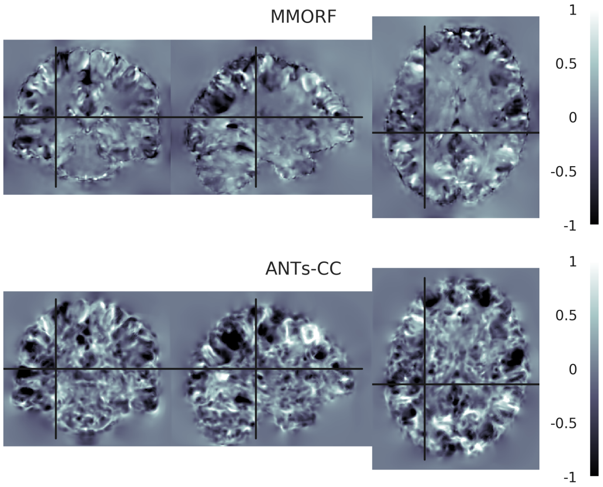


**Log-Jacobian determinant spatial maps - subject 10 to 04**

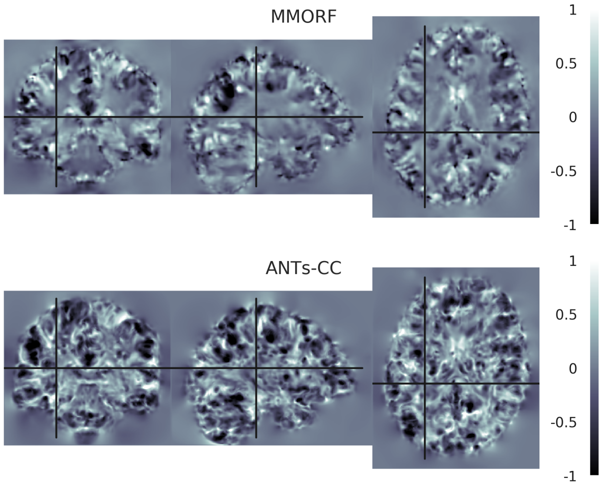


**Log-Jacobian determinant spatial maps - subject 13 to 04**

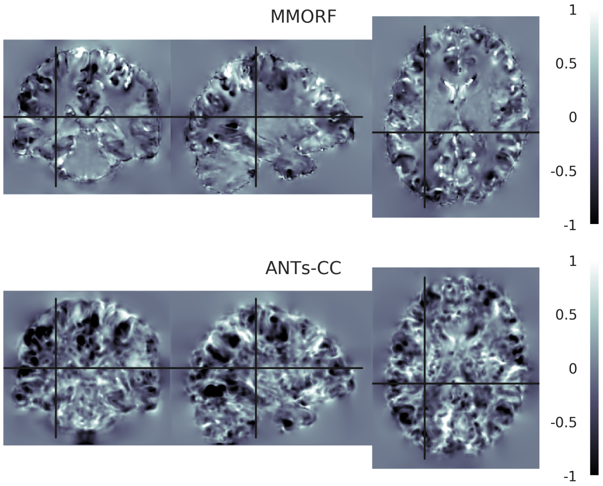

**Log-Jacobian determinant spatial maps - subject 14 to 04**

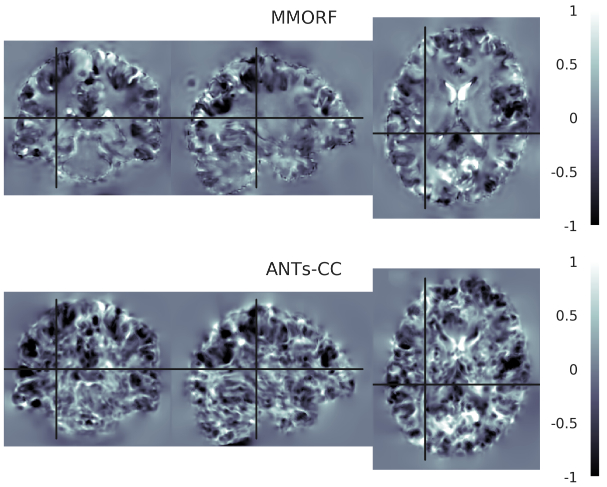


**Log-Jacobian determinant spatial maps - subject 15 to 04**

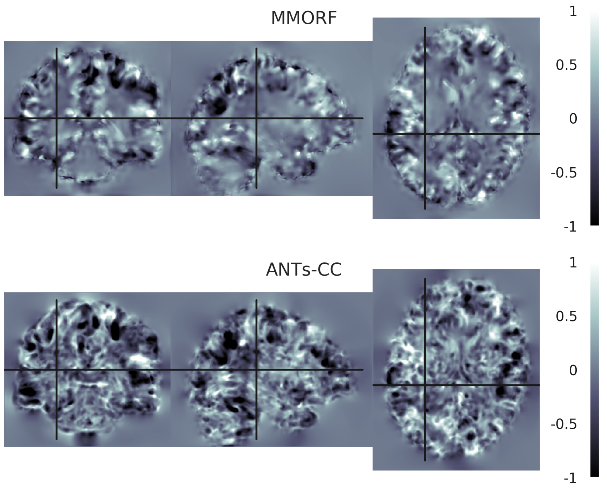


**Log-Jacobian determinant spatial maps - subject 16 to 04**

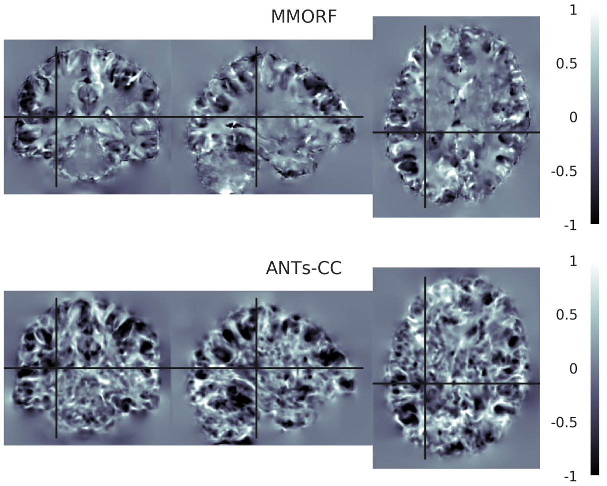

#### 5. Reference Subject 05

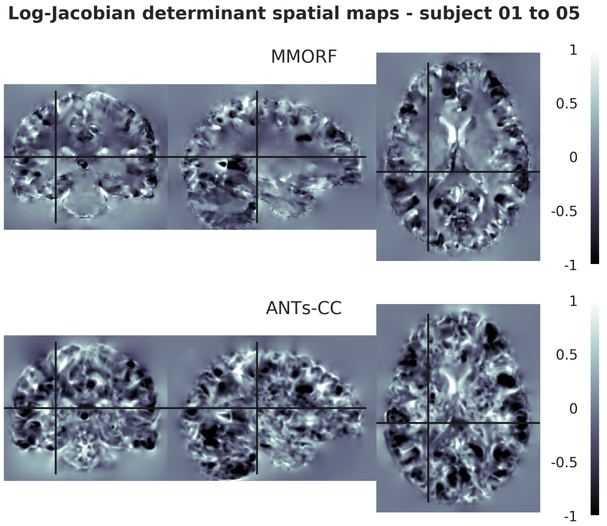


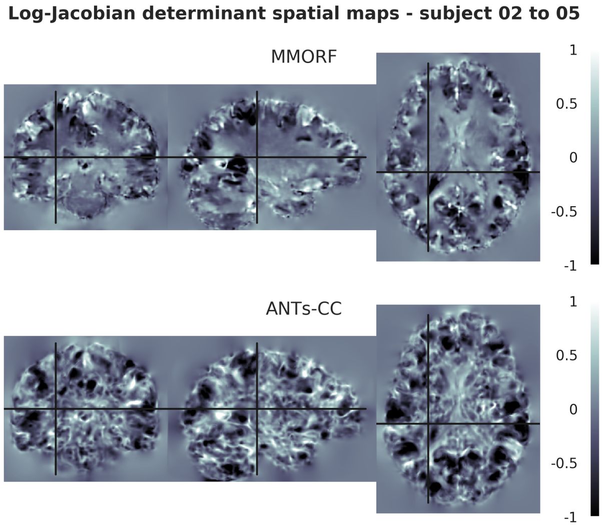


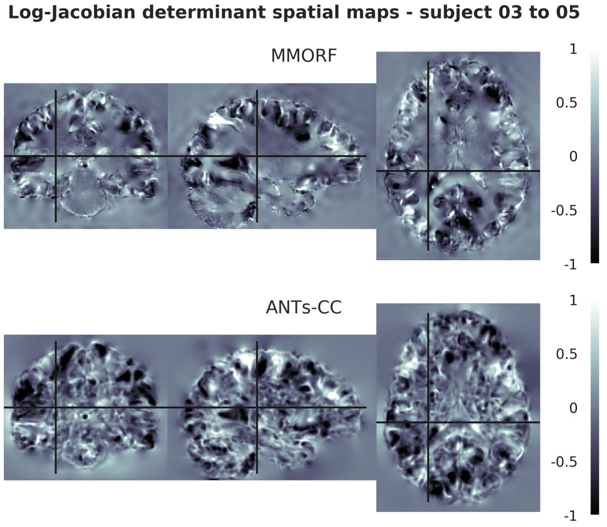


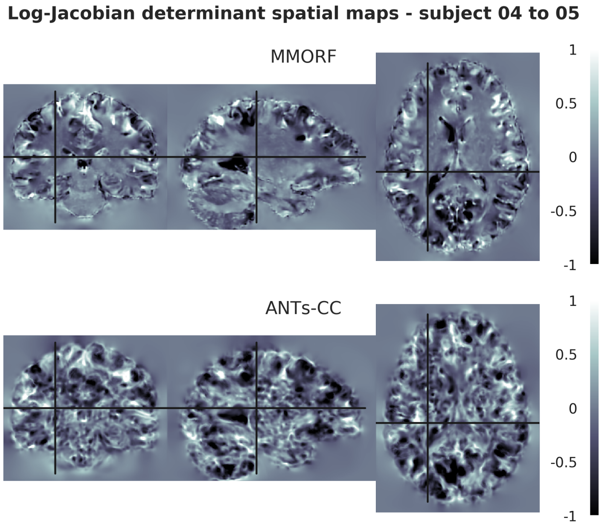


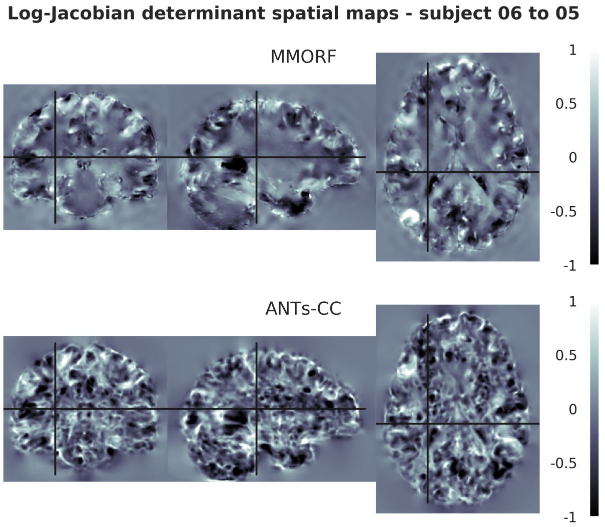


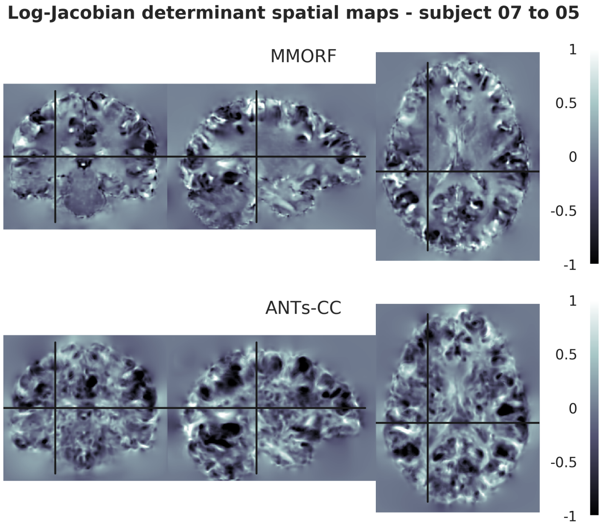

**Log-Jacobian determinant spatial maps - subject 08 to 05**

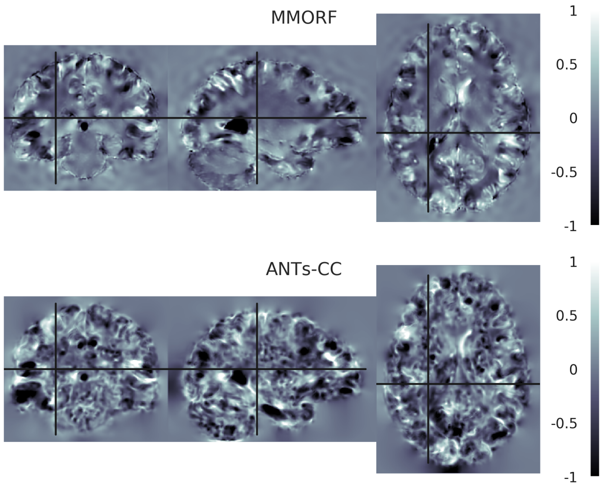


**Log-Jacobian determinant spatial maps - subject 09 to 05**

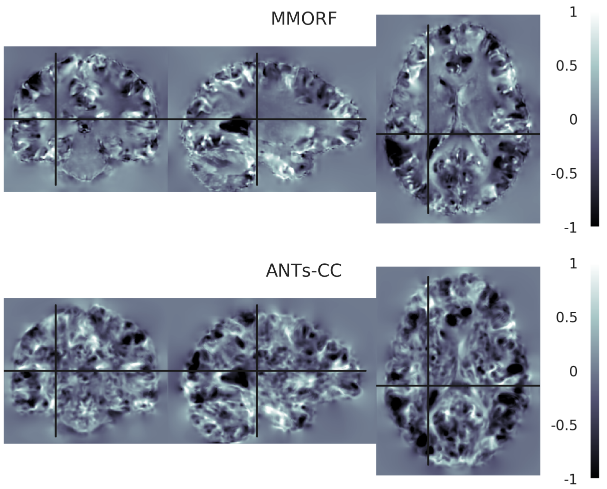


**Log-Jacobian determinant spatial maps - subject 10 to 05**

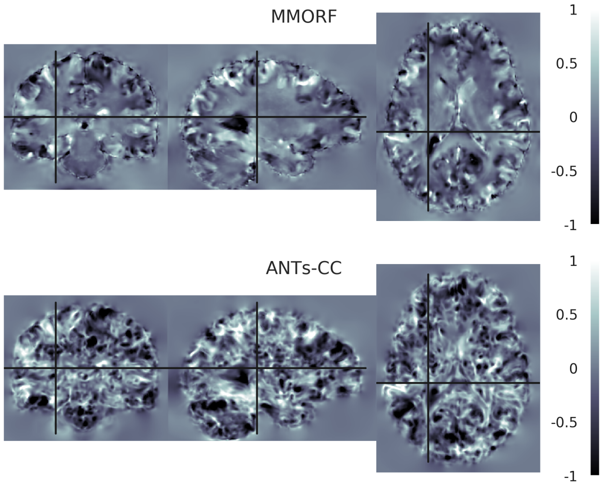


**Log-Jacobian determinant spatial maps - subject 11 to 05**

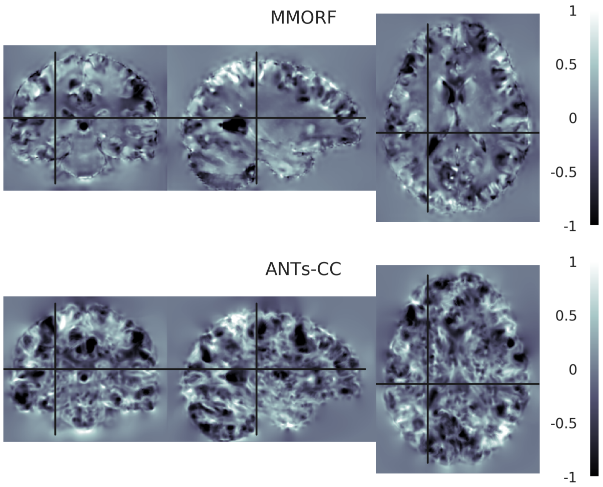


**Log-Jacobian determinant spatial maps - subject 12 to 05**

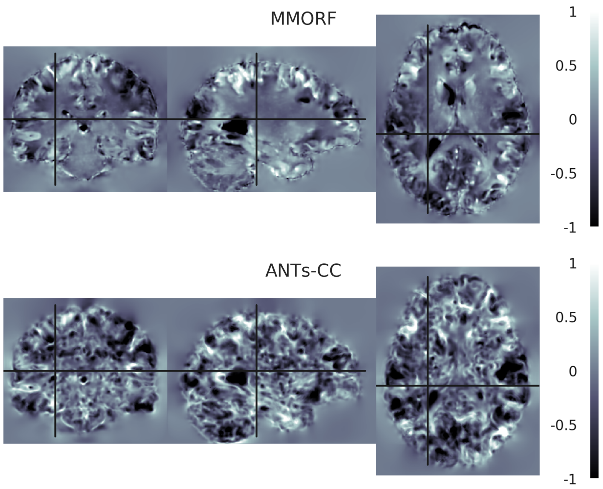


**Log-Jacobian determinant spatial maps - subject 13 to 05**

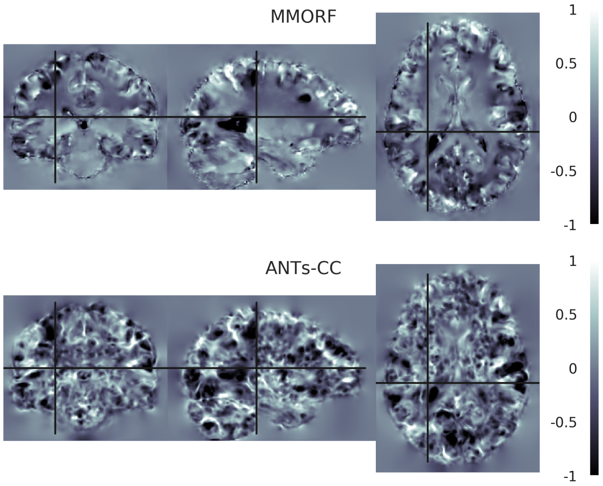

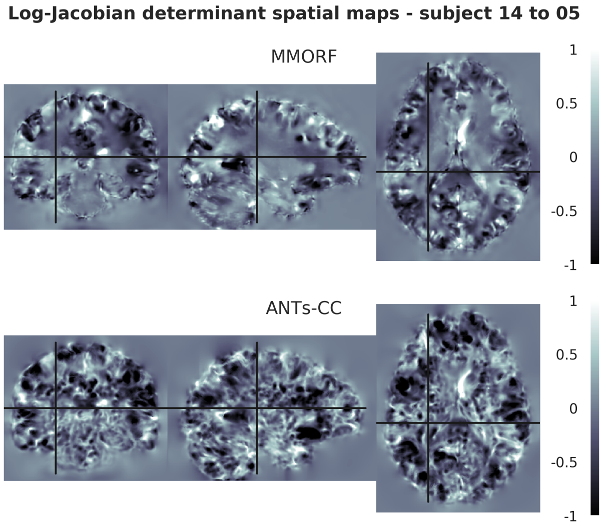
Log-Jacobian determinant spatial maps - subject 14 to 05
MMORF
ANTs-CC
1
0.5
0
-0.5
-1

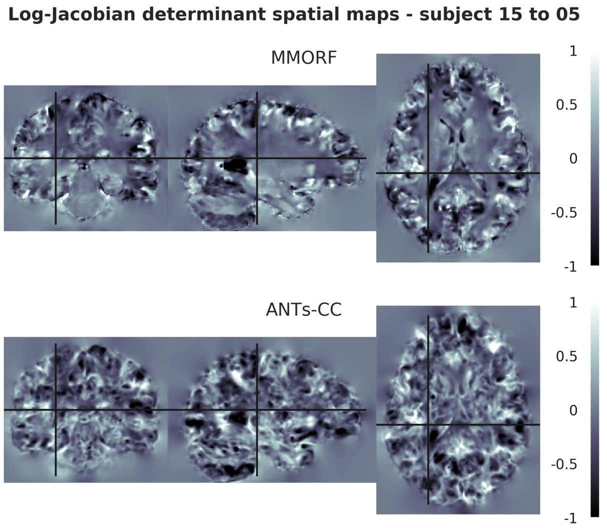
Log-Jacobian determinant spatial maps - subject 15 to 05
MMORF
ANTs-CC
1
0.5
0
-0.5
-1

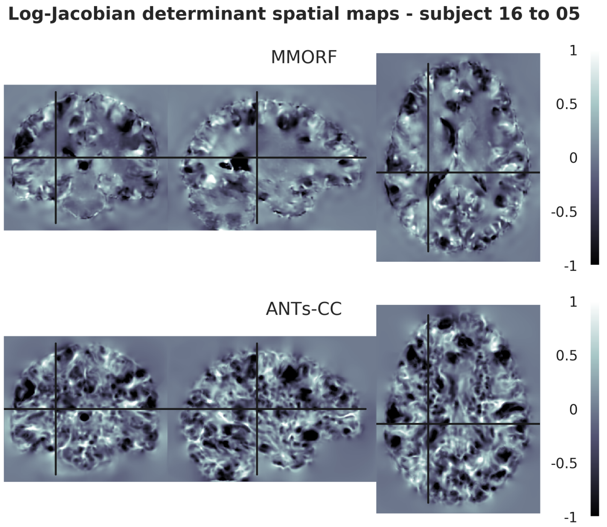
Log-Jacobian determinant spatial maps - subject 16 to 05
MMORF
ANTs-CC
1
0.5
0
-0.5
-1

#### 6. Reference Subject 06

**Log-Jacobian determinant spatial maps - subject 01 to 06**

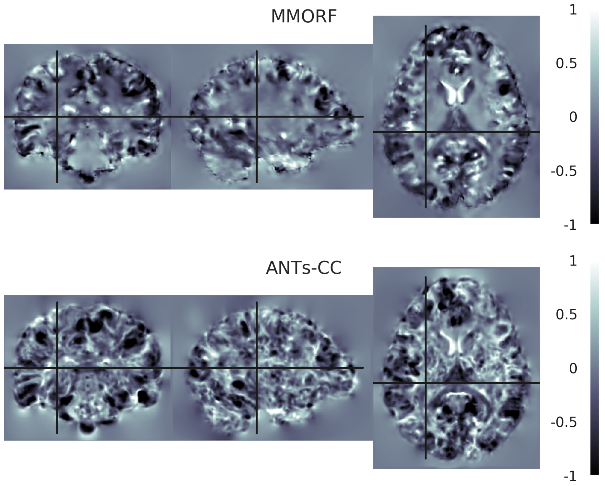


**Log-Jacobian determinant spatial maps - subject 02 to 06**

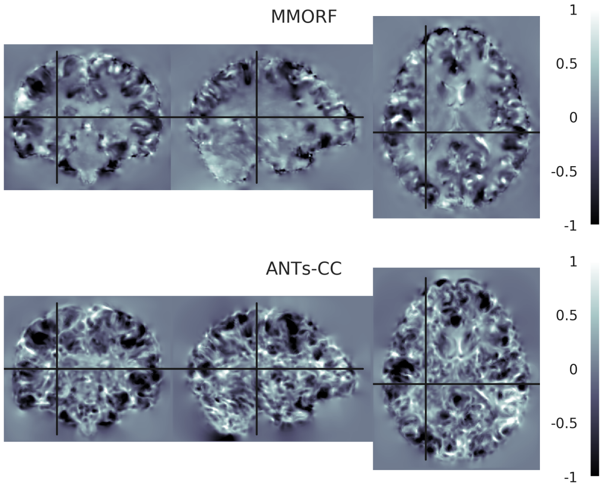


**Log-Jacobian determinant spatial maps - subject 03 to 06**

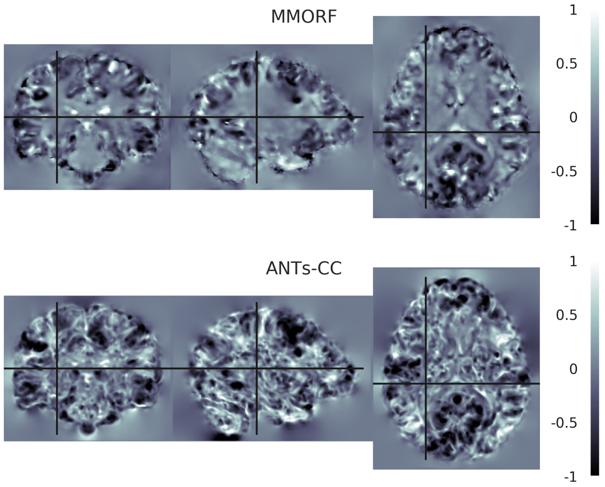


**Log-Jacobian determinant spatial maps - subject 04 to 06**

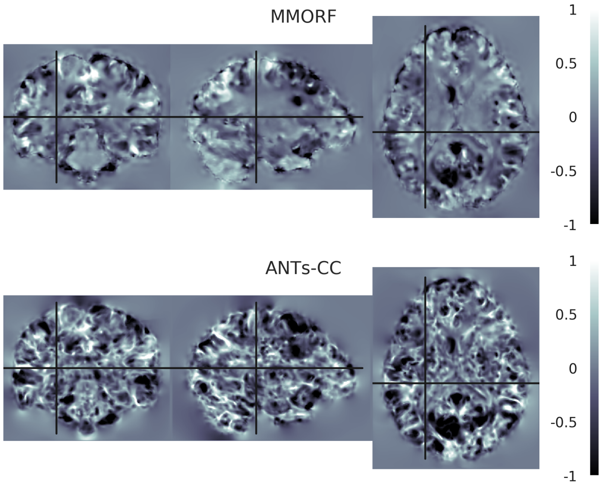


**Log-Jacobian determinant spatial maps - subject 05 to 06**

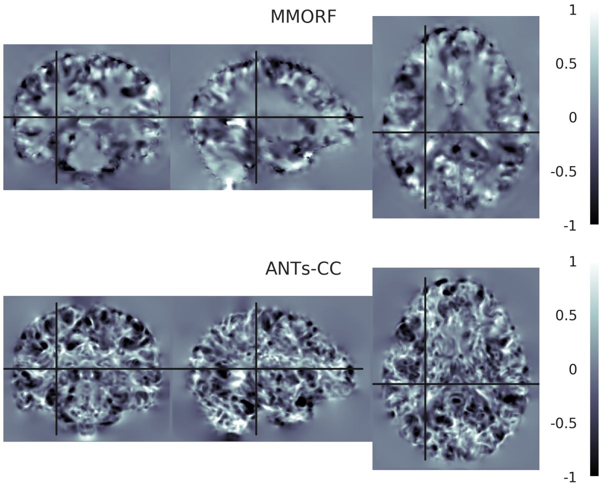


**Log-Jacobian determinant spatial maps - subject 07 to 06**

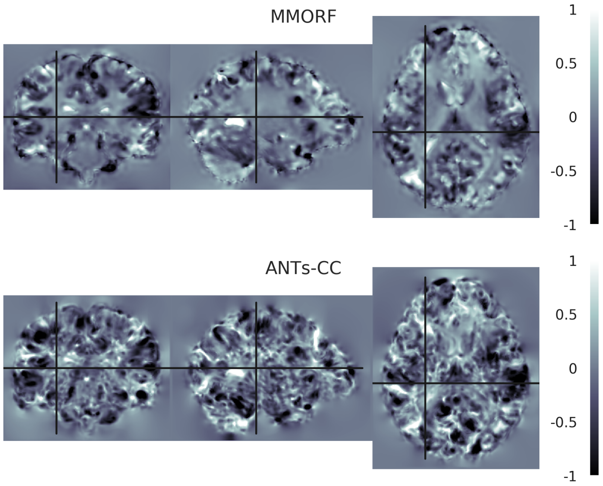

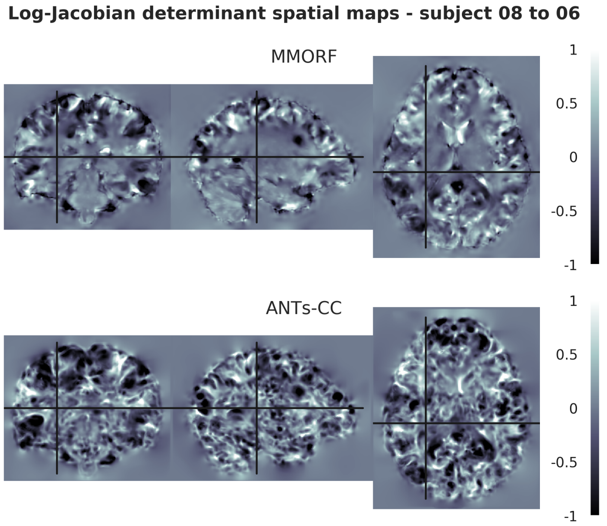
Log-Jacobian determinant spatial maps - subject 08 to 06
MMORF
ANTs-CC
1
0.5
0
-0.5
-1

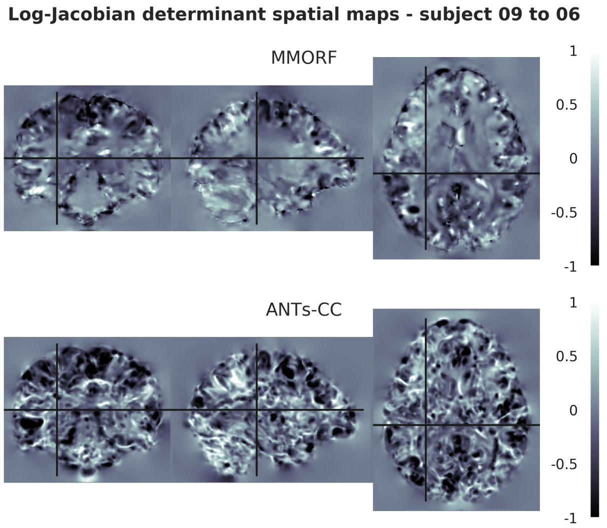
Log-Jacobian determinant spatial maps - subject 09 to 06
MMORF
ANTs-CC
1
0.5
0
-0.5
-1

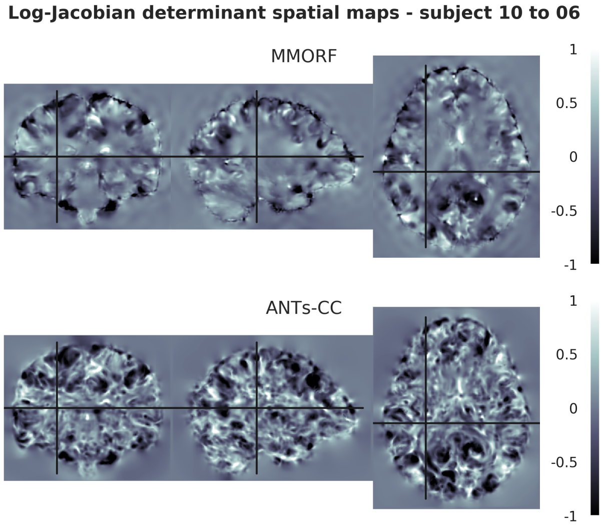
Log-Jacobian determinant spatial maps - subject 10 to 06
MMORF
ANTs-CC
1
0.5
0
-0.5
-1

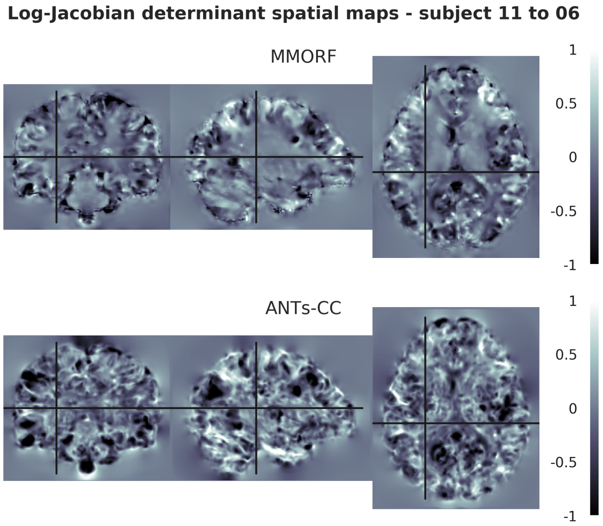
Log-Jacobian determinant spatial maps - subject 11 to 06
MMORF
ANTs-CC
1
0.5
0
-0.5
-1

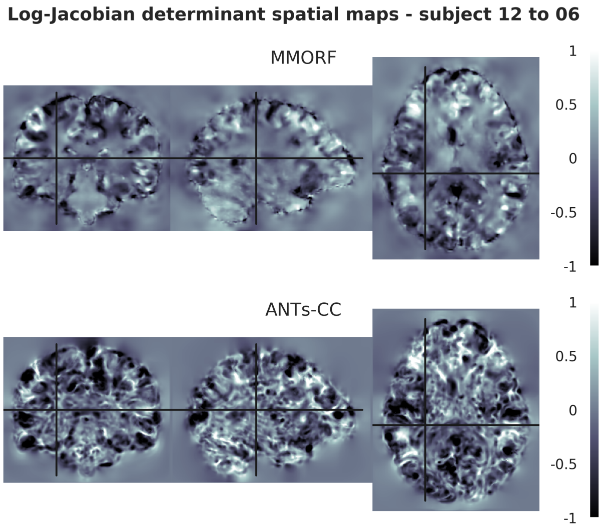
Log-Jacobian determinant spatial maps - subject 12 to 06
MMORF
ANTs-CC
1
0.5
0
-0.5
-1

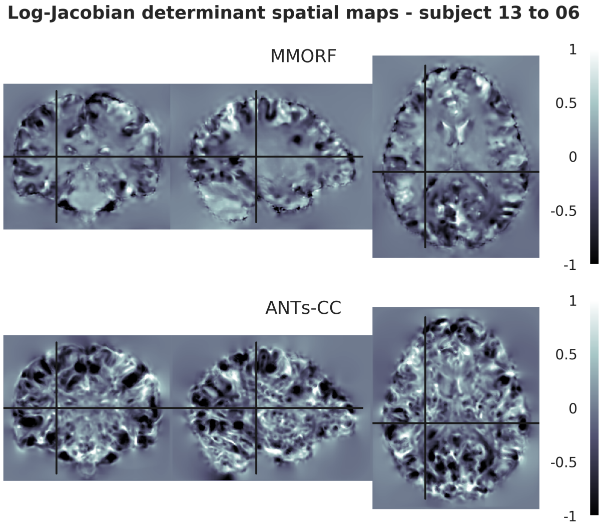
Log-Jacobian determinant spatial maps - subject 13 to 06
MMORF
ANTs-CC
1
0.5
0
-0.5
-1

**Log-Jacobian determinant spatial maps - subject 14 to 06**

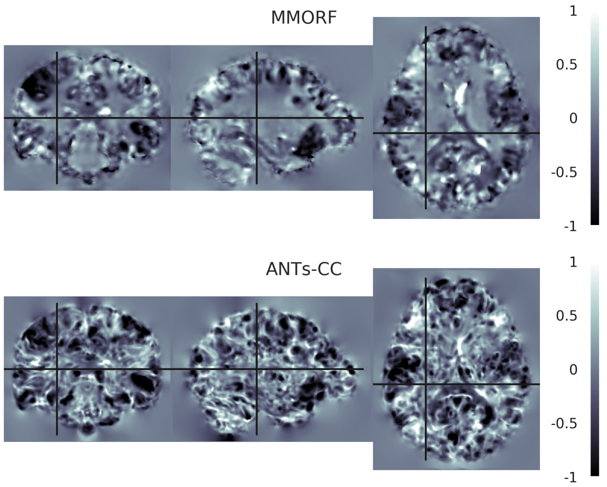


**Log-Jacobian determinant spatial maps - subject 15 to 06**

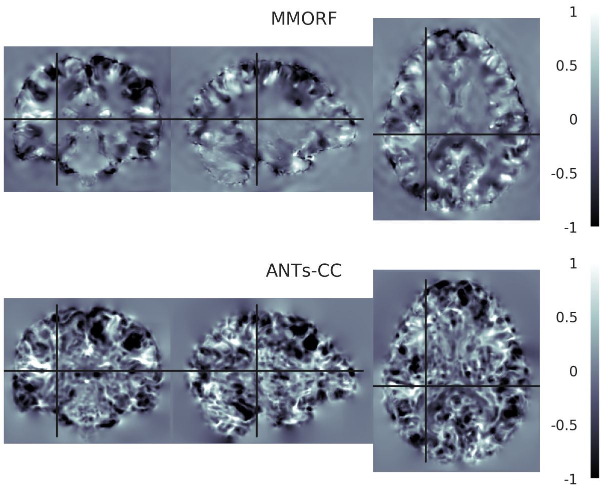


**Log-Jacobian determinant spatial maps - subject 16 to 06**

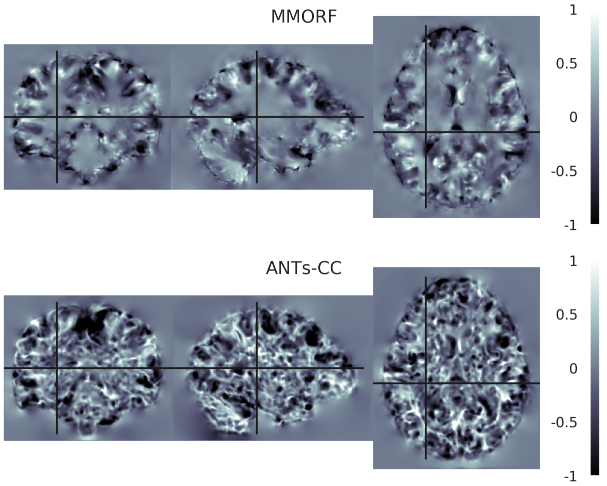

#### 7. Reference Subject 07

**Log-Jacobian determinant spatial maps - subject 01 to 07**

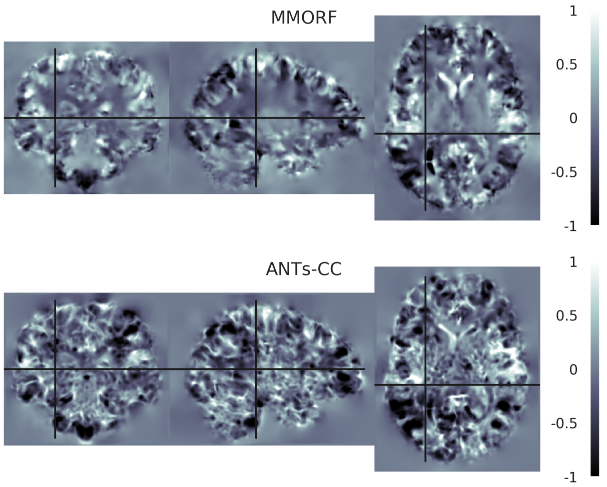


**Log-Jacobian determinant spatial maps - subject 02 to 07**

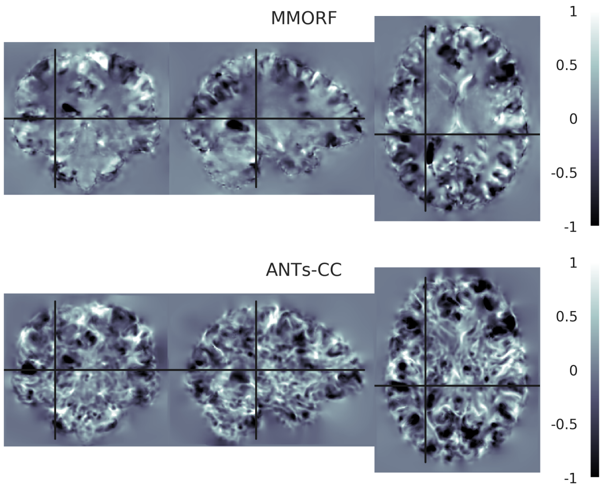


**Log-Jacobian determinant spatial maps - subject 03 to 07**

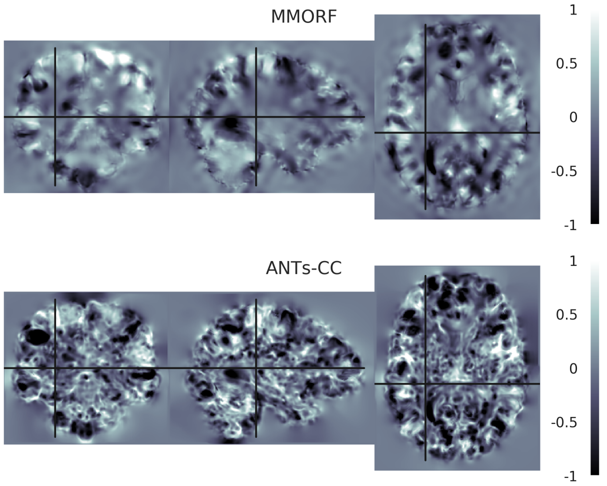


**Log-Jacobian determinant spatial maps - subject 04 to 07**

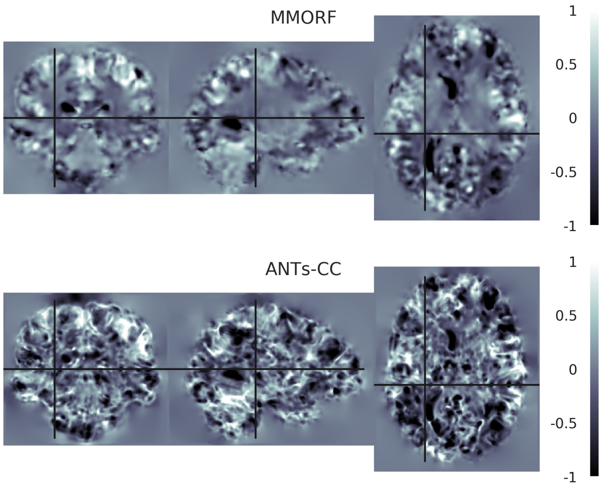


**Log-Jacobian determinant spatial maps - subject 05 to 07**

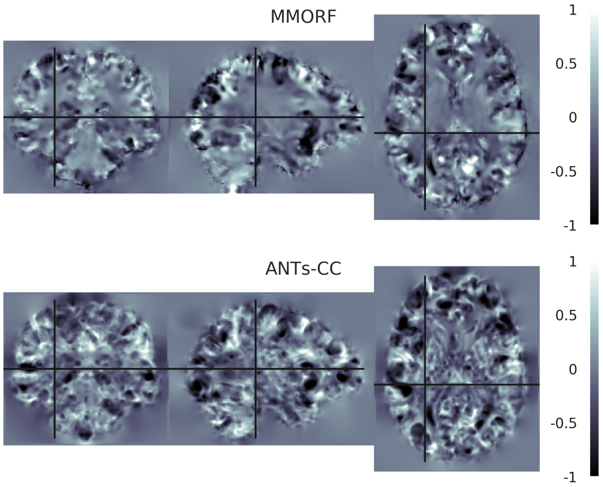


**Log-Jacobian determinant spatial maps - subject 06 to 07**

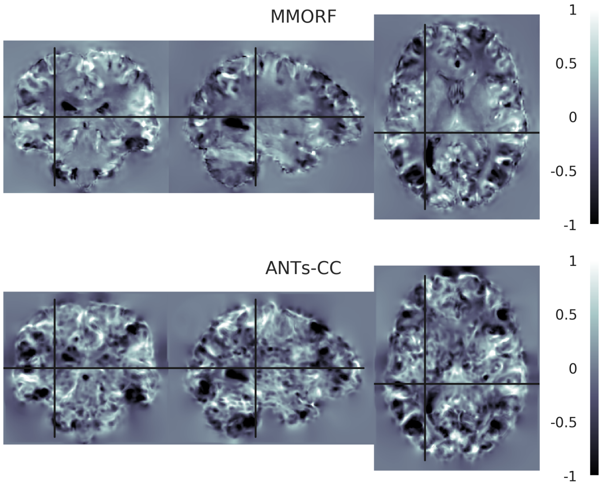

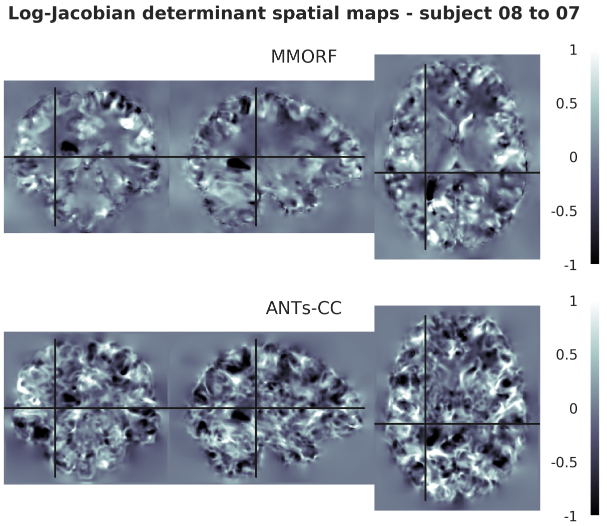
Log-Jacobian determinant spatial maps - subject 08 to 07
MMORF
ANTs-CC
1
0.5
0
-0.5
-1

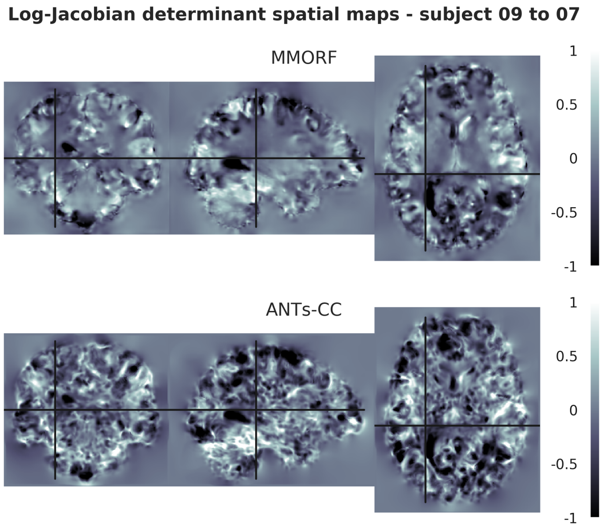
Log-Jacobian determinant spatial maps - subject 09 to 07
MMORF
ANTs-CC
1
0.5
0
-0.5
-1

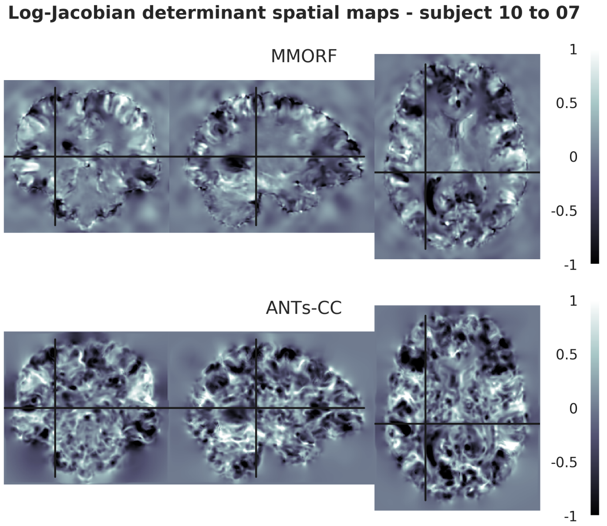
Log-Jacobian determinant spatial maps - subject 10 to 07
MMORF
ANTs-CC
1
0.5
0
-0.5
-1

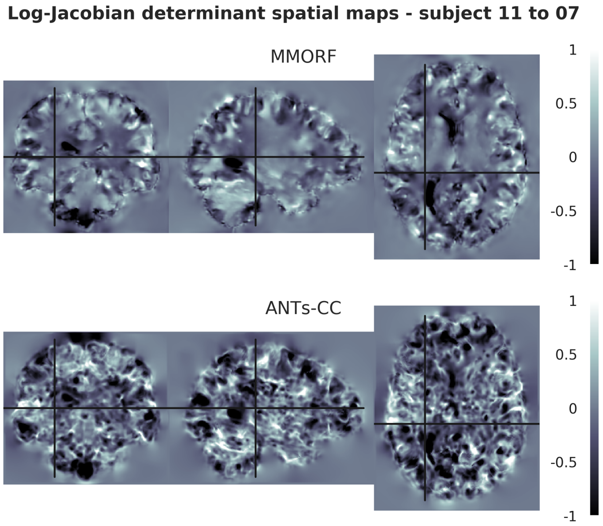
Log-Jacobian determinant spatial maps - subject 11 to 07
MMORF
ANTs-CC
1
0.5
0
-0.5
-1

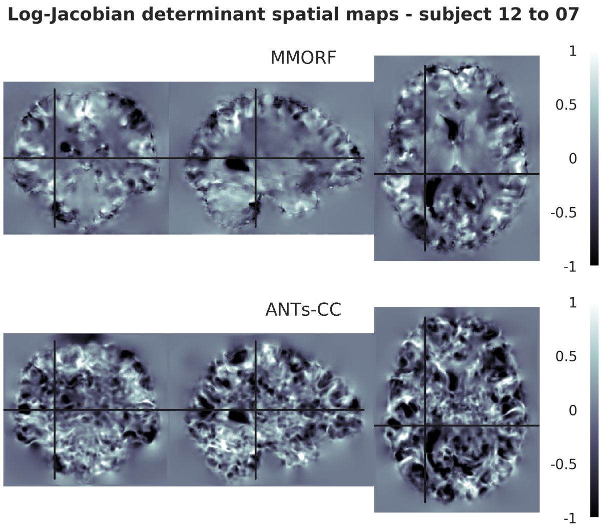
Log-Jacobian determinant spatial maps - subject 12 to 07
MMORF
ANTs-CC
1
0.5
0
-0.5
-1

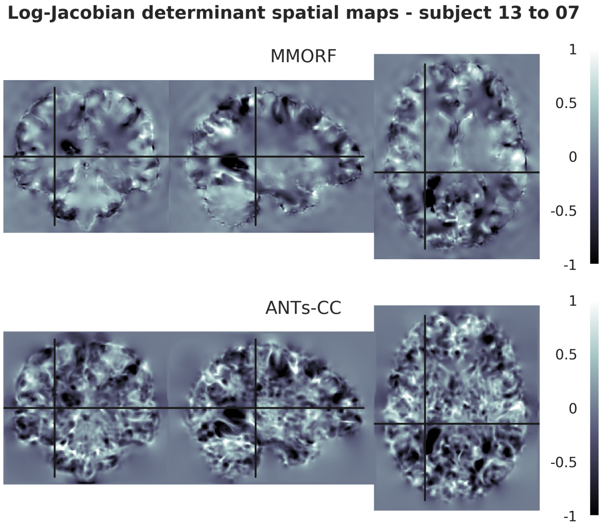
Log-Jacobian determinant spatial maps - subject 13 to 07
MMORF
ANTs-CC
1
0.5
0
-0.5
-1

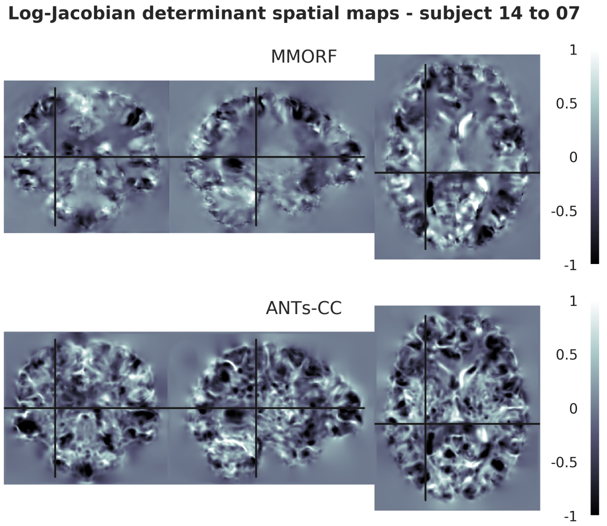
Log-Jacobian determinant spatial maps - subject 14 to 07
MMORF
ANTs-CC
1
0.5
0
-0.5
-1

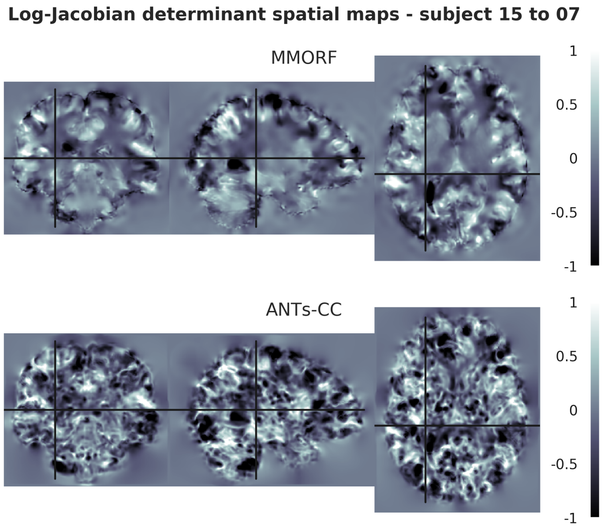
Log-Jacobian determinant spatial maps - subject 15 to 07
MMORF
ANTs-CC
1
0.5
0
-0.5
-1

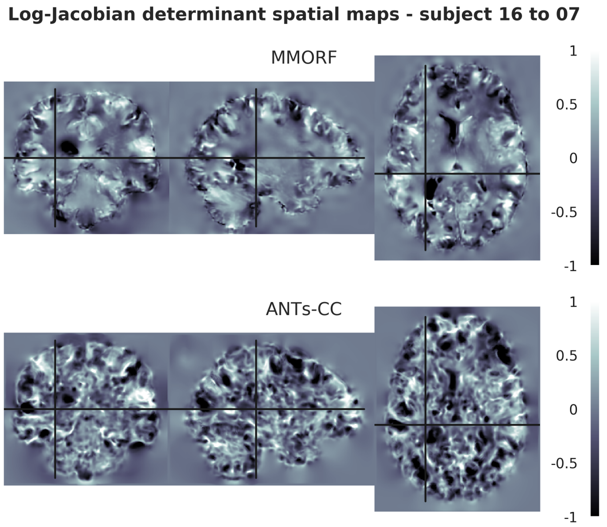
Log-Jacobian determinant spatial maps - subject 16 to 07
MMORF
ANTs-CC
1
0.5
0
-0.5
-1

#### 8. Reference Subject 08

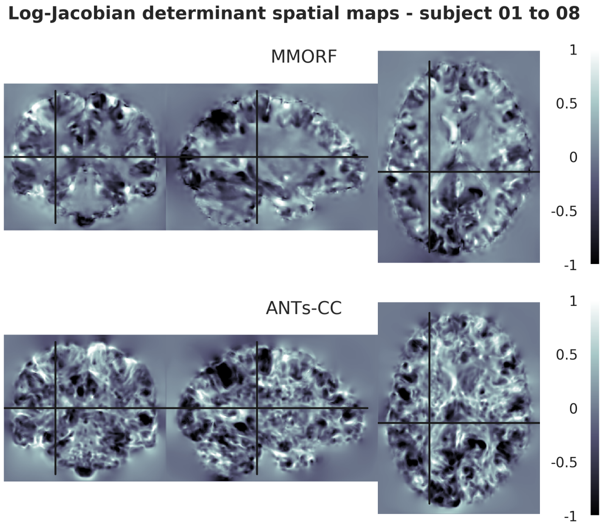


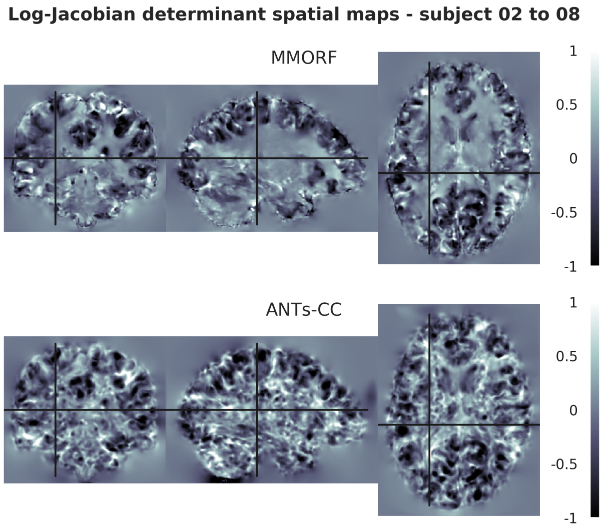


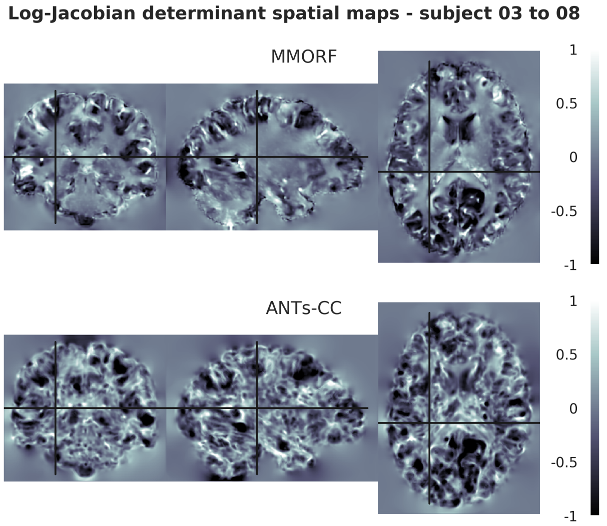


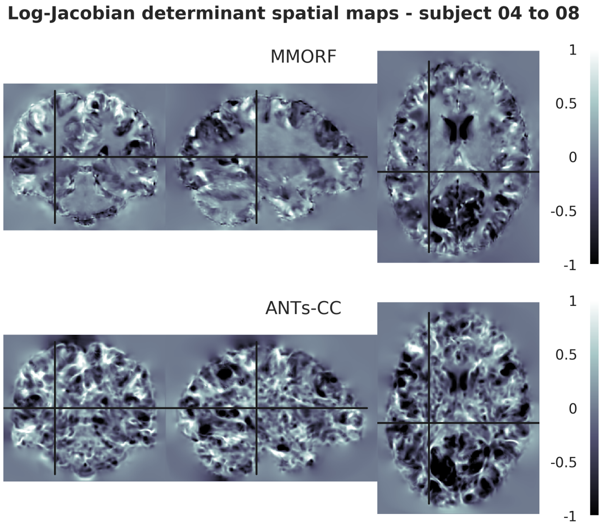


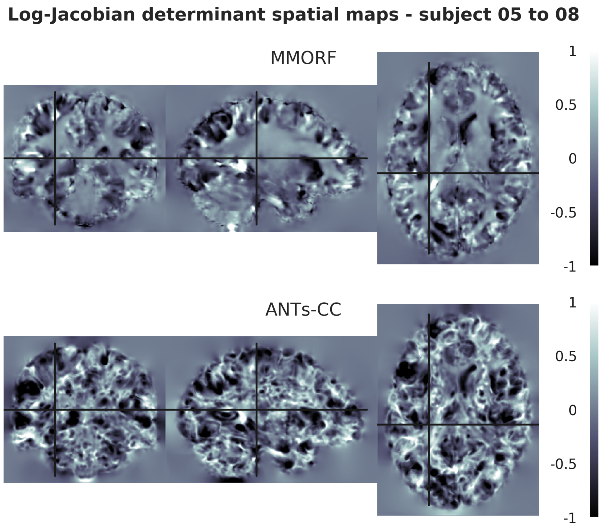


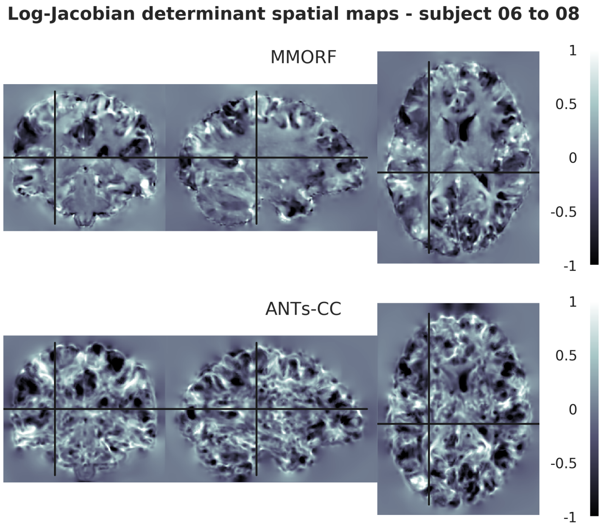

**Log-Jacobian determinant spatial maps - subject 07 to 08**

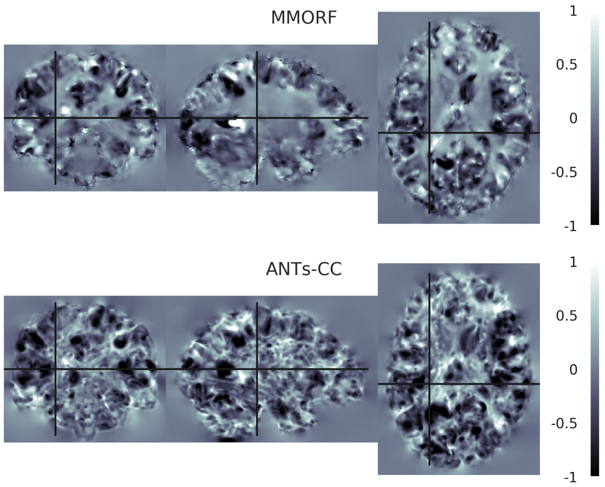


**Log-Jacobian determinant spatial maps - subject 09 to 08**

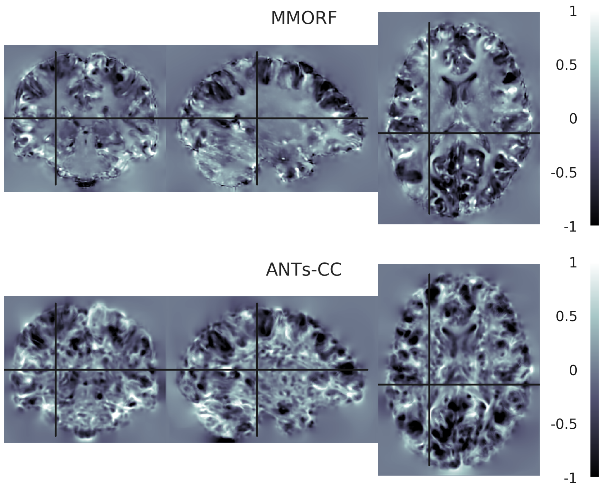


**Log-Jacobian determinant spatial maps - subject 10 to 08**

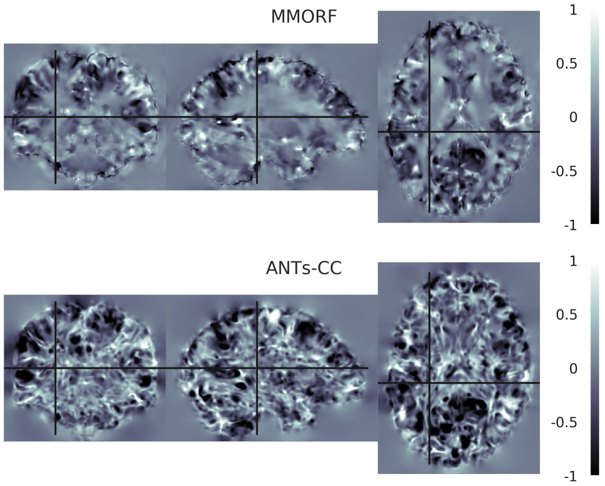


**Log-Jacobian determinant spatial maps - subject 11 to 08**

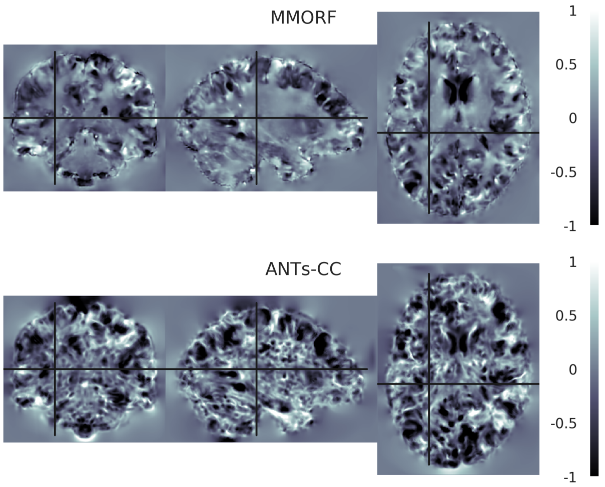


**Log-Jacobian determinant spatial maps - subject 12 to 08**

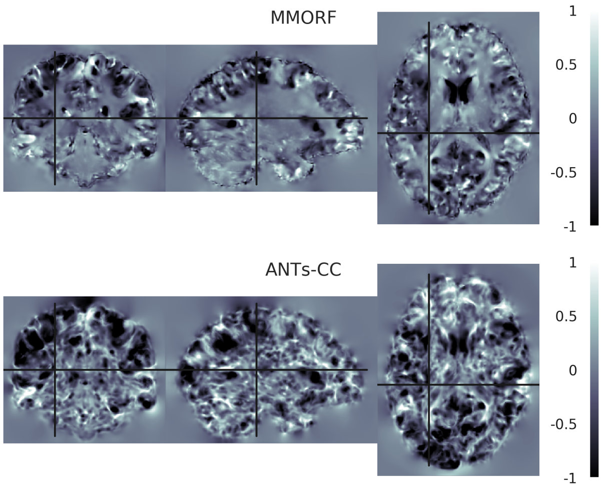


**Log-Jacobian determinant spatial maps - subject 13 to 08**

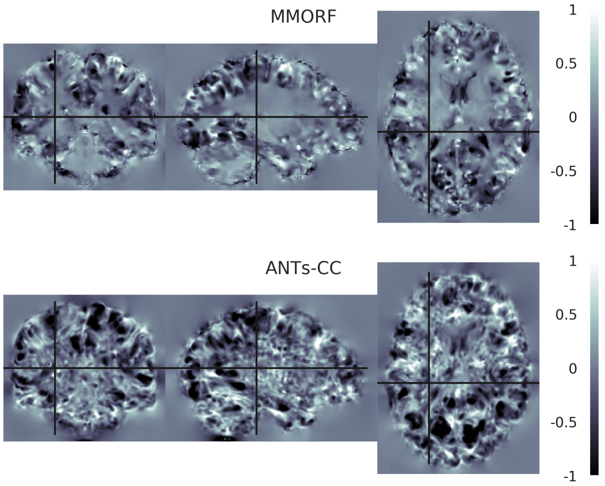

**Log-Jacobian determinant spatial maps - subject 14 to 08**

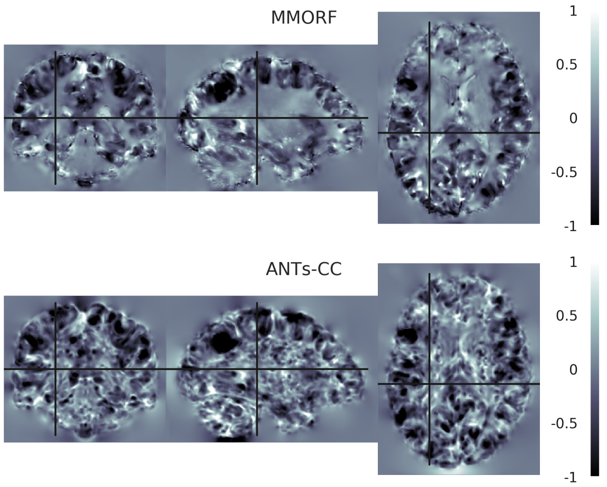


**Log-Jacobian determinant spatial maps - subject 15 to 08**

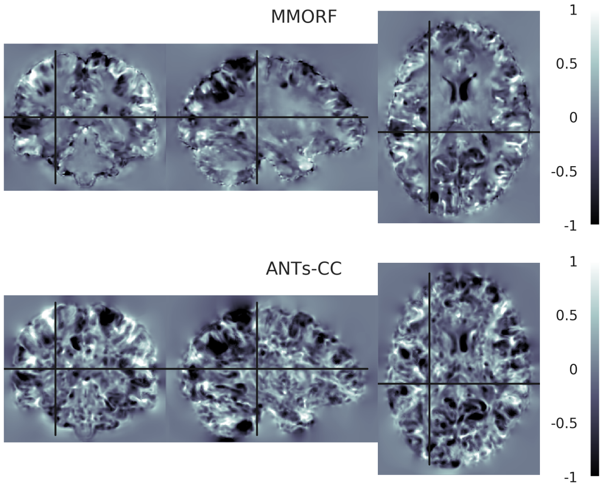


**Log-Jacobian determinant spatial maps - subject 16 to 08**

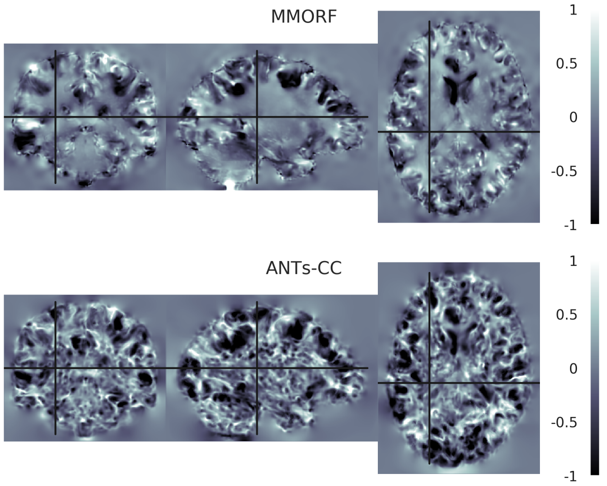

#### 9. Reference Subject 09

**Log-Jacobian determinant spatial maps - subject 01 to 09**

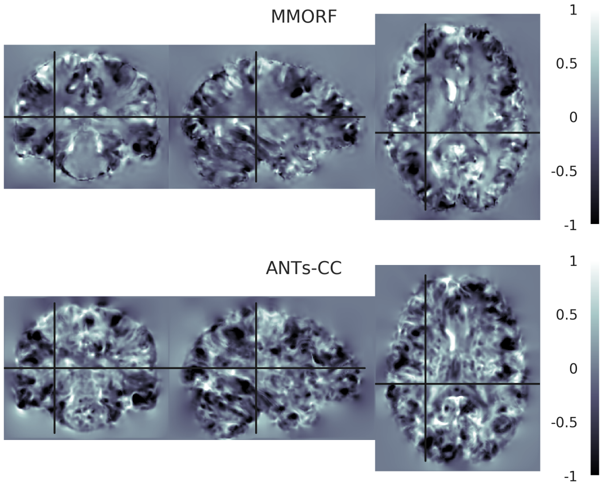


**Log-Jacobian determinant spatial maps - subject 02 to 09**

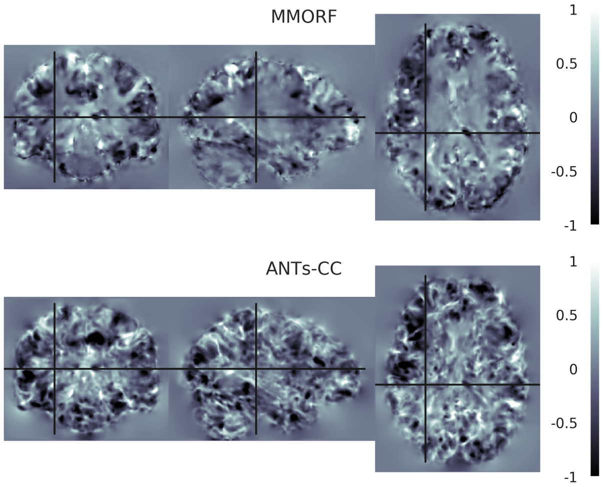


**Log-Jacobian determinant spatial maps - subject 03 to 09**

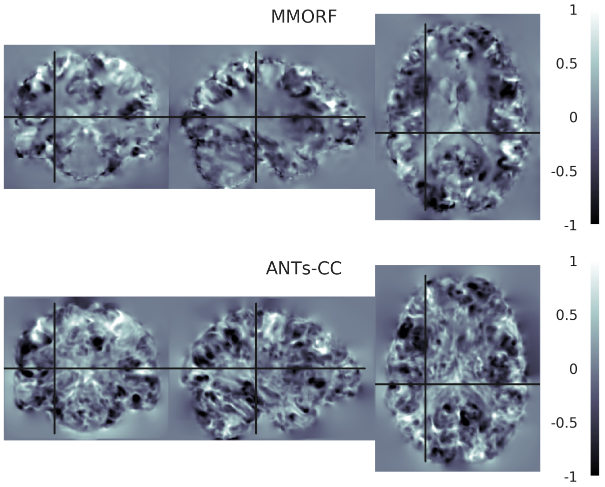


**Log-Jacobian determinant spatial maps - subject 04 to 09**

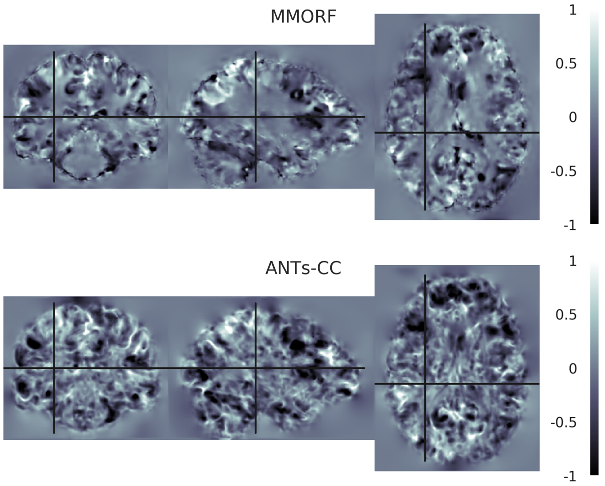


**Log-Jacobian determinant spatial maps - subject 05 to 09**

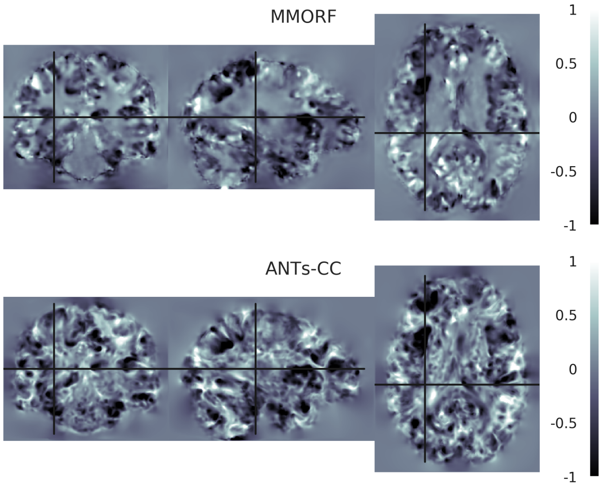


**Log-Jacobian determinant spatial maps - subject 06 to 09**

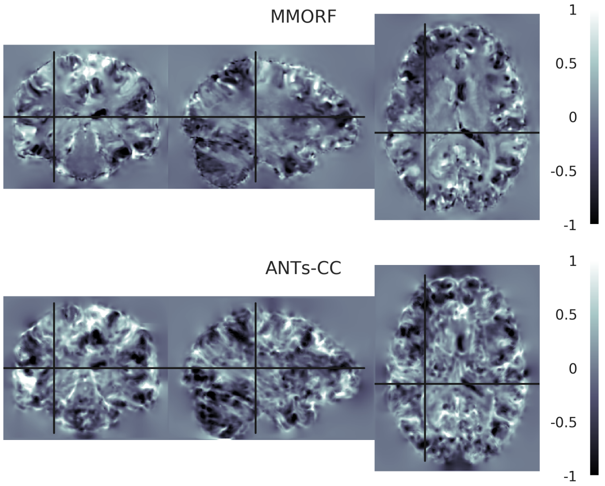

**Log-Jacobian determinant spatial maps - subject 07 to 09**

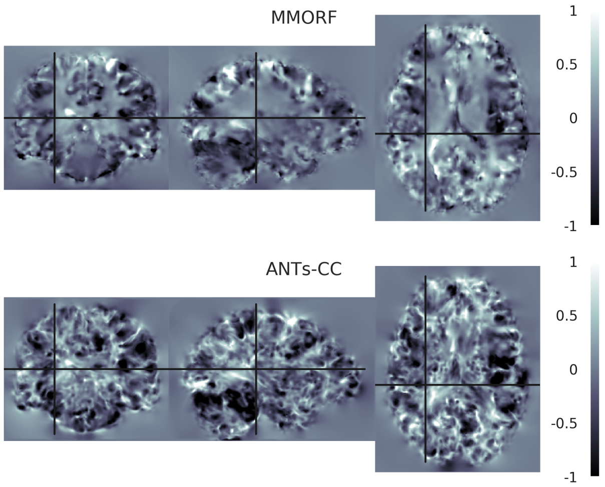


**Log-Jacobian determinant spatial maps - subject 08 to 09**

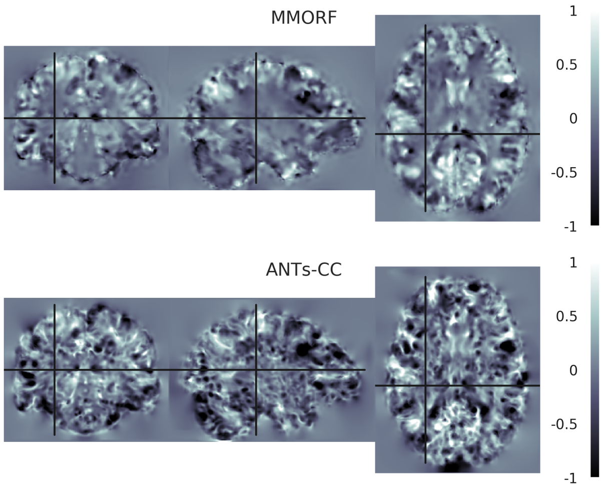


**Log-Jacobian determinant spatial maps - subject 10 to 09**

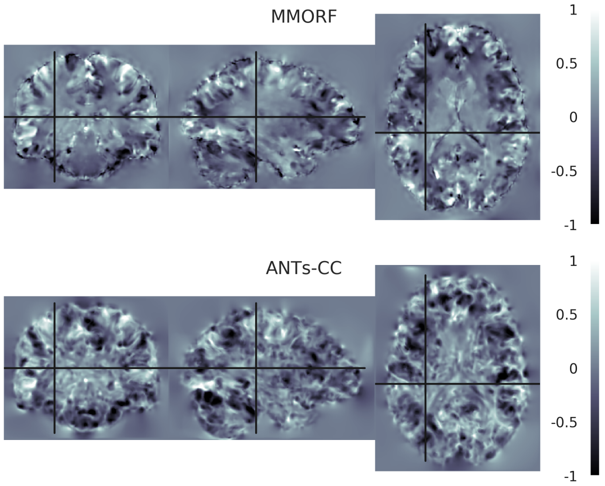


**Log-Jacobian determinant spatial maps - subject 11 to 09**

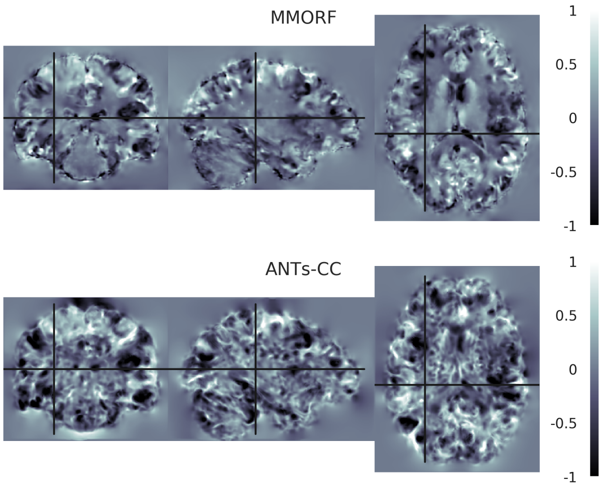


**Log-Jacobian determinant spatial maps - subject 12 to 09**

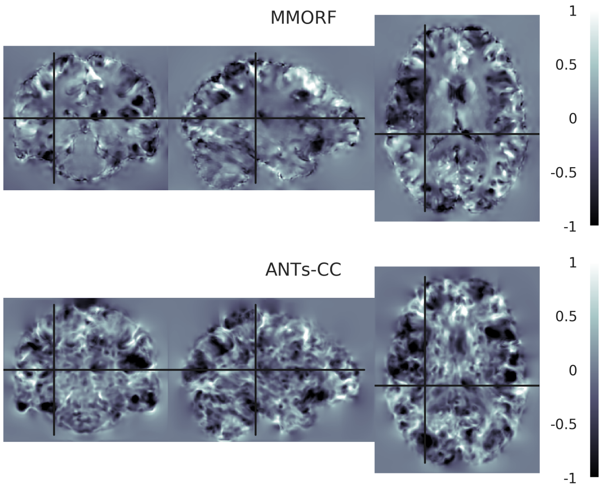


**Log-Jacobian determinant spatial maps - subject 13 to 09**

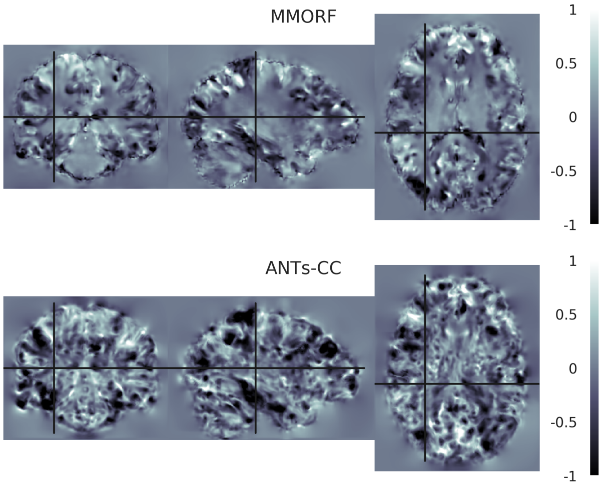

**Log-Jacobian determinant spatial maps - subject 14 to 09**

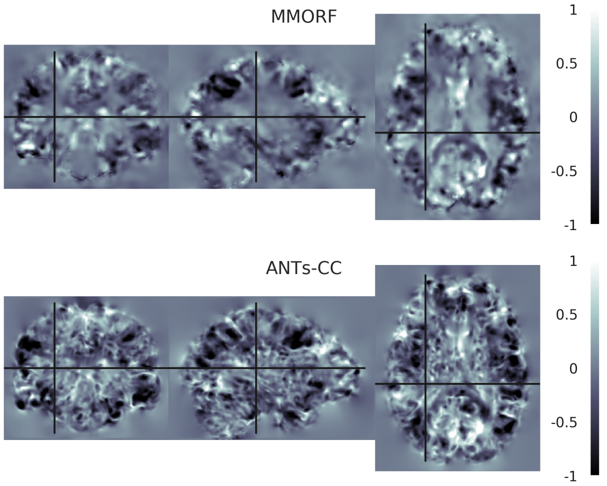


**Log-Jacobian determinant spatial maps - subject 15 to 09**

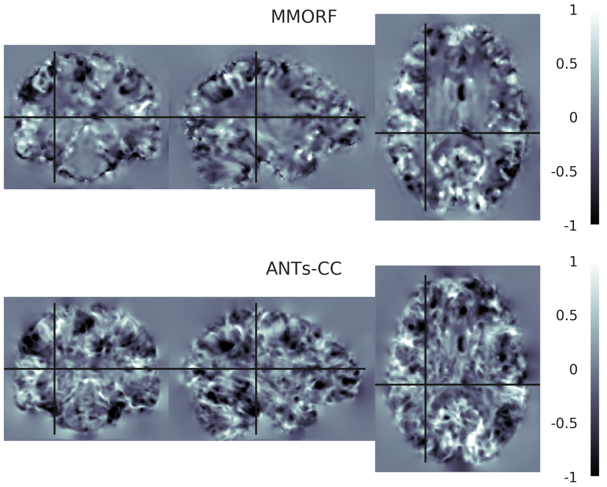


**Log-Jacobian determinant spatial maps - subject 16 to 09**

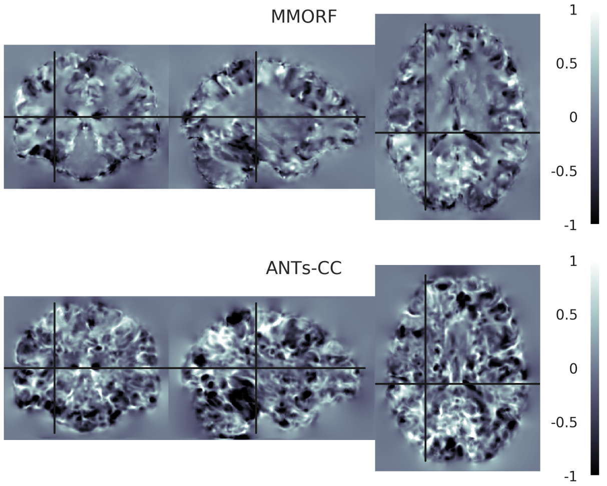

#### 10. Reference Subject 10

**Log-Jacobian determinant spatial maps - subject 01 to 10**

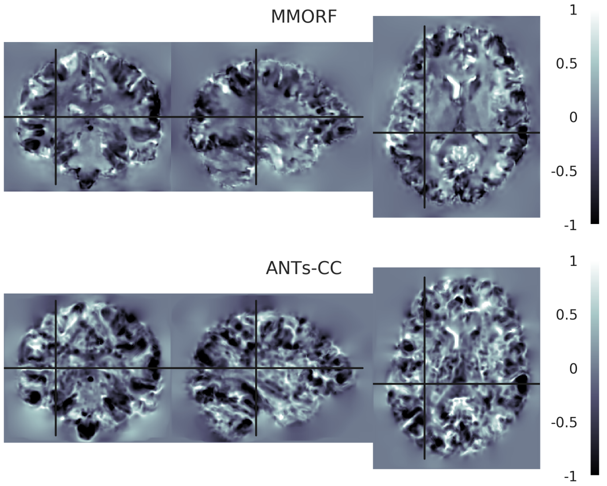


**Log-Jacobian determinant spatial maps - subject 02 to 10**

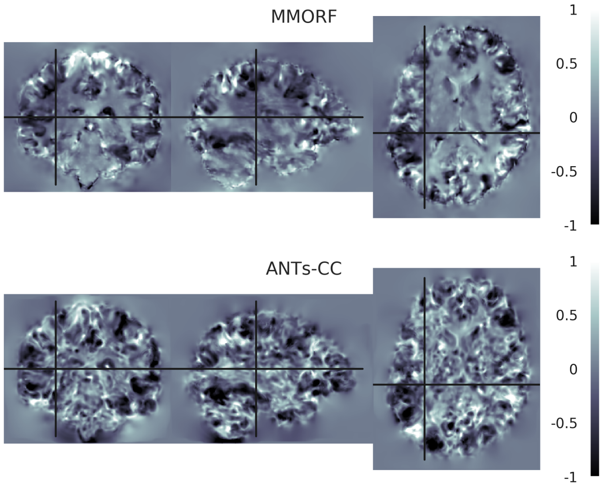


**Log-Jacobian determinant spatial maps - subject 03 to 10**

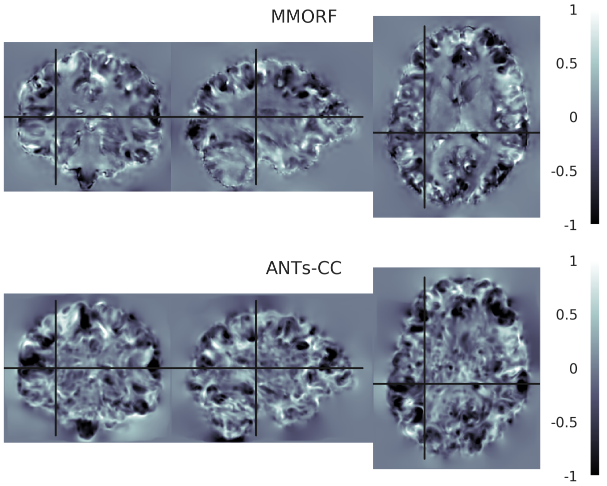


**Log-Jacobian determinant spatial maps - subject 04 to 10**

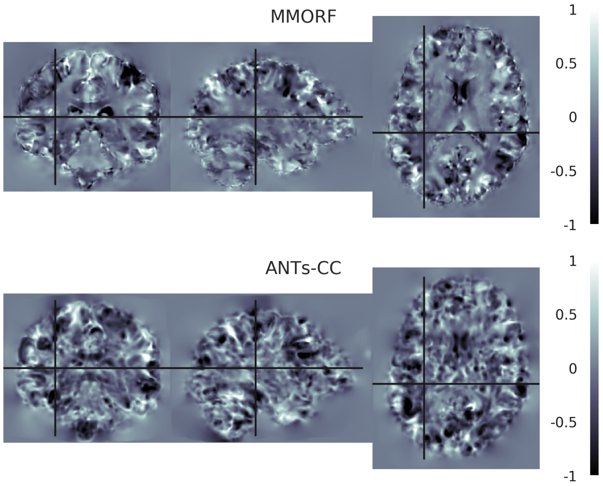


**Log-Jacobian determinant spatial maps - subject 05 to 10**

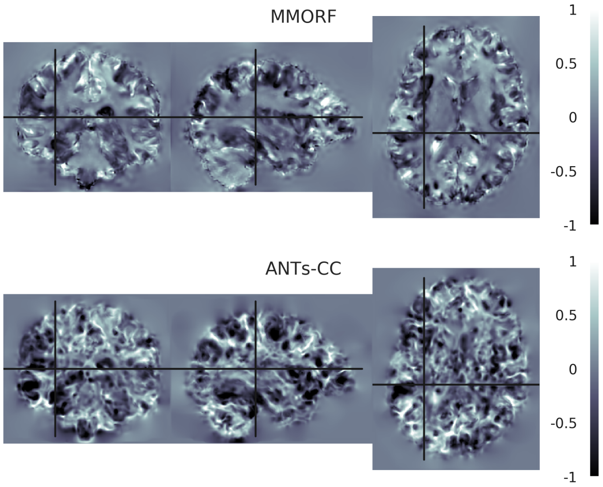


**Log-Jacobian determinant spatial maps - subject 06 to 10**

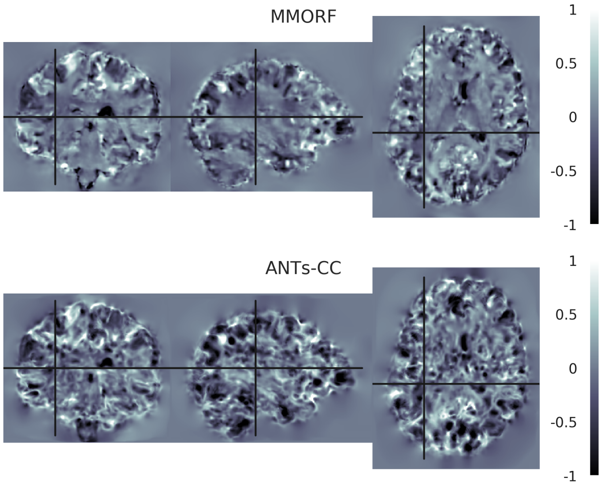

**Log-Jacobian determinant spatial maps - subject 07 to 10**

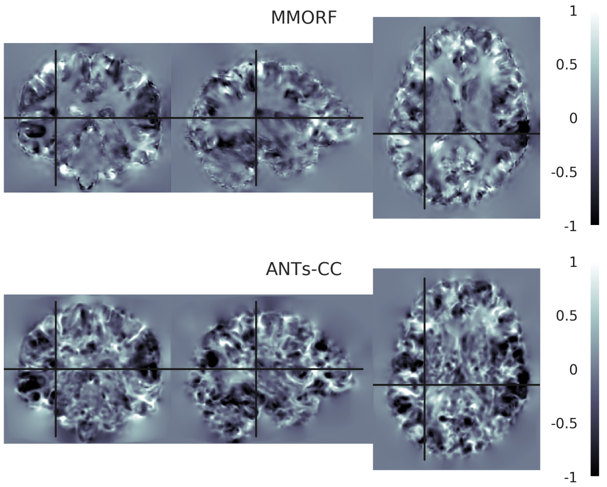


**Log-Jacobian determinant spatial maps - subject 08 to 10**

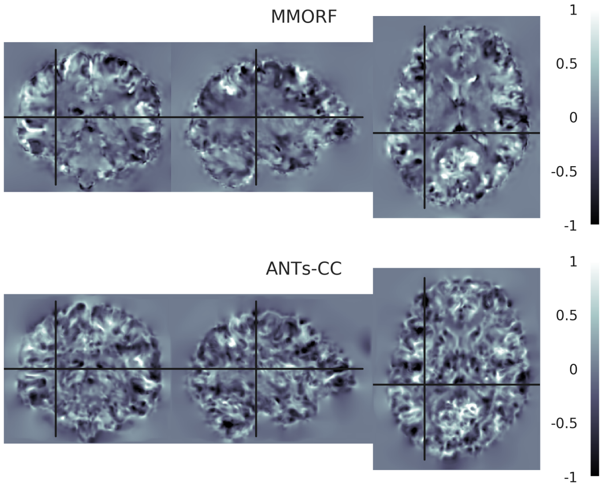


**Log-Jacobian determinant spatial maps - subject 09 to 10**

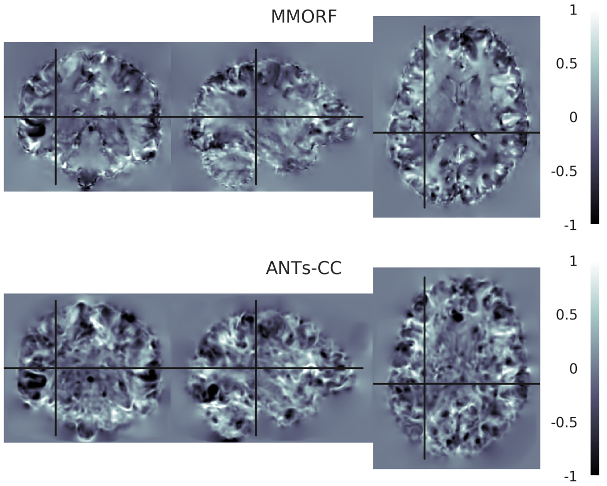


**Log-Jacobian determinant spatial maps - subject 11 to 10**

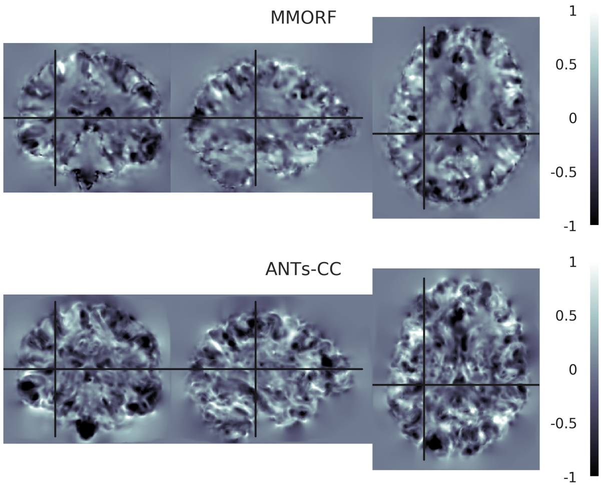


**Log-Jacobian determinant spatial maps - subject 12 to 10**

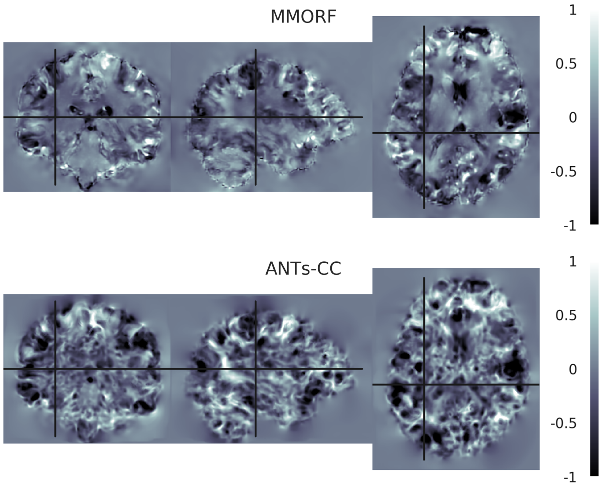


**Log-Jacobian determinant spatial maps - subject 13 to 10**

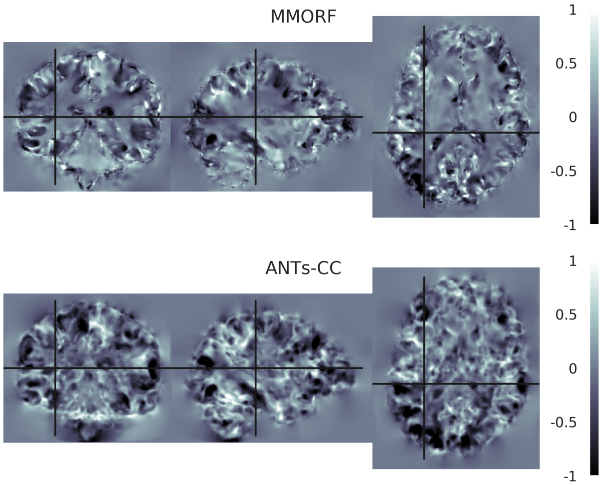

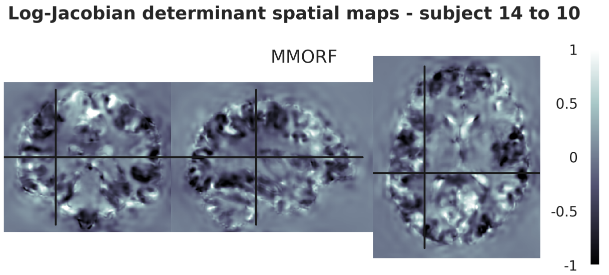
Log-Jacobian determinant spatial maps - subject 14 to 10
MMORF
ANTs-CC
1
0.5
0
-0.5
-1

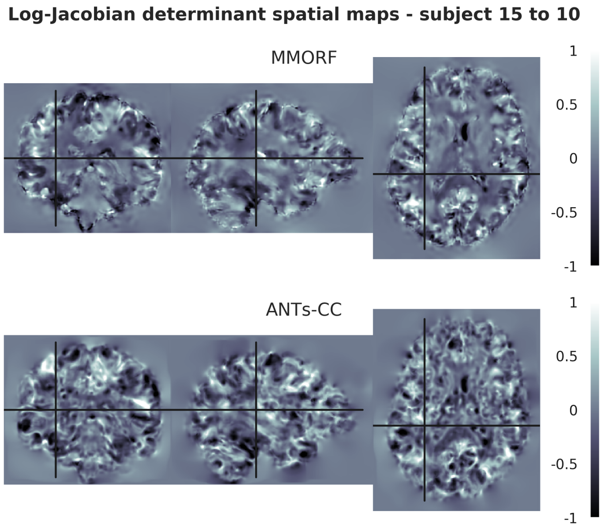
Log-Jacobian determinant spatial maps - subject 15 to 10
MMORF
ANTs-CC
1
0.5
0
-0.5
-1

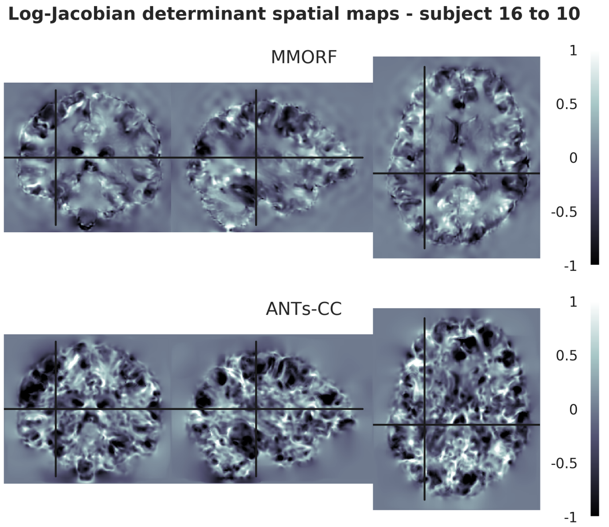
Log-Jacobian determinant spatial maps - subject 16 to 10
MMORF
ANTs-CC
1
0.5
0
-0.5
-1

#### 11. Reference Subject 11

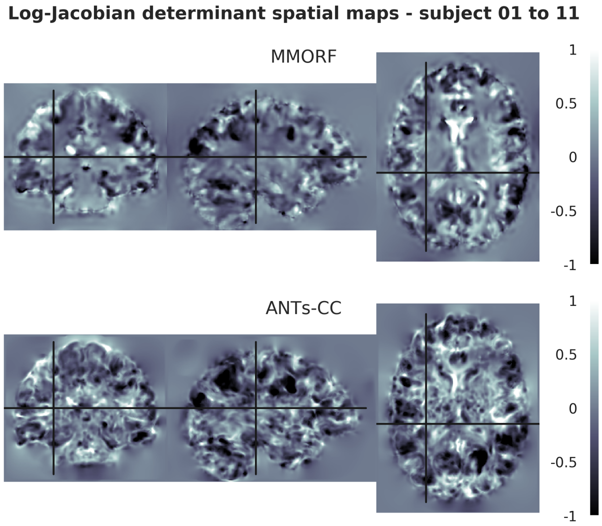


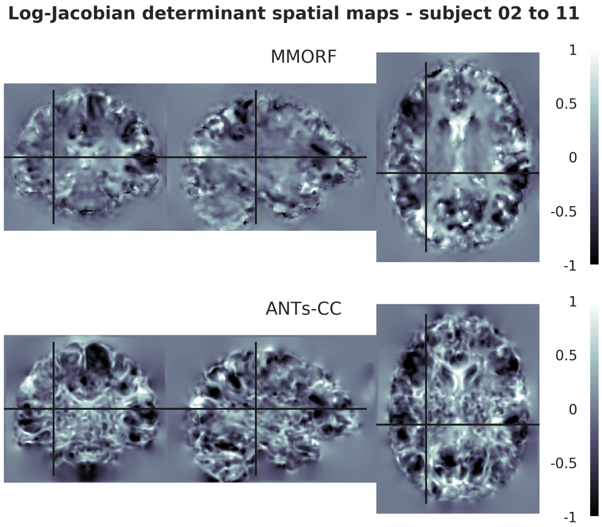


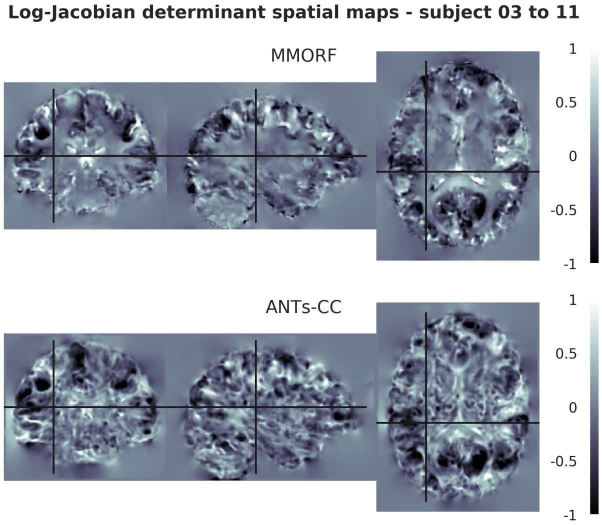


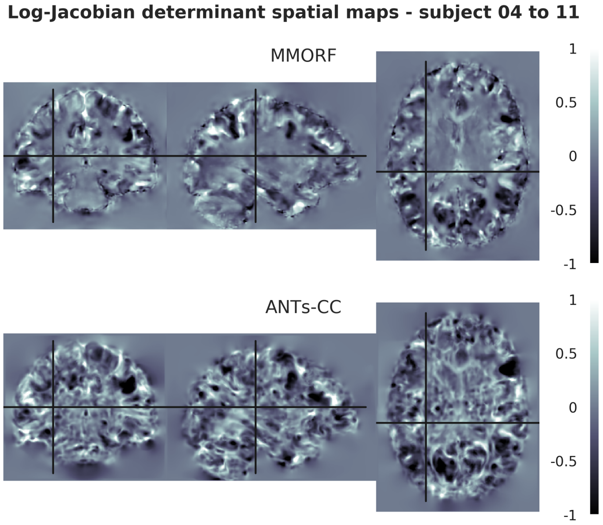


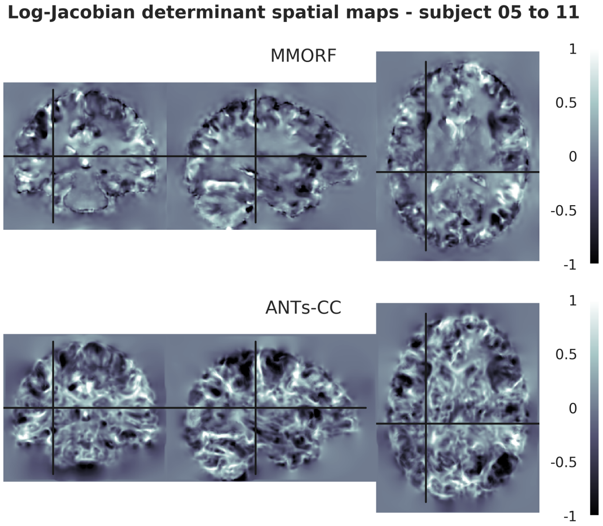


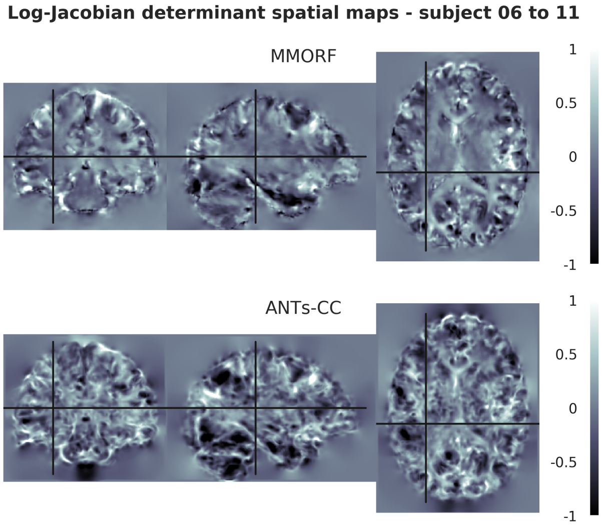

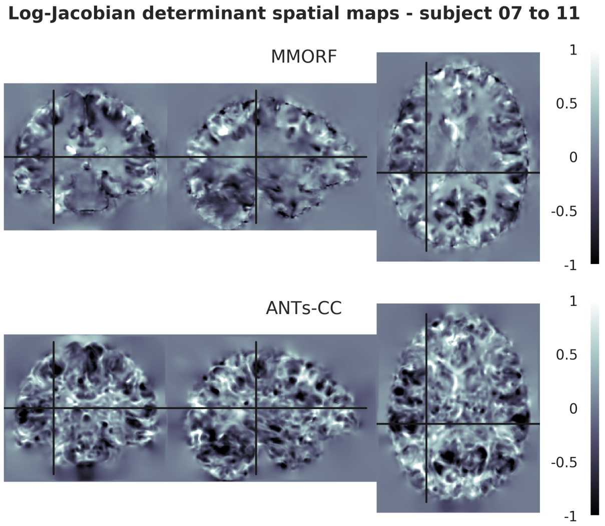
Log-Jacobian determinant spatial maps - subject 07 to 11
MMORF
ANTs-CC
1
0.5
0
-0.5
-1

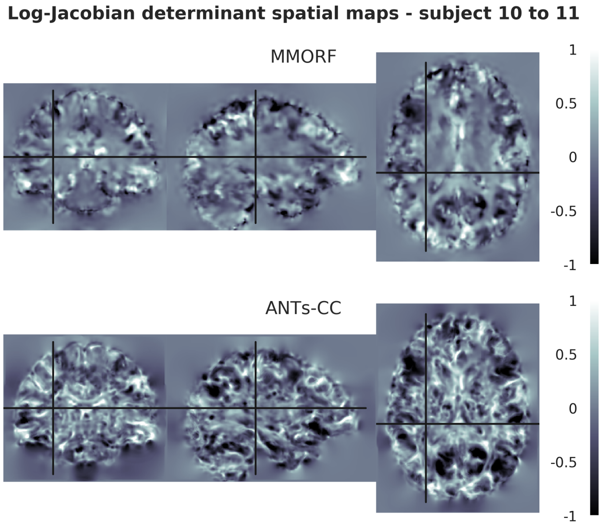
Log-Jacobian determinant spatial maps - subject 10 to 11
MMORF
ANTs-CC
1
0.5
0
-0.5
-1

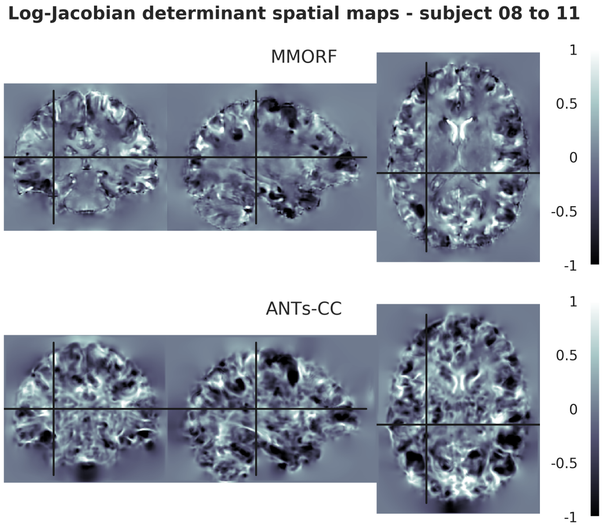
Log-Jacobian determinant spatial maps - subject 08 to 11
MMORF
ANTs-CC
1
0.5
0
-0.5
-1

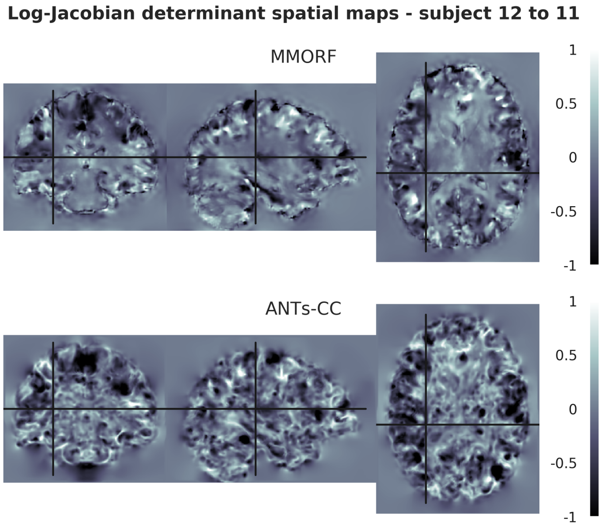
Log-Jacobian determinant spatial maps - subject 12 to 11
MMORF
ANTs-CC
1
0.5
0
-0.5
-1

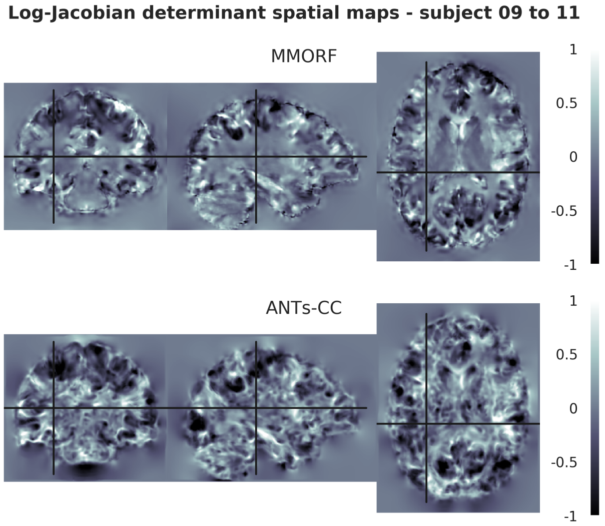
Log-Jacobian determinant spatial maps - subject 09 to 11
MMORF
ANTs-CC
1
0.5
0
-0.5
-1

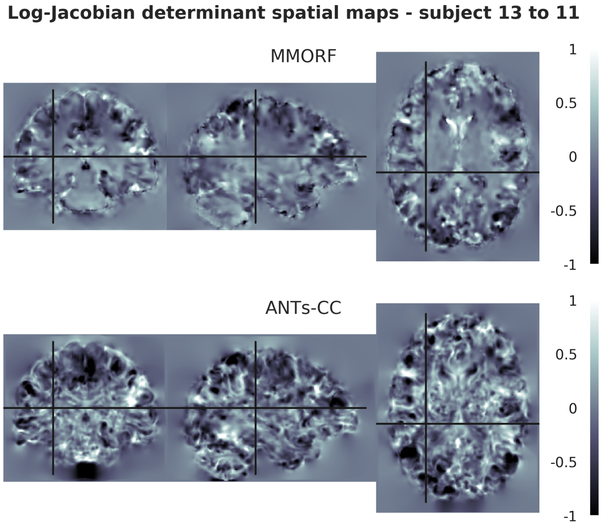
Log-Jacobian determinant spatial maps - subject 13 to 11
MMORF
ANTs-CC
1
0.5
0
-0.5
-1

**Log-Jacobian determinant spatial maps - subject 14 to 11**

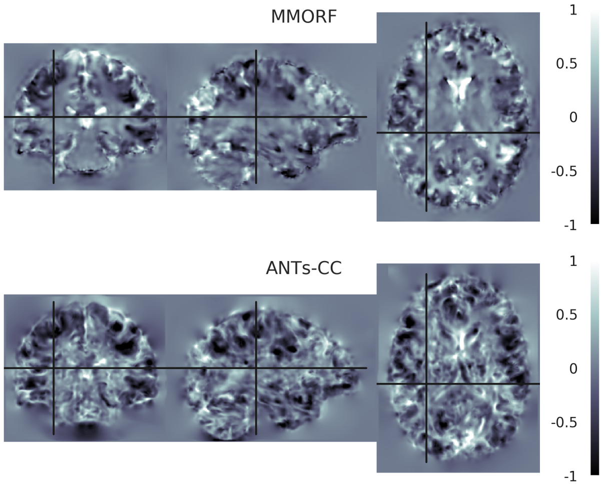


**Log-Jacobian determinant spatial maps - subject 15 to 11**

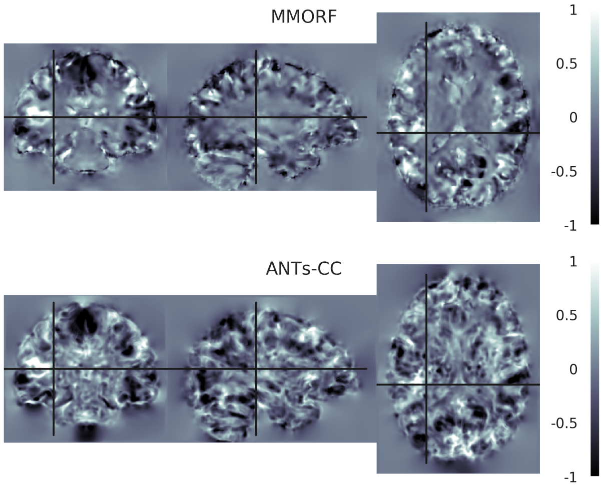


**Log-Jacobian determinant spatial maps - subject 16 to 11**

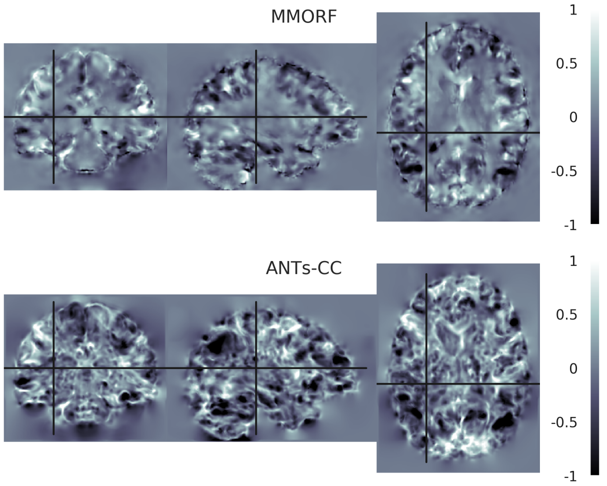

#### 12. Reference Subject 12

**Log-Jacobian determinant spatial maps - subject 01 to 12**

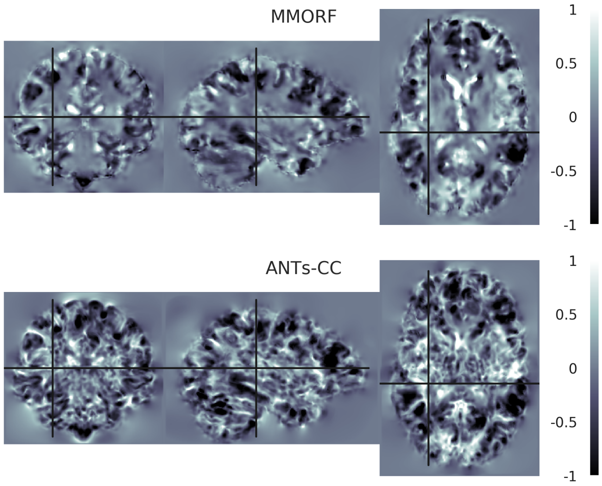


**Log-Jacobian determinant spatial maps - subject 02 to 12**

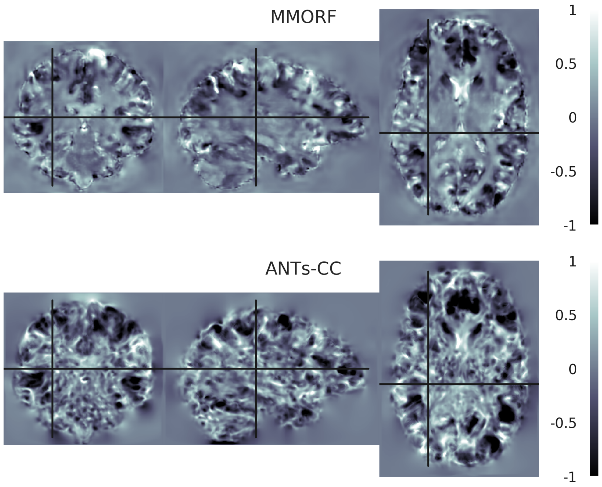


**Log-Jacobian determinant spatial maps - subject 03 to 12**

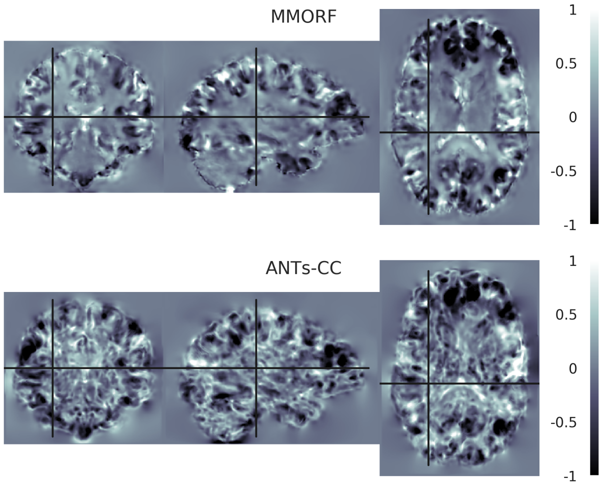


**Log-Jacobian determinant spatial maps - subject 04 to 12**

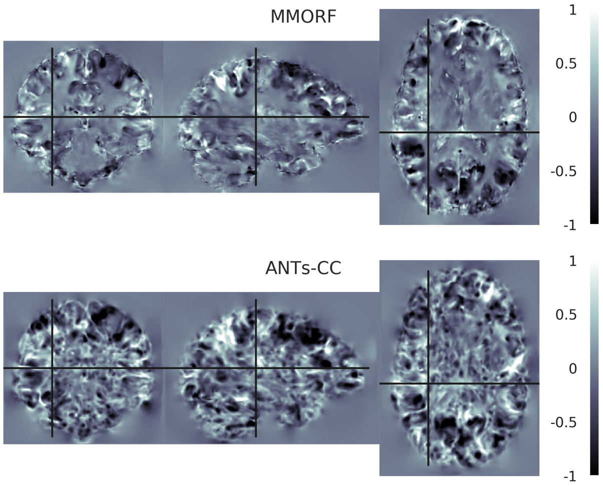


**Log-Jacobian determinant spatial maps - subject 05 to 12**

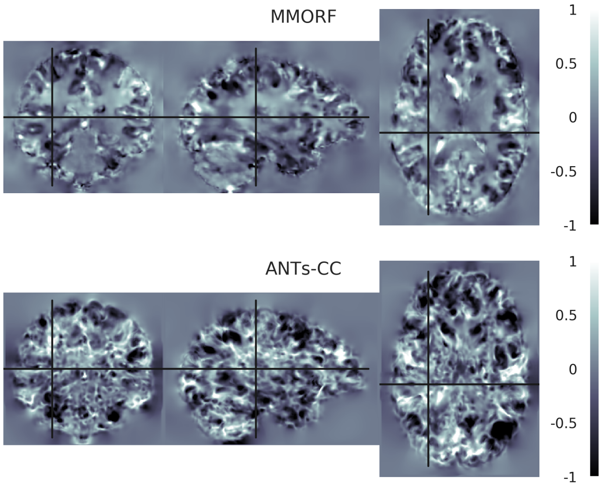


**Log-Jacobian determinant spatial maps - subject 06 to 12**

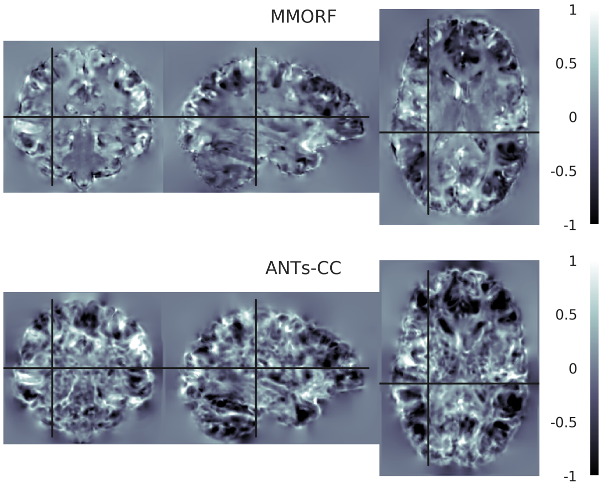

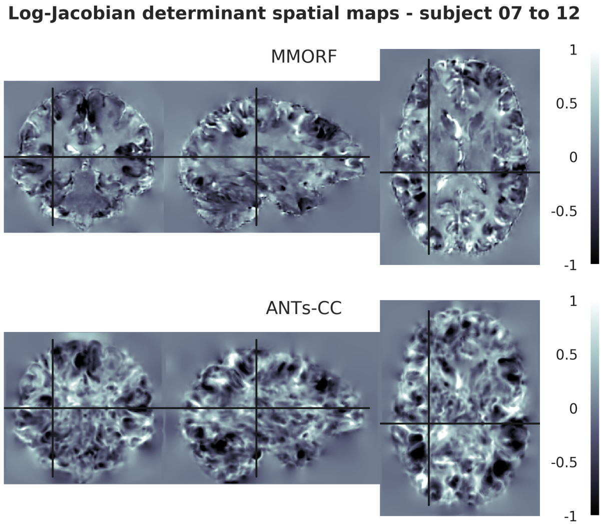
Log-Jacobian determinant spatial maps - subject 07 to 12
MMORF
ANTs-CC
1
0.5
0
-0.5
-1

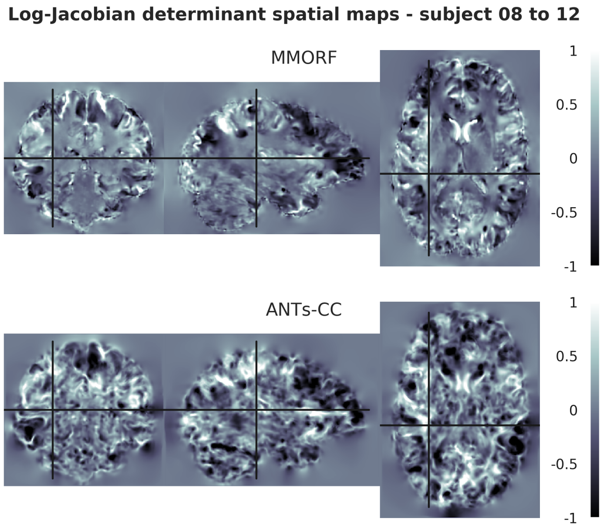
Log-Jacobian determinant spatial maps - subject 08 to 12
MMORF
ANTs-CC
1
0.5
0
-0.5
-1

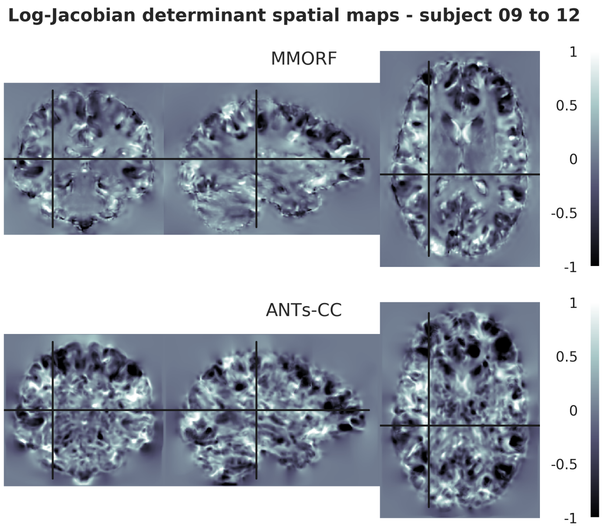
Log-Jacobian determinant spatial maps - subject 09 to 12
MMORF
ANTs-CC
1
0.5
0
-0.5
-1

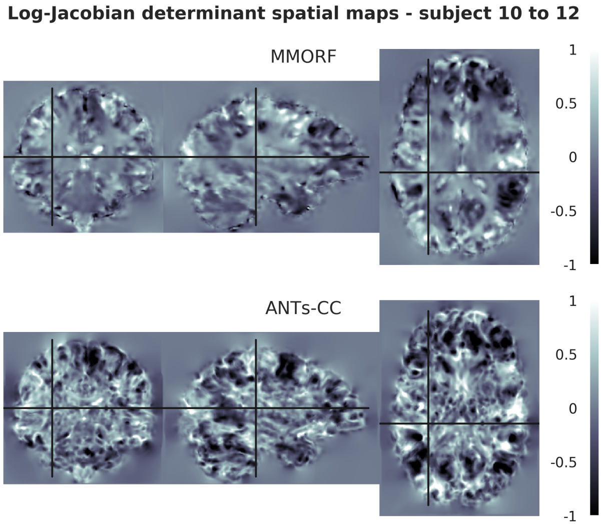
Log-Jacobian determinant spatial maps - subject 10 to 12
MMORF
ANTs-CC
1
0.5
0
-0.5
-1

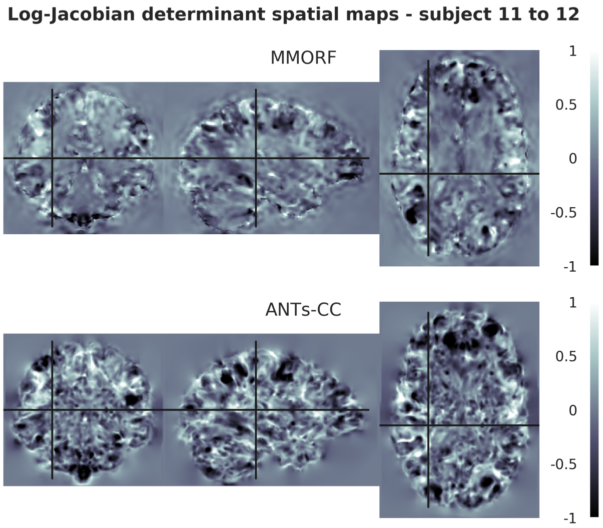
Log-Jacobian determinant spatial maps - subject 11 to 12
MMORF
ANTs-CC
1
0.5
0
-0.5
-1

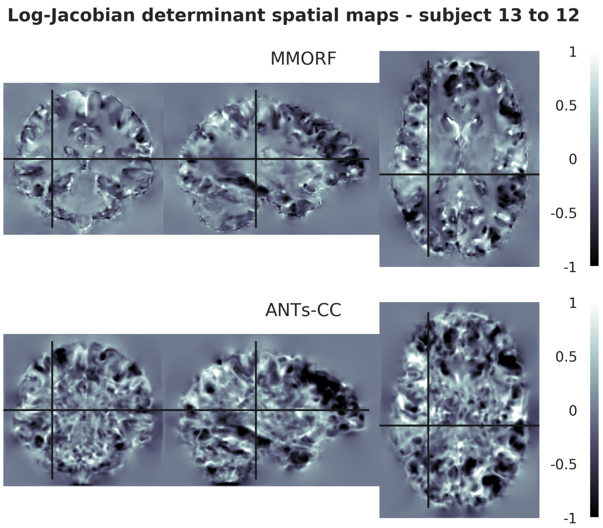
Log-Jacobian determinant spatial maps - subject 13 to 12
MMORF
ANTs-CC
1
0.5
0
-0.5
-1

**Log-Jacobian determinant spatial maps - subject 14 to 12**

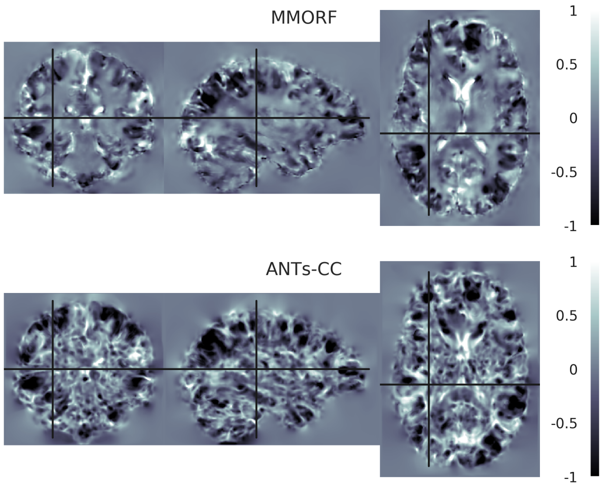


**Log-Jacobian determinant spatial maps - subject 15 to 12**

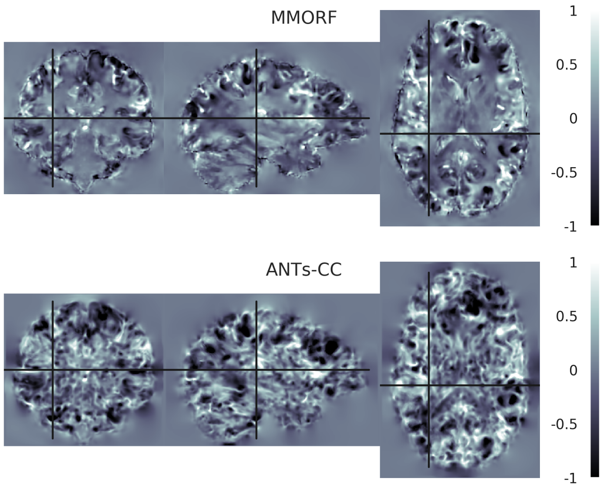


**Log-Jacobian determinant spatial maps - subject 16 to 12**

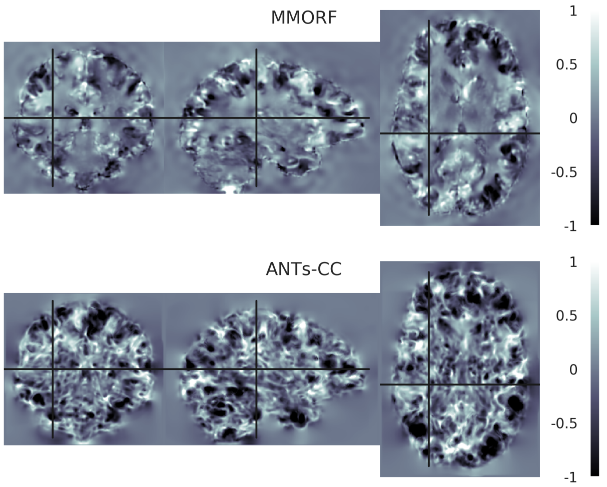

#### 13. Reference Subject 13

**Log-Jacobian determinant spatial maps - subject 01 to 13**

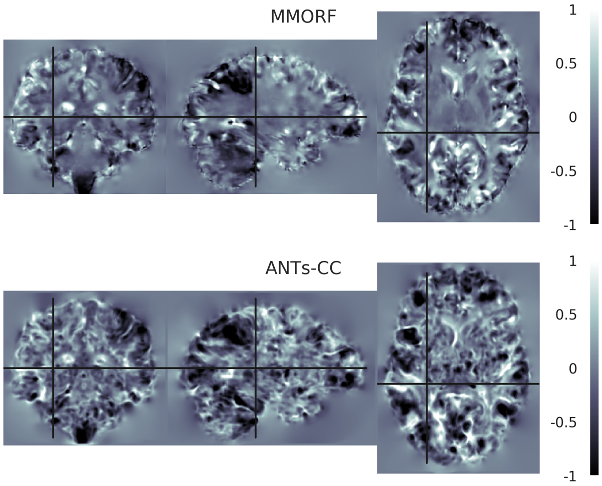


**Log-Jacobian determinant spatial maps - subject 02 to 13**

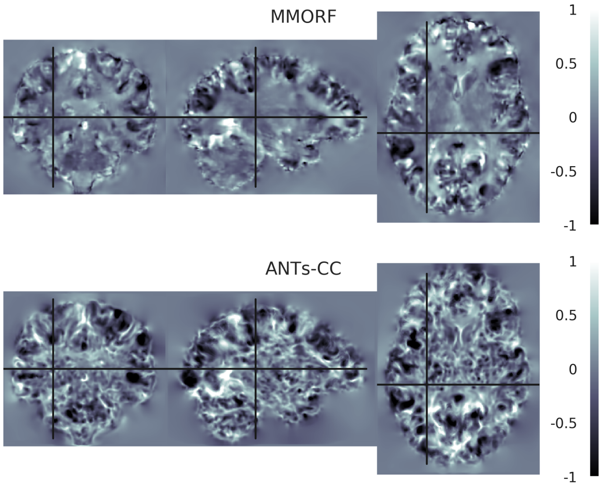


**Log-Jacobian determinant spatial maps - subject 03 to 13**

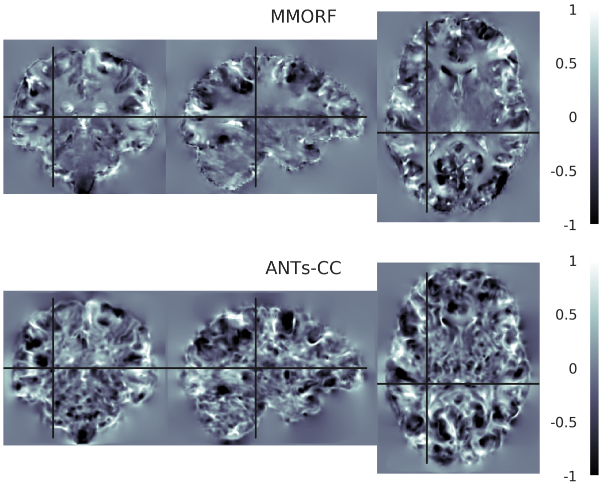


**Log-Jacobian determinant spatial maps - subject 04 to 13**

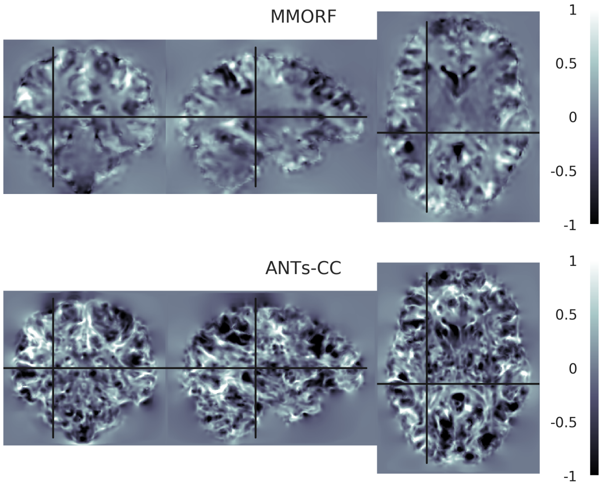


**Log-Jacobian determinant spatial maps - subject 05 to 13**

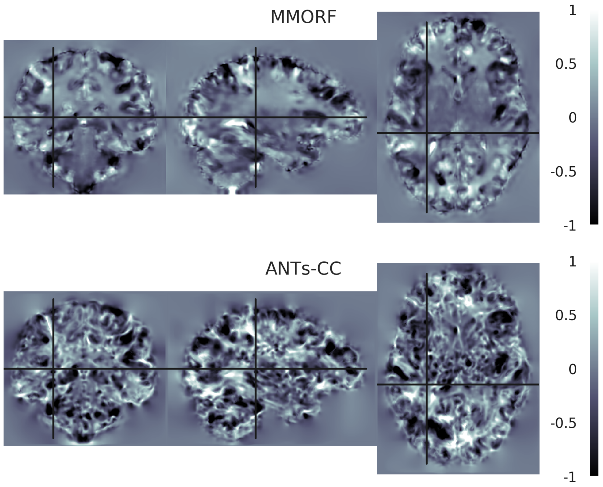


**Log-Jacobian determinant spatial maps - subject 06 to 13**

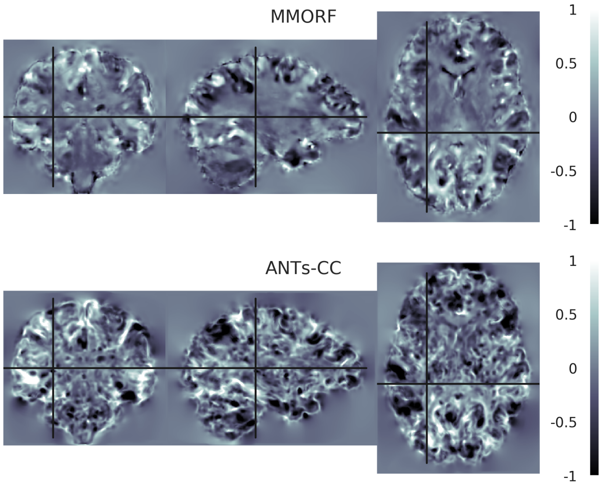

**Log-Jacobian determinant spatial maps - subject 07 to 13**

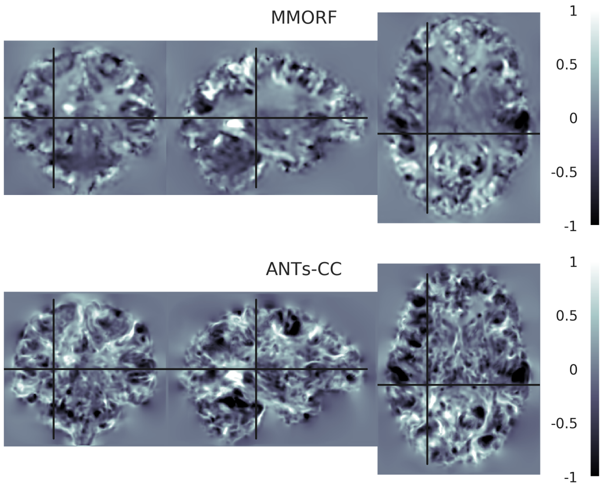


**Log-Jacobian determinant spatial maps - subject 08 to 13**

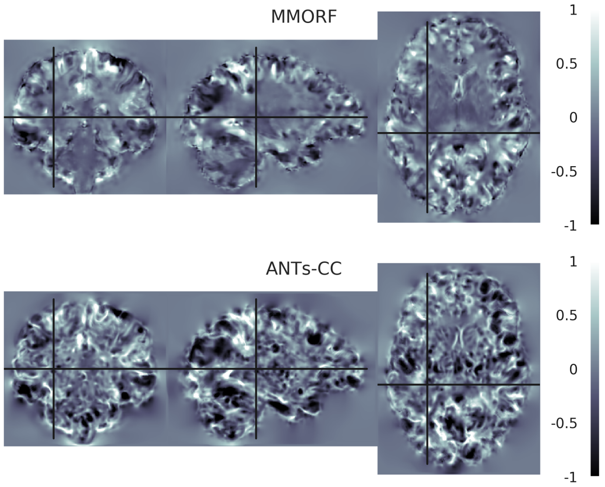


**Log-Jacobian determinant spatial maps - subject 09 to 13**

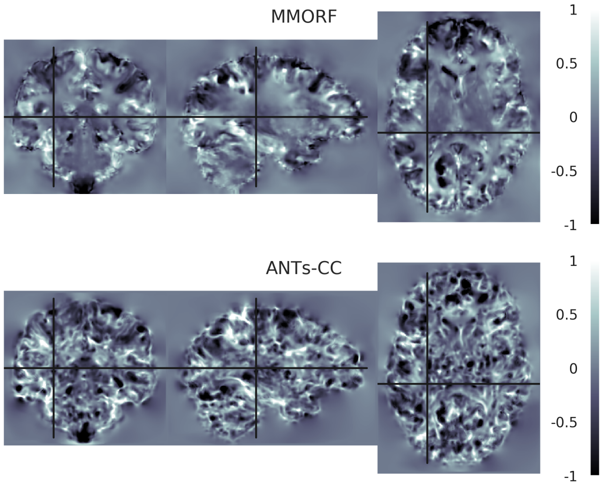


**Log-Jacobian determinant spatial maps - subject 10 to 13**

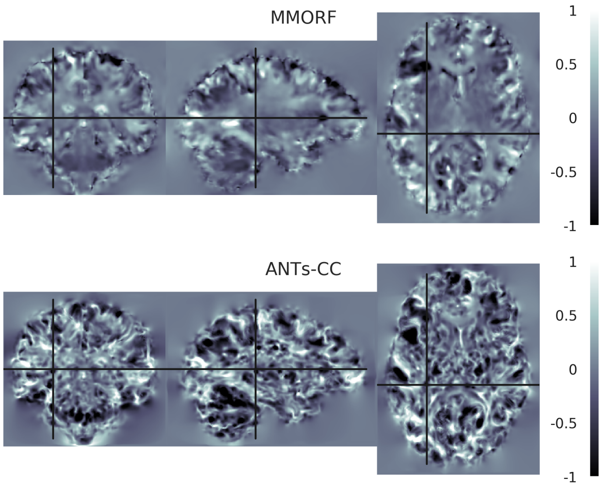


**Log-Jacobian determinant spatial maps - subject 11 to 13**

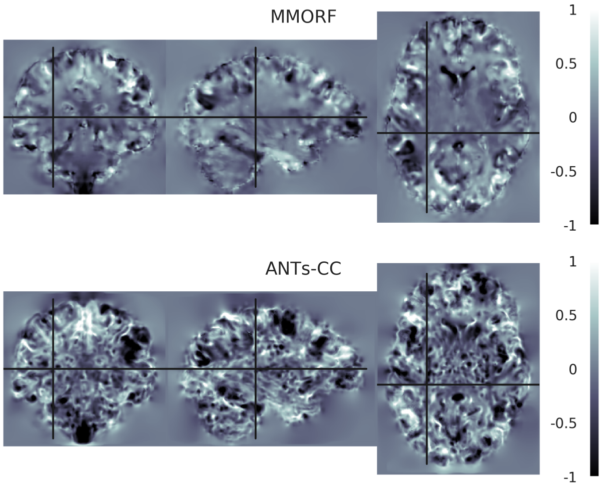


**Log-Jacobian determinant spatial maps - subject 12 to 13**

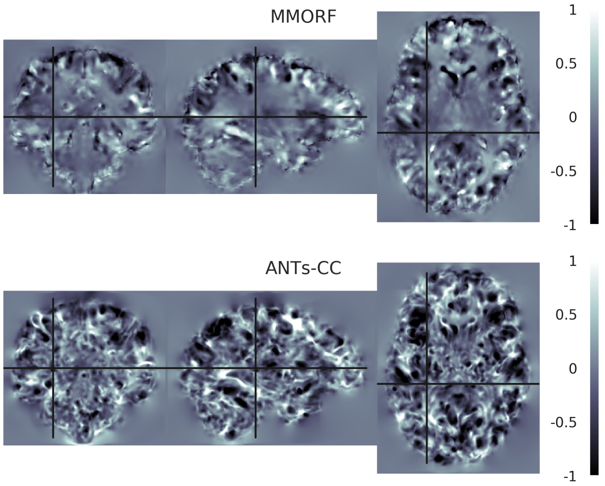

**Log-Jacobian determinant spatial maps - subject 14 to 13**

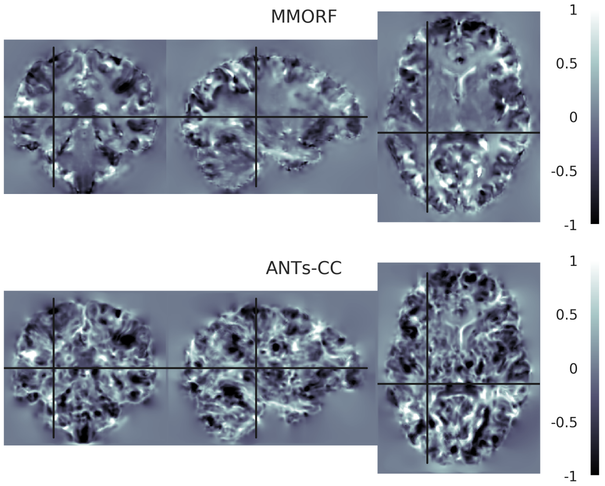


**Log-Jacobian determinant spatial maps - subject 15 to 13**

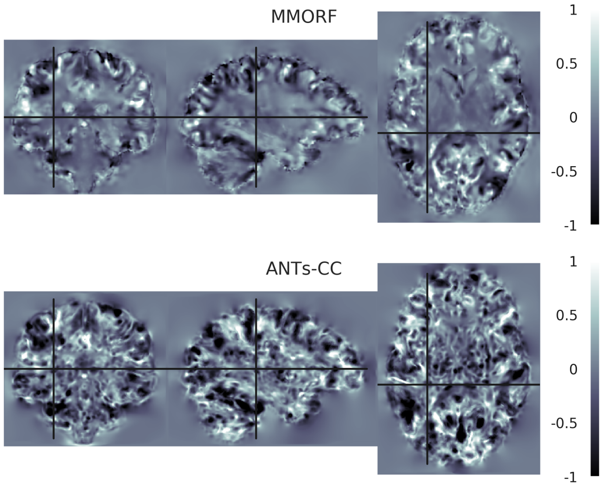


**Log-Jacobian determinant spatial maps - subject 16 to 13**

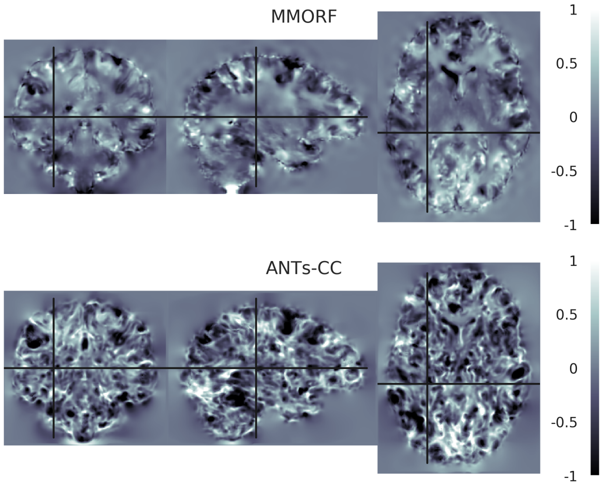

#### 14. Reference Subject 14

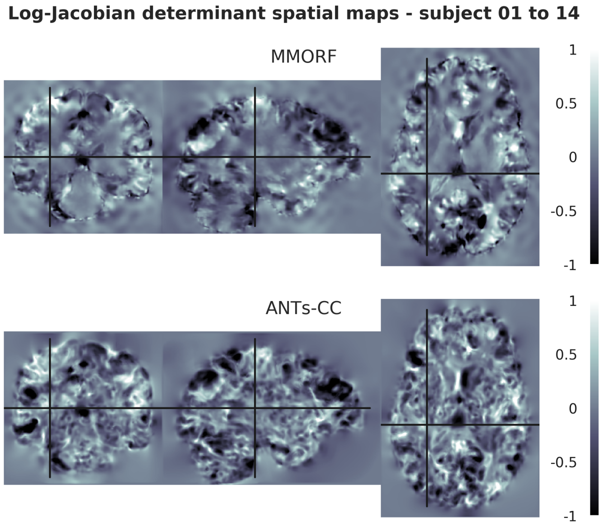


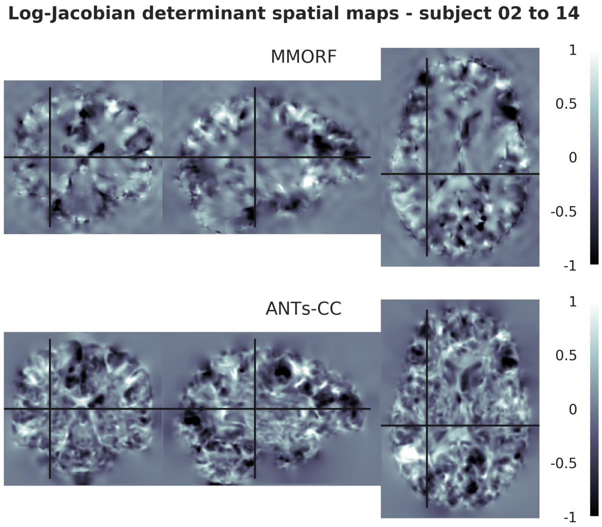


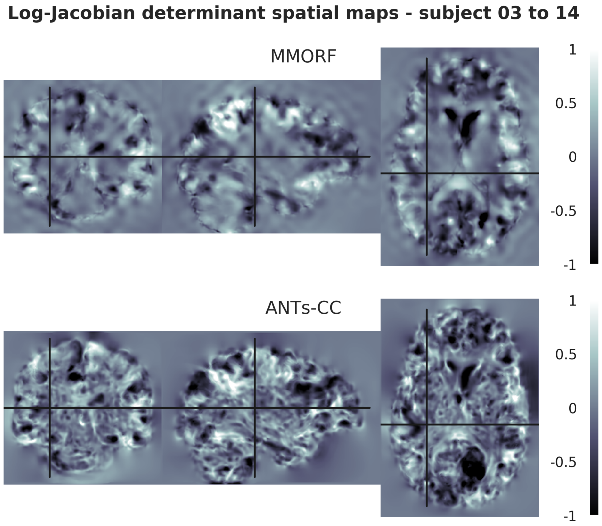


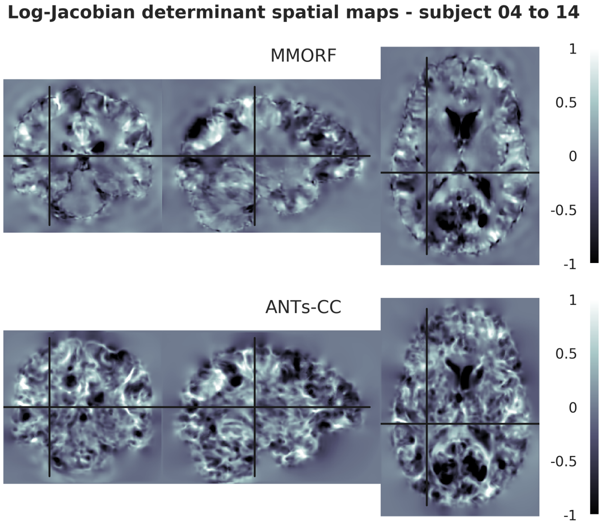


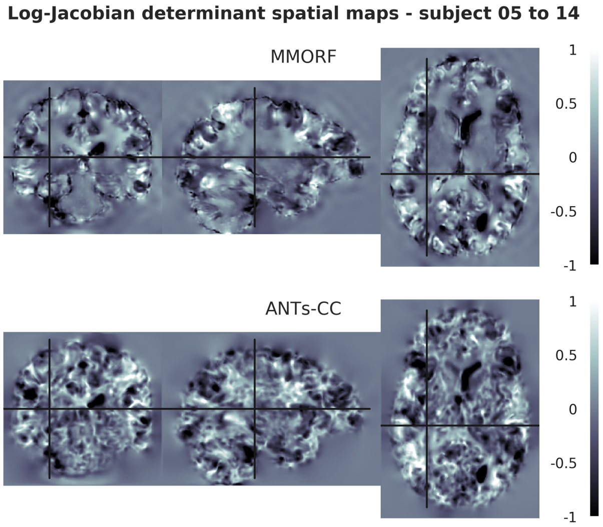


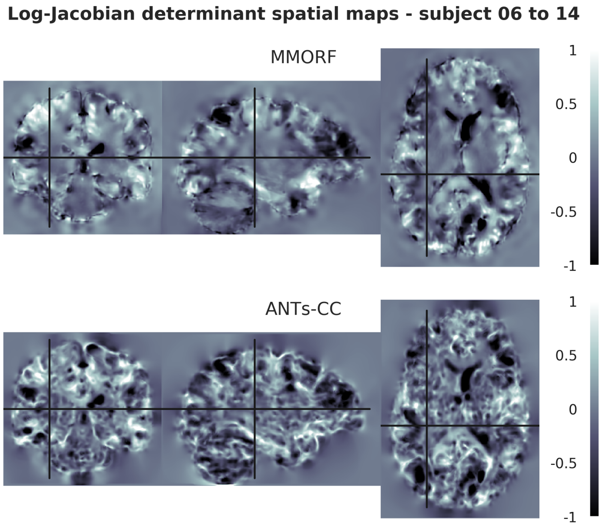

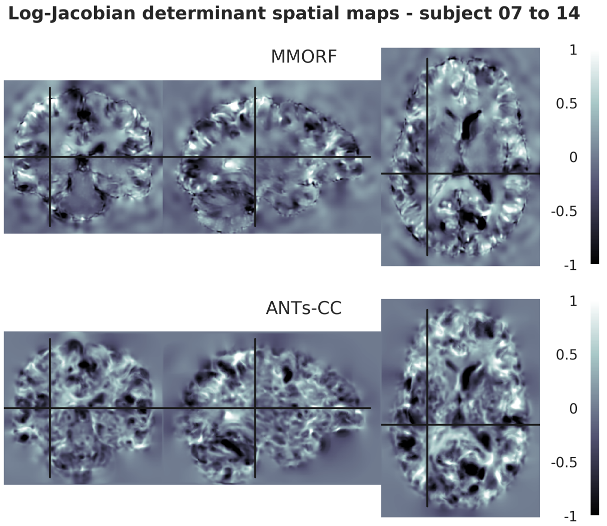
Log-Jacobian determinant spatial maps - subject 07 to 14
MMORF
ANTs-CC
1
0.5
0
-0.5
-1

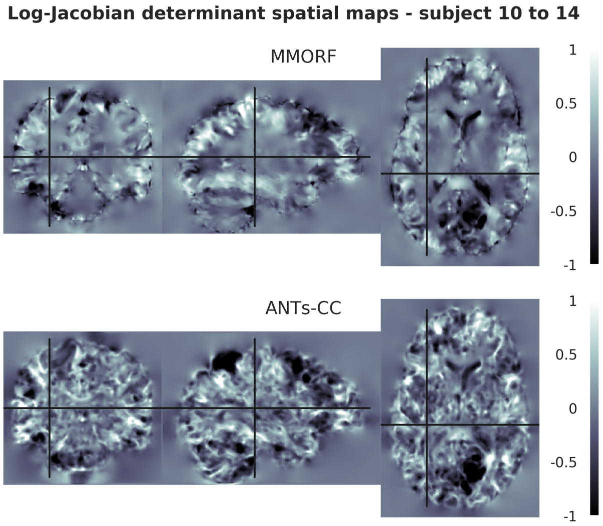
Log-Jacobian determinant spatial maps - subject 10 to 14
MMORF
ANTs-CC
1
0.5
0
-0.5
-1

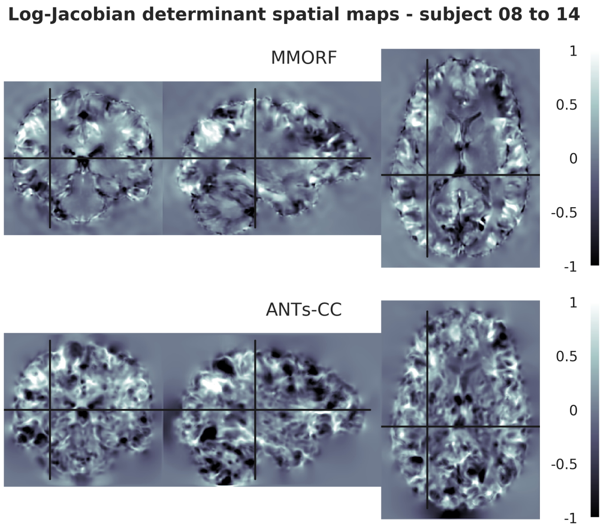
Log-Jacobian determinant spatial maps - subject 08 to 14
MMORF
ANTs-CC
1
0.5
0
-0.5
-1

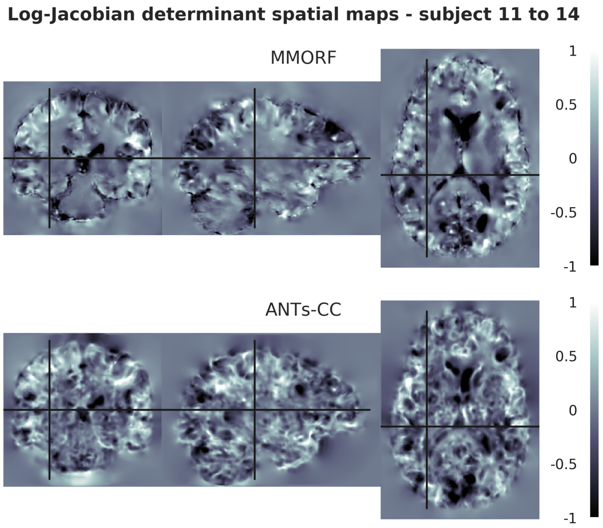
Log-Jacobian determinant spatial maps - subject 11 to 14
MMORF
ANTs-CC
1
0.5
0
-0.5
-1

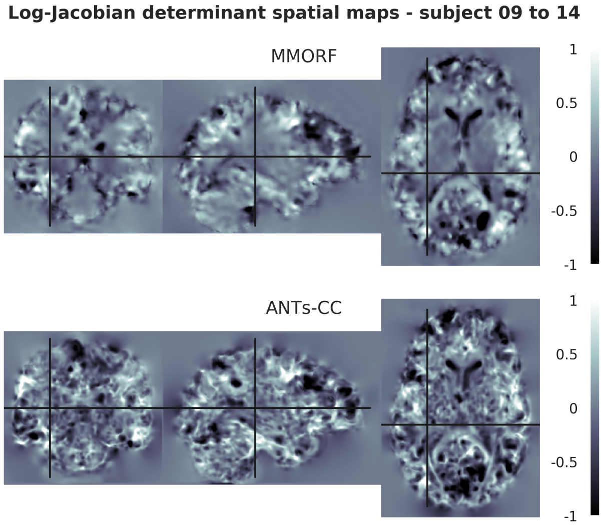
Log-Jacobian determinant spatial maps - subject 09 to 14
MMORF
ANTs-CC
1
0.5
0
-0.5
-1

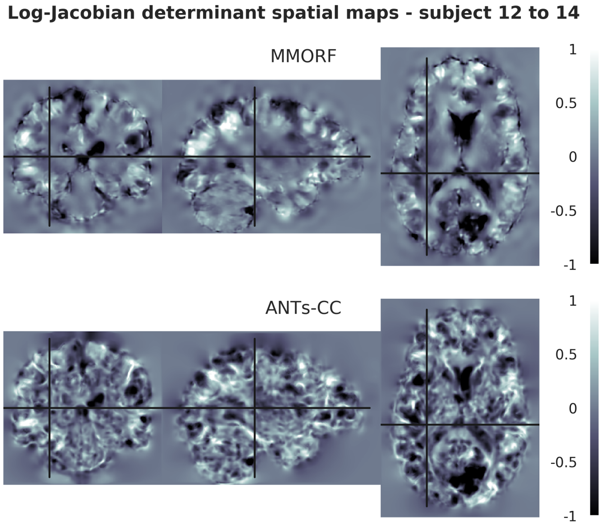
Log-Jacobian determinant spatial maps - subject 12 to 14
MMORF
ANTs-CC
1
0.5
0
-0.5
-1

**Log-Jacobian determinant spatial maps - subject 13 to 14**

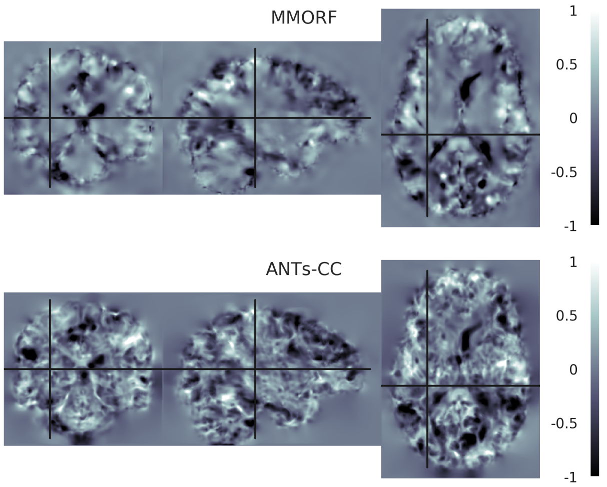


**Log-Jacobian determinant spatial maps - subject 15 to 14**

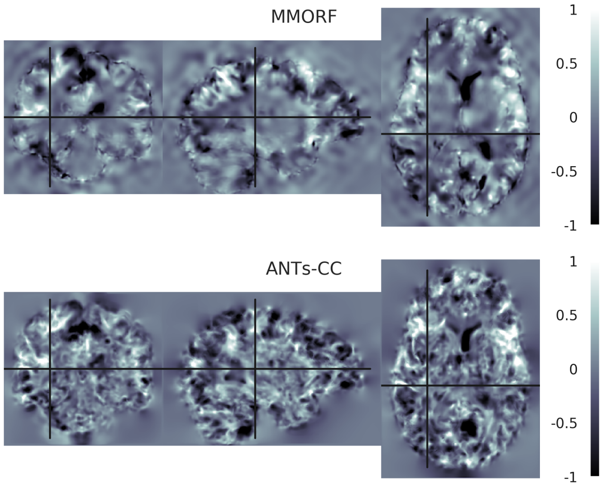


**Log-Jacobian determinant spatial maps - subject 16 to 14**

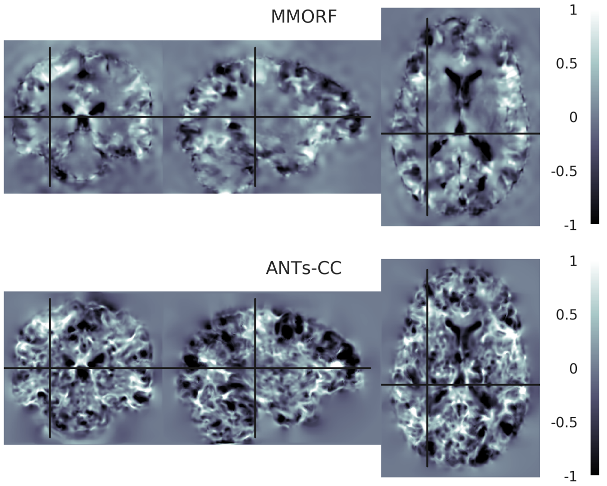

#### 15. Reference Subject 15

**Log-Jacobian determinant spatial maps - subject 01 to 15**

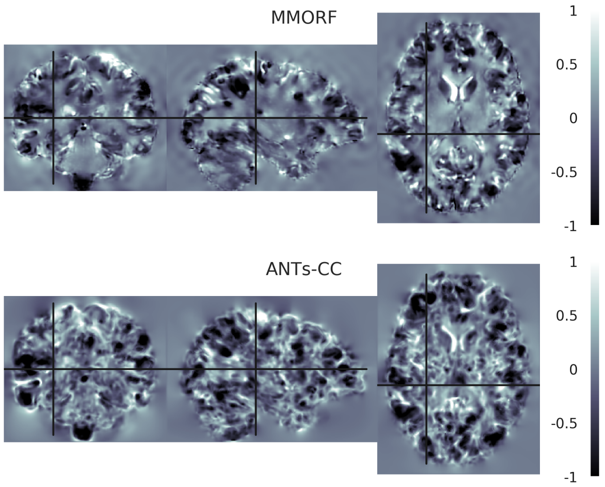


**Log-Jacobian determinant spatial maps - subject 02 to 15**

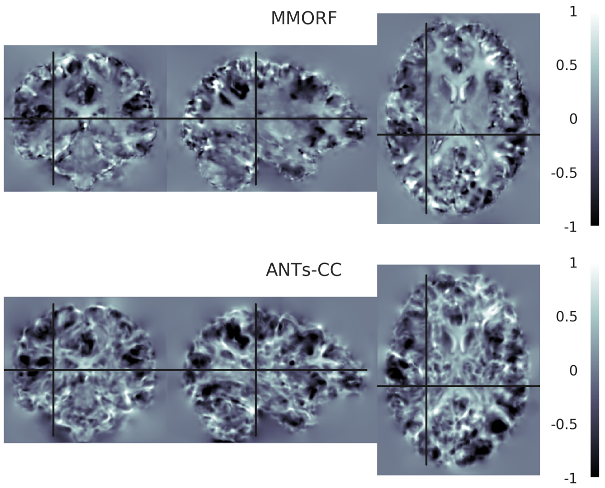


**Log-Jacobian determinant spatial maps - subject 03 to 15**

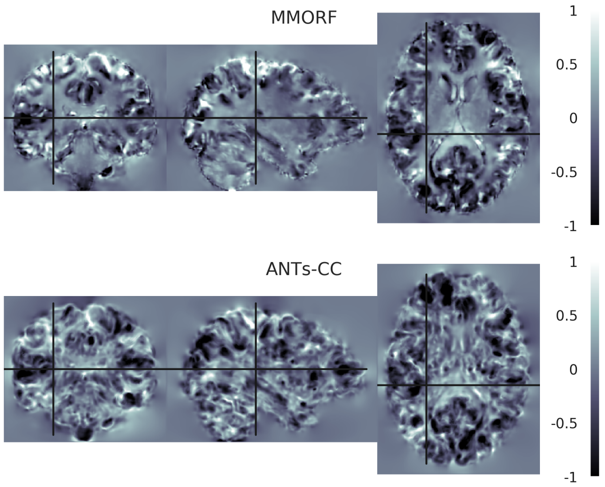


**Log-Jacobian determinant spatial maps - subject 04 to 15**

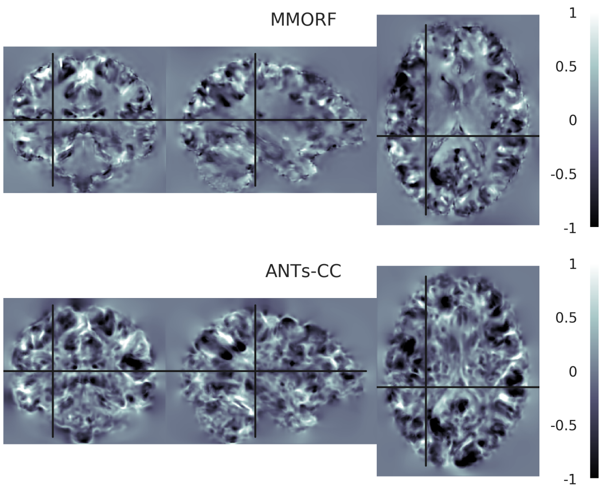


**Log-Jacobian determinant spatial maps - subject 05 to 15**

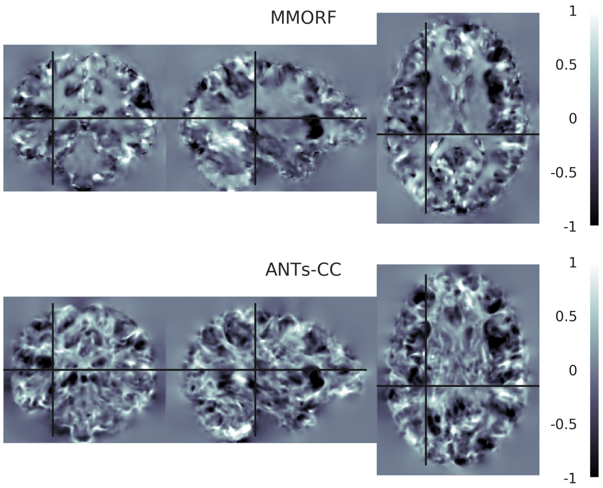


**Log-Jacobian determinant spatial maps - subject 06 to 15**

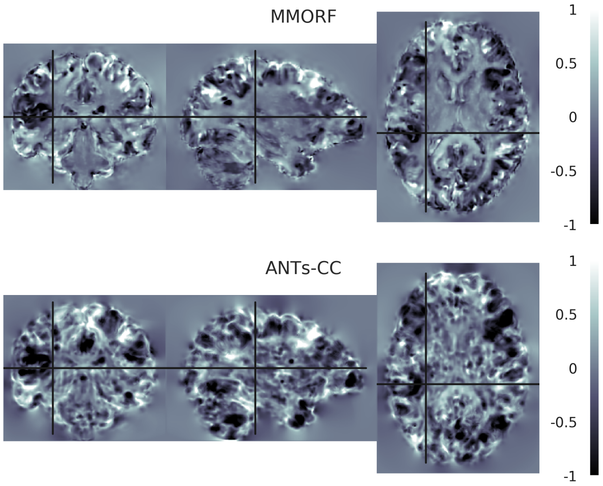

**Log-Jacobian determinant spatial maps - subject 07 to 15**

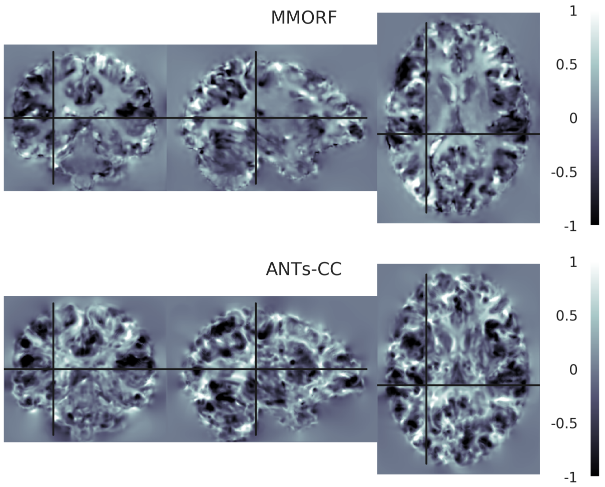


**Log-Jacobian determinant spatial maps - subject 08 to 15**

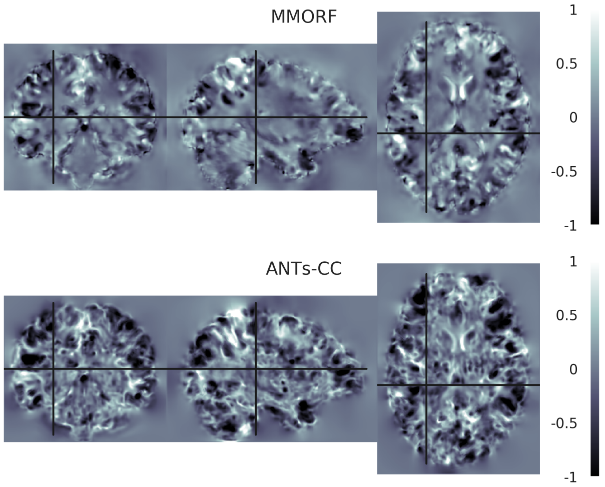


**Log-Jacobian determinant spatial maps - subject 09 to 15**

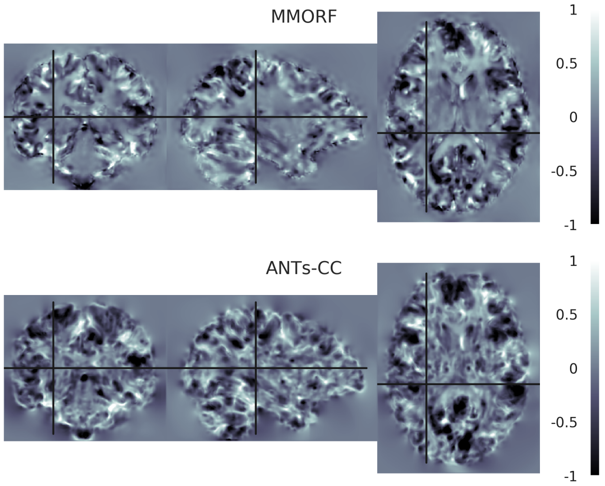


**Log-Jacobian determinant spatial maps - subject 10 to 15**

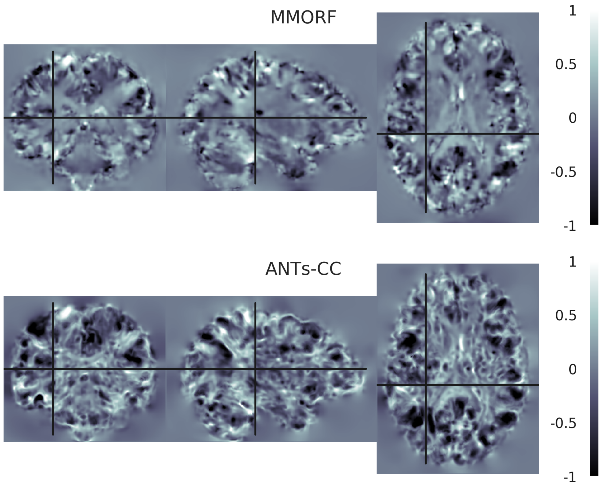


**Log-Jacobian determinant spatial maps - subject 11 to 15**

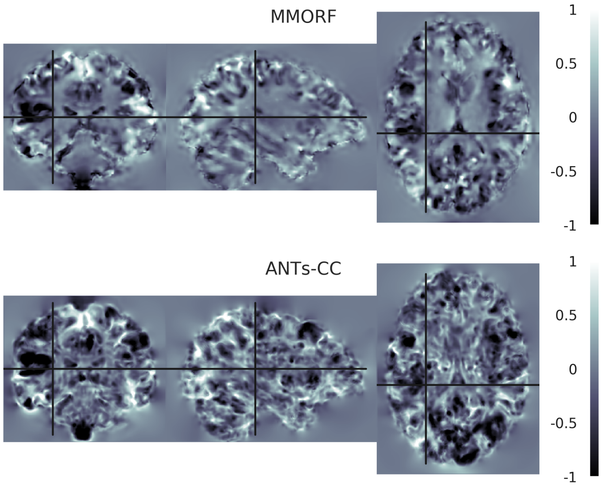


**Log-Jacobian determinant spatial maps - subject 12 to 15**

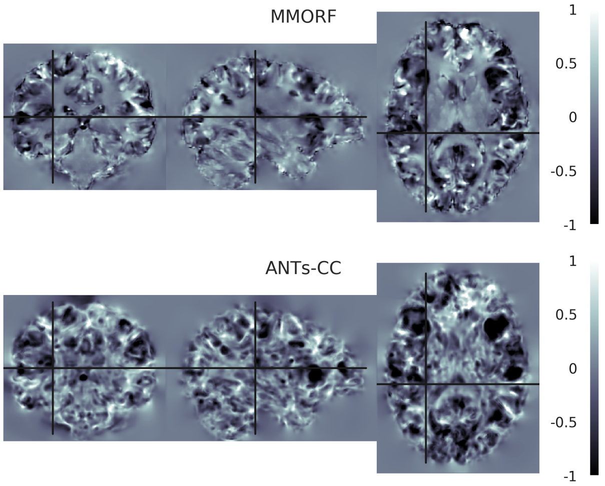

**Log-Jacobian determinant spatial maps - subject 13 to 15**

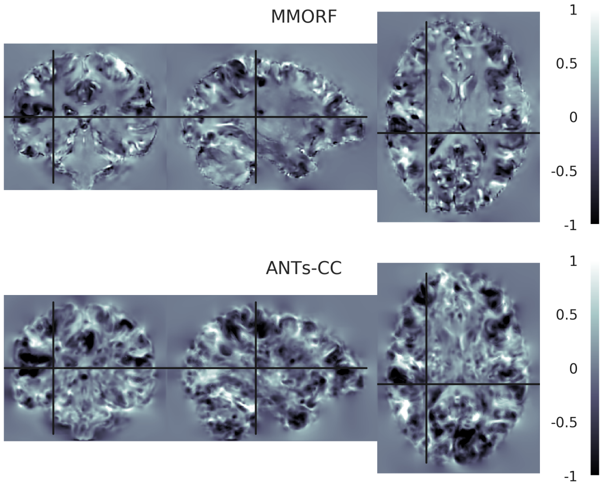


**Log-Jacobian determinant spatial maps - subject 14 to 15**

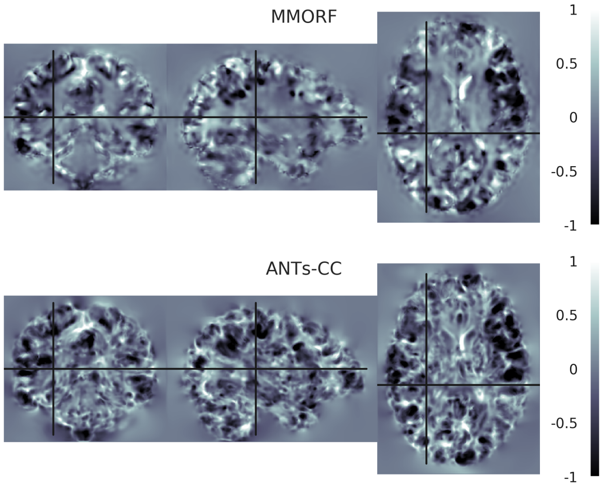


**Log-Jacobian determinant spatial maps - subject 16 to 15**

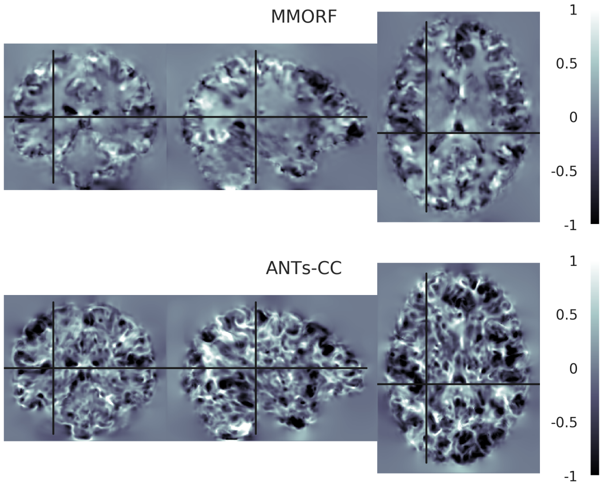

#### 16. Reference Subject 16

**Log-Jacobian determinant spatial maps - subject 01 to 16**

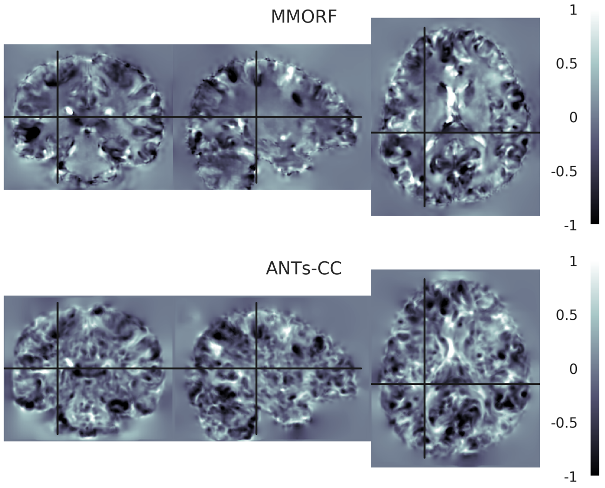


**Log-Jacobian determinant spatial maps - subject 02 to 16**

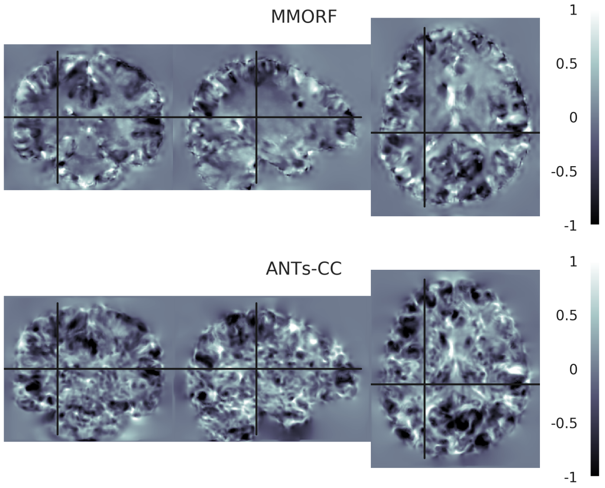


**Log-Jacobian determinant spatial maps - subject 03 to 16**

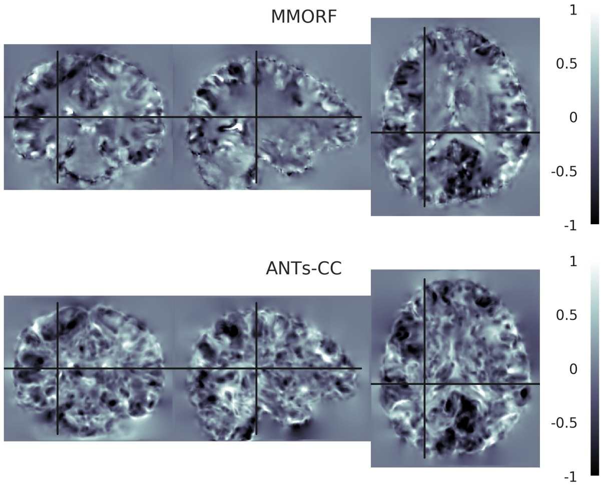


**Log-Jacobian determinant spatial maps - subject 04 to 16**

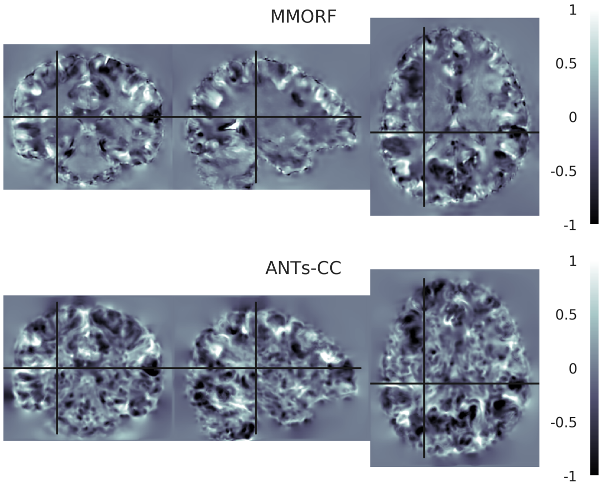


**Log-Jacobian determinant spatial maps - subject 05 to 16**

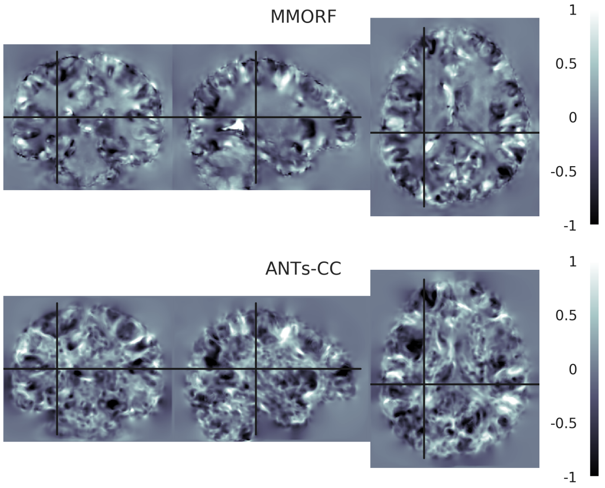


**Log-Jacobian determinant spatial maps - subject 06 to 16**

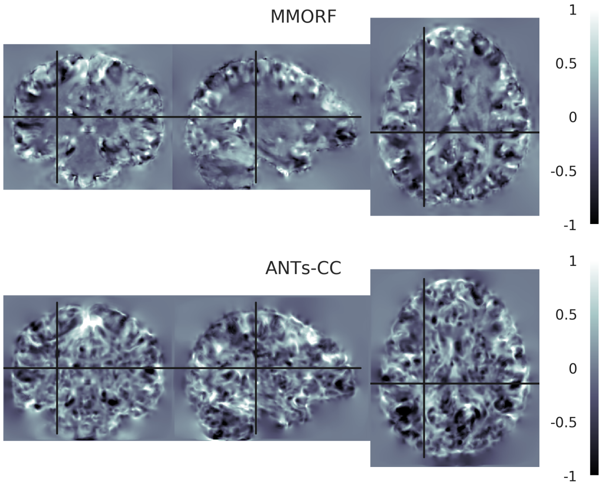

**Log-Jacobian determinant spatial maps - subject 07 to 16**

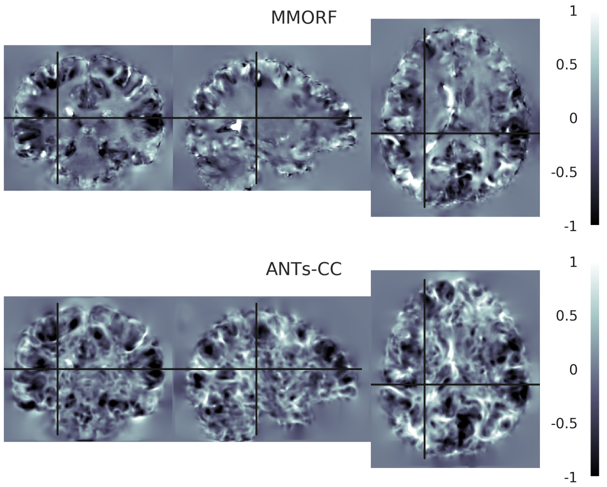


**Log-Jacobian determinant spatial maps - subject 08 to 16**

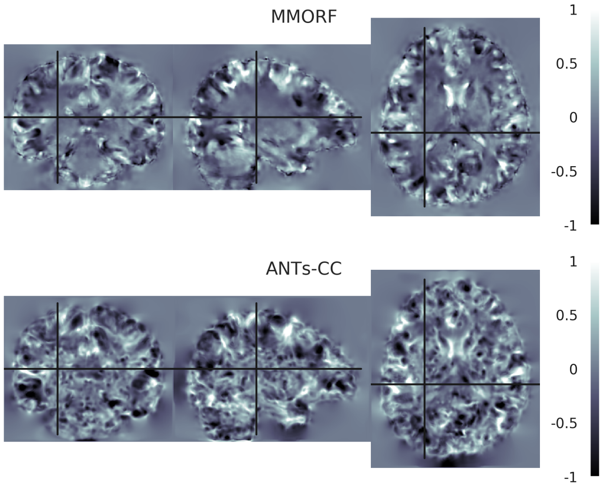


**Log-Jacobian determinant spatial maps - subject 09 to 16**

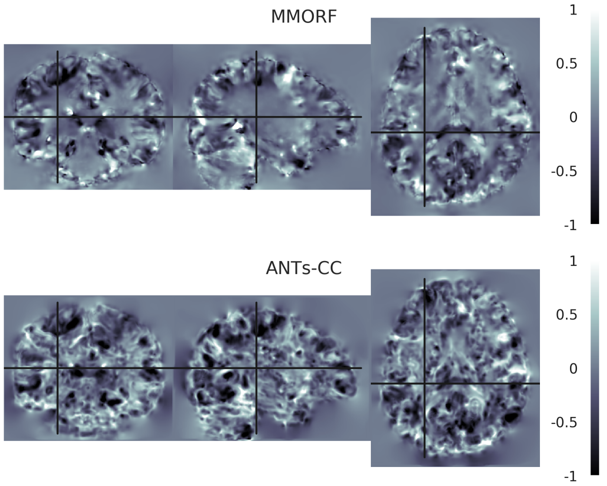


**Log-Jacobian determinant spatial maps - subject 10 to 16**

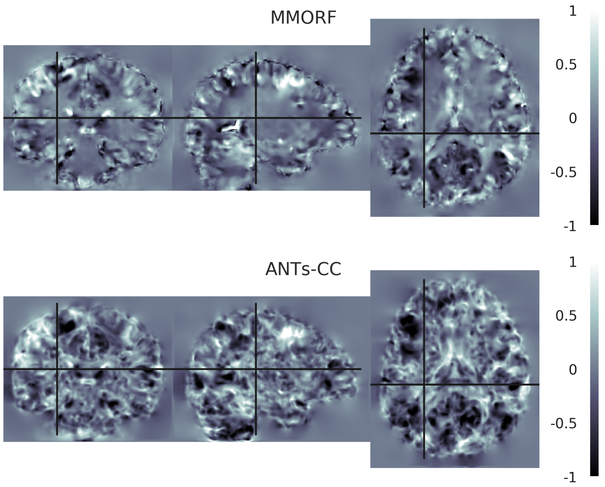


**Log-Jacobian determinant spatial maps - subject 11 to 16**

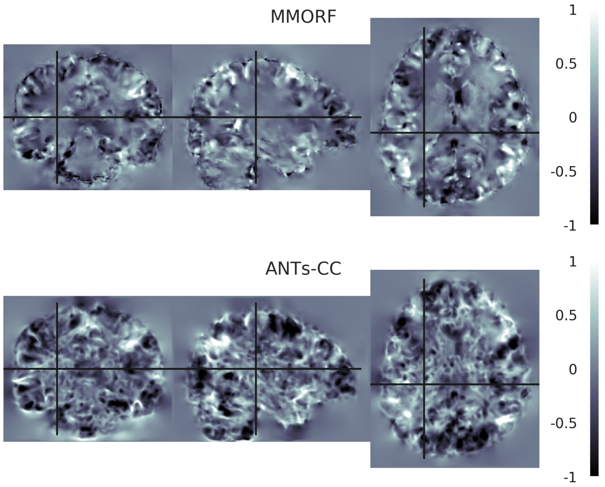


**Log-Jacobian determinant spatial maps - subject 12 to 16**

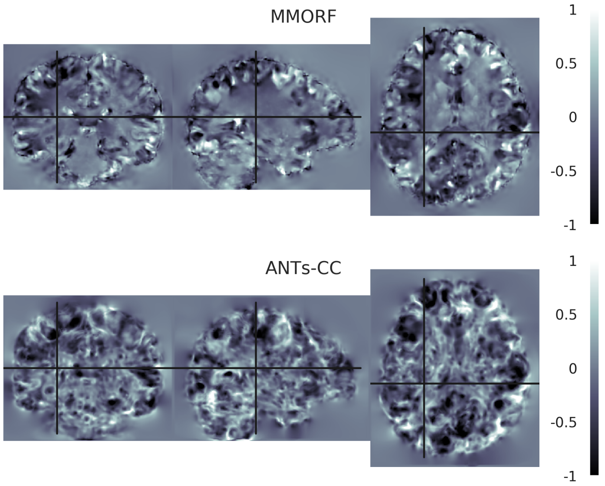

**Log-Jacobian determinant spatial maps - subject 13 to 16**

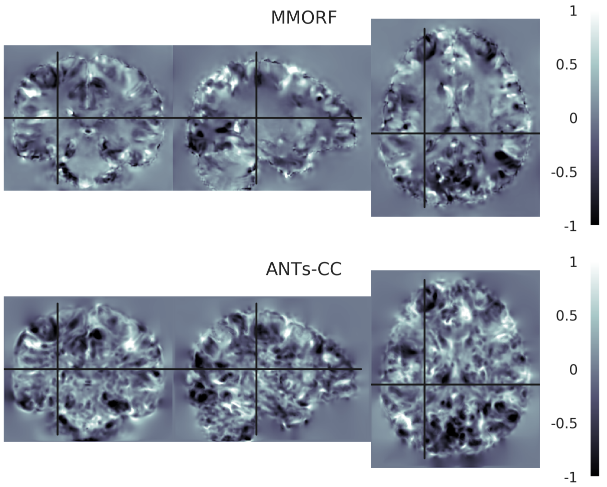


**Log-Jacobian determinant spatial maps - subject 14 to 16**

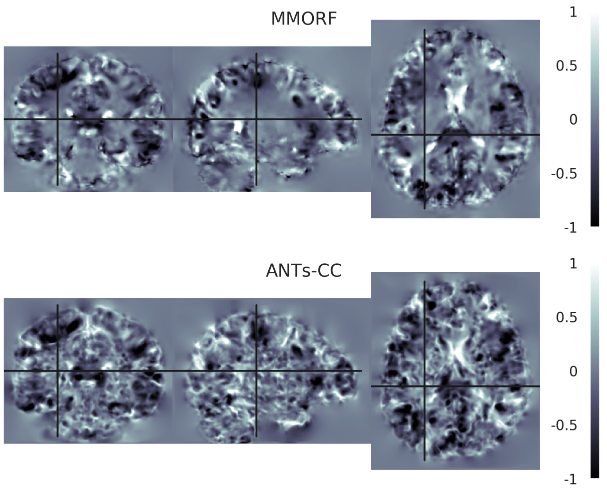


**Log-Jacobian determinant spatial maps - subject 15 to 16**

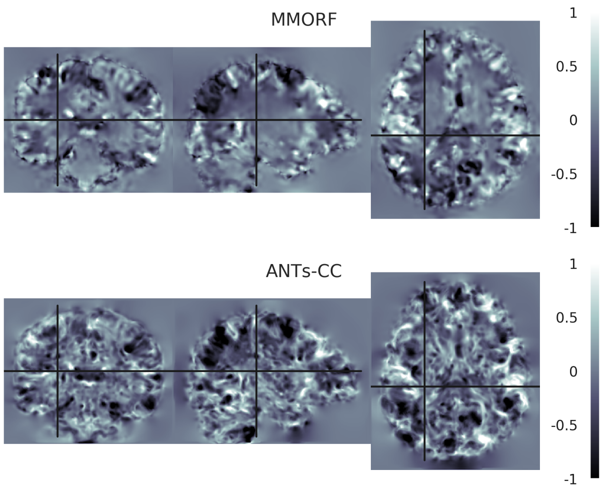
